# Injury Causes Altered Metabolism including O_2_ Consumption in Bovine and Human Chondrocytes

**DOI:** 10.64898/2026.02.13.705805

**Authors:** Aidan J. Gregory, Priyanka P. Brahmachary, Molly E. Piazza, William S. Rockwell, Erik Myers, Mark Greenwood, Ross P. Carlson, Ronald K. June

## Abstract

Traumatic joint injuries both disrupt chondrocyte metabolism and increase the risk for post-traumatic osteoarthritis. Yet the relationships between trauma, altered metabolism, and cartilage degradation remains unclear. This study compares the metabolic responses of bovine (normal) and osteoarthritic (OA) chondrocytes to physiological and injurious mechanical stimuli under normoxic (20% O_2_) and hypoxic (5% O_2_) conditions. Using primary chondrocytes encapsulated in agarose, physiological and injurious mechanical stimulation, targeted metabolomic profiling of central carbon metabolites, and O_2_ saturation measurements, we find that healthy bovine chondrocytes exhibit robust, time-dependent adaptation to mechanical stimuli, whereas OA chondrocytes display a blunted response, particularly under injury conditions. Injurious mechanical stimuli led to altered O_2_ consumption and glutamine accumulation, suggesting disrupted respiration and reduced protein synthesis hypothesized to be a result of altered mitochondrial metabolism in OA cells. These findings underscore the role of mechanical cues in chondrocyte metabolism and inform future studies aimed at identifying metabolic targets relevant to post-traumatic osteoarthritis progression.

## Introduction

Osteoarthritis (OA) is characterized by the progressive degradation of articular cartilage and is the leading global cause of disability, affecting over 600 million people worldwide [1, 2]. Traumatic joint injuries increase risk for the development of osteoarthritis: isolated anterior cruciate ligament (ACL) injuries account for an estimated 12% of all OA cases, while multi-tissue injuries account for 48% of cases [3]. This form of OA, known as post-traumatic osteoarthritis (PTOA), develops from joint trauma, often due to altered joint mechanics and abnormal cellular responses [4]. Despite the high prevalence of PTOA, cellular mechanisms linking injury to cartilage degradation are not fully understood.

Articular cartilage provides a smooth, low friction surface that facilitates joint movement while distributing mechanical loads in the joint [5]. Chondrocytes, the sole cell in articular cartilage, are responsible for synthesizing and maintaining the extracellular and pericellular matrices (ECM and PCM, respectively). Unlike most tissues, articular cartilage is avascular and exists in a hypoxic environment with O_2_ tension below ∼5% (0.05 atm) [6]. Importantly, synthesis of key amino acid precursors for both the ECM and PCM stem from central carbon metabolism, which relies on O_2_ for much of its cellular energy generation. *In vivo*, the ECM and PCM transmit mechanical forces to the chondrocytes. This mechanotransduction influences chondrocyte metabolism and matrix homeostasis. However, excessive mechanical forces, such as those that develop during traumatic joint injury, can shift chondrocyte metabolism toward matrix catabolism, reducing their ability to regulate and maintain cartilage integrity [7, 8].

Central carbon metabolism – which includes glycolysis, the pentose phosphate pathway (PPP), and the tricarboxylic acid (TCA) cycle – synthesizes the precursors to non-essential amino acids. This plays a critical role in maintaining articular cartilage by supplying the precursors needed for synthesis of ECM and PCM molecules. Dysregulation of central metabolism can lead to an imbalance between anabolic and catabolic processing of the ECM and PCM proteins. In OA cartilage, chondrocytes exhibit increased matrix catabolic activity leading to net matrix degradation [9, 10]. Healthy chondrocytes, however, generally maintain matrix homeostasis, which is supported by physiological mechanical stimulation [11]. It is unclear how mechanical injury alters central metabolism and how this is perturbed in OA.

Previous studies have characterized macroscopic and microscopic tissue changes following joint injury [12, 13]. However, the chondrocyte metabolic response, particularly that involving central carbon metabolism, remains poorly defined. This knowledge gap limits our ability to develop targeted interventions to increase metabolic production of non-essential amino acid precursors needed to preserve cartilage health post-injury. Therefore, the objective of this study is to define the metabolic responses of chondrocytes to mechanical injury. We hypothesized that injurious mechanical stimulation would induce a metabolic shift favoring fermentation, characterized by an increase in lactate production and reduced oxygen consumption, whereas physiological stimulation will produce metabolites that maintain matrix homeostasis.

This study uses primary bovine and human OA chondrocytes embedded in high-stiffness agarose. Independent variables include physiological or injurious compression and normoxic or hypoxic tissue culture. Dependent variables include O_2_ consumption and concentrations of central metabolites measured using targeted metabolomics. By identifying metabolic changes associated with an *in vitro* mechanical injury model, this study seeks to provide insight into chondrocyte mechanotransduction to inform potential future therapeutic strategies for PTOA prevention.

## Methods

### Chondrocyte Harvest and Sample Preparation

Primary human osteoarthritic chondrocytes were obtained from discarded arthroplasty tissue from Stage-IV osteoarthritis patients (n=10, 5 female and 5 male, age 65±10 years, mean±standard deviation) undergoing total joint replacement under IRB approval. As a normal control, bovine chondrocytes were harvested from knee joints of 18-22 month old cattle obtained from a local abattoir. For chondrocyte isolation articular cartilage was digested with Type I Collagenase (2 mg/mL, Gibco, Waltham, MA, USA) for 14 h at 37 ºC using a previously established protocol [14]. Isolated chondrocytes were cultured in white Dulbecco’s Modified Eagle’s medium (DMEM, Gibco, Waltham, MA, USA) supplemented with 4.5 g/L glucose, 2 mM L-glutamine,110 mg/L sodium pyruvate (Sigma-Alrich), 10% Fetal Bovine Serum (FBS, Bio-Techne), and 1% PenStrep (Penicillin 10,000 I.U./mL, Streptomycin 10,000 µg/mL, Sigma-Aldrich) in 5% CO_2_ at 37 ºC and first passage cells were used for experiments.

To mimic the PCM in an *in vitro* 3-D model, articular chondrocytes were suspended in 4.5% wt/vol type VIIA low-gelling temperature agarose using previously described methods [12, 15]. A total of 90 gels from n=10 bovine samples, and 90 gels from n=10 ten human OA donors were created. After trypsinization, chondrocytes were washed with 1X PBS and resuspended in DMEM (devoid of Phenol Red) and mixed with low melting temperature agarose Type VII-A (Sigma Aldrich) at a final concentration of 4.5% wt/vol with ∼500,000 cells/gel and allowed to solidify for 10 min. Agarose gels were placed in a 24-well tissue culture plate (Falcon) with 2 mL custom DMEM media containing 4.5 g/L glucose, 2 mM L-glutamine, 110 mg/L sodium pyruvate, 10% FBS, and 1% PenStrep and cultured overnight in 5% CO_2_ at 37ºC.

Because phenol red interferes with mass spectrometry (see below), before mechanical stimulation gels were equilibrated in phenol-red-free (white) DMEM supplemented with 10% FBS, 1% PenStrep, 4.5 g/L glucose, 2 mM glutamine, and 110 mg/L sodium pyruvate for 24 hours under standard tissue culture conditions (normoxic 20% O_2_ or hypoxic 5% O_2_ with 5% CO_2_ at 37°C) [15]. 1 hour prior to mechanical stimulation samples were submerged in 5 mL of Phosphate Buffered Saline (PBS).

### *In vitro* Mechanical Stimulation and Injury

Gels from each donor were divided into 3 mechanical stimulation groups: injury, physiological, and control. A strain energy calculation was performed to determine the necessary loading conditions — the magnitude of compressive strain, strain rate, and number of loading cycles. Grodzinsky *et al* previously established an *in vitro* injury model for explants of articular cartilage by applying a compressive strain of 65% at a strain rate of 4 s^-1^. This induces macroscopic damage to cartilage explants that is consistent with injury [12]. In pilot studies, we observed that agarose samples fracture at absolute strains greater than 30% (a 5% prestrain followed by additional compressive strain beyond 25%). Using a custom-built loading device (Figure 1A), a preload of 5% static compressive strain was applied for 15 minutes [16]. To simulate the Grodzinsky injury model, a total strain of 65% was then applied over multiple cycles: gels were subjected to additional 25% strain for 5 cycles using a triangle waveform at a frequency of 6.6 Hz with a strain rate of 4 s^-1^. For the normal loading control, the same total strain of 65% was achieved using 42 cycles at 5% strain amplitude at a frequency of 1.1 Hz (strain rate of 0.346 s^-1^). The study also included an unloaded control group.

**Figure 1.**
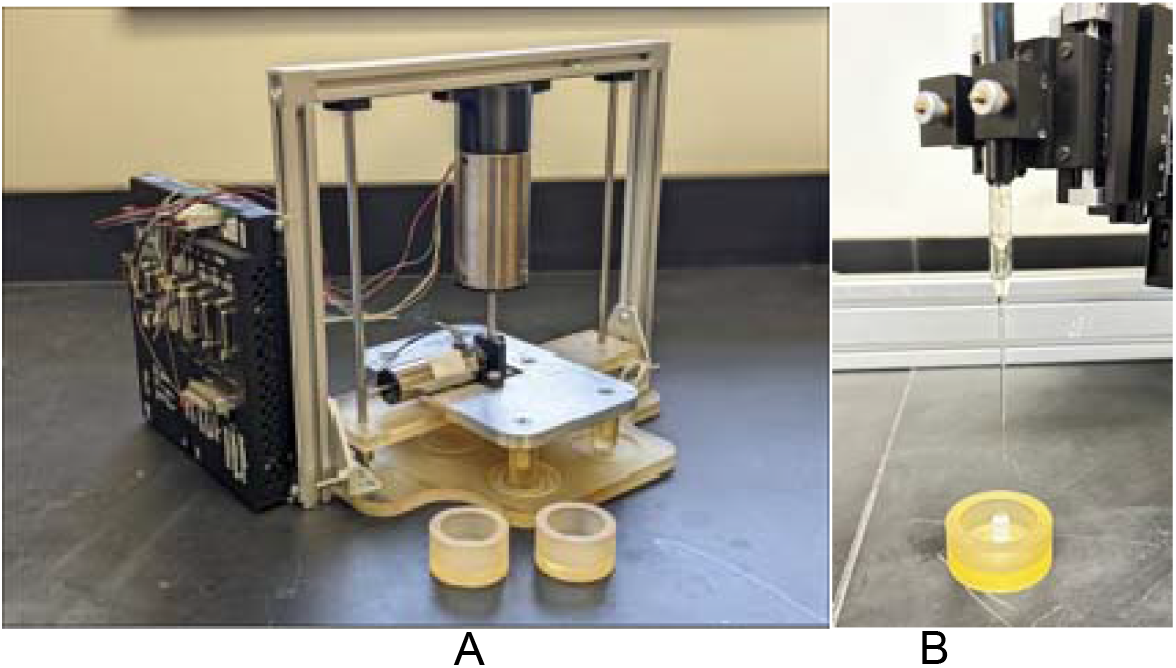
(A) Custom built loading machine (B) Probing a hydrogel with UNISENSE O_2_ probe

### Measurement of O_2_ Consumption

Mitochondria utilize redox carriers generated in the TCA cycle to generate ATP. TCA cycle fluxes are often linked to O_2_ respiration therefore mitochondrial O_2_ consumption was estimated by measuring sample O_2_ saturation[8]. Measurements permitted characterization of differences in respiration magnitude (decrease in soluble O_2_) and overall metabolic activity between groups. A single gel per donor was processed for each timepoint to ensure accurate readings across experimental groups. After loading, hydrogels were bisected. One half of each gel was reserved for metabolomic profiling by flash freezing in liquid nitrogen while the remaining half was immediately probed for O_2_ saturation. To characterize time-dependent metabolic shifts after mechanical stimulation, hydrogels from all experimental groups were reintroduced to their respective culture conditions (*i*.*e*. normoxic or hypoxic) in white DMEM (as described above) for 0, 1, 4, or 24 hours prior to O_2_testing and metabolomic processing.

O_2_ testing was performed using 50 micrometer O_2_ probes (Figure 1B, UNISENSE, Aarhus, Denmark) and logger software. Probe locations were estimated to be at the center of the sample with a tolerance of ±1 mm for both radial and axial directions. After probe insertion, the minimum value of O_2_saturation over a 15 second period was used.

### Metabolite extraction

After mechanical stimulation (or unloaded control), samples were flash frozen and pulverized before storage at –80°C for 12 hours. Samples were then homogenized using a temperature-controlled CryoMill (RETSCH, Haan, Germany) for a total of 12 minutes, and subsequently immersed in 5 mL of 70:30 methanol:acetone with 1% formic acid for 12 hours. All reagents for metabolite extraction were HPLC-grade or higher. Samples and buffer were centrifuged at 4000 rpm for 10 minutes, and supernatants were removed by pipetting. To further purify metabolites, the supernatants were vacuum concentrated at 45°C until only sediment remained. Samples were then stored at –80°C until mass spectrometry [15].

### Metabolomic Profiling

To investigate how mechanical stimulation impacts central carbon metabolism, a targeted panel of key central metabolites was developed (Supplemental Table 1, {Myers, 2025 #23}). These included glucose, pyruvate, lactate, oxaloacetate, and glutamine. Glucose, the primary substrate for glycolysis, and pyruvate, a critical glycolytic product, were quantified to assess glycolytic activity. Pyruvate serves as a metabolic branching point, converting to either lactic acid or acetyl-CoA, which feeds the TCA cycle. The TCA cycle can produce oxaloacetate, a precursor for lysine, which contributes to synthesis of collagen, a key constituent of both the PCM and ECM. Additionally, glutamine and proline are precursors for collagen. Collagens (type II and VI) are critical for the structural integrity of the ECM and PCM, highlighting the importance of these metabolites in supporting cartilage function under mechanical stimuli.

To quantify metabolomic changes, a targeted metabolomics approach was employed. Calibration standards were prepared using each of the central metabolites of interest at concentrations of 1, 2, 10, 50, and 100 µg/µL suspended in 50 mL of 50:50 acetonitrile:HPLC water. Experimental samples were suspended in 10 µL of the same solvent and vortexed for one minute [15].

Metabolite analysis was performed using high-performance liquid chromatography-mass spectrometry (HPLC-MS) operating in positive ionization mode (Waters, Synapt, C18 column). The instrument operated at a resolution of 20 ppm, maintaining a mass accuracy of 5 ppm. Metabolites were identified and separated based on the corresponding m/z value and retention time through Agilent Mass Hunter Quantitative software. Quantification was achieved by comparing sample metabolite intensities to calibration curves from the prepared standards.

### Statistical Analysis of O_2_ and Metabolic Data

To reduce potential biases and ensure robust experimental design, samples from each donor were randomly assigned to each loading group (control, injury, and physiological) and across all timepoints (0, 1, 4, and 24 hours). Statistical analyses were performed using GraphPad Prism (v10) for O_2_ data and R {Team, 2025 #24} for metabolite data. O_2_ data were analyzed using either one-way (factor levels of mechanical stimulus within either human OA or bovine samples) or two-way (factors of mechanical stimulus and cell types, either human OA or bovine) ANOVA with intragroup comparisons using Tukey’s correction for multiple comparisons based on final models.

All metabolite concentration data were normalized to the total ion intensity (TII) to correct for variations in extraction and instrument performance. Metabolites were then classified into three categories based on detection frequency (<33% (3 bovine and 6 human metabolites), between 33 – 80% (5 bovine and 6 human metabolites), and >80% (4 bovine and 0 human metabolites)). If a metabolite was detected in less than 1/3 of samples the metabolite was excluded from all statistical analysis. For metabolites detected in 33 to 80% of all samples, logistic regression was used with a dependent variable of detection rate, but subject (either donor or bovine sample) was removed because the high number of non-detects lead to covariate patterns that perfectly predicted the response. Factors were loading type (injurious or physiological) and time (1, 4, and 24 hours). A second logistic regression model was run separately for the 0-hour timepoint, (*i*.*e*., samples analyzed immediately after loading) that included only the factor of loading type (injurious, physiological, or unloaded control).

If the metabolite was detected in over 80% of all samples, linear models were created, including factors for loading type, time, and donor, as well as their interactions. A stepdown model simplification approach was applied: the factor with the greatest nonsignificant p-value was removed, starting with any interactions, and then the model was re-fit. A separate linear model was created at time point 0, immediately after loading, including loading type and donor as factors. The same stepdown approach was used to simplify the model. Tukey’s multiple correction was used to compare levels of factors that were retained in the final models. Type-II tests were used for both metabolite and O_2_ models.

To compare relationships between O_2_ saturation and metabolite levels, differences were calculated by subtracting the baseline control levels from experimental values on a per-donor basis for all loading conditions and timepoints. This analysis was only performed for metabolites with detection rates above 33% using logistic regression with increases defined as differences above 0.1 μg/μL/TII and negligible increases defined as differences below 0.1 μg/μL/TII. Logistic regression models based on the change in O_2_ saturation (often modeled with a quadratic polynomial due to observed curved patterns) and its interaction with hypoxia/normoxia were considered, along with donor if non-detect rates were low using the same stepdown approach as above.

## Results

### O_2_ Saturation in Normoxic Culture

O_2_ is required for mitochondrial respiration to process TCA cycle products therefore, we measured O_2_ saturation levels within the hydrogels using microelectrodes. At baseline and after both normal (e.g. physiological) and injurious mechanical stimulation, there were no detectable differences in O_2_ saturation between hydrogels from unloaded control bovine samples and OA human donors (Figures 2A-B).

**Figure 2.**
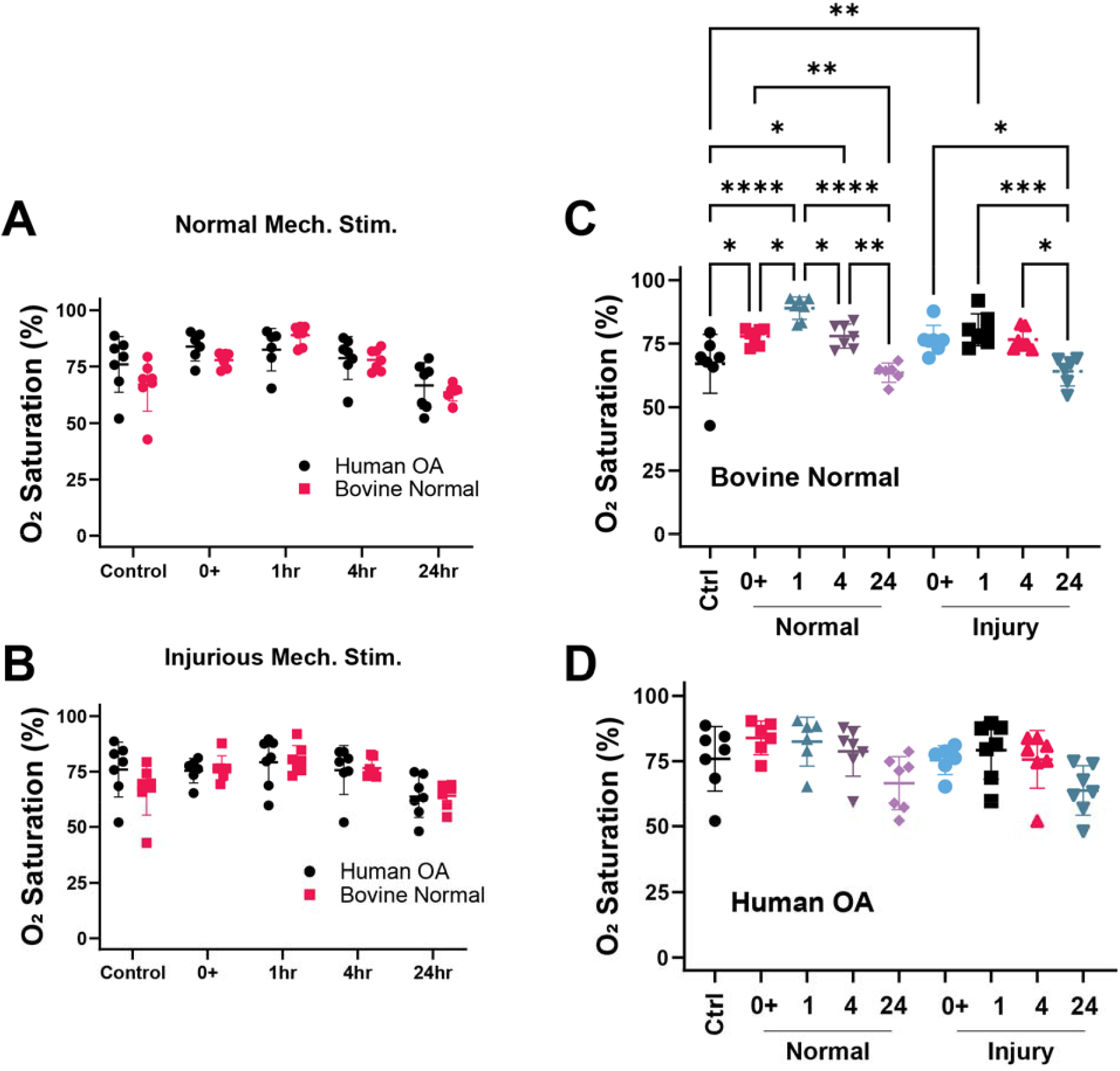
O_2_ saturation (%) levels of chondrocytes after normal and injurious mechanical stimulation in normoxic O_2_ levels with means (wide line) and 95% confidence intervals (error bars). (A) Comparison of bovine and human OA chondrocytes after normal mechanical stimulation. (B) Comparison of bovine and human OA chondrocytes after injurious mechanical stimulation. (C) Response of bovine chondrocytes after both normal and injurious mechanical stimulation. (D) Response of human OA chondrocytes after both normal and injurious mechanical stimulation. Data in panels A and B analyzed by 2-way ANOVA comparing the means of each group. Data in panels C and D analyzed using 1-way ANOVA. Statistical significance is denoted by the following code for Tukey’s adjusted p-values: p < 0.05 = *, p < 0.01 = **, p < 0.001= ***.

Hydrogels from bovine samples (n=6-10) showed a clear O_2_ saturation response to both physiological and injurious mechanical stimuli (Figure 2C). O_2_ saturation increased in both loading conditions at 1 hour post loading compared to control suggesting reduced metabolic activity then decreased at later timepoints when compared to 1 hour (all tests for differences had p < 0.01) suggesting time dependent changes in metabolic activity for normal bovine chondrocytes. In contrast, hydrogels from human OA chondrocytes did not show changes in O_2_ saturation in response to physiological or injurious mechanical stimuli over time suggesting no metabolic response to these stimuli in normoxic conditions (Figure 2D).

### O_2_ Saturation in Hypoxic Culture

Similar to normoxic conditions, hypoxic culture resulted in no baseline differences in O_2_ saturation between bovine normal and human OA chondrocytes (Figure 3A). For samples subjected to both normal and injurious mechanical stimulation, there were no differences in O_2_ saturation between normal bovine and human OA donors at any time point (Figure 3A-B).

**Figure 3.**
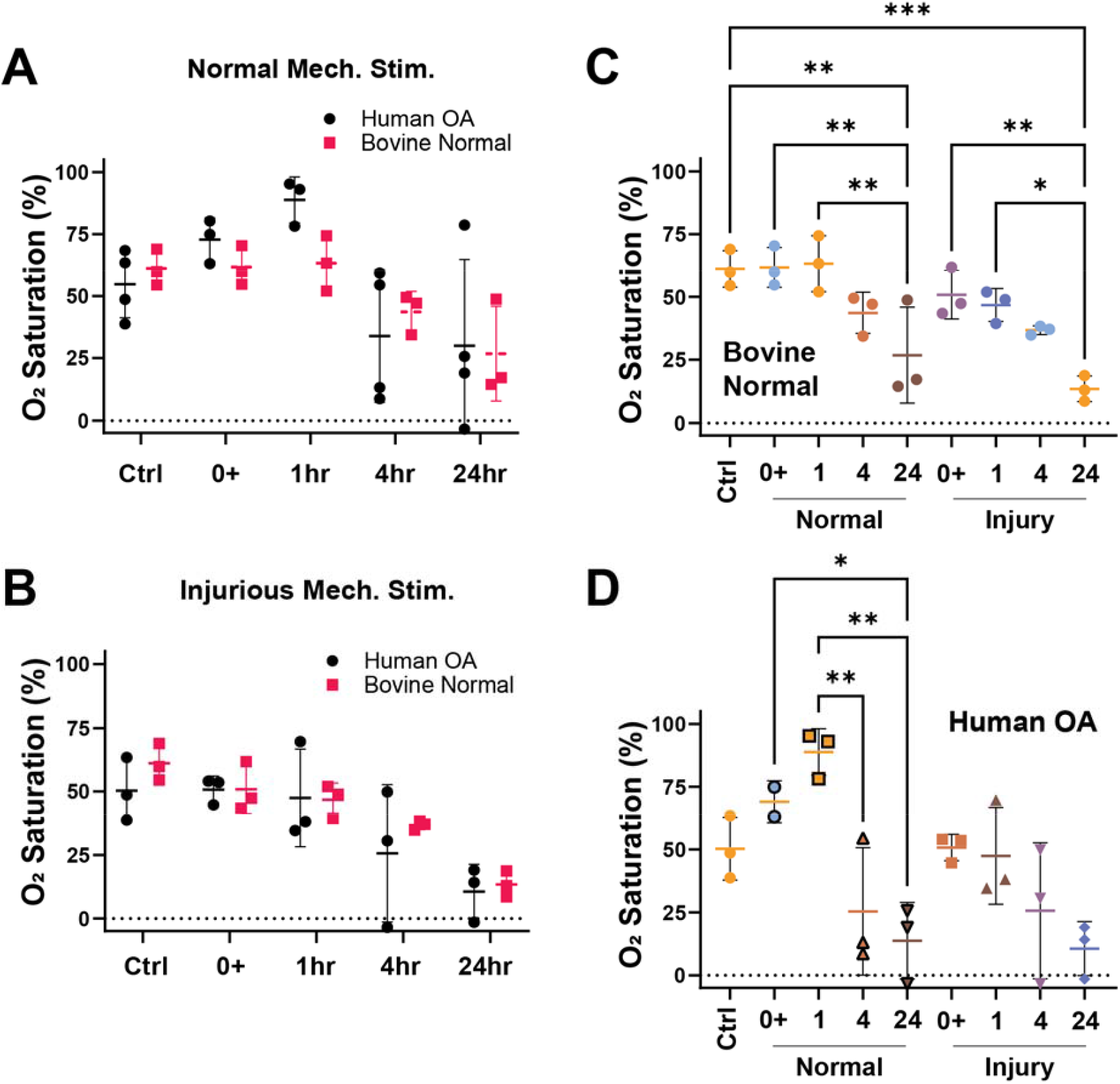
O_2_ saturation (%) levels of chondrocytes after normal and injurious mechanical stimulation in hypoxic oxygen levels with means (wide line) and 95% confidence intervals (error bars).. (A) Comparison of bovine and human OA chondrocytes after normal mechanical stimulation. (B) Comparison of bovine and human OA chondrocytes after injurious mechanical stimulation. (C) Response of bovine chondrocytes after both normal and injurious mechanical stimulation. (D) Response of human OA chondrocytes after both normal and injurious mechanical stimulation. Panels A and B compare the means of each group using 2-way ANOVA. Panels C and D compare the means of each group using 1-way ANOVA. Statistical significance is denoted by the following code for Tukey’s adjusted p-values: p < 0.05 = *, p < 0.01 = **, p < 0.001= ***.

Hydrogels from bovine samples subjected to normal mechanical compression showed a significant decrease in O_2_ saturation at 24 hours when compared to unloaded controls indicating an increase in metabolic activity with stimulation (p<0.01, Figure 3C). Bovine O_2_ saturation at 24-hours was also significantly less than that at 0+-, 1-, and 4-hour timepoints after normal mechanical stimulation (all p<0.01, Figure 3C). After 24 hours, bovine samples subjected to injurious compression showed a significant decrease in O_2_ saturation compared to both unloaded controls (p<0.01) and 1-hour post-loading (p<0.05, Figure 3C) suggesting the cells were mounting a vigorous metabolic response to the insult.

O_2_ saturation for hydrogels from human OA donors was affected by normal mechanical stimulation. O_2_ levels were lower after 24 hours of mechanical stimulation compared to 0 (p<0.05) and 1 hour (p<0.01, Figure 3D) timepoints. Furthermore, O_2_ levels after 4 hours were significantly lower than after 1 hour (p<0.01, Figure 3D). In contrast to normal chondrocytes from bovine samples, there were no detectable differences in samples from human OA donors subjected to injurious stimulation when compared across timepoints.

### Central Metabolite Response to Injurious Loading in Normoxia and Hypoxia

Hydrogels from bovine donors (n=10) showed a greater than 50% detection rate in the 6 metabolites of interest (Supplemental Figure 1). Hydrogels from OA donors (n = 10) showed a greater the 50% detection rate in only 3 metabolites (Supplemental Figure 1). Full statistical results available in Supplemental File.

Healthy bovine samples subjected to injury had lower levels of glucose compared to normal loading (p = 0.029, Figure 4A), while there were no detectable differences in glucose levels for human OA samples (Figure 4B). The glucose data is consistent with the O_2_ data suggesting the healthy cells mount a robust metabolic response to mechanical stimulation while the metabolism of the OA cells is largely unresponsive to mechanical stimulation. Furthermore, there was greater probability of increased glutamine in normoxia than hypoxia when accounting for donor and O_2_ saturation differences (p = 0.0100, Figure 5). Finally, in human OA samples there was a higher detection rate of fructose-6-phosphate after normal mechanical stimulation compared to injurious (p=0.0302, Supplemental Figure 2).

**Figure 4.**
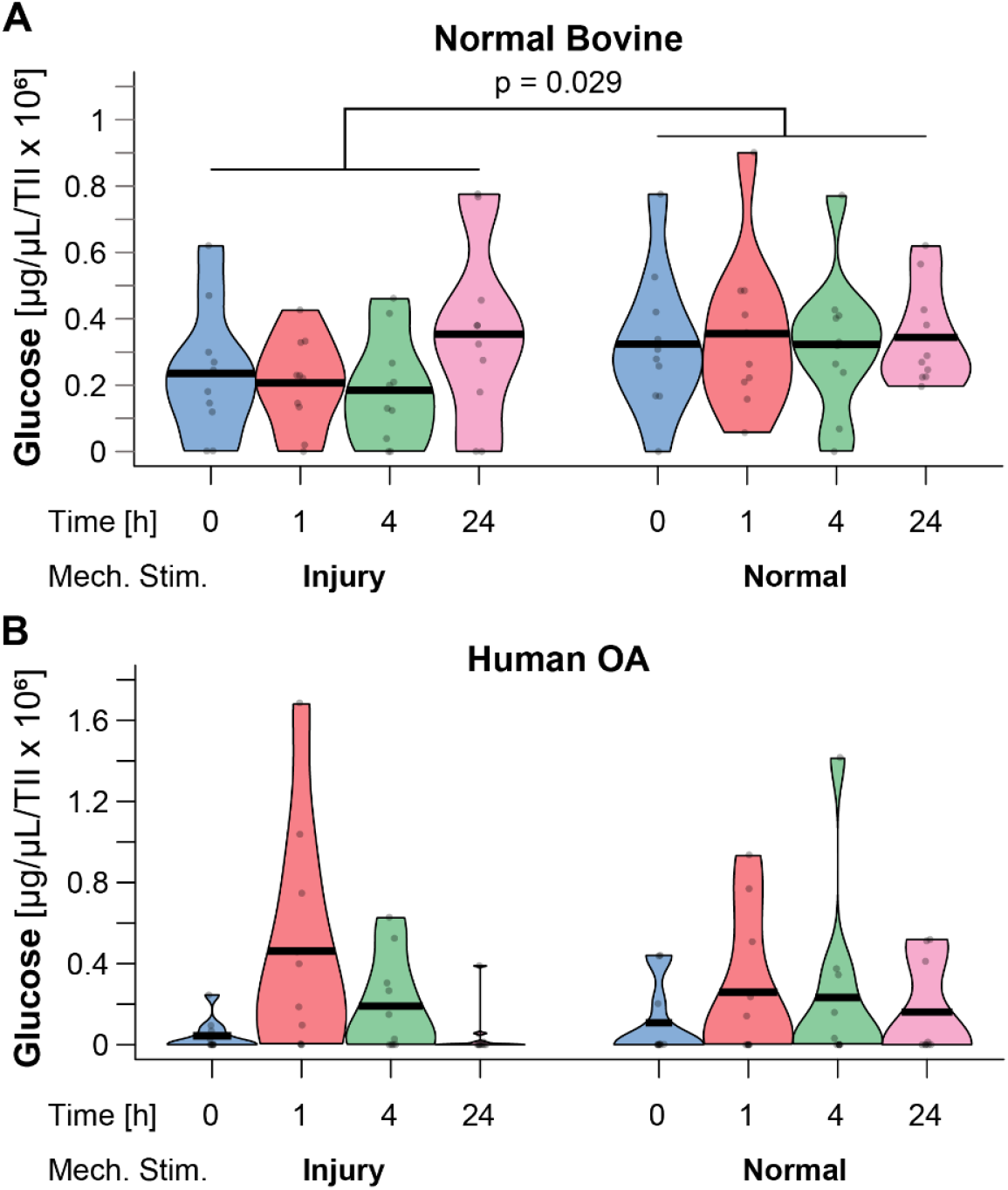
Glucose abundance TIC corrected. (A) Bovine (B) Human OA. Bovine glucose was lower under injurious mechanical stimulation than normal p = 0.029.

**Figure 5.**
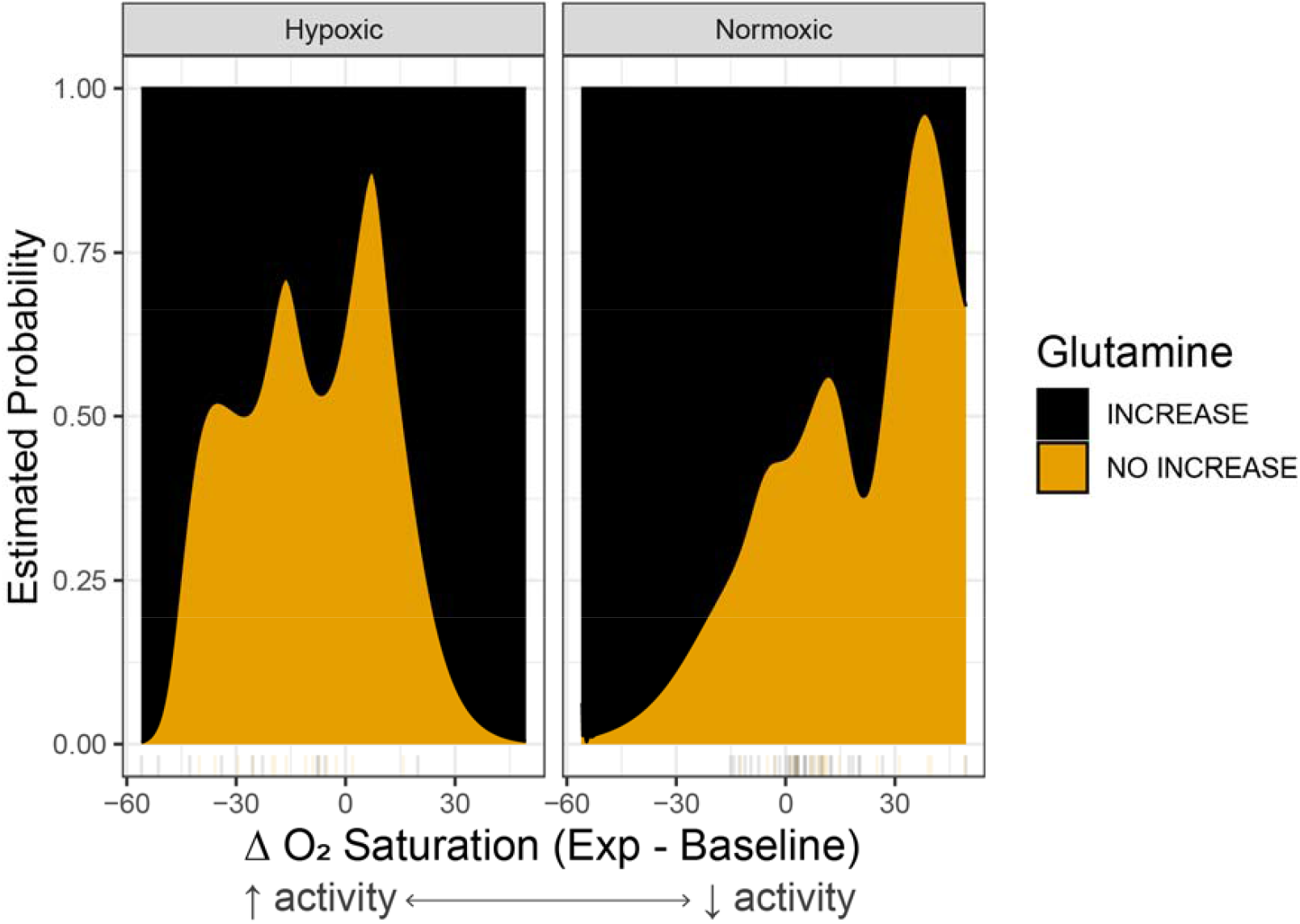
Estimated probability of glutamine increase as a function of change in O_2_ saturation by condition. The final logistic regression model suggests higher probability of increased glutamine in normoxia than hypoxia when accounting for O_2_ saturation differences (two-sided p = 0.010). After dropping the interaction, there is strong evidence of a difference based on hypoxia/normoxia, Chi-squared(1) =6.63, p = 0.01, accounting for change in O_2_ saturation, with the estimated probability of glutamine increase higher in the normoxia group than in hypoxia.

## Discussion

This study aims to evaluate how central metabolism of normal and OA chondrocytes responds to normal and injurious mechanical stimulation. These experiments integrate mechanical compression with metabolic readouts, including O_2_ levels which can be correlated to respiration activity and metabolite profiles. By comparing normal chondrocytes with OA chondrocytes in both normoxia and hypoxia, this study provides insight into how chondrocytes respond to loading in both healthy and diseased states. Both strain magnitude and strain rate were designed to model physiologically relevant (e.g. normal) and injurious conditions *in vitro*. By characterizing the metabolic state of chondrocytes through quantitative metabolomics and O_2_ consumption data, this paper shows that normal chondrocytes display metabolic plasticity in response to different loading conditions that is diminished in OA cells. These data provide new insight into how the metabolism of OA chondrocytes is affected by normal and injurious loading and may have key implications for understanding OA progression.

### Injury Alters O_2_ Consumption of OA and Bovine Chondrocytes

This study shows metabolic distinctions between OA and normal chondrocytes subjected to injurious and normal levels of mechanical stimulation. Targeted metabolomics and O_2_ consumption data found that OA chondrocytes exhibit a minimal metabolic response to injury compared to normal bovine chondrocytes. Normal bovine chondrocytes showed robust and time-dependent metabolic alterations under both injurious and normal loading likely associated with healthy healing mechanisms. Notably, in normoxic conditions, bovine chondrocytes responded metabolically as early as one hour post loading, with sustained activity across timepoints (Figure 2C), consistent with a substantial metabolic capacity for mechanotransduction [17].

In contrast, OA chondrocytes displayed minimal changes in O_2_ saturation and associated metabolic activity under normoxic injury conditions (Figure 3C), indicating a potential mechanotransduction impairment and/or mitochondrial alterations [18]. This suggests caution in using normoxia for culture of articular chondrocytes. However, under hypoxic conditions – more reflective of the *in vivo* chondrocyte environment – human OA cells showed decreased O_2_ consumption as early as 4 hours post loading (Figure 3D) whereas bovine cells did not exhibit a metabolic response until 24 hours (Figure 3C). This suggests that human OA chondrocytes shift toward fermentation and lactate production, a hallmark of OA cell metabolism [19].

Furthermore, the suppressed response of OA chondrocytes to mechanical injury may reflect either limited mitochondrial energy reserves and/or a metabolic phenotype incapable of responding to mechanical queues, as supported by previous findings in OA cartilage [20]. Together these data highlight the importance of O_2_ metabolism as a holistic functional readout of mechanotransduction and suggest that therapeutic targeting of mitochondrial metabolism could restore chondrocyte responsiveness in OA.

### Relationship between environmental O_2_and glutamine

The glutamine level probability curves differed between normoxia and hypoxia suggesting a different metabolic regulation as a function of O_2_. Glutamine serves as an important metabolic substrate as well as a key intermediate in the synthesis of many amino acids and therefore protein synthesis [21]. The glutamine differences between normoxia and hypoxia are not surprising given the importance of O_2_ for cellular energy generation and TCA cycle fluxes. The data highlight challenges of interpreting data from *in vitro* cultures that are not grown under appropriate O_2_ tension. Glutamine levels can also increase under stress in highly proliferating cells [23]. However, chondrocytes in agarose are not highly proliferating suggesting that the observed increase in probability of glutamine increase in normoxia may be either a stress response or to promote collagen synthesis. There were no detected differences in glutamine levels between any mechanical stimuli in this study.

### Study limitations

This study has important limitations. First, although bovine chondrocytes provide a consistent source of non-OA chondrocytes, they do not fully capture the behavior of human chondrocytes and are an imperfect normal control. Species-specific differences may influence the observed bovine responses to mechanical stimuli and hypoxia. Second, the *in vitro* injury model, while enabling controlled application of mechanical loads, does not fully represent the complex physiological environment of the joint, which includes synovial fluid and immune response. Because of the mechanical properties of agarose, the injurious load was applied over several cycles. While the use of the normal mechanical stimuli also applied the same total strain energy, it is possible that the injury protocol may miss important aspects of the *in vitro* injury response to. Finally, the interpretations of metabolite levels assume pseudo steady-state flux, which may not be accurate in the context of acute cellular stress or injury.

## Supporting information

Supplemental Statistics

## Acknowledgements

This study was supported by research grants from the NIH (NIAMS R01AR073964 and R01AR081489) and NSF (CMMI 1554708).

## Supplemental Material

- Supplemental Table 1: O_2_ Saturation Data.
- Supplementary Table 2: Metabolomic Profiling of Central Metabolites
- Supplemental Methods: Statistical Analysis Code

**Supplemental Figure 1.**
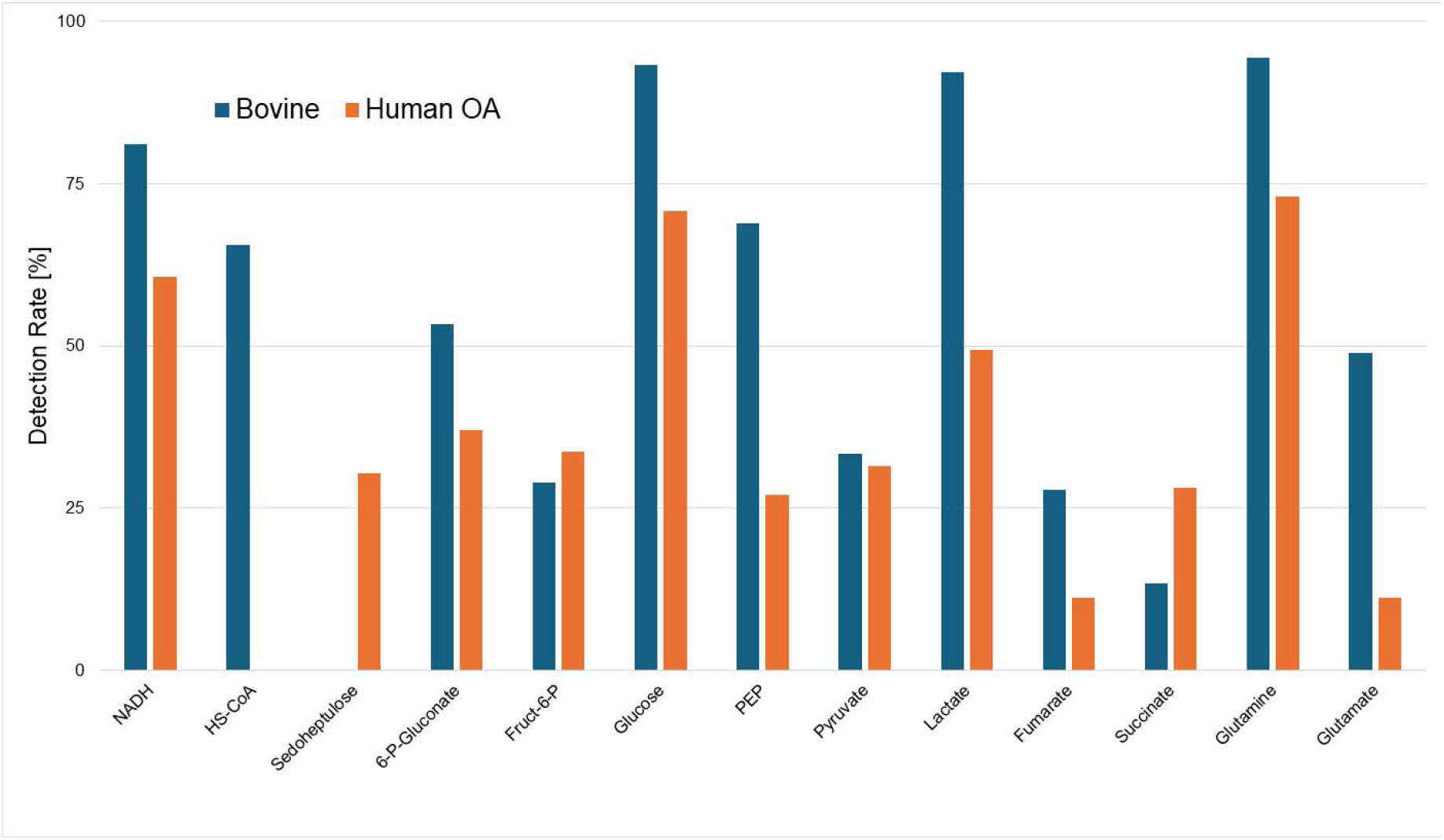
Detection rates for central metabolites in bovine and human OA samples.

**Supplemental Figure 2.**
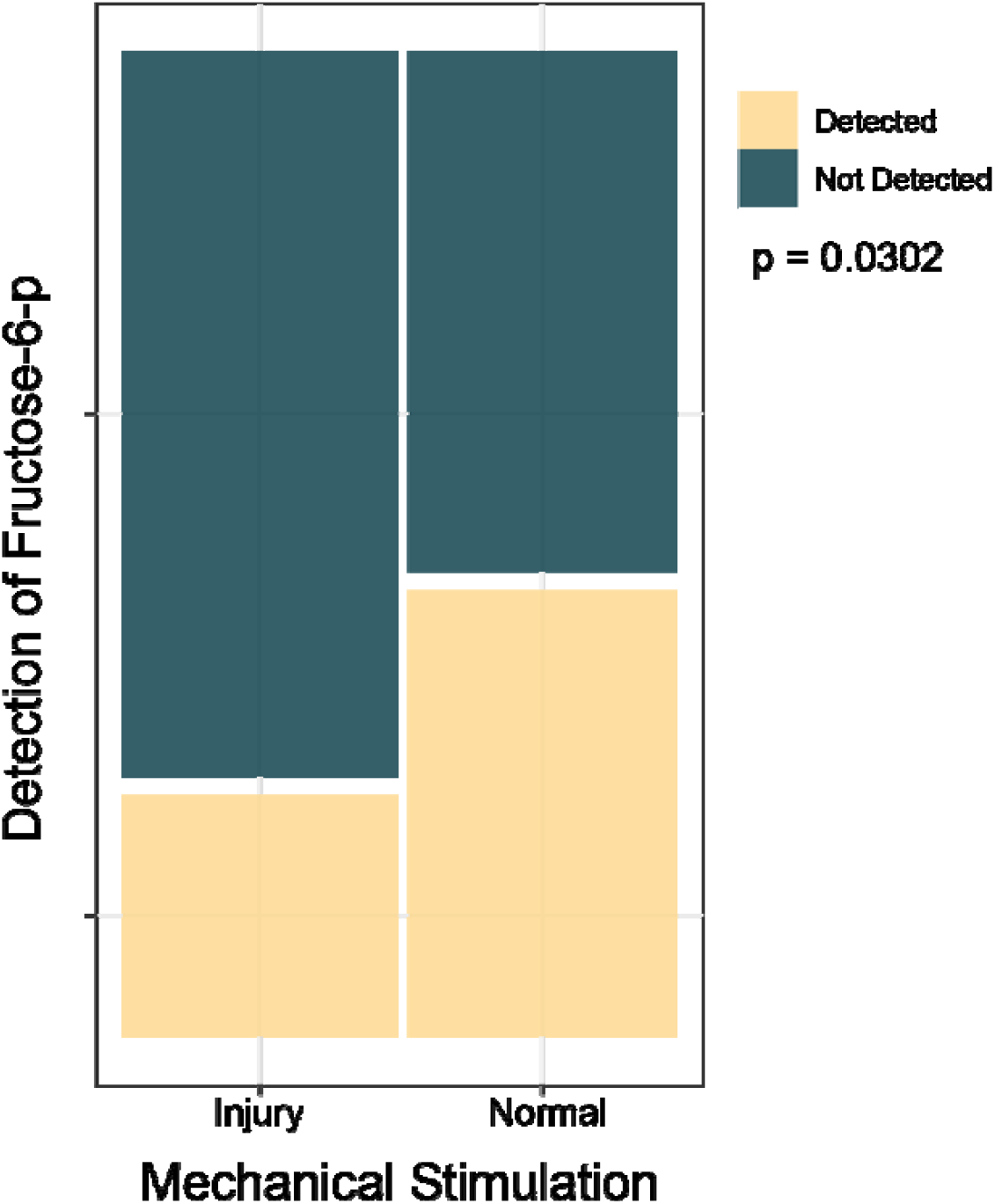
Increased detection of Fructose-6-phosphate after normal mechanical stimulation compared to injurious mechanical stimulation in human OA chondrocytes.

