## Supplemental Statistics for "Injury Causes Altered Metabolism including O_2_ Consumption in Bovine and Human Chondrocytes"

### Statistical Analysis

OA Metabolites vs time and loading

```
library(readxl)
TICsetOA <- read_excel("Injury_ML_HeatMap_OA_withTIC.xlsx")
dim(TICsetOA)

## [1] 90 50

tally(Time ~ Label, data = TICsetOA)

##       Label
## Time    C  I  N <NA>
##   0    10 10 10    0
##   1     0 10 10    0
##   4     0 10 10    0
##   24    0 10  9    0
##   <NA>  0  0  0    1

TICsetOA <- TICsetOA %>% mutate(TimeF = factor(Time),
                                Label.Time = str_c(Label, TimeF))
```

Analysis for metabolite abundance compared to loading condition and time is split into two types of analysis. The first is at all time points for the injury and normal loading and the second is only at time 0 for control, injury and normal. The NA is a sample that underwent normal loading conditions and at the 24 hour time point but was not able to be run on the mass spectrometer.

Fructose-6-Phosphate

The original full logistic regression model includes an interaction between time and loading as well as the donor. In this model a “success” is defined by the metabolite being detected in the sample. The model was simplified to just include time and loading without an interaction as both donor and the interaction showed little to no evidence of an impact on metabolite abundance.

```
TICsetOAR <- TICsetOA %>% dplyr::filter(Label != "C") %>%
  mutate(Fructose6B = factor(Fructose6B),
         Donor = factor(Donor),
         TimeF = factor(Time))

#pdf("OA_Fructose_Loading.pdf",width=5, height=5)
Fru<- TICsetOAR %>%
  ggplot() + scale_fill_paletteer_d("nationalparkcolors::Acadia")  +
  geom_mosaic(aes(x=product(Label),fill=Fructose6B),offset=0.02)

Fru
```

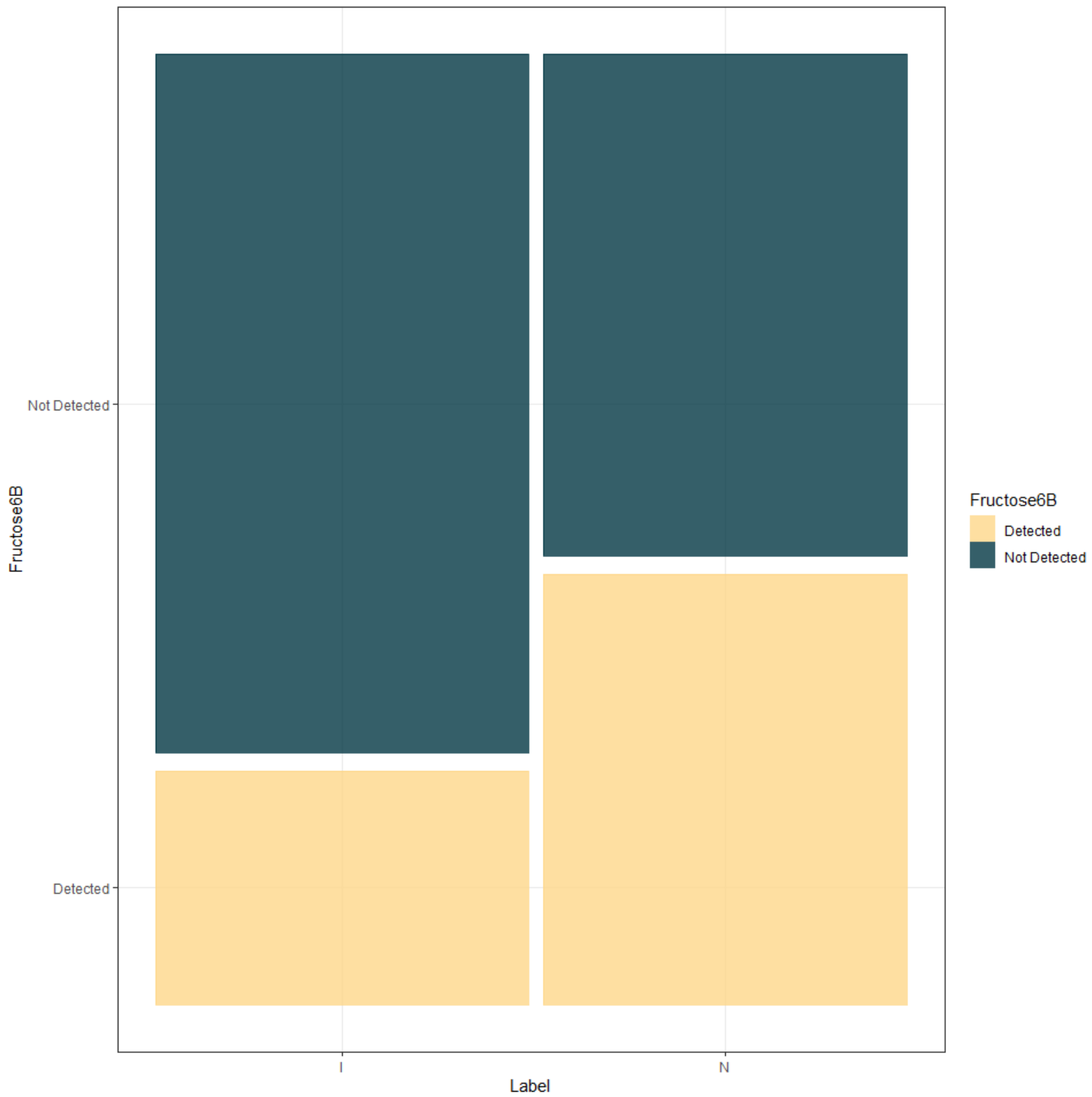


```
TICsetOAR <- TICsetOA %>% dplyr::filter(Label != "C") %>%
  mutate(Fructose6B = factor(Fructose6B),
         Donor = factor(Donor))

glm1 <- glm(relevel(Fructose6B,"Not Detected") ~ Label*TimeF+Donor,
data=TICsetOAR, family = "binomial")
Anova(glm1, type="III")

## Analysis of Deviance Table (Type III tests)
##
## Response: relevel(Fructose6B, "Not Detected")
##             LR Chisq Df Pr(>Chisq)
## Label         0.0000  1     1.0000
## TimeF         3.2933  3     0.3486
## Donor        12.4501  9     0.1891
## Label:TimeF   5.1968  3     0.1579
```

```r
summary(glm1)

##
## Call:
## glm(formula = relevel(Fructose6B, "Not Detected") ~ Label * TimeF +
##     Donor, family = "binomial", data = TICsetOAR)
##
## Coefficients:
##                  Estimate Std. Error z value Pr(>|z|)
## (Intercept)    -2.228e+00  1.258e+00  -1.771   0.0765
## LabelN         -2.100e-15  1.208e+00   0.000   1.0000
## TimeF1          1.188e+00  1.124e+00   1.056   0.2909
## TimeF4          6.436e-01  1.147e+00   0.561   0.5746
## TimeF24        -9.148e-01  1.396e+00  -0.655   0.5122
## Donor2          7.646e-01  1.250e+00   0.612   0.5408
## Donor3          7.646e-01  1.250e+00   0.612   0.5408
## Donor4         -1.035e+00  1.481e+00  -0.698   0.4849
## Donor5          1.432e+00  1.237e+00   1.157   0.2472
## Donor6         -5.609e-01  1.517e+00  -0.370   0.7117
## Donor7          2.081e+00  1.256e+00   1.657   0.0974
## Donor8         -7.733e-16  1.305e+00   0.000   1.0000
## Donor9         -1.729e-15  1.305e+00   0.000   1.0000
## Donor10         2.081e+00  1.256e+00   1.657   0.0974
## LabelN:TimeF1   1.533e+00  1.603e+00   0.956   0.3390
## LabelN:TimeF4   9.392e-16  1.619e+00   0.000   1.0000
## LabelN:TimeF24  3.312e+00  1.837e+00   1.803   0.0713
##
## (Dispersion parameter for binomial family taken to be 1)
##
##     Null deviance: 102.723  on 78  degrees of freedom
## Residual deviance:  75.637  on 62  degrees of freedom
## AIC: 109.64
##
## Number of Fisher Scoring iterations: 5

plot(allEffects(glm1), type = "response", grid = T)
```

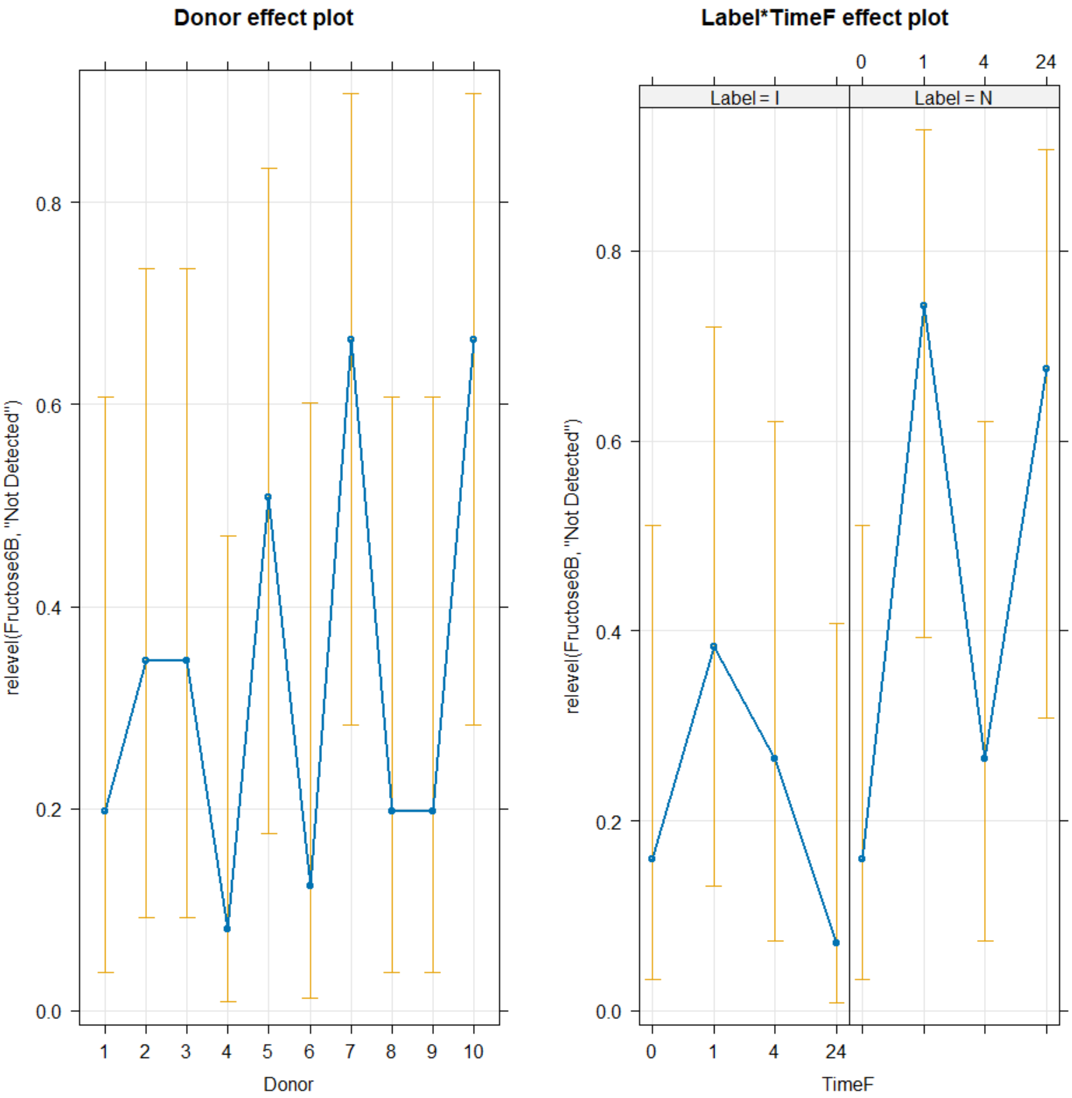


```
glm2 <- glm(relevel(Fructose6B,"Not Detected") ~ Label+TimeF+Donor,
data=TICsetOAR, family = "binomial")
Anova(glm2)

## Analysis of Deviance Table (Type II tests)
##
## Response: relevel(Fructose6B, "Not Detected")
##       LR Chisq Df Pr(>Chisq)
## Label   4.7001  1    0.03016
## TimeF   7.0747  3    0.06956
## Donor  11.9145  9    0.21817

#Can't add donor - causes separation here

summary(glm2)
```

```
##
## Call:
## glm(formula = relevel(Fructose6B, "Not Detected") ~ Label + TimeF +
##     Donor, family = "binomial", data = TICsetOAR)
##
## Coefficients:
##               Estimate Std. Error z value Pr(>|z|)
## (Intercept) -2.797e+00  1.139e+00  -2.455   0.0141
## LabelN       1.181e+00  5.626e-01   2.099   0.0358
## TimeF1       1.987e+00  8.209e-01   2.421   0.0155
## TimeF4       6.592e-01  8.218e-01   0.802   0.4224
## TimeF24      1.003e+00  8.201e-01   1.223   0.2213
## Donor2       6.915e-01  1.188e+00   0.582   0.5606
## Donor3       6.915e-01  1.188e+00   0.582   0.5606
## Donor4      -9.589e-01  1.428e+00  -0.672   0.5018
## Donor5       1.301e+00  1.177e+00   1.105   0.2691
## Donor6      -7.139e-01  1.455e+00  -0.491   0.6237
## Donor7       1.909e+00  1.198e+00   1.593   0.1112
## Donor8       6.346e-16  1.244e+00   0.000   1.0000
## Donor9       2.361e-16  1.244e+00   0.000   1.0000
## Donor10      1.909e+00  1.198e+00   1.593   0.1112
##
## (Dispersion parameter for binomial family taken to be 1)
##
##     Null deviance: 102.723  on 78  degrees of freedom
## Residual deviance:  80.834  on 65  degrees of freedom
## AIC: 108.83
##
## Number of Fisher Scoring iterations: 4

plot(allEffects(glm2), type = "response", grid = T)
```

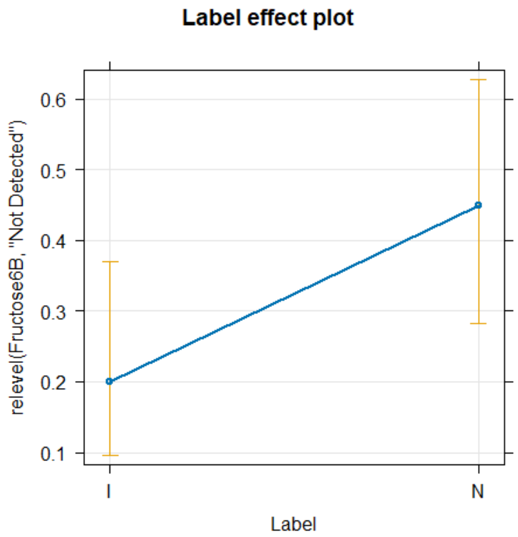


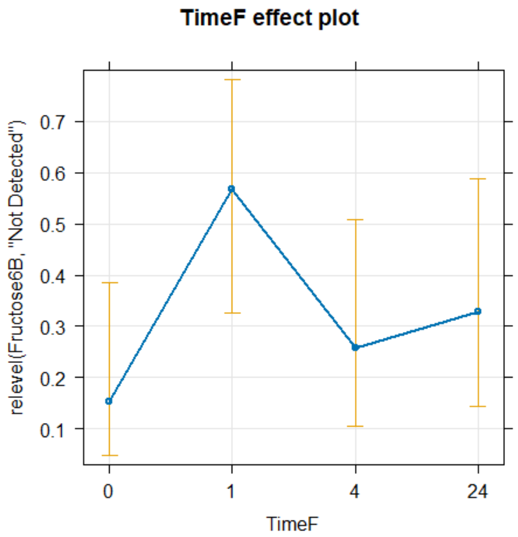


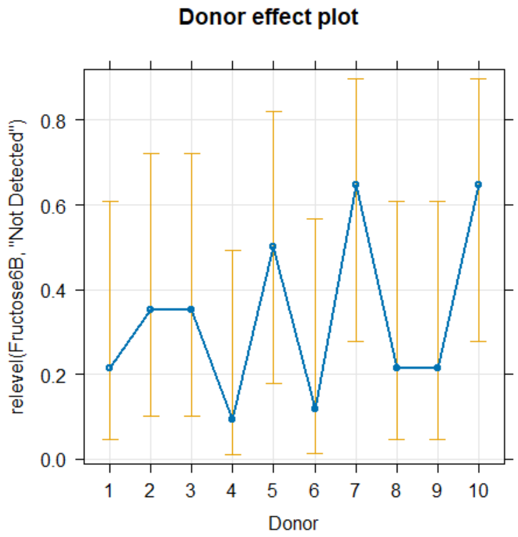


```
confint(glm2)

##                  2.5 %     97.5 %
## (Intercept) -5.3155384 -0.7610275
## LabelN       0.1110587  2.3396441
## TimeF1       0.4604681  3.7244469
## TimeF4      -0.9329975  2.3481498
## TimeF24     -0.5670577  2.7026975
## Donor2      -1.6241522  3.1810140
## Donor3      -1.6241522  3.1810140
## Donor4      -4.2857055  1.7696581
## Donor5      -0.9359352  3.8043040
## Donor6      -4.0792577  2.0691575
## Donor7      -0.3252864  4.4836840
## Donor8      -2.5378189  2.5378189
## Donor9      -2.5378189  2.5378189
## Donor10     -0.3252864  4.4836840
```

```r
pdf("OA_Glucose_Loading.pdf",width=6, height=4)
pirateplot(formula=GlucoseTT~TimeF+Label, data=TICsetOAR, theme=3, inf.b.o =
0,inf.f.o = 0, ylab="TIC Corrected Glucose*10e6", main="Glucose Concentration
OA")
```

OA Glutamine Plot

```r
ggintplot(response="Glutamineplot", groupvar = c("Label", "TimeF"), array =
F, data = TICsetOA)
```

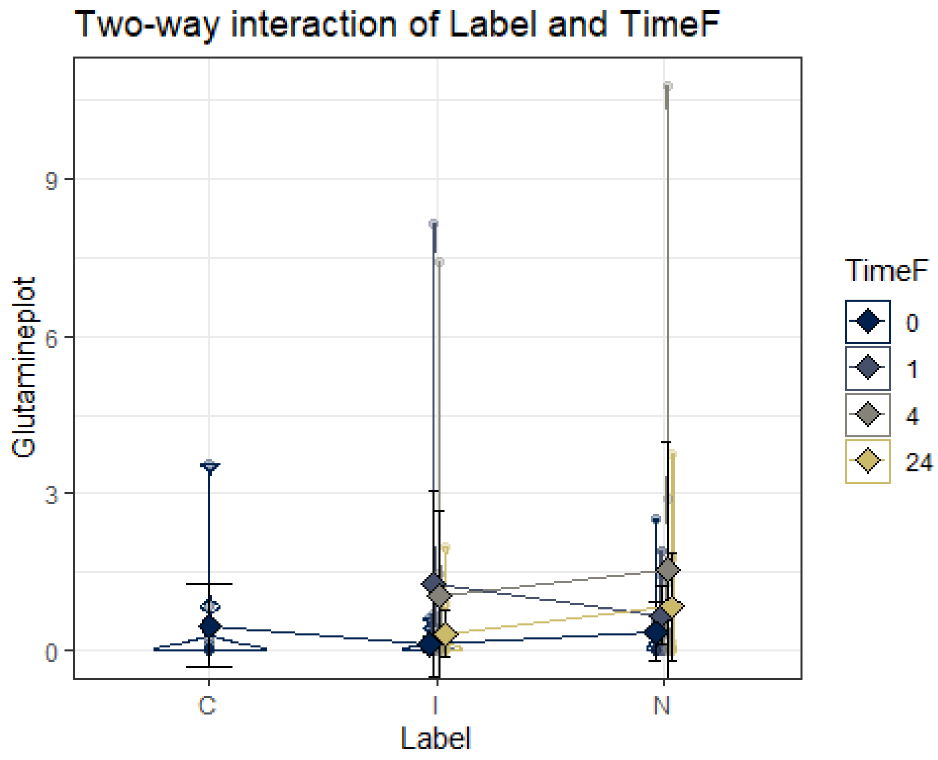


Bovine metabolite vs time and loading

```r
library(readxl)
TICset <- read_excel("InjuryBovine_With_TIC.xlsx")
TICset <- TICset %>% mutate(Donor = factor(Donor))
dim(TICset)

## [1] 90 51

tally(Time ~ Label, data = TICset)

##      Label
## Time   C  I  N
##    0  10 10 10
##    1   0 10 10
##    4   0 10 10
##    24  0 10 10
```

```r
TICset <- TICset %>% mutate(TimeF = factor(Time),
                            Label.Time = str_c(Label, TimeF))
```

For bovine metabolites the same approach was taken above to split the analysis into two groups starting with loading and normal at all time points followed by loading, normal and control at time point 0.

Glutamine

The full model includes and interaction between Loading and Time as well as donor as a random variable. The interaction was simplified out of the model as there was no evidence of its impact of the metabolite abundance.A log transformation was done on this data to meet the constant variance assumption.

```r
TICsetR <- TICset %>% dplyr::filter(Label != "C")

ggintplot(response="GlutamineT", groupvar = c("Label", "TimeF"), array = F,
data = TICsetR)
```

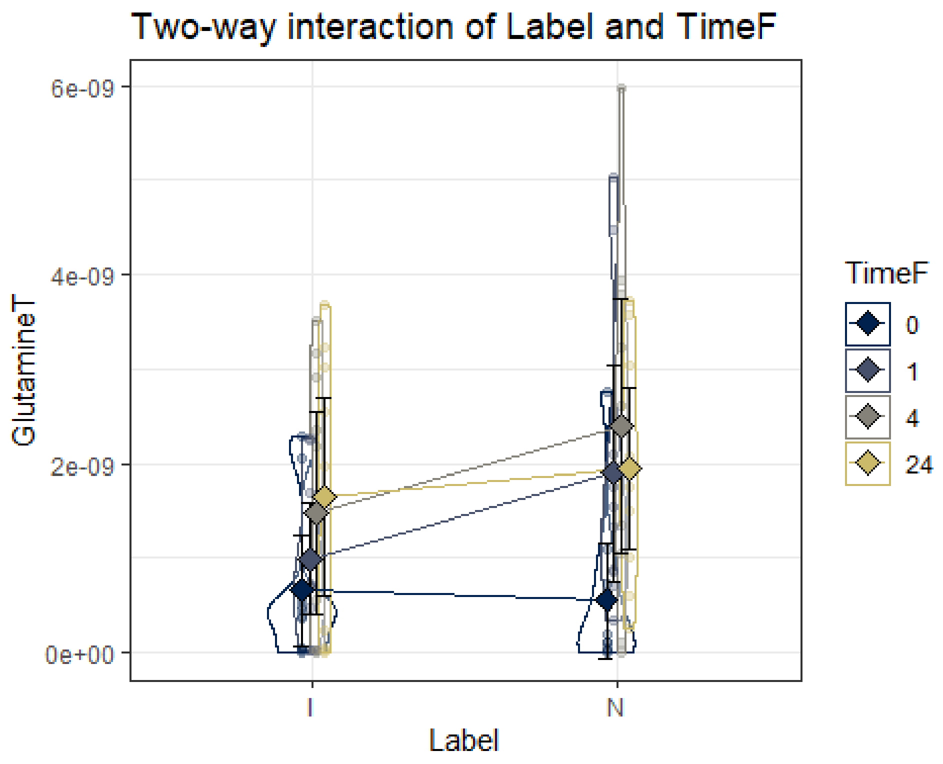


```r
lmer1 <- lmer(GlutamineTlog ~ Label*TimeF+(1|Donor) , data=TICsetR)
Anova(lmer1,test.statistic = "F")

## Analysis of Deviance Table (Type II Wald F tests with Kenward-Roger df)
##
## Response: GlutamineTlog
##                  F Df Df.res  Pr(>F)
## Label       3.5930  1     63 0.06261
```

```
## TimeF       2.5787  3     63 0.06144
## Label:TimeF 0.8822  3     63 0.45524

lm1 <- lmer(GlutamineTlog ~ Label+TimeF+(1|Donor) , data=TICsetR)
Anova(lm1,test.statistic = "F")

## Analysis of Deviance Table (Type II Wald F tests with Kenward-Roger df)
##
## Response: GlutamineTlog
##            F Df Df.res  Pr(>F)
## Label 3.6124  1     66 0.06172
## TimeF 2.5926  3     66 0.05999

summary(lm1)

## Linear mixed model fit by REML ['lmerMod']
## Formula: GlutamineTlog ~ Label + TimeF + (1 | Donor)
##    Data: TICsetR
##
## REML criterion at convergence: 203
##
## Scaled residuals:
##     Min      1Q  Median      3Q     Max
## -2.4857 -0.5044  0.2531  0.6707  1.5929
##
## Random effects:
##  Groups   Name        Variance Std.Dev.
##  Donor    (Intercept) 0.01857  0.1363
##  Residual             0.70169  0.8377
## Number of obs: 80, groups:  Donor, 10
##
## Fixed effects:
##             Estimate Std. Error t value
## (Intercept)  -9.9073     0.2138 -46.338
## LabelN        0.3560     0.1873   1.901
## TimeF1        0.6357     0.2649   2.400
## TimeF4        0.4522     0.2649   1.707
## TimeF24       0.6426     0.2649   2.426
##
## Correlation of Fixed Effects:
##         (Intr) LabelN TimeF1 TimeF4
## LabelN  -0.438
## TimeF1  -0.619  0.000
## TimeF4  -0.619  0.000  0.500
## TimeF24 -0.619  0.000  0.500  0.500

#If need Tukey's in interaction model:

TukeyRes2 <- emmeans(lm1, pairwise ~ TimeF, adjust = "tukey")
TukeyRes2
```

```
## $emmeans
##  TimeF emmean    SE   df lower.CL upper.CL
##  0      -9.73 0.192 59.2   -10.11    -9.34
##  1      -9.09 0.192 59.2    -9.48    -8.71
##  4      -9.28 0.192 59.2    -9.66    -8.89
##  24     -9.09 0.192 59.2    -9.47    -8.70
##
## Results are averaged over the levels of: Label
## Degrees-of-freedom method: kenward-roger
## Confidence level used: 0.95
##
## $contrasts
##  contrast         estimate    SE df t.ratio p.value
##  TimeF0 - TimeF1  -0.63571 0.265 66  -2.400  0.0870
##  TimeF0 - TimeF4  -0.45221 0.265 66  -1.707  0.3282
##  TimeF0 - TimeF24 -0.64259 0.265 66  -2.426  0.0821
##  TimeF1 - TimeF4   0.18350 0.265 66   0.693  0.8995
##  TimeF1 - TimeF24 -0.00688 0.265 66  -0.026  1.0000
##  TimeF4 - TimeF24 -0.19038 0.265 66  -0.719  0.8893
##
## Results are averaged over the levels of: Label
## Degrees-of-freedom method: kenward-roger
## P value adjustment: tukey method for comparing a family of 4 estimates
```

```
confint(TukeyRes2)
```

```
## $emmeans
##  TimeF emmean    SE   df lower.CL upper.CL
##  0      -9.73 0.192 59.2   -10.11    -9.34
##  1      -9.09 0.192 59.2    -9.48    -8.71
##  4      -9.28 0.192 59.2    -9.66    -8.89
##  24     -9.09 0.192 59.2    -9.47    -8.70
##
## Results are averaged over the levels of: Label
## Degrees-of-freedom method: kenward-roger
## Confidence level used: 0.95
##
## $contrasts
##  contrast         estimate    SE df lower.CL upper.CL
##  TimeF0 - TimeF1  -0.63571 0.265 66   -1.334   0.0625
##  TimeF0 - TimeF4  -0.45221 0.265 66   -1.150   0.2460
##  TimeF0 - TimeF24 -0.64259 0.265 66   -1.341   0.0556
##  TimeF1 - TimeF4   0.18350 0.265 66   -0.515   0.8817
##  TimeF1 - TimeF24 -0.00688 0.265 66   -0.705   0.6913
##  TimeF4 - TimeF24 -0.19038 0.265 66   -0.889   0.5078
##
## Results are averaged over the levels of: Label
## Degrees-of-freedom method: kenward-roger
## Confidence level used: 0.95
## Conf-level adjustment: tukey method for comparing a family of 4 estimates
```

```r
plot(TukeyRes2, comparison = T) + coord_flip()
```

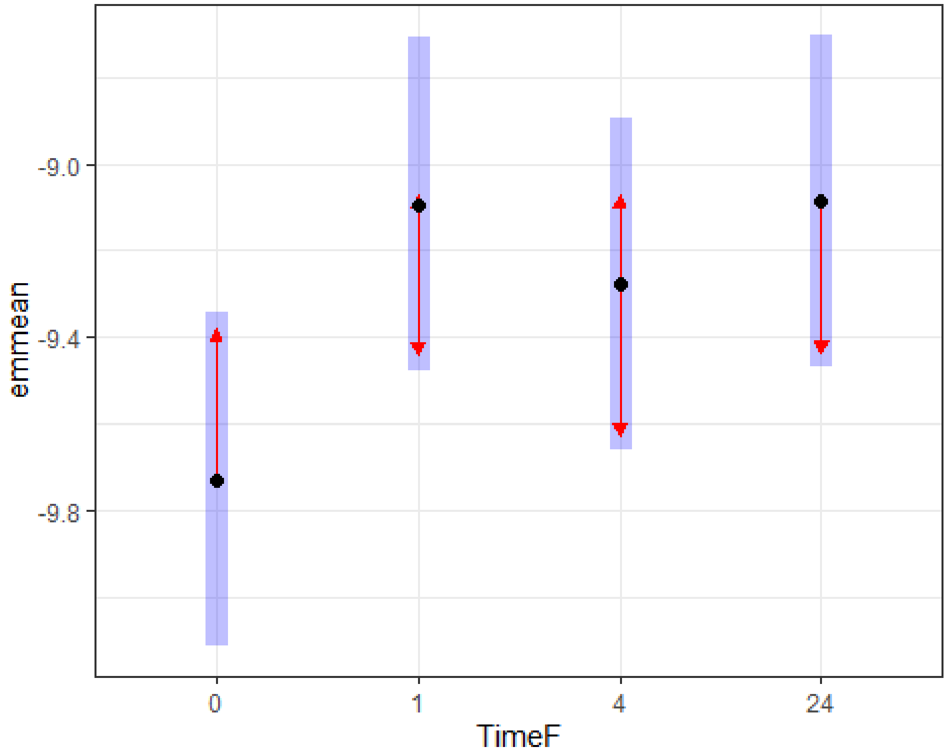


```r
multcomp::cld(TukeyRes2, Letters = LETTERS)

##  TimeF emmean    SE   df lower.CL upper.CL .group
##  0      -9.73 0.192 59.2   -10.11    -9.34  A
##  4      -9.28 0.192 59.2    -9.66    -8.89  A
##  1      -9.09 0.192 59.2    -9.48    -8.71  A
##  24     -9.09 0.192 59.2    -9.47    -8.70  A
##
## Results are averaged over the levels of: Label
## Degrees-of-freedom method: kenward-roger
## Confidence level used: 0.95
## P value adjustment: tukey method for comparing a family of 4 estimates
## significance level used: alpha = 0.05
## NOTE: If two or more means share the same grouping symbol,
##       then we cannot show them to be different.
##       But we also did not show them to be the same.

plot(allEffects(lm1), grid = T)
```

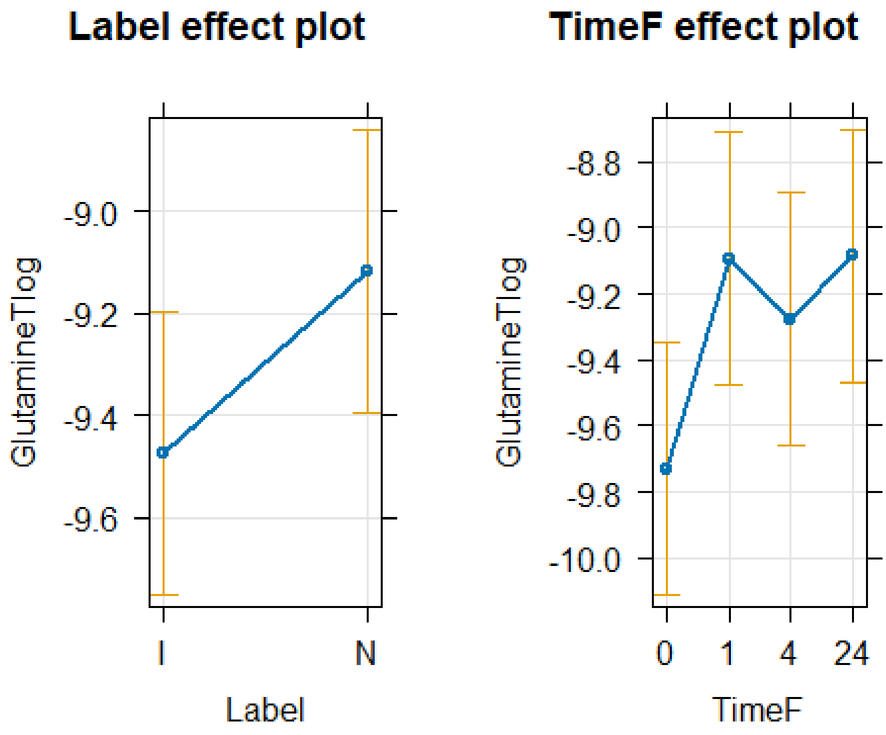


```r
resid_panel(lm1)
```

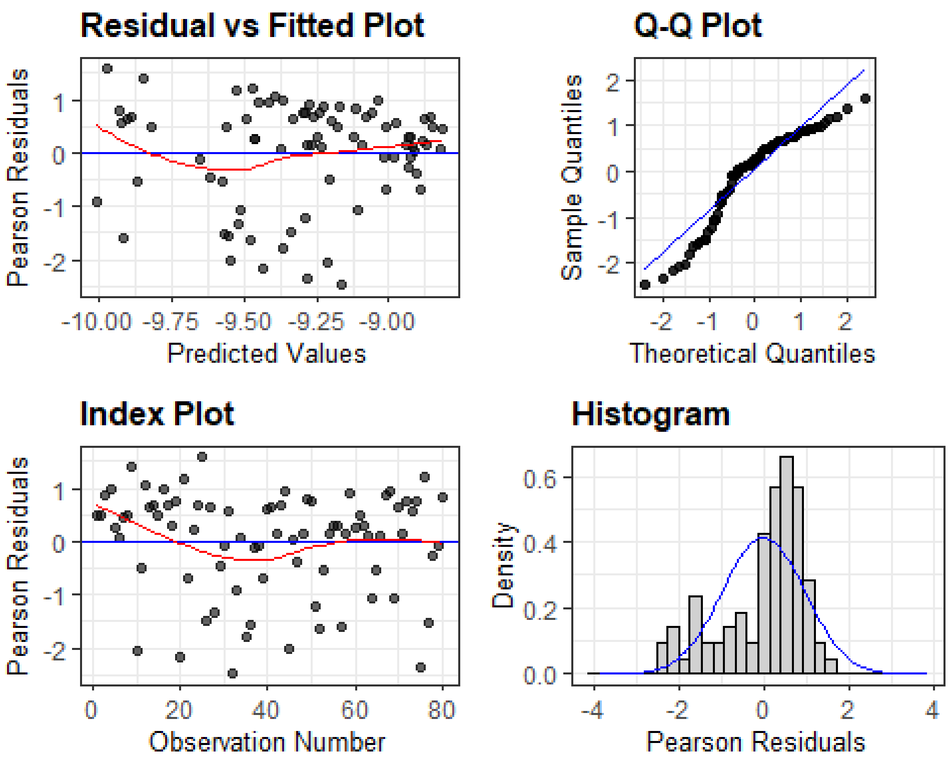

```
#plot(allEffects(lm1, residuals = T), x.var = "TimeF:Label")
#plot(allEffects(lm1), x.var = "Label", grid = T)
```

Bovine Glutamine plot

```
#pdf("Bovine_Glutamine_Time.pdf",width=6, height=4)
pirateplot(formula=GlutamineT~Label+Time, data=TICsetR, theme=3, inf.b.o = 
0,inf.f.o = 0, ylab="Logged TIC Corrected Glutamine", main="Glutamine 
Concentration",)
text(x=1,y=8,"A")
text(x=2,y=8,"A")
text(x=3,y=8,"A")
text(x=4,y=8,"A")
```

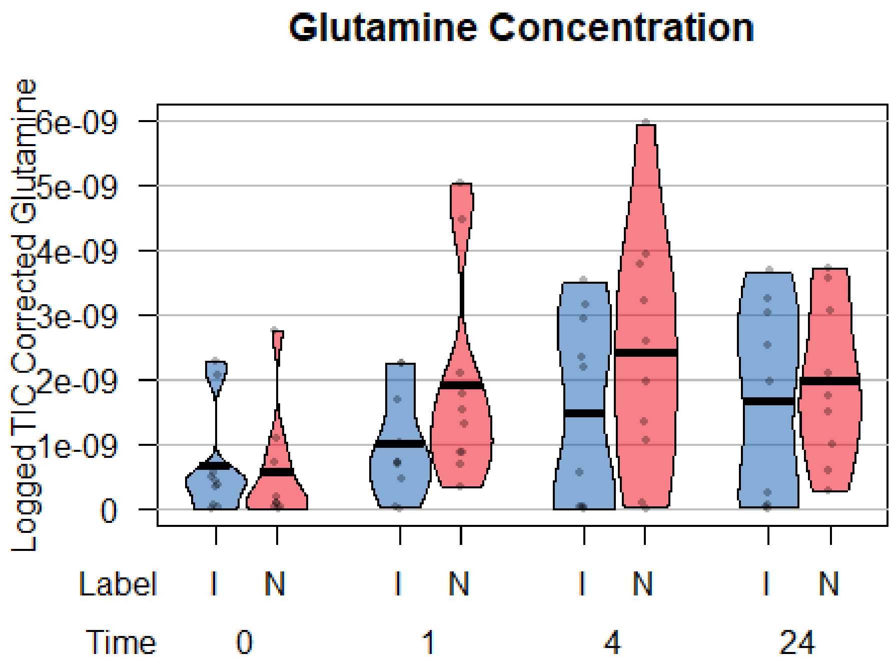


Glucose

```
#pdf("Bovine_Glucose_Loading.pdf",width=6, height=4)
pirateplot(formula=GlucoseT~TimeF+Label, data=TICsetR, theme=3, inf.b.o = 
0,inf.f.o = 0, ylab="TIC Corrected Glucose*10e6", main="Glucose Concentration 
p=0.029",ylim=c(0,1.1))
text(x=2.5,y=0.95,"A")
segments(x0=1,y0=.85,x1=4,y1=0.85)
text(x=7.5,y=1.05,"B")
segments(x0=6,y0=0.95,x1=9,y1=0.95)
```

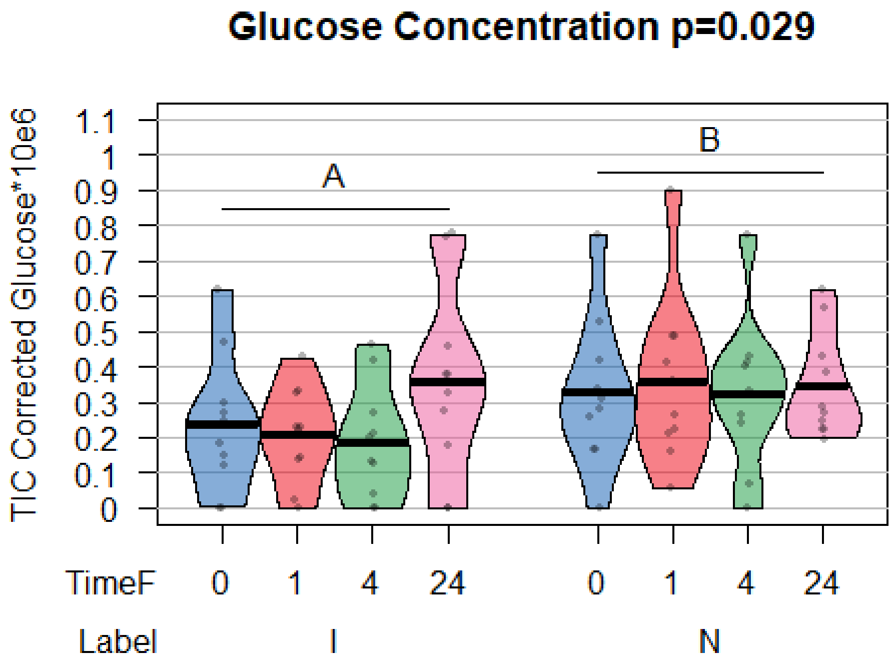


```
ggintplot(response="GlucoseT", groupvar = c("Label", "TimeF"), array = F,
data = TICsetR)
```

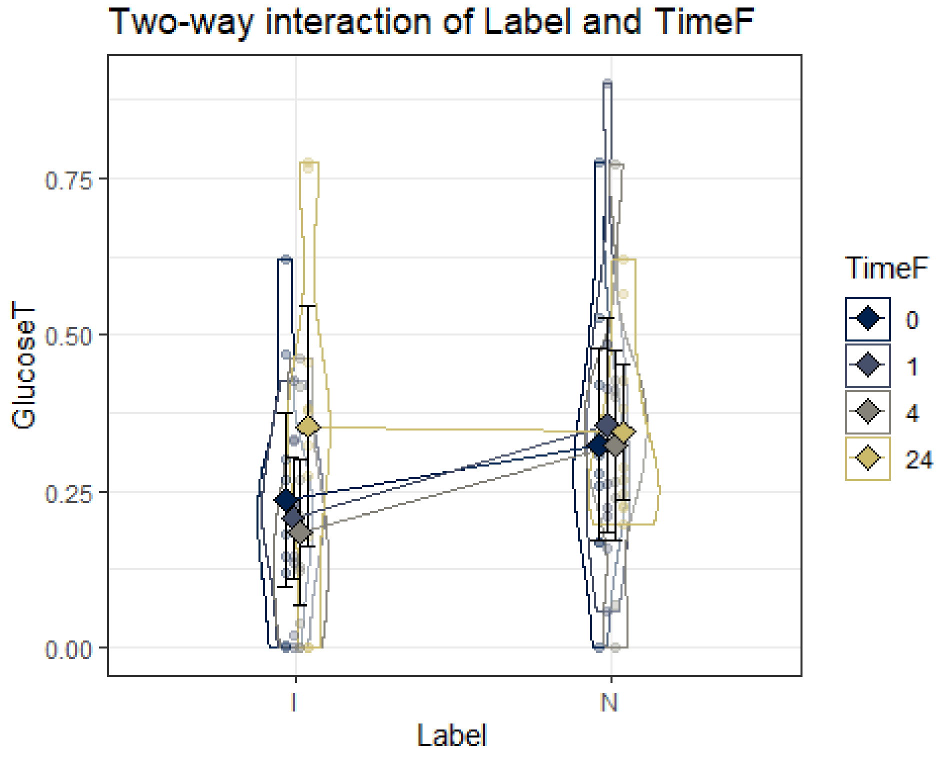

```r
lmer1 <- lmer(GlucoseT ~ Label*TimeF+(1|Donor) , data=TICsetR)
Anova(lmer1,test.statistic = "F")

## Analysis of Deviance Table (Type II Wald F tests with Kenward-Roger df)
##
## Response: GlucoseT
##                   F Df Df.res  Pr(>F)
## Label       4.9282  1     63 0.03003
## TimeF       0.9793  3     63 0.40824
## Label:TimeF 0.7667  3     63 0.51695

lm1 <- lmer(GlucoseT ~ Label+TimeF+(1|Donor) , data=TICsetR)
Anova(lm1,test.statistic = "F")

## Analysis of Deviance Table (Type II Wald F tests with Kenward-Roger df)
##
## Response: GlucoseT
##             F Df Df.res  Pr(>F)
## Label 4.9810  1     66 0.02903
## TimeF 0.9898  3     66 0.40311

#If need Tukey's in interaction model:

confint(lm1)

##                    2.5 %     97.5 %
## .sig01       0.01979456 0.15350840
## .sigma       0.15203258 0.21196033
## (Intercept)  0.13257372 0.33537306
## LabelN       0.01238147 0.17045599
## TimeF1      -0.11043947 0.11311166
## TimeF4      -0.13765561 0.08589552
## TimeF24     -0.04239747 0.18115366

summary(lm1)

## Linear mixed model fit by REML ['lmerMod']
## Formula: GlucoseT ~ Label + TimeF + (1 | Donor)
##    Data: TICsetR
##
## REML criterion at convergence: -18.1
##
## Scaled residuals:
##      Min       1Q   Median       3Q      Max
## -1.53007 -0.63029 -0.09068  0.42817  2.56116
##
## Random effects:
##  Groups   Name        Variance Std.Dev.
##  Donor    (Intercept) 0.00676  0.08222
##  Residual             0.03356  0.18319
## Number of obs: 80, groups:  Donor, 10
```

```
##
## Fixed effects:
##              Estimate Std. Error t value
## (Intercept)  0.233973   0.052663   4.443
## LabelN       0.091419   0.040961   2.232
## TimeF1       0.001336   0.057928   0.023
## TimeF4      -0.025880   0.057928  -0.447
## TimeF24      0.069378   0.057928   1.198
##
## Correlation of Fixed Effects:
##         (Intr) LabelN TimeF1 TimeF4
## LabelN  -0.389
## TimeF1  -0.550  0.000
## TimeF4  -0.550  0.000  0.500
## TimeF24 -0.550  0.000  0.500  0.500
```

```
plot(allEffects(lm1), grid = T)
```

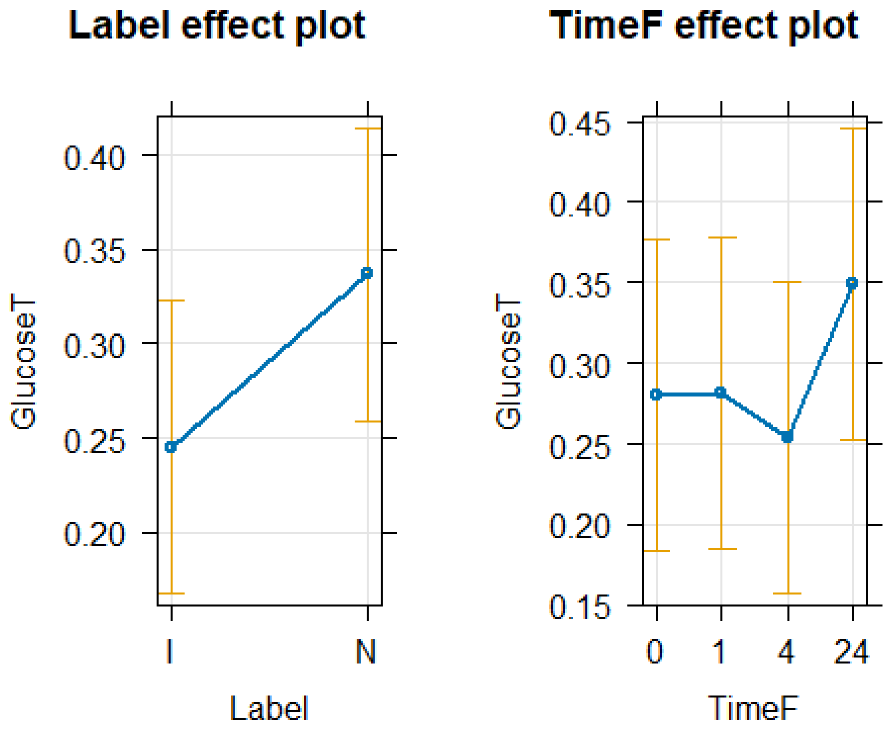


```
resid_panel(lm1)
```

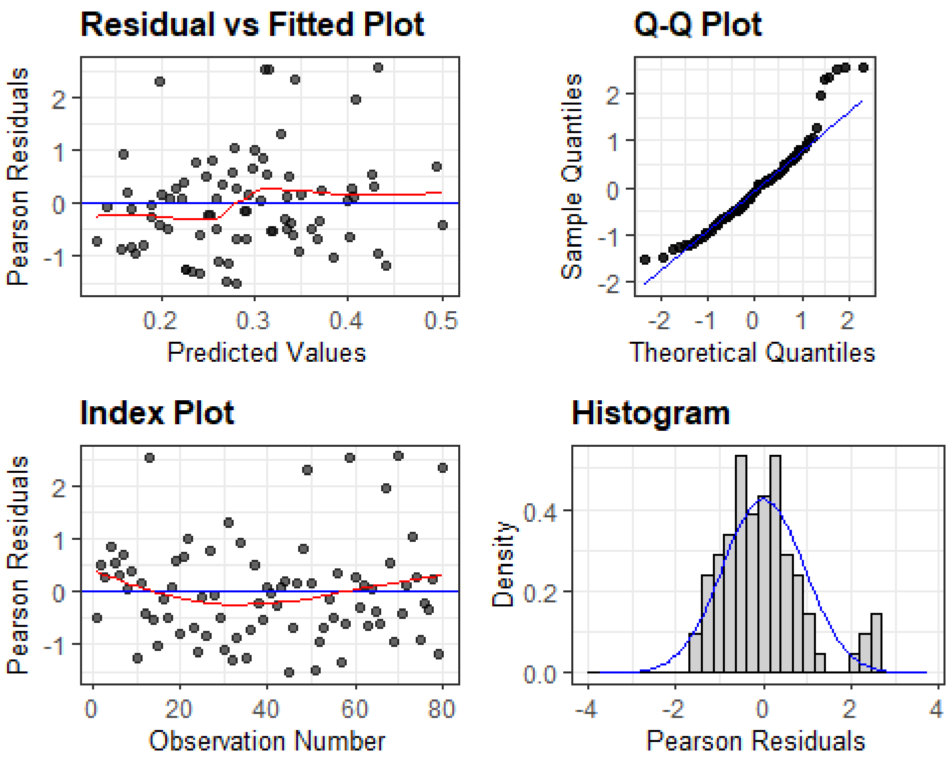


```r
#plot(allEffects(lm1, residuals = T), x.var = "TimeF:Label")
#plot(allEffects(lm1), x.var = "Label", grid = T)
```

Here we switch to the second type of analysis looking at the three loading conditions just at time 0.

Glutamate There was no evidence to support an impact of loading condition on glutamate concentration at time point 0.

```r
TICsetR2 <- TICset %>% dplyr::filter(TimeF == "0") %>%
  mutate(GlutemicAcidB = factor(GlutemicAcidB))


glm2 <- glm(GlutemicAcidB ~ Label+Donor, data=TICsetR2, family = "binomial")

#Can't add donor - causes separation here
Anova(glm2,test.statistic = "F")

## Analysis of Deviance Table (Type II tests)
##
## Response: GlutemicAcidB
## Error estimate based on Pearson residuals
##
##            Sum Sq Df F value Pr(>F)
## Label      2.7485  2  1.0278 0.3779
## Donor      9.5270  9  0.7917 0.6280
## Residuals 24.0680 18
```

```r
summary(glm2)

##
## Call:
## glm(formula = GlutemicAcidB ~ Label + Donor, family = "binomial",
##     data = TICsetR2)
##
## Coefficients:
##                Estimate Std. Error z value Pr(>|z|)
## (Intercept) -1.834e+00  1.544e+00  -1.188    0.235
## LabelI       1.841e+00  1.199e+00   1.536    0.124
## LabelN       1.261e+00  1.171e+00   1.077    0.282
## Donor5      -2.538e-15  1.827e+00   0.000    1.000
## Donor6       1.581e+00  1.856e+00   0.852    0.394
## Donor7       1.581e+00  1.856e+00   0.852    0.394
## Donor8       1.581e+00  1.856e+00   0.852    0.394
## Donor9      -1.790e+01  3.555e+03  -0.005    0.996
## Donor10     -1.790e+01  3.555e+03  -0.005    0.996
## Donor11     -5.329e-15  1.827e+00   0.000    1.000
## Donor12     -4.636e-15  1.827e+00   0.000    1.000
## Donor13     -3.651e-15  1.827e+00   0.000    1.000
##
## (Dispersion parameter for binomial family taken to be 1)
##
##     Null deviance: 39.429  on 29  degrees of freedom
## Residual deviance: 27.804  on 18  degrees of freedom
## AIC: 51.804
##
## Number of Fisher Scoring iterations: 17

plot(allEffects(glm2), type = "response", grid = T)
```

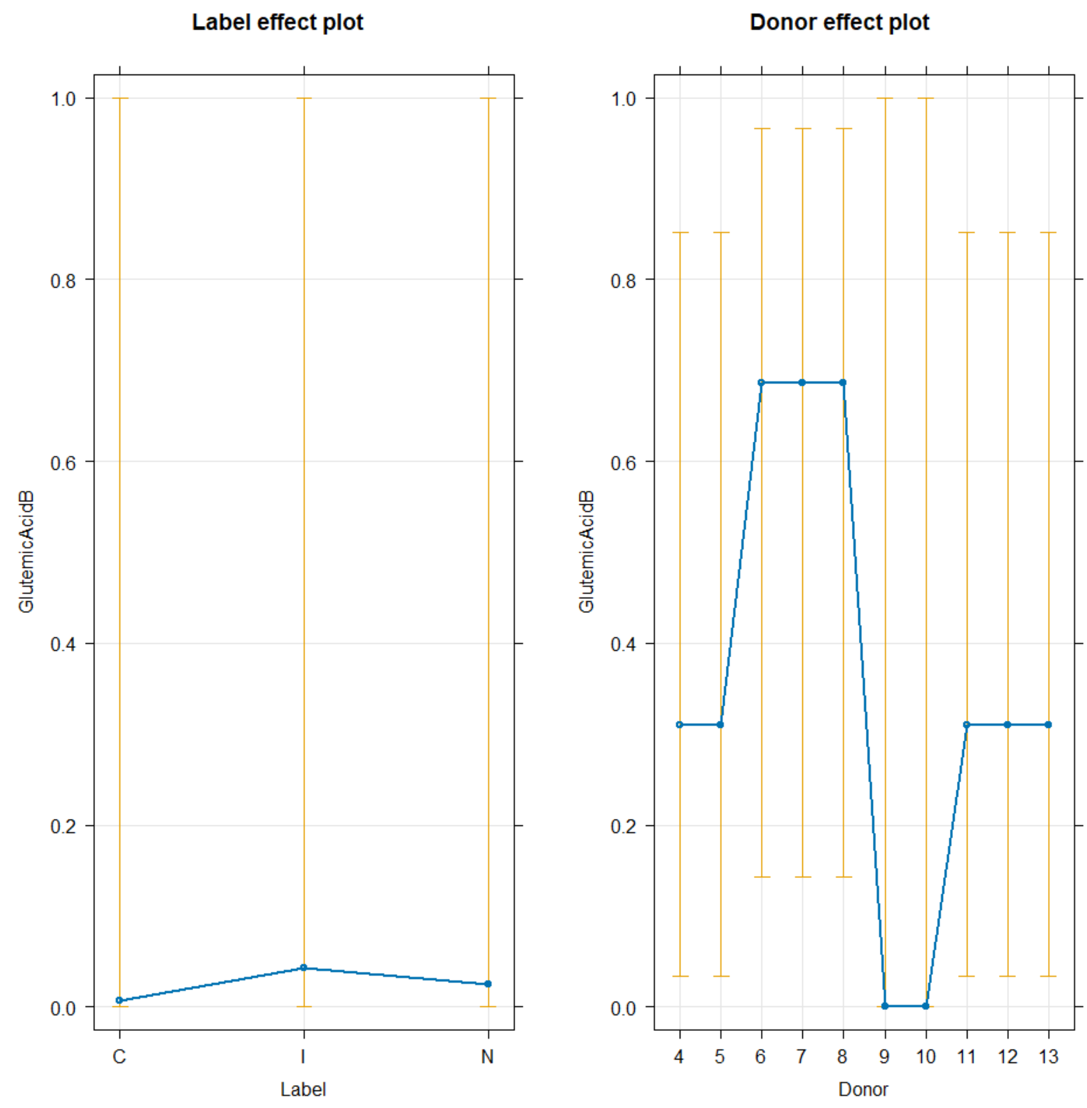


Metabolites vs oxygen consumption.

For the following analysis both the metabolite value and oxygen consumption is reported as change from the control of the same donor. Each donor was assigned to either a hypoxic or normoxic environment. Non-detects were considered 0s for this analysis.

```
library(readxl)

TICsetOAO <- read_excel("OAO2data.xlsx")
dim(TICsetOAO)

## [1] 79 24

tally(Oxygen~Donor, data=TICsetOAO)

##             Donor
## Oxygen       1  2  3  4  5  6  7  8  9  10
```

```
##    Hypoxic  0 0 0 0 8 7 8 0 0  0
##    Normoxic 8 8 8 8 0 0 0 8 8  8

 TICsetOAO <- TICsetOAO %>% drop_na(O2data) # Remove samples where O2 reading
could not be obtained
 dim(TICsetOAO)

## [1] 75 24
```

For the following generalized linear models the full model includes an interaction between oxygen consumption and the hypoxic vs normoxic enviornment. This interaction was removed from the model if there was not enough evidence of the interaction having an impact on the metabolite abundance.

OA Glutamine

```
G1<- TICsetOAO %>%
  ggplot(aes(x=O2data, fill=GlutamineBi))  + facet_wrap(~Oxygen)+
  geom_density(position='fill')+ scale_fill_colorblind()+
geom_rug(aes(col=GlutamineBi),alpha=0.1) + scale_color_colorblind()

G1
```

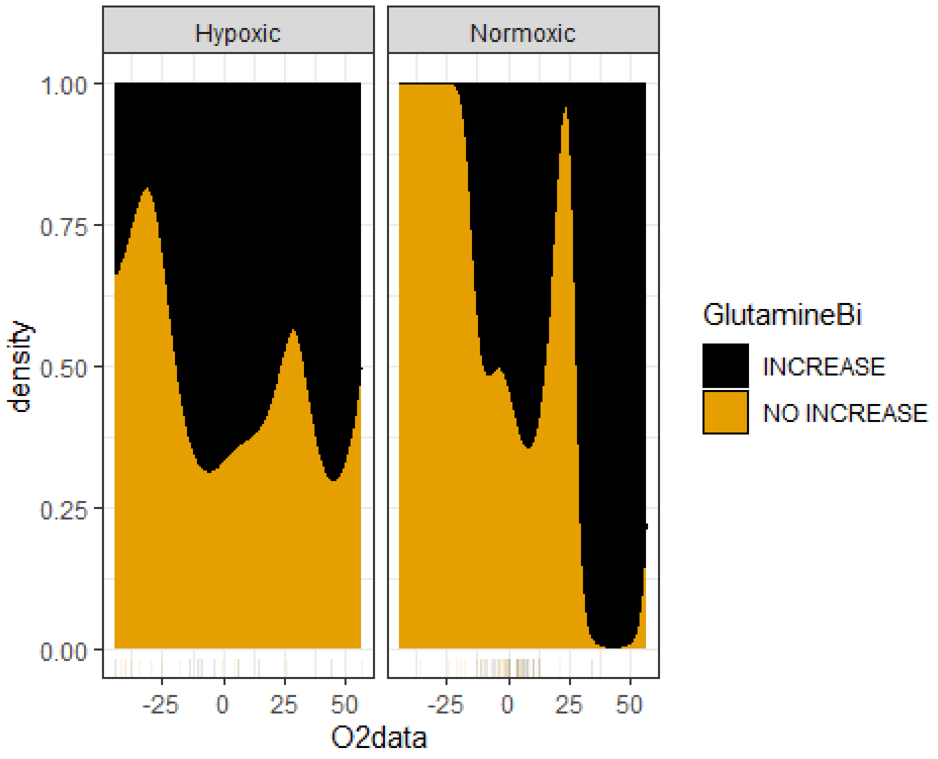


```
D1<- TICsetOAO %>%
  ggplot() + scale_fill_paletteer_d("nationalparkcolors::Acadia")  +
  geom_mosaic(aes(x=product(Donor),fill=GlutamineBi), offset=0.02)
D1
```

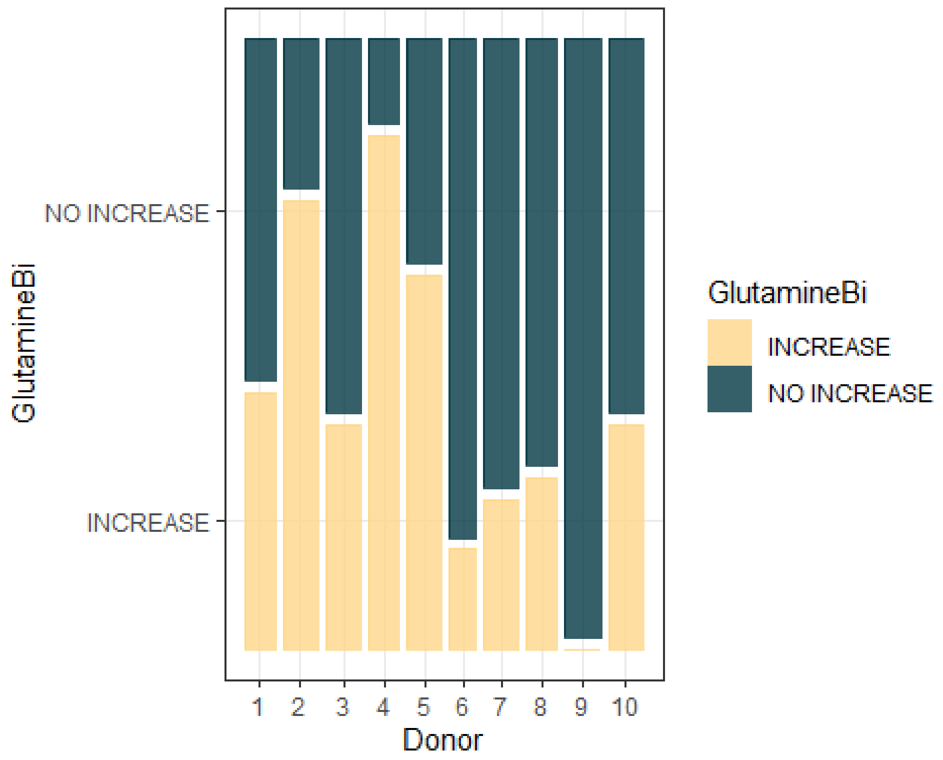


```
TICsetOAO <- TICsetOAO %>% mutate(GlutamineBi = factor(GlutamineBi),
                                  Timef=factor(Time),
                                  Donor=factor(Donor))
glm1 <- glm(GlutamineBi ~ O2data*Oxygen, data=TICsetOAO, family = "binomial")
#Donor could not be included in the model due to the pattern of binary
responses creating separation

summary(glm1)

##
## Call:
## glm(formula = GlutamineBi ~ O2data * Oxygen, family = "binomial",
##     data = TICsetOAO)
##
## Coefficients:
##                         Estimate Std. Error z value Pr(>|z|)
## (Intercept)              0.44160    0.46616   0.947    0.343
## O2data                  -0.01504    0.01662  -0.905    0.366
## OxygenNormoxic          -0.17887    0.54871  -0.326    0.744
## O2data:OxygenNormoxic -0.04259    0.03401  -1.252    0.210
##
## (Dispersion parameter for binomial family taken to be 1)
##
##     Null deviance: 101.707  on 74  degrees of freedom
## Residual deviance:  95.963  on 71  degrees of freedom
## AIC: 103.96
```

```
##
## Number of Fisher Scoring iterations: 4

plot(allEffects(glm1), type = "response", grid = T)
```

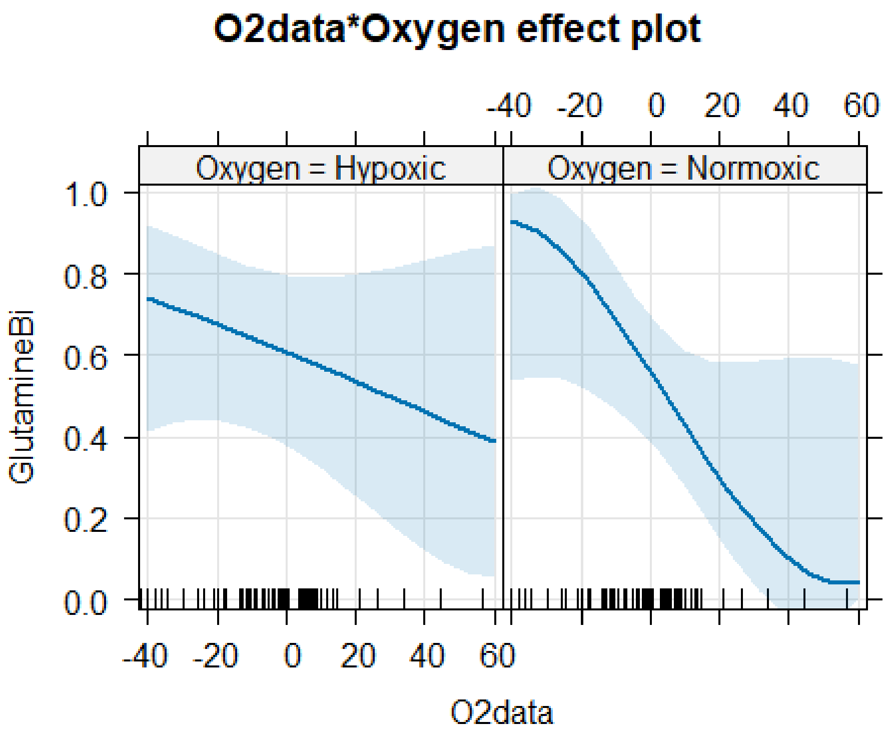


```
plot(allEffects(glm1, residuals=T), type = "link", grid = T)
```

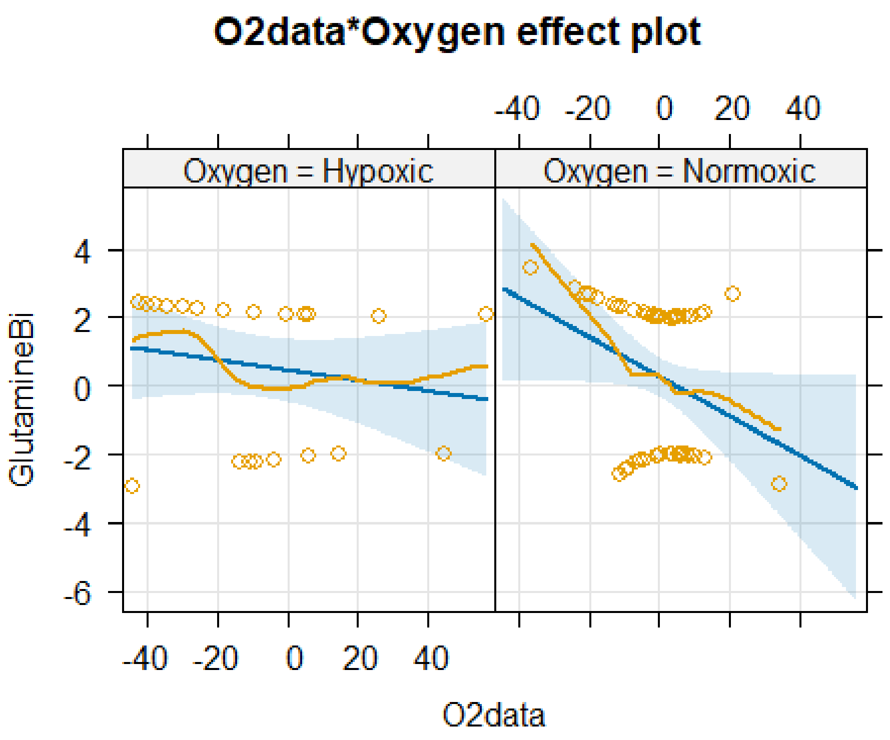


```
confint(glm1)

##                               2.5 %     97.5 %
## (Intercept)              -0.4764182 1.39104103
## O2data                   -0.0508869 0.01715832
## OxygenNormoxic           -1.2773689 0.90020910
## O2data:OxygenNormoxic -0.1146590 0.02106286

glm2 <- glm(GlutamineBi ~ O2data+Oxygen, data=TICsetOAO, family = "binomial")

summary(glm2)

##
## Call:
## glm(formula = GlutamineBi ~ O2data + Oxygen, family = "binomial",
##     data = TICsetOAO)
##
## Coefficients:
##                Estimate Std. Error z value Pr(>|z|)
## (Intercept)     0.36345    0.47967   0.758   0.4486
## O2data         -0.02722    0.01487  -1.831   0.0672
## OxygenNormoxic -0.10645    0.55539  -0.192   0.8480
##
## (Dispersion parameter for binomial family taken to be 1)
##
##     Null deviance: 101.707  on 74  degrees of freedom
## Residual deviance:  97.646  on 72  degrees of freedom
```

```
## AIC: 103.65
##
## Number of Fisher Scoring iterations: 4
```

```
plot(allEffects(glm2), type = "response", grid = T)
```

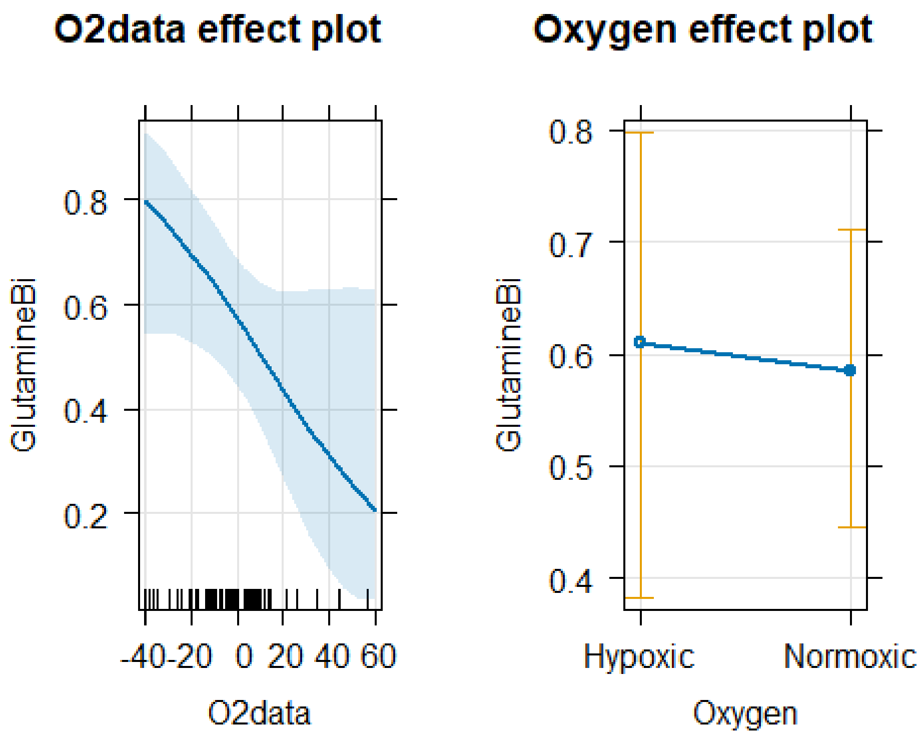


```
plot(allEffects(glm2, residuals=T), type = "link", grid = T)
```

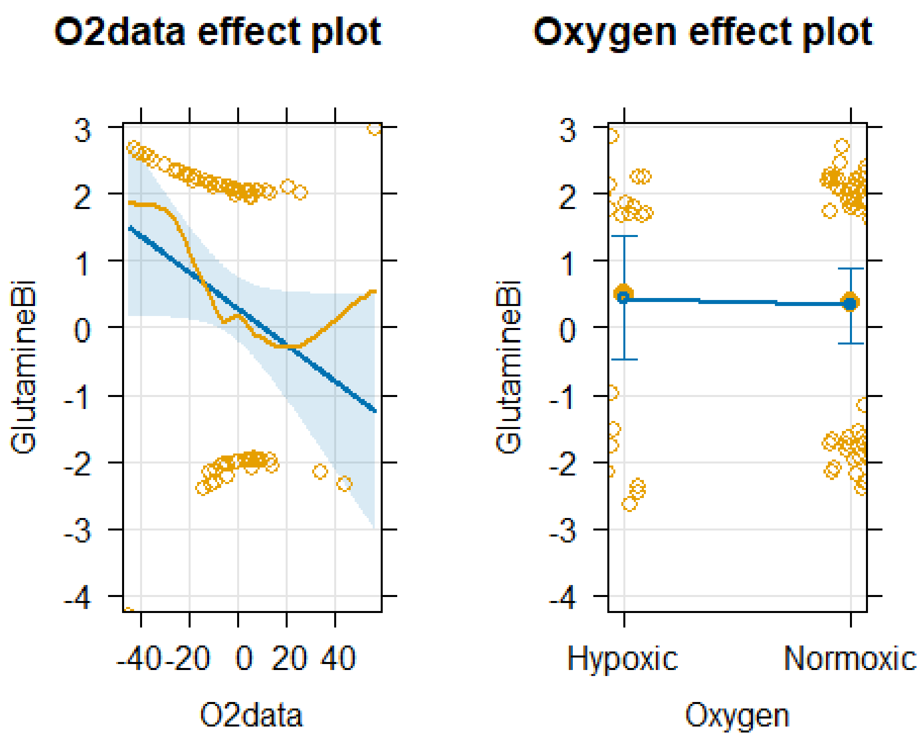


```
confint(glm2)

##                        2.5 %       97.5 %
## (Intercept)     -0.58284230 1.3359110747
## O2data          -0.05917204 0.0003491336
## OxygenNormoxic -1.21619167 0.9889693997
```

OA Glucose

```
G2<- TICsetOAO %>%
  ggplot(aes(x=O2data, fill=GlucoseBi))  + facet_wrap(~Oxygen)+
  geom_density(position='fill')+ scale_fill_colorblind()+
geom_rug(aes(col=GlucoseBi),alpha=0.1) + scale_color_colorblind()

G2
```

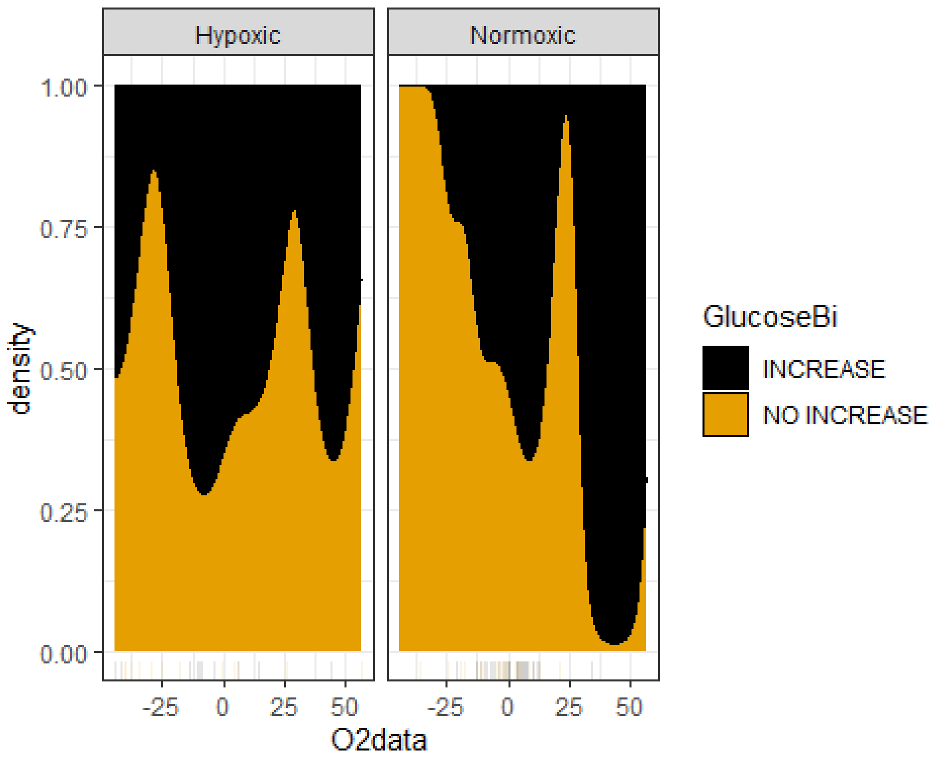


```
D2<- TICsetOAO %>%
  ggplot() + scale_fill_paletteer_d("nationalparkcolors::Acadia")  +
  geom_mosaic(aes(x=product(Donor),fill=GlucoseBi), offset=0.02)
D2
```

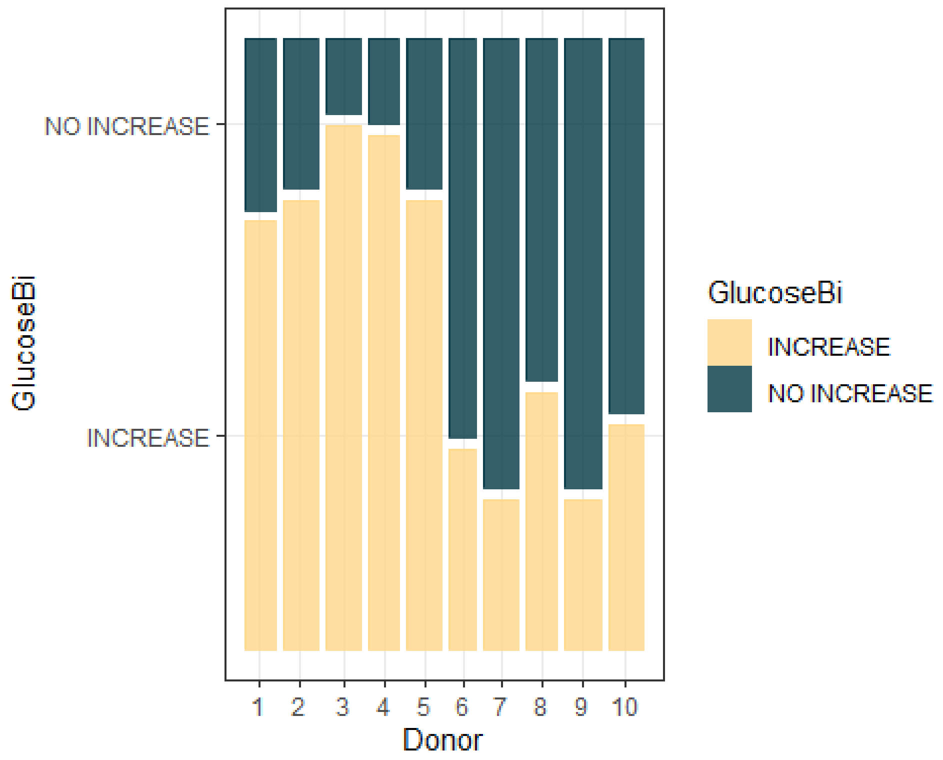


```
TICsetOAO <- TICsetOAO %>% mutate(GlucoseBi = factor(GlucoseBi),
                                  Timef=factor(Time),
                                  Donor=factor(Donor))
glm1 <- glm(GlucoseBi ~ O2data*Oxygen, data=TICsetOAO, family = "binomial")
#Donor could not be included in the model due to the pattern of binary
responses creating separation

summary(glm1)

##
## Call:
## glm(formula = GlucoseBi ~ O2data * Oxygen, family = "binomial",
##     data = TICsetOAO)
##
## Coefficients:
##                          Estimate Std. Error z value Pr(>|z|)
## (Intercept)              0.130691   0.451667   0.289    0.772
## O2data                  -0.005759   0.015800  -0.364    0.715
## OxygenNormoxic          -0.607353   0.538751  -1.127    0.260
## O2data:OxygenNormoxic -0.044632   0.032154  -1.388    0.165
##
## (Dispersion parameter for binomial family taken to be 1)
##
##     Null deviance: 102.889  on 74  degrees of freedom
## Residual deviance:  97.651  on 71  degrees of freedom
## AIC: 105.65
```

```
##
## Number of Fisher Scoring iterations: 4
```

```r
plot(allEffects(glm1), type = "response", grid = T)
```

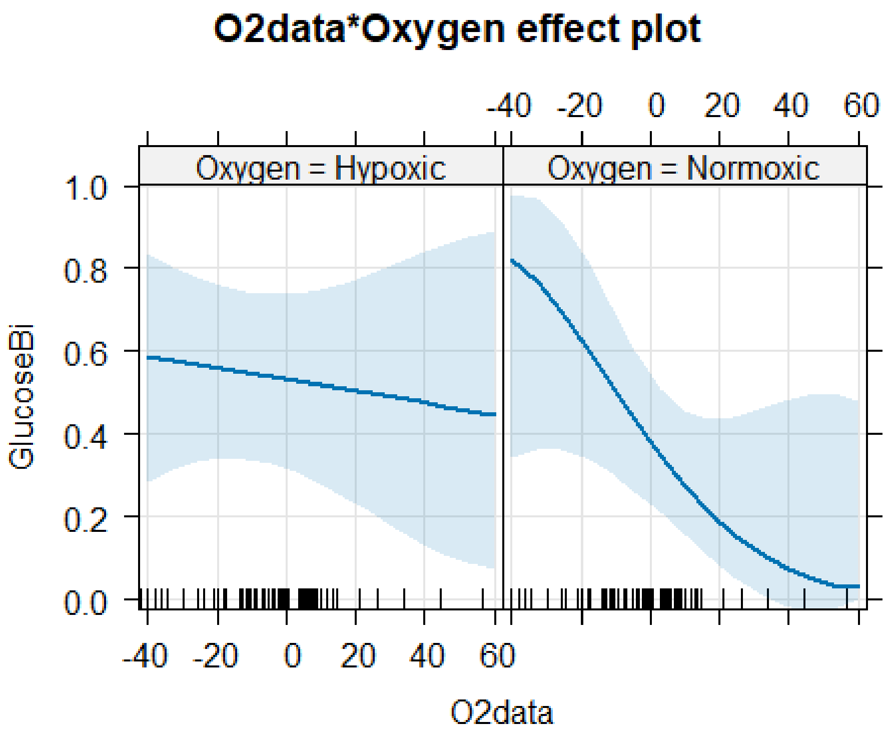


```r
plot(allEffects(glm1, residuals=T), type = "link", grid = T)
```

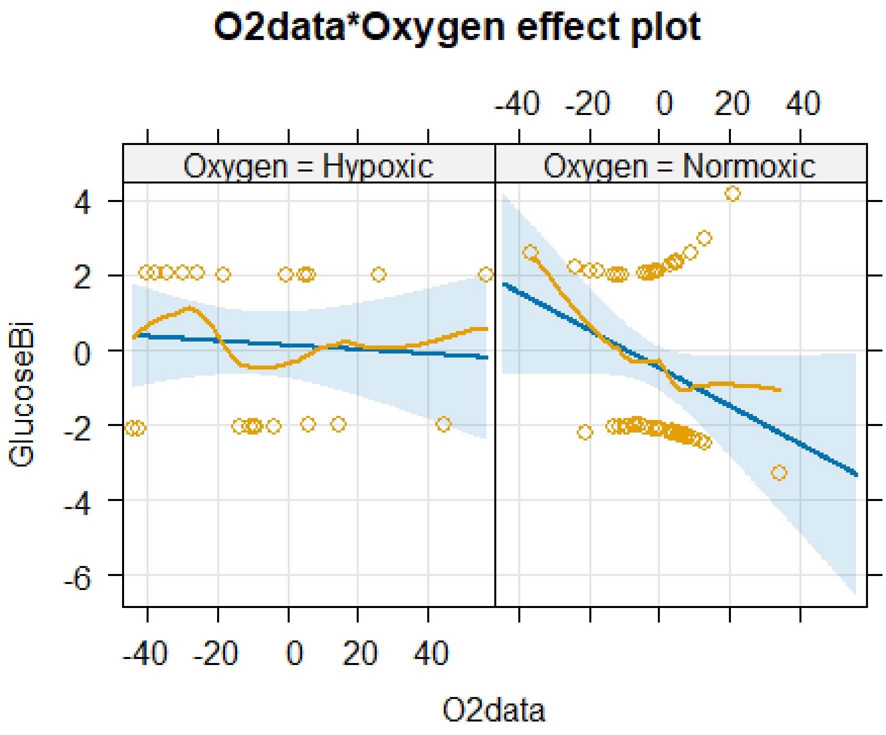


```
confint(glm1)

##                              2.5 %     97.5 %
## (Intercept)             -0.77223748 1.03715104
## O2data                  -0.03868886 0.02578670
## OxygenNormoxic          -1.68146170 0.45523051
## O2data:OxygenNormoxic -0.11198035 0.01599515

glm2 <- glm(GlucoseBi ~ O2data+Oxygen, data=TICsetOAO, family = "binomial")

summary(glm2)

##
## Call:
## glm(formula = GlucoseBi ~ O2data + Oxygen, family = "binomial",
##     data = TICsetOAO)
##
## Coefficients:
##                Estimate Std. Error z value Pr(>|z|)
## (Intercept)     0.02124    0.45809   0.046    0.963
## O2data         -0.01813    0.01395  -1.300    0.194
## OxygenNormoxic -0.45699    0.53495  -0.854    0.393
##
## (Dispersion parameter for binomial family taken to be 1)
##
##     Null deviance: 102.889  on 74  degrees of freedom
## Residual deviance:  99.704  on 72  degrees of freedom
```

```
## AIC: 105.7
##
## Number of Fisher Scoring iterations: 4
```

```
plot(allEffects(glm2), type = "response", grid = T)
```

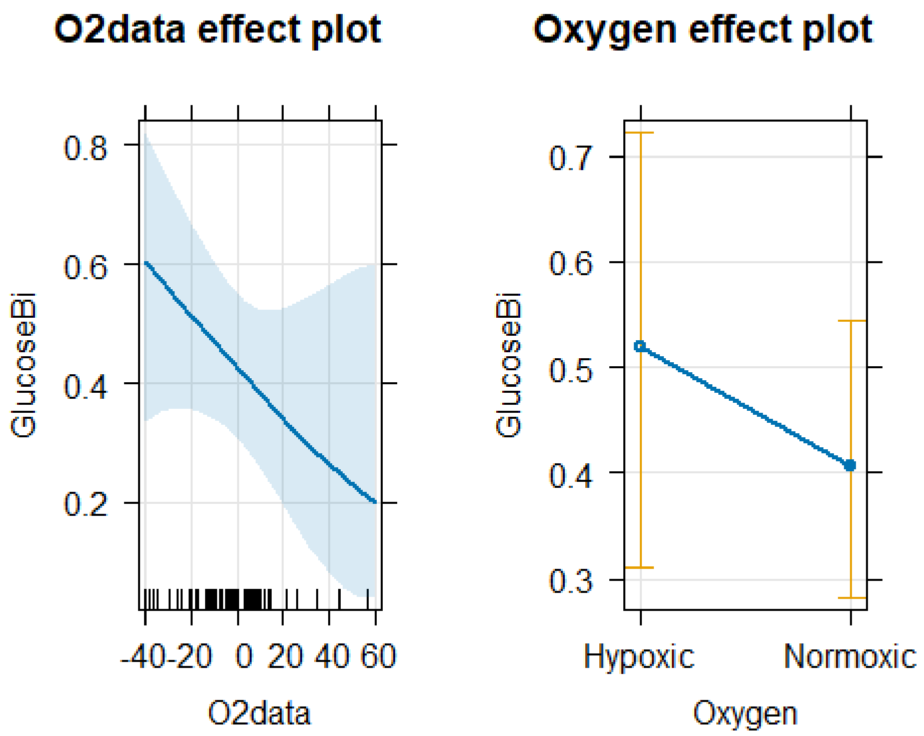


```
plot(allEffects(glm2, residuals=T), type = "link", grid = T)
```

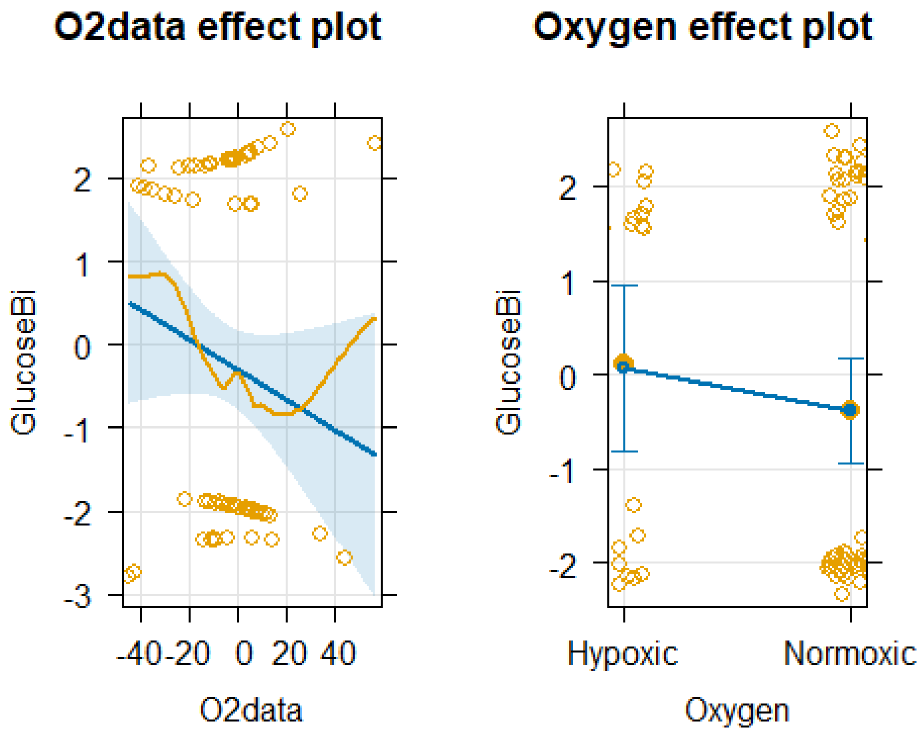


```
confint(glm2)

##                         2.5 %      97.5 %
## (Intercept)     -0.90358135 0.925417167
## O2data          -0.04749893 0.008302312
## OxygenNormoxic -1.51282175 0.606061479
```

Bovine data set

```
library(readxl)

TICsetO <- read_excel("BovineO2data.xlsx")
dim(TICsetO)

## [1] 80 25

tally(Oxygen~Donor, data=TICsetO)

##             Donor
## Oxygen      4 5 6 7 8 9 10 11 12 13
##   Hypoxic   8 8 0 0 8 0  0  0  0  0
##   Normoxic 0 0 8 8 0 8  8  8  8  8

 TICsetO <- TICsetO %>% drop_na(O2data) # Remove samples where O2 reading
could not be obtained
 dim(TICsetO)

## [1] 78 25
```

Bovine Glutamine Due to curvature in the oxygen consumption data a quadratic transformation is done in order to meet the linearity assumption.

```
G3<- TICsetO %>%
  ggplot(aes(x=O2data, fill=GlutamineBi))  +facet_wrap(~Oxygen)+
  geom_density(position='fill',bw=5)+ scale_fill_colorblind()+
geom_rug(aes(col=GlutamineBi),alpha=0.1) + scale_color_colorblind()

G3
```

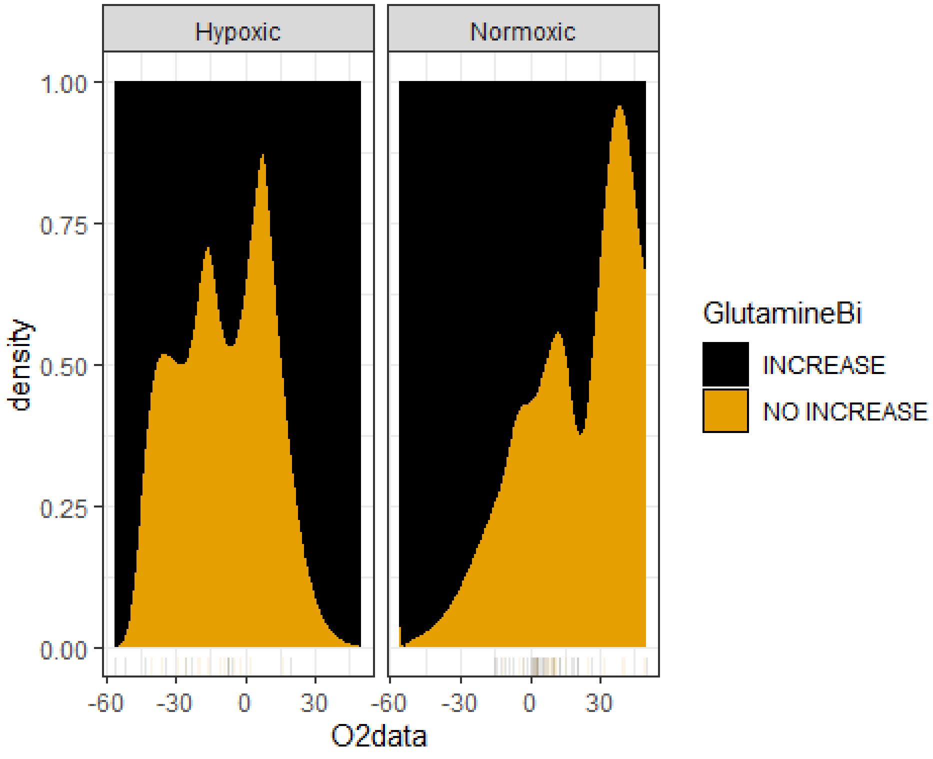


```
D3<- TICsetO %>%
  ggplot() + scale_fill_paletteer_d("nationalparkcolors::Acadia")  +
  geom_mosaic(aes(x=product(Donor),fill=GlutamineBi), offset=0.02)
D3
```

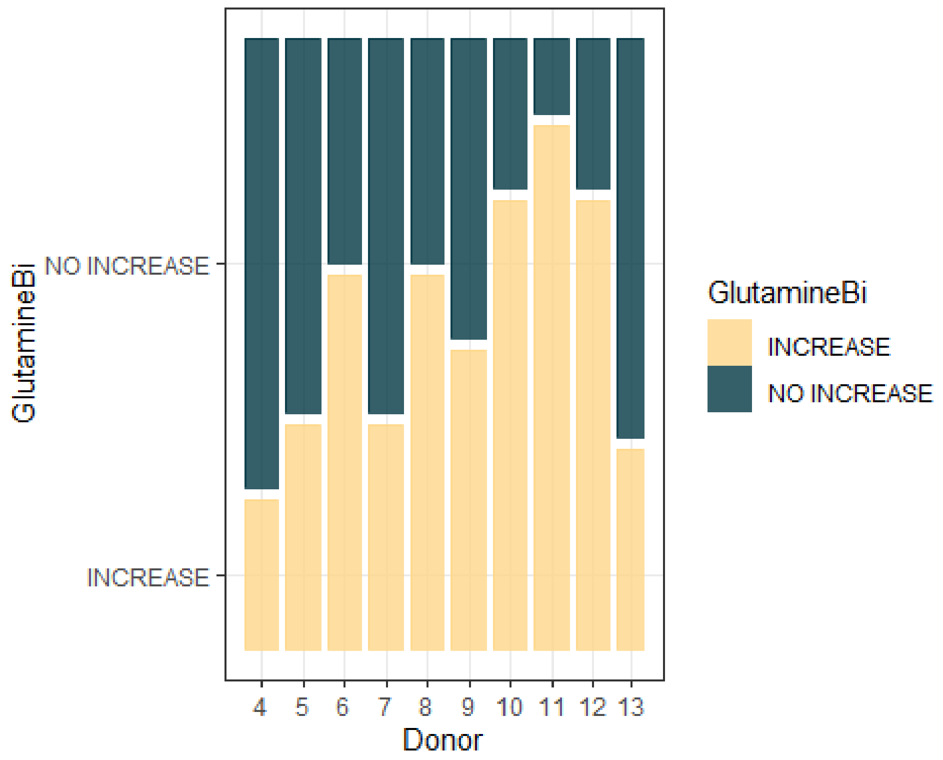


```r
TICsetO <- TICsetO %>% mutate(GlutamineBi = factor(GlutamineBi),
                                  Timef=factor(Time),
                                  Donor=factor(Donor))
glm1 <- glm(GlutamineBi ~ poly(O2data,2)*Oxygen, data=TICsetO, family =
"binomial") #Donor could not be included in the model due to the pattern of
binary responses creating separation
Anova(glm1)
```

```
## Analysis of Deviance Table (Type II tests)
##
## Response: GlutamineBi
##                          LR Chisq Df Pr(>Chisq)
## poly(O2data, 2)            4.5431  2    0.10315
## Oxygen                     6.6293  1    0.01003
## poly(O2data, 2):Oxygen     2.6055  2    0.27178
```

```r
summary(glm1)
```

```
##
## Call:
## glm(formula = GlutamineBi ~ poly(O2data, 2) * Oxygen, family = "binomial",
##     data = TICsetO)
##
## Coefficients:
##                                   Estimate Std. Error z value Pr(>|z|)
## (Intercept)                         0.1860     0.7170   0.259    0.795
## poly(O2data, 2)1                   -4.3742     6.7757  -0.646    0.519
```

```
## poly(O2data, 2)2                  -9.9638     6.6155  -1.506    0.132
## OxygenNormoxic                    -1.0653     0.9071  -1.174    0.240
## poly(O2data, 2)1:OxygenNormoxic  12.5499    10.3397   1.214    0.225
## poly(O2data, 2)2:OxygenNormoxic   8.2984     8.9153   0.931    0.352
##
## (Dispersion parameter for binomial family taken to be 1)
##
##     Null deviance: 107.309  on 77  degrees of freedom
## Residual deviance:  97.624  on 72  degrees of freedom
## AIC: 109.62
##
## Number of Fisher Scoring iterations: 4
```

```
plot(allEffects(glm1), type = "response", grid = T)
```

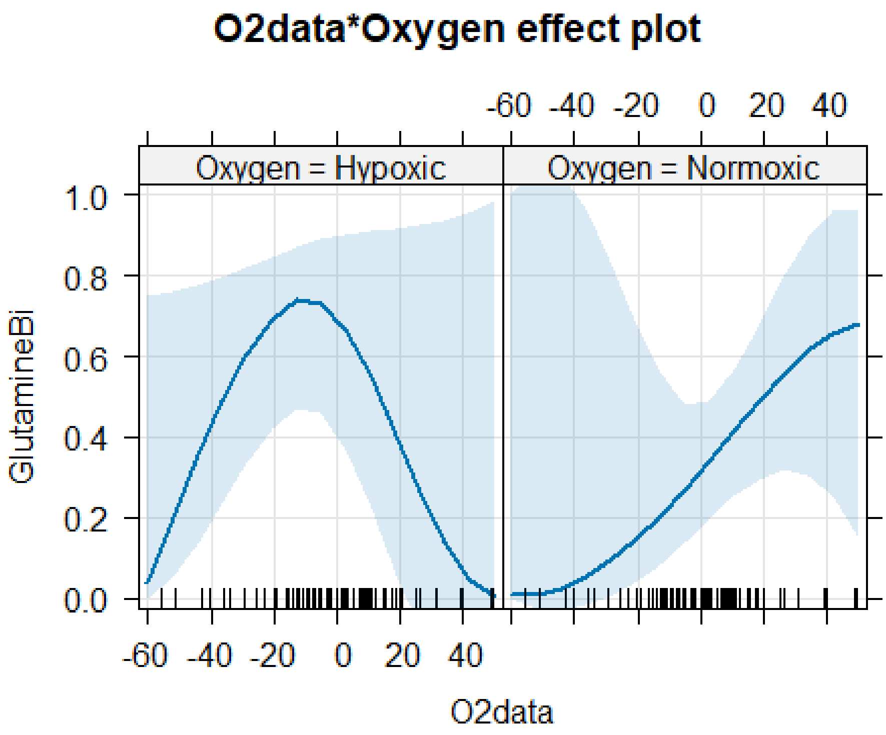


```
plot(allEffects(glm1, residuals=T), type = "link", grid = T)
```

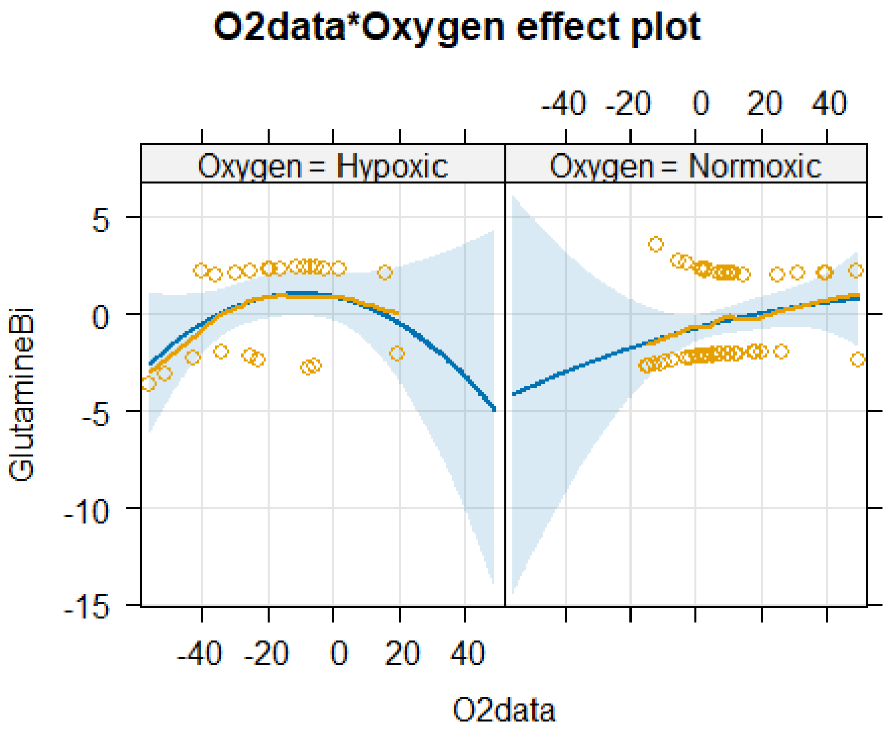


```r
confint(glm1)

##                                           2.5 %     97.5 %
## (Intercept)                           -1.418842  1.6701648
## poly(O2data, 2)1                     -19.771559  9.4037683
## poly(O2data, 2)2                     -25.194564  2.0128585
## OxygenNormoxic                        -2.923560  0.8098893
## poly(O2data, 2)1:OxygenNormoxic       -7.193235 34.4774299
## poly(O2data, 2)2:OxygenNormoxic       -8.779803 27.1139985

glm2 <- glm(GlutamineBi ~ poly(O2data,2)+Oxygen, data=TICsetO, family =
"binomial")
Anova(glm2)

## Analysis of Deviance Table (Type II tests)
##
## Response: GlutamineBi
##                 LR Chisq Df Pr(>Chisq)
## poly(O2data, 2)   4.5431  2    0.10315
## Oxygen            6.6293  1    0.01003

summary(glm2)

##
## Call:
## glm(formula = GlutamineBi ~ poly(O2data, 2) + Oxygen, family = "binomial",
##     data = TICsetO)
##
```

```
## Coefficients:
##                    Estimate Std. Error z value Pr(>|z|)
## (Intercept)          0.9998     0.5558   1.799   0.0721
## poly(O2data, 2)1     5.7927     2.8548   2.029   0.0424
## poly(O2data, 2)2    -0.9838     2.2304  -0.441   0.6592
## OxygenNormoxic      -1.7478     0.7199  -2.428   0.0152
##
## (Dispersion parameter for binomial family taken to be 1)
##
##     Null deviance: 107.31  on 77  degrees of freedom
## Residual deviance: 100.23  on 74  degrees of freedom
## AIC: 108.23
##
## Number of Fisher Scoring iterations: 4
```

```r
plot(allEffects(glm2), type = "response", grid = T)
```

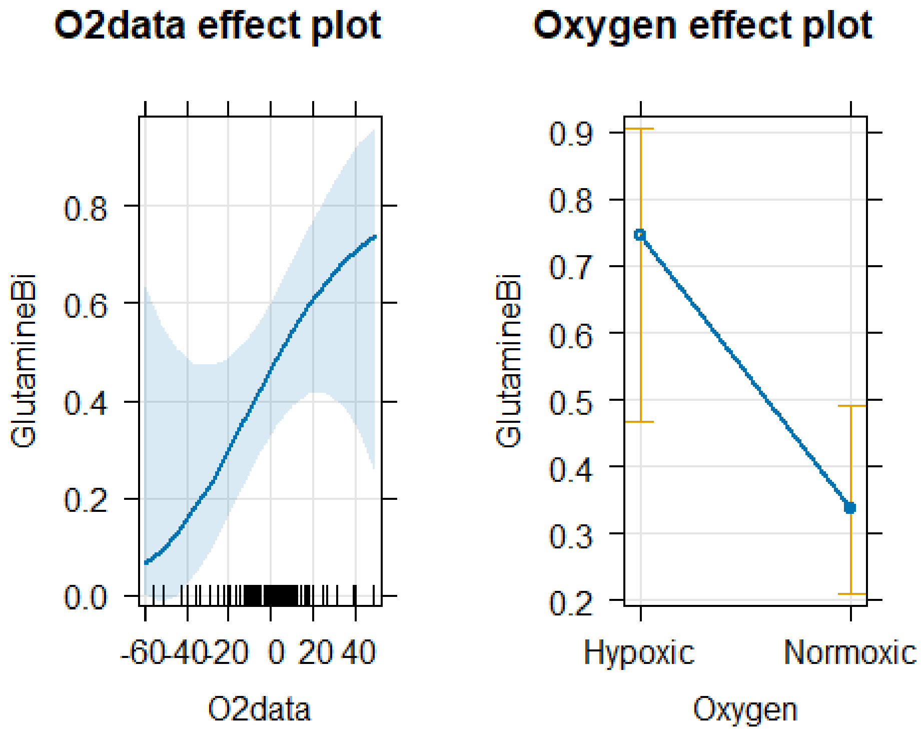


```r
plot(allEffects(glm2, residuals=T), type = "link", grid = T)
```

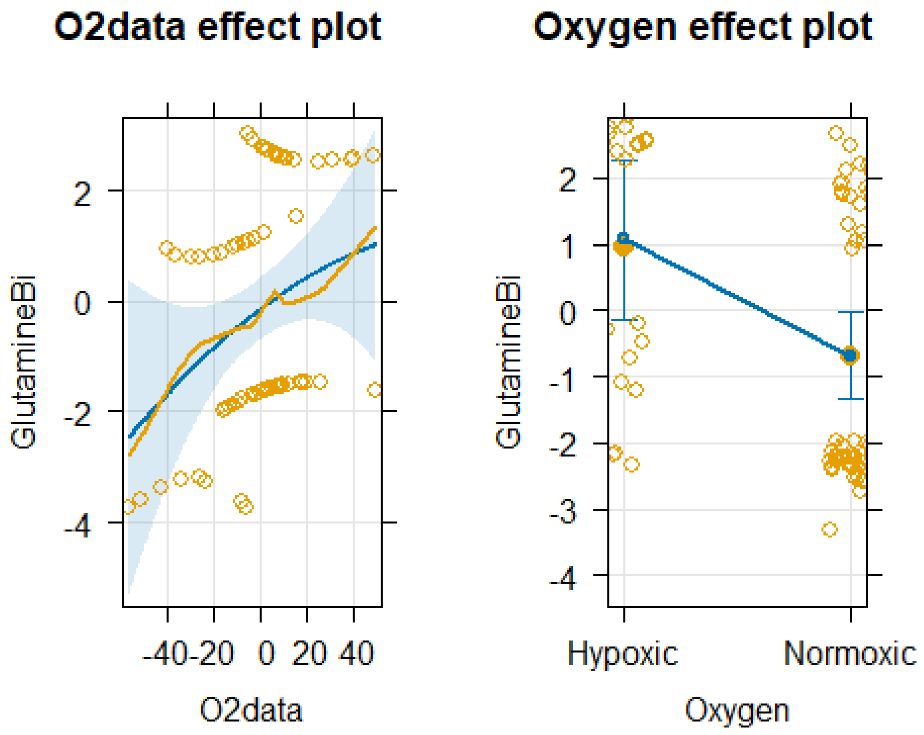


```
confint(glm2)

##                              2.5 %     97.5 %
## (Intercept)           -0.03544524  2.1822965
## poly(O2data, 2)1       0.44760440 11.8194297
## poly(O2data, 2)2      -5.64560816  3.3729529
## OxygenNormoxic        -3.26572432 -0.4039885

pdf("Bovine_Glutamine_Oxygen.pdf",width=6, height=4)
G6<- TICsetO %>%
  ggplot(aes(x=O2data, fill=GlutamineBi))  +facet_wrap(~Oxygen)+
  geom_density(position='fill',bw=5)+ scale_fill_colorblind()+
geom_rug(aes(col=GlutamineBi),alpha=0.1) + scale_color_colorblind()

G6

D3<- TICsetO %>%
  ggplot() + scale_fill_paletteer_d("nationalparkcolors::Acadia")  +
  geom_mosaic(aes(x=product(Oxygen),fill=GlutamineBi), offset=0.02)
D3
```

Bovine Glucose For this model there is strong evidence of an interaction between the quadratic of the Oxygen consumption and the oxygen environment therefor the interaction is left in the final model.

```
G4<- TICsetO %>%
  ggplot(aes(x=O2data, fill=GlucoseBi))  + facet_wrap(~Oxygen)+
```

```
  geom_density(position='fill',bw=5)+ scale_fill_colorblind()+
geom_rug(aes(col=GlucoseBi),alpha=0.1) + scale_color_colorblind()

G4
```

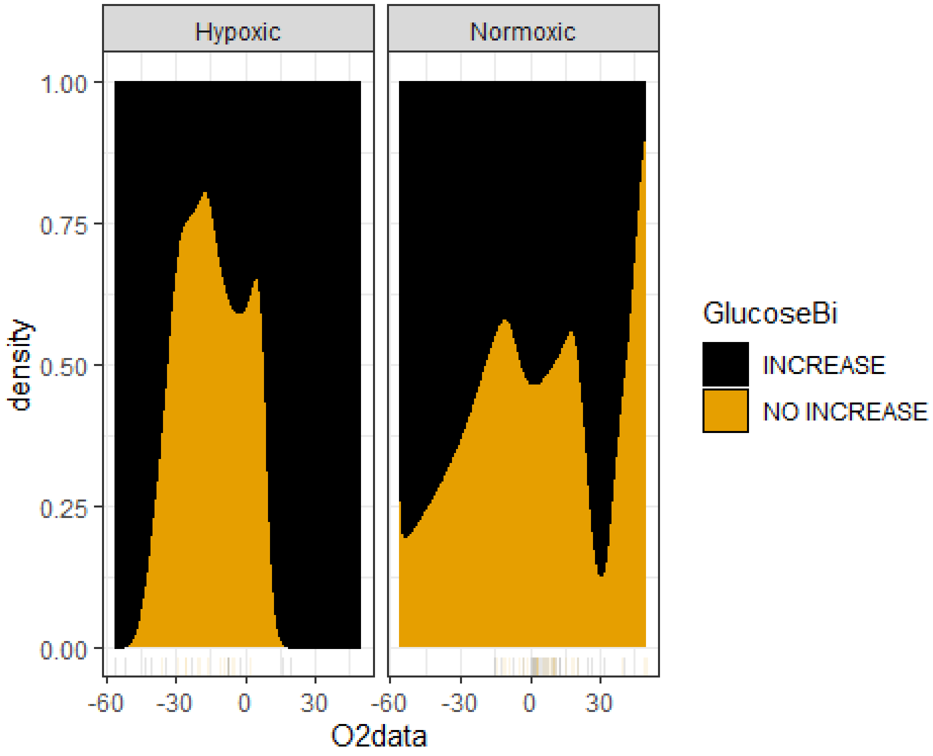


```
D4<- TICsetO %>%
  ggplot() + scale_fill_paletteer_d("nationalparkcolors::Acadia")  +
  geom_mosaic(aes(x=product(Donor),fill=GlucoseBi), offset=0.02)
D4
```

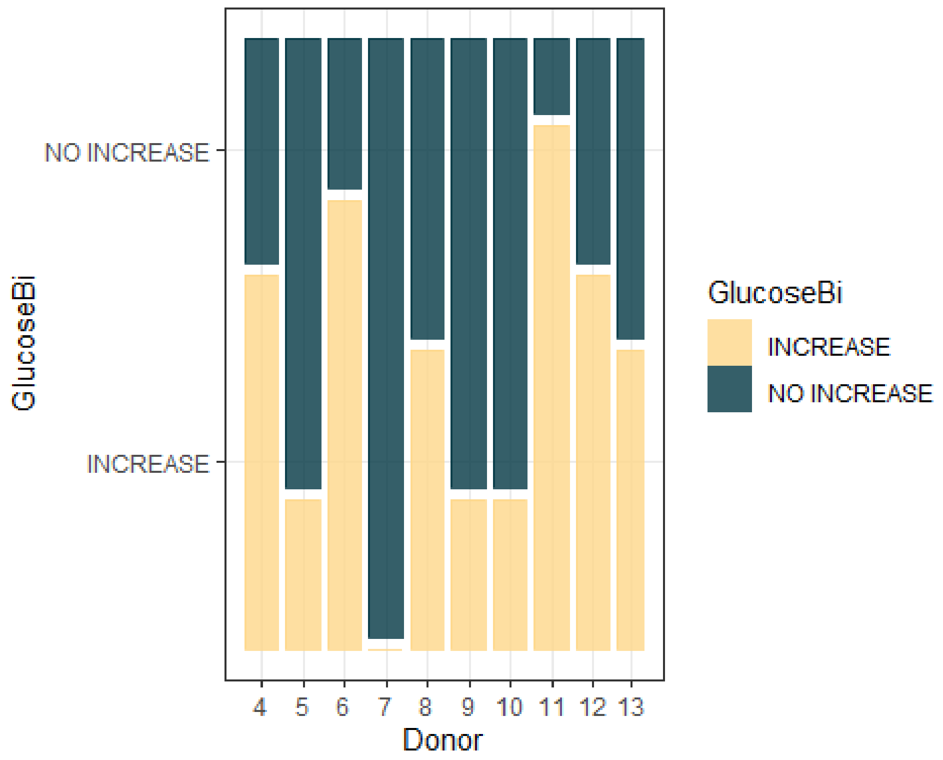


```
TICsetO <- TICsetO %>% mutate(GlucoseBi = factor(GlucoseBi),
                                  Timef=factor(Time),
                                  Donor=factor(Donor))
glm1 <- glm(GlucoseBi ~ poly(O2data,2)*Oxygen, data=TICsetO, family =
"binomial") #Donor could not be included in the model due to the pattern of
binary responses creating separation

Anova(glm1)

## Analysis of Deviance Table (Type II tests)
##
## Response: GlucoseBi
##                        LR Chisq Df Pr(>Chisq)
## poly(O2data, 2)          1.3759  2   0.502595
## Oxygen                   0.3598  1   0.548640
## poly(O2data, 2):Oxygen  10.1695  2   0.006191

summary(glm1)

##
## Call:
## glm(formula = GlucoseBi ~ poly(O2data, 2) * Oxygen, family = "binomial",
##     data = TICsetO)
##
## Coefficients:
##                                   Estimate Std. Error z value Pr(>|z|)
## (Intercept)                         -2.201      2.046  -1.076   0.2820
```

```
## poly(O2data, 2)1                    -27.803     19.615  -1.417   0.1564
## poly(O2data, 2)2                    -32.596     18.419  -1.770   0.0768
## OxygenNormoxic                        2.647      2.103   1.258   0.2082
## poly(O2data, 2)1:OxygenNormoxic      23.002     20.806   1.106   0.2689
## poly(O2data, 2)2:OxygenNormoxic      37.464     19.259   1.945   0.0517
##
## (Dispersion parameter for binomial family taken to be 1)
##
##     Null deviance: 107.669  on 77  degrees of freedom
## Residual deviance:  96.122  on 72  degrees of freedom
## AIC: 108.12
##
## Number of Fisher Scoring iterations: 6
```

```r
plot(allEffects(glm1), type = "response", grid = T)
```

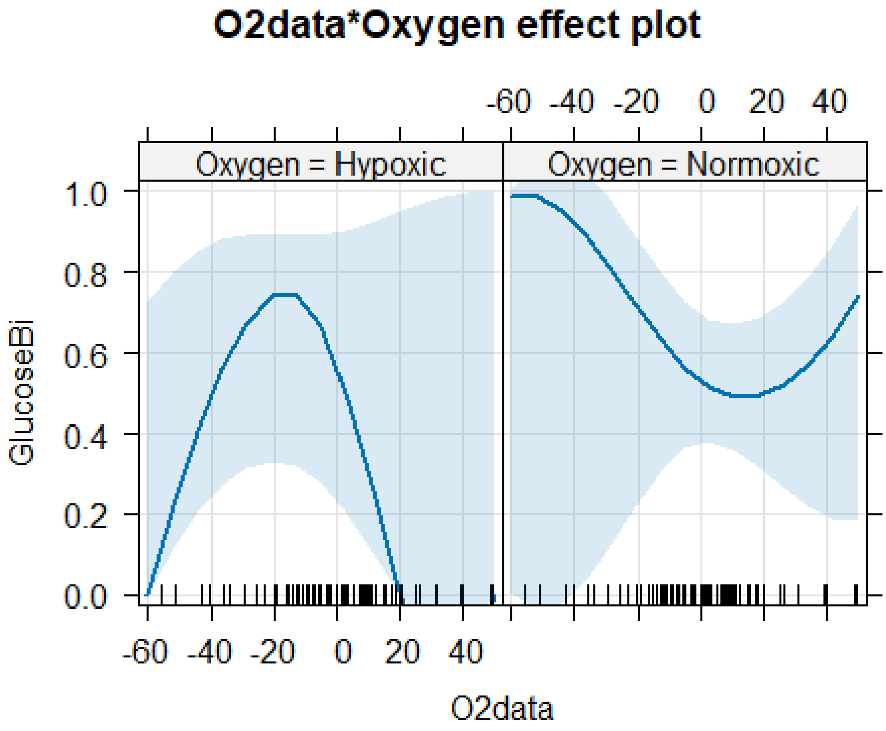


```r
plot(allEffects(glm1, residuals=T), type = "link", grid = T)
```

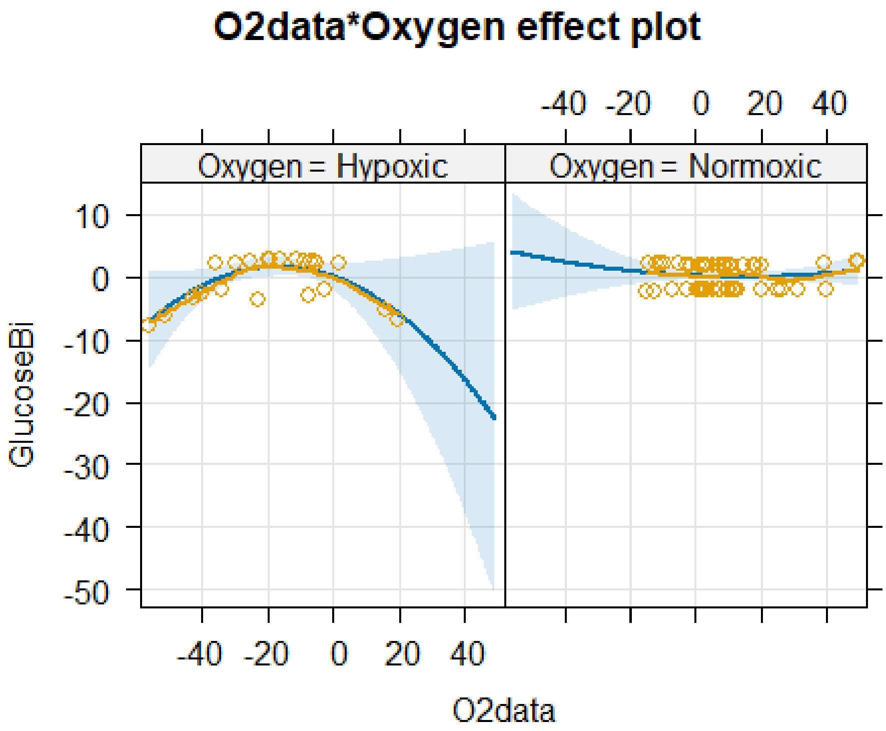


```
confint(glm1)

##                                       2.5 %     97.5 %
## (Intercept)                      -7.2756750  0.412948
## poly(O2data, 2)1                -76.5559846 -2.876844
## poly(O2data, 2)2                -79.5573990 -8.766040
## OxygenNormoxic                   -0.2044453  7.796778
## poly(O2data, 2)1:OxygenNormoxic  -6.9122955 73.288712
## poly(O2data, 2)2:OxygenNormoxic  10.3606218 85.513154
```

### TIC_Testing

Molly Piazza

2025-10-13

The following were all run to screen the metabolites for signs of differences that could be weeded out with the more in depth tests that were done above.

```
library(readxl)
TICset <- read_excel("InjuryBovine_With_TIC.xlsx")
TICset <- TICset %>% mutate(Donor = factor(Donor))
dim(TICset)

## [1] 90 51
```

#### Possible model:

```
tally(Time ~ Label, data = TICset)

##     Label
## Time  C  I  N
##    0  10 10 10
##    1   0 10 10
##    4   0 10 10
##    24  0 10 10

TICset <- TICset %>% mutate(TimeF = factor(Time),
                            Label.Time = str_c(Label, TimeF))
```

#### Just Time 0 observations for three group comparison:

Group based comparisons

Glutamate

```
TICsetR2 <- TICset %>% dplyr::filter(TimeF == "0") %>%
  mutate(GlutemicAcidB = factor(GlutemicAcidB))

glm2 <- glm(GlutemicAcidB ~ Label, data=TICsetR2, family = "binomial")

#Can't add donor - causes separation here
Anova(glm2)

## Analysis of Deviance Table (Type II tests)
##
## Response: GlutemicAcidB
```

```
##       LR Chisq Df Pr(>Chisq)
## Label   2.0982  2     0.3502
```

```r
summary(glm2)
```

```
##
## Call:
## glm(formula = GlutemicAcidB ~ Label, family = "binomial", data = TICsetR2)
##
## Coefficients:
##             Estimate Std. Error z value Pr(>|z|)
## (Intercept)  -1.3863     0.7906  -1.754   0.0795 .
## LabelI        1.3863     1.0124   1.369   0.1709
## LabelN        0.9808     1.0206   0.961   0.3365
## ---
## Signif. codes:  0 '***' 0.001 '**' 0.01 '*' 0.05 '.' 0.1 ' ' 1
##
## (Dispersion parameter for binomial family taken to be 1)
##
##     Null deviance: 39.429  on 29  degrees of freedom
## Residual deviance: 37.331  on 27  degrees of freedom
## AIC: 43.331
##
## Number of Fisher Scoring iterations: 4
```

```r
plot(allEffects(glm2), type = "response", grid = T)
```

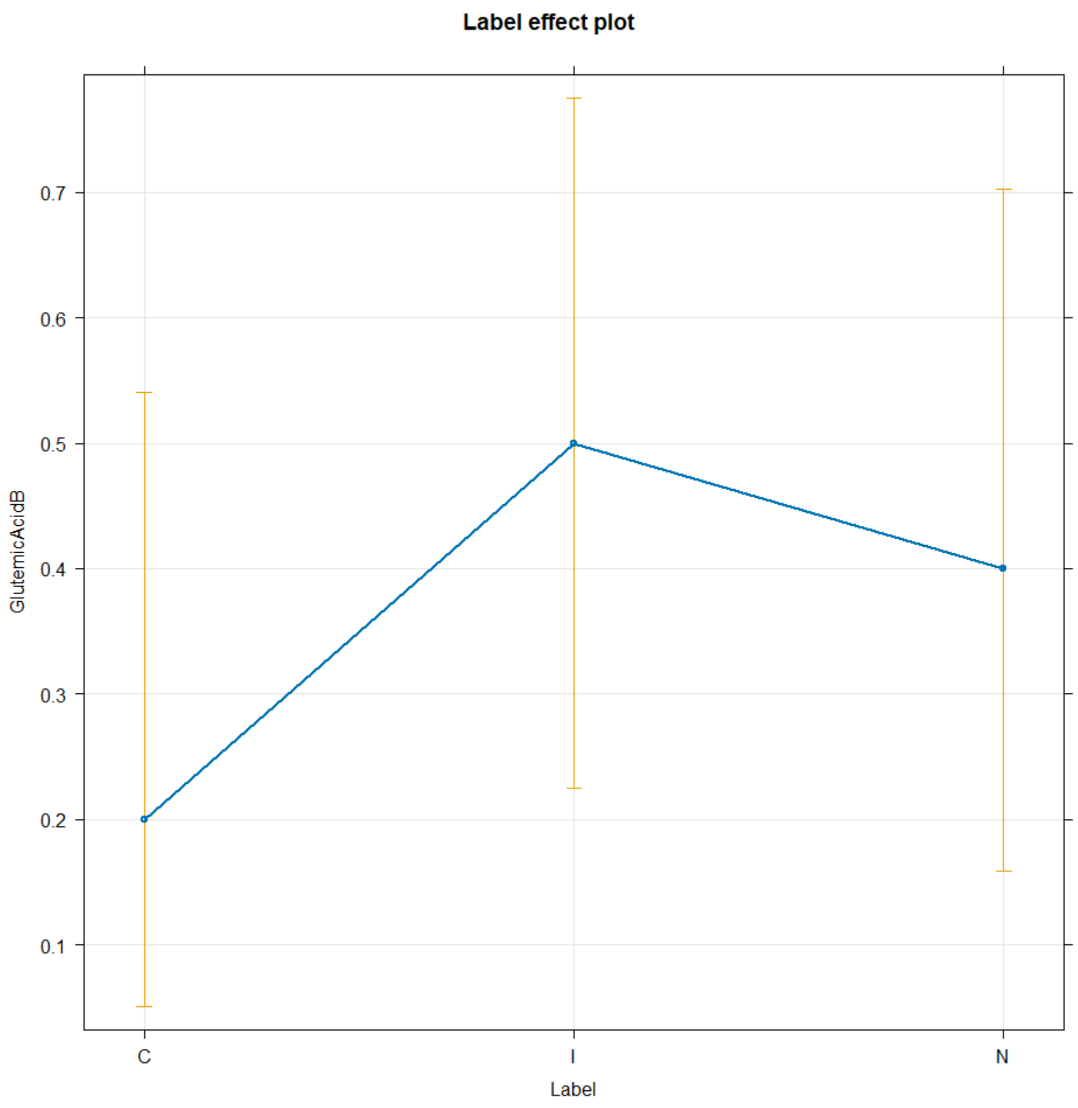


HSCOA

```
TICsetR2 <- TICset %>% dplyr::filter(TimeF == "0") %>%
  mutate(HSCOAB = factor(HSCOAB))

glm2 <- glm(HSCOAB ~ Label, data=TICsetR2, family = "binomial")

#Can't add donor - causes separation here
Anova(glm2)

## Analysis of Deviance Table (Type II tests)
##
## Response: HSCOAB
##       LR Chisq Df Pr(>Chisq)
## Label   2.7728  2       0.25

summary(glm2)
```

```
##
## Call:
## glm(formula = HSCOAB ~ Label, family = "binomial", data = TICsetR2)
##
## Coefficients:
##             Estimate Std. Error z value Pr(>|z|)
## (Intercept)   0.0000     0.6325   0.000    1.000
## LabelI       -1.3863     1.0124  -1.369    0.171
## LabelN       -1.3863     1.0124  -1.369    0.171
##
## (Dispersion parameter for binomial family taken to be 1)
##
##     Null deviance: 36.652  on 29  degrees of freedom
## Residual deviance: 33.879  on 27  degrees of freedom
## AIC: 39.879
##
## Number of Fisher Scoring iterations: 4
```

```
plot(allEffects(glm2), type = "response", grid = T)
```

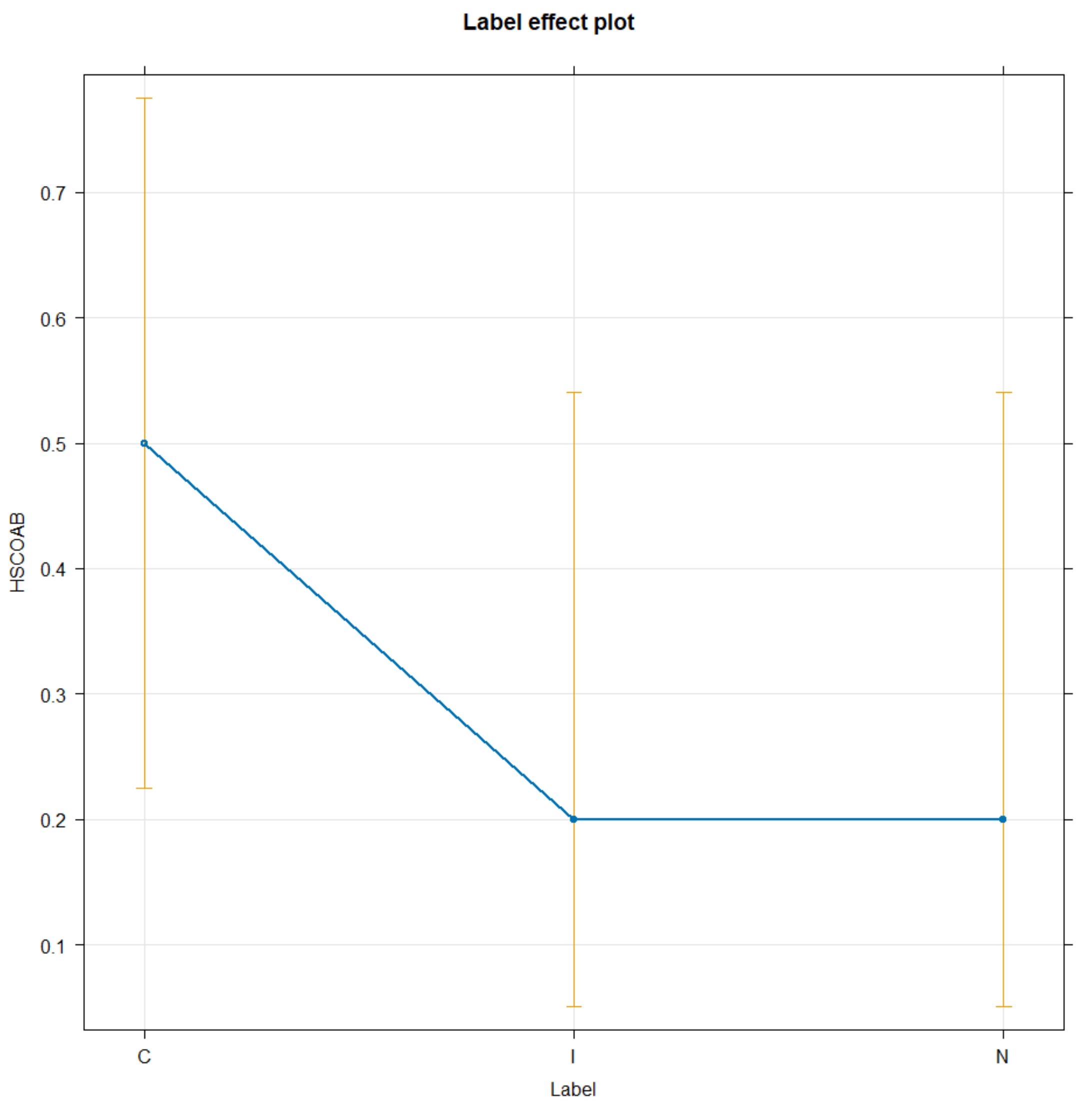


6-Phosphogluconate

```
TICsetR2 <- TICset %>% dplyr::filter(TimeF == "0") %>%
  mutate(SixPhosphoB = factor(SixPhosphoB))

glm2 <- glm(SixPhosphoB ~ Label, data=TICsetR2, family = "binomial")

#Can't add donor - causes separation here
Anova(glm2)

## Analysis of Deviance Table (Type II tests)
##
## Response: SixPhosphoB
##          LR Chisq Df Pr(>Chisq)
## Label -7.1054e-15  2          1

summary(glm2)
```

```
##
## Call:
## glm(formula = SixPhosphoB ~ Label, family = "binomial", data = TICsetR2)
##
## Coefficients:
##               Estimate Std. Error z value Pr(>|z|)
## (Intercept) -4.055e-01  6.455e-01  -0.628     0.53
## LabelI       5.266e-16  9.129e-01   0.000     1.00
## LabelN       0.000e+00  9.129e-01   0.000     1.00
##
## (Dispersion parameter for binomial family taken to be 1)
##
##     Null deviance: 40.381  on 29  degrees of freedom
## Residual deviance: 40.381  on 27  degrees of freedom
## AIC: 46.381
##
## Number of Fisher Scoring iterations: 4

plot(allEffects(glm2), type = "response", grid = T)
```

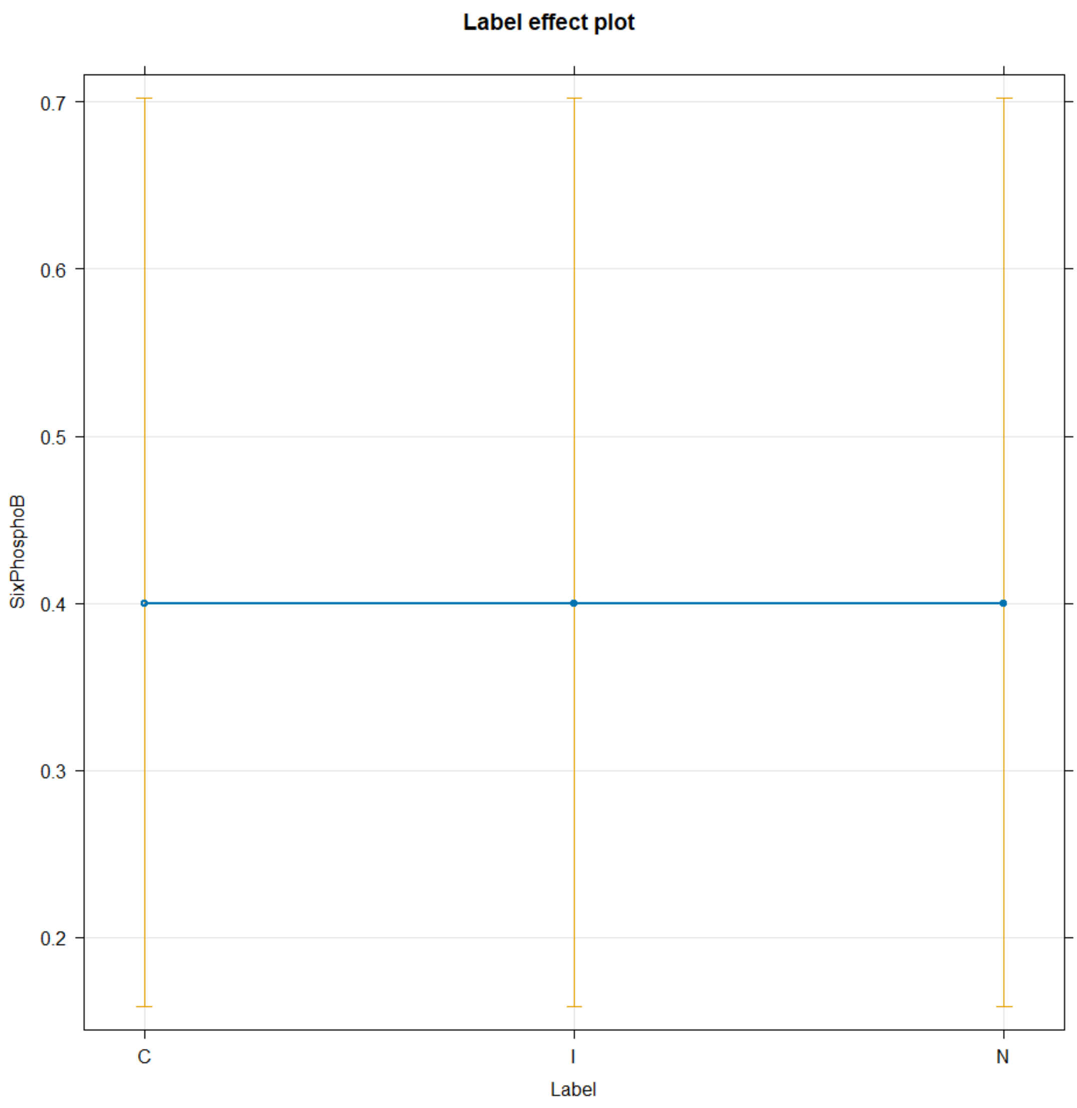


PEP

```r
TICsetR2 <- TICset %>% dplyr::filter(TimeF == "0") %>%
  mutate(PhosphoenolB = factor(PhosphoenolB))

glm2 <- glm(PhosphoenolB ~ Label, data=TICsetR2, family = "binomial")

#Can't add donor - causes separation here
Anova(glm2)

## Analysis of Deviance Table (Type II tests)
##
## Response: PhosphoenolB
##       LR Chisq Df Pr(>Chisq)
## Label   2.1026  2     0.3495

summary(glm2)
```

```
##
## Call:
## glm(formula = PhosphoenolB ~ Label, family = "binomial", data = TICsetR2)
##
## Coefficients:
##             Estimate Std. Error z value Pr(>|z|)
## (Intercept)   0.0000     0.6325   0.000    1.000
## LabelI       -0.8473     0.9361  -0.905    0.365
## LabelN       -1.3863     1.0124  -1.369    0.171
##
## (Dispersion parameter for binomial family taken to be 1)
##
##     Null deviance: 38.191  on 29  degrees of freedom
## Residual deviance: 36.088  on 27  degrees of freedom
## AIC: 42.088
##
## Number of Fisher Scoring iterations: 4
```

```
plot(allEffects(glm2), type = "response", grid = T)
```

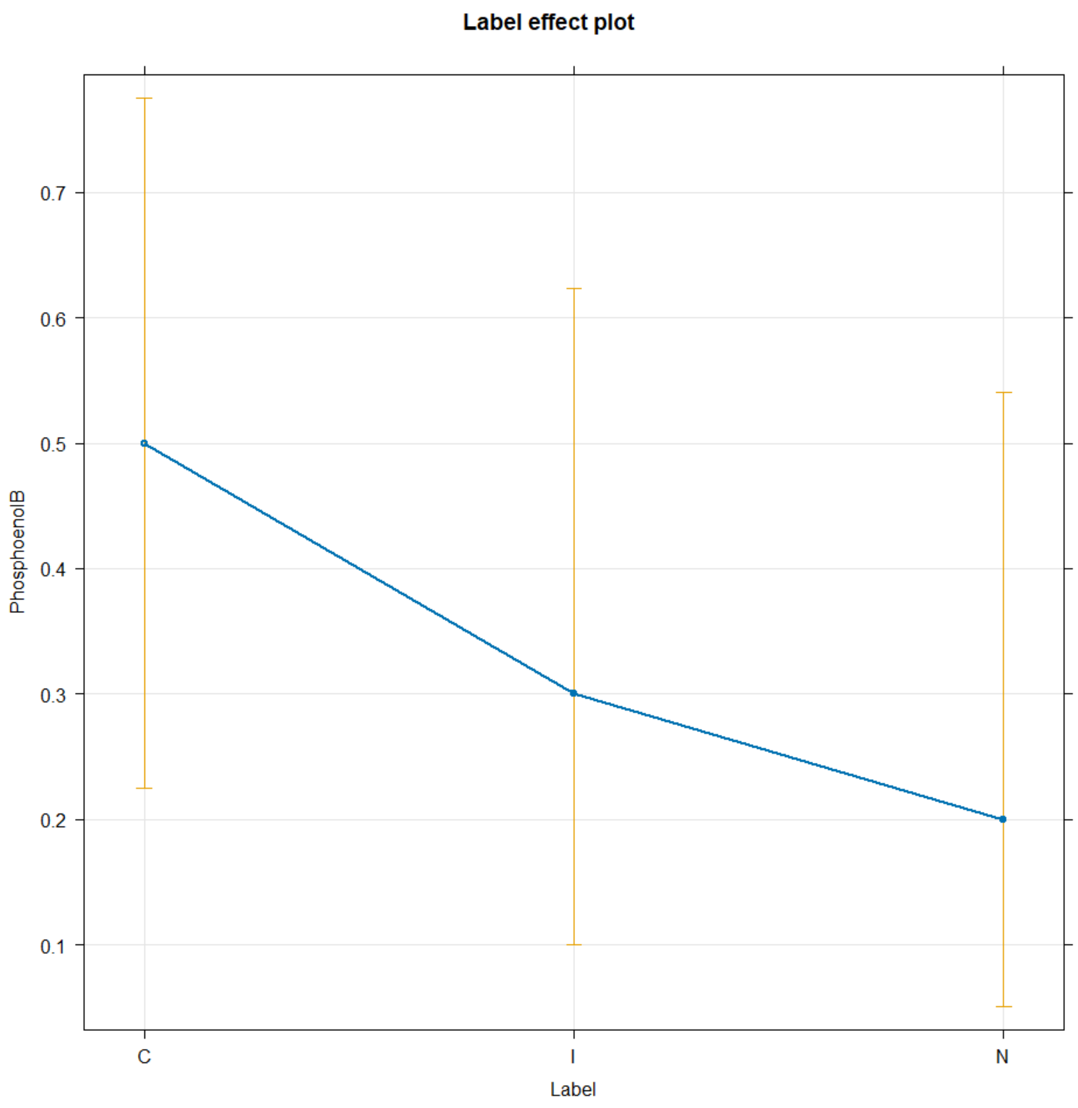


Pyruvic Acid

```
TICsetR2 <- TICset %>% dplyr::filter(TimeF == "0") %>%
  mutate(PyruvicB = factor(PyruvicB))

glm2 <- glm(PyruvicB ~ Label, data=TICsetR2, family = "binomial")

#Can't add donor - causes separation here
Anova(glm2)

## Analysis of Deviance Table (Type II tests)
##
## Response: PyruvicB
##       LR Chisq Df Pr(>Chisq)
## Label   7.2877  2    0.02615 *
## ---
## Signif. codes:  0 '***' 0.001 '**' 0.01 '*' 0.05 '.' 0.1 ' ' 1
```

```r
summary(glm2)

##
## Call:
## glm(formula = PyruvicB ~ Label, family = "binomial", data = TICsetR2)
##
## Coefficients:
##              Estimate Std. Error z value Pr(>|z|)
## (Intercept)    0.8473     0.6901   1.228    0.220
## LabelI        19.7188  5606.8353   0.004    0.997
## LabelN        19.7188  5606.8354   0.004    0.997
##
## (Dispersion parameter for binomial family taken to be 1)
##
##     Null deviance: 19.505  on 29  degrees of freedom
## Residual deviance: 12.217  on 27  degrees of freedom
## AIC: 18.217
##
## Number of Fisher Scoring iterations: 19

plot(allEffects(glm2), type = "response", grid = T)
```

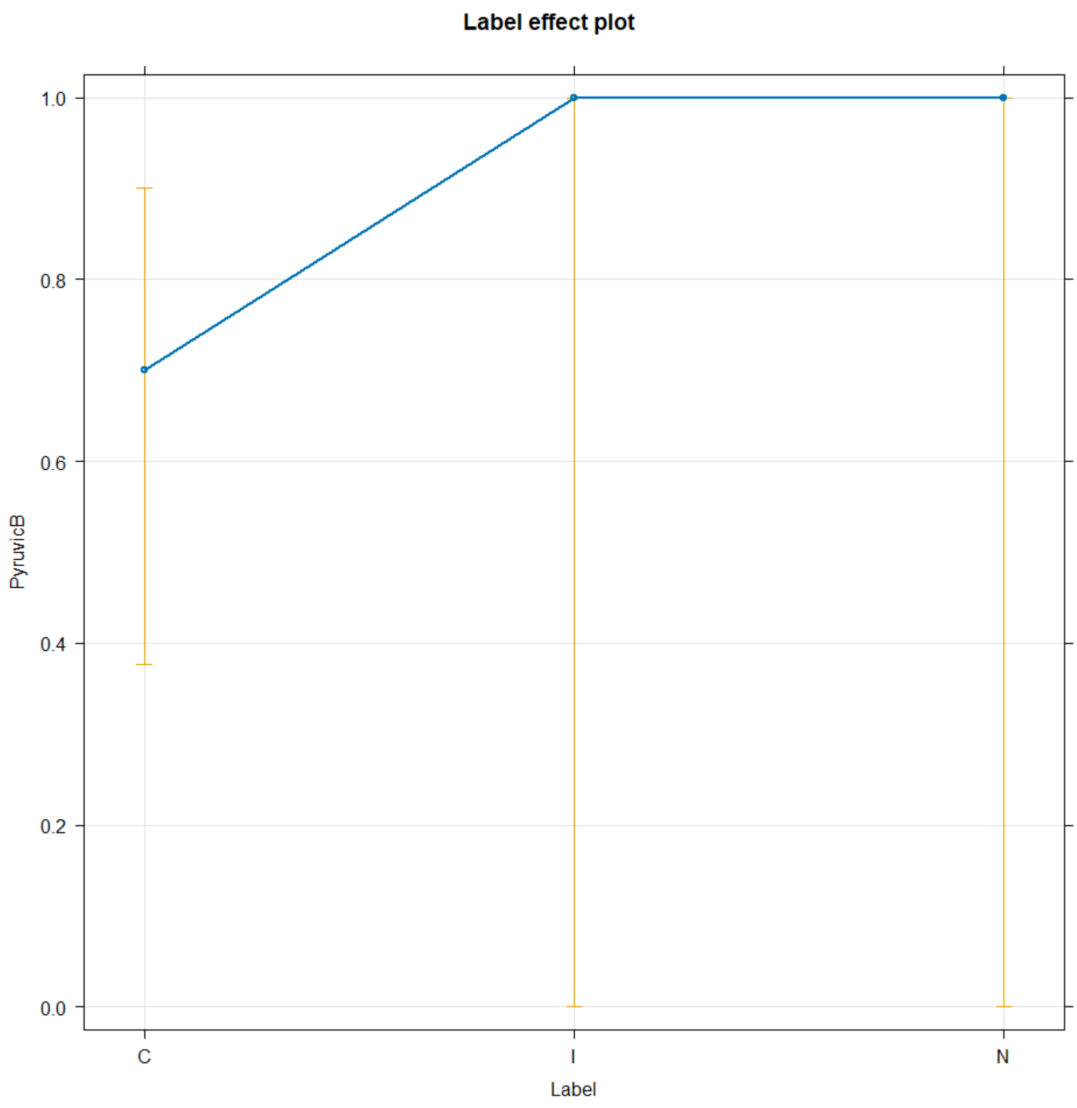


Group Comparison over 80%

Logged NADH

```
lm0 <- lm(NADHT ~ Label+Donor, data=TICsetR2)
Anova(lm0)

## Anova Table (Type II tests)
##
## Response: NADHT
##            Sum Sq Df F value Pr(>F)
## Label     0.24119  2  2.1575 0.1446
## Donor     0.25153  9  0.5000 0.8556
## Residuals 1.00612 18

resid_panel(lm0, "R")
```

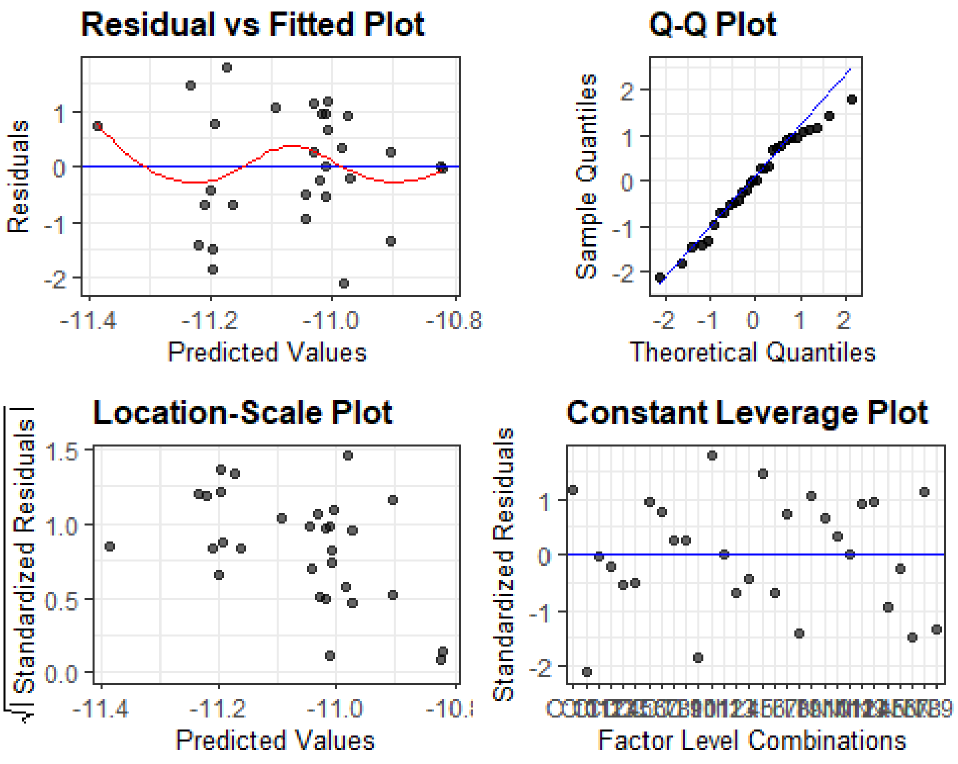


```
summary(lm0)

##
## Call:
## lm(formula = NADHT ~ Label + Donor, data = TICsetR2)
##
## Residuals:
##      Min       1Q   Median       3Q      Max
## -0.38811 -0.11881  0.00168  0.15899  0.32703
##
## Coefficients:
##               Estimate Std. Error t value Pr(>|t|)
## (Intercept) -11.009341   0.149527 -73.628   <2e-16 ***
## LabelI       -0.191051   0.105732  -1.807   0.0875 .
## LabelN       -0.001700   0.105732  -0.016   0.9874
## Donor5       -0.033530   0.193038  -0.174   0.8640
## Donor6       -0.008766   0.193038  -0.045   0.9643
## Donor7       -0.185287   0.193038  -0.960   0.3498
## Donor8       -0.020365   0.193038  -0.105   0.9171
## Donor9        0.104918   0.193038   0.544   0.5934
## Donor10       0.003240   0.193038   0.017   0.9868
## Donor11       0.026983   0.193038   0.140   0.8904
## Donor12       0.189963   0.193038   0.984   0.3381
## Donor13       0.036970   0.193038   0.192   0.8503
## ---
## Signif. codes:  0 '***' 0.001 '**' 0.01 '*' 0.05 '.' 0.1 ' ' 1
##
```

```
## Residual standard error: 0.2364 on 18 degrees of freedom
## Multiple R-squared:  0.3287, Adjusted R-squared:  -0.08148
## F-statistic: 0.8014 on 11 and 18 DF,  p-value: 0.6384
```

```
plot(allEffects(lm0), grid = T)
```

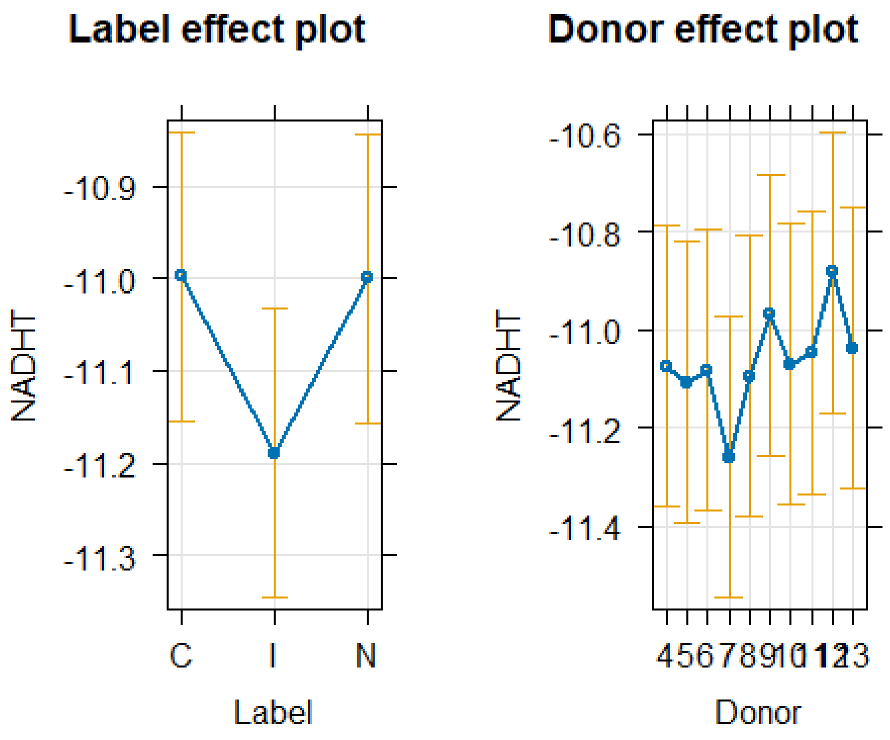


```
lm1 <- lm(NADHT ~ Label, data=TICsetR2)
Anova(lm1)
```

```
## Anova Table (Type II tests)
##
## Response: NADHT
##            Sum Sq Df F value  Pr(>F)
## Label     0.24119  2   2.589 0.09362 .
## Residuals 1.25766 27
## ---
## Signif. codes:  0 '***' 0.001 '**' 0.01 '*' 0.05 '.' 0.1 ' ' 1
```

```
resid_panel(lm1, "R")
```

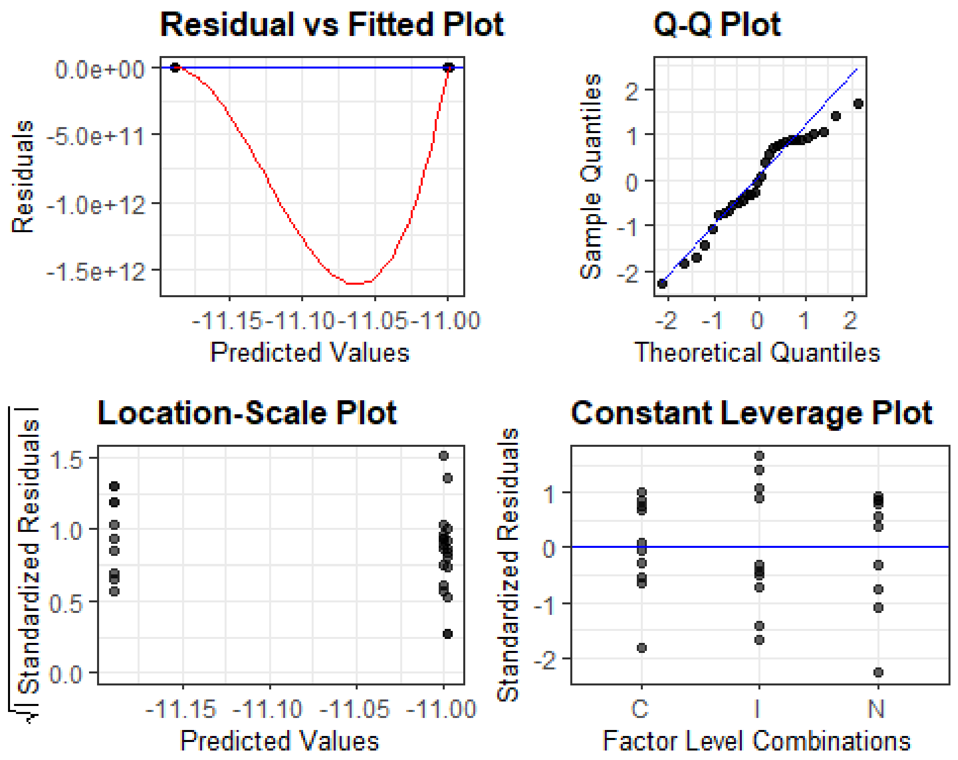


```
summary(lm1)

##
## Call:
## lm(formula = NADHT ~ Label, data = TICsetR2)
##
## Residuals:
##      Min       1Q   Median       3Q      Max
## -0.46706 -0.12837  0.00017  0.17690  0.34260
##
## Coefficients:
##             Estimate Std. Error  t value Pr(>|t|)
## (Intercept) -10.99793    0.06825 -161.143   <2e-16 ***
## LabelI       -0.19105    0.09652   -1.979    0.058 .
## LabelN       -0.00170    0.09652   -0.018    0.986
## ---
## Signif. codes:  0 '***' 0.001 '**' 0.01 '*' 0.05 '.' 0.1 ' ' 1
##
## Residual standard error: 0.2158 on 27 degrees of freedom
## Multiple R-squared:  0.1609, Adjusted R-squared:  0.09876
## F-statistic: 2.589 on 2 and 27 DF,  p-value: 0.09362

plot(allEffects(lm1), grid = T)
```

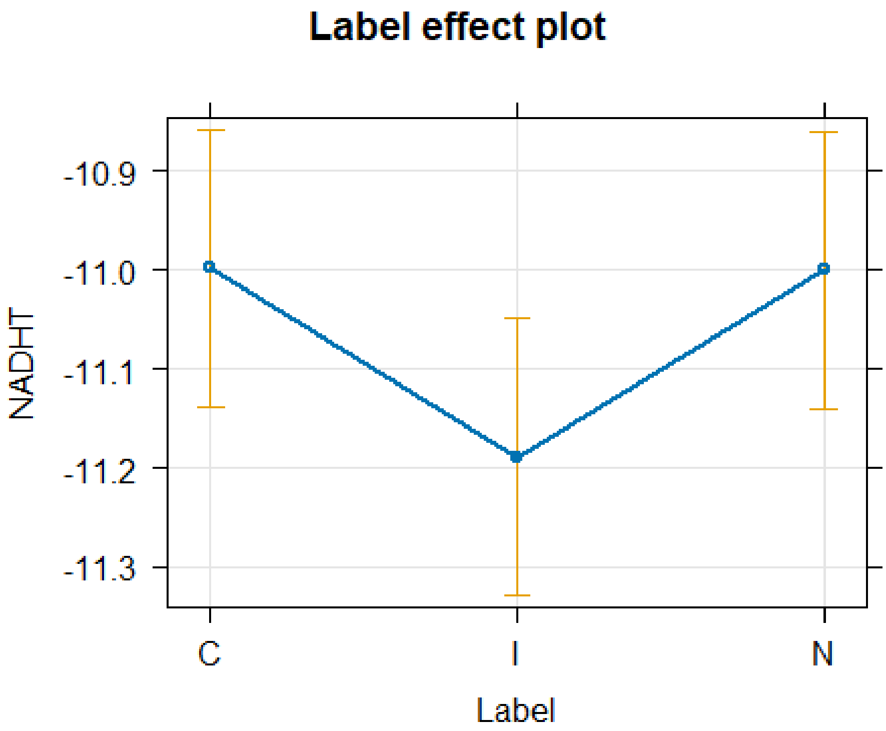


```
TukeyRes <- emmeans(lm1, pairwise ~ Label, adjust = "tukey")
TukeyRes

## $emmeans
##  Label emmean     SE df lower.CL upper.CL
##  C      -11.0 0.0682 27    -11.1    -10.9
##  I      -11.2 0.0682 27    -11.3    -11.0
##  N      -11.0 0.0682 27    -11.1    -10.9
##
## Confidence level used: 0.95
##
## $contrasts
##  contrast estimate     SE df t.ratio p.value
##  C - I      0.1911 0.0965 27   1.979  0.1367
##  C - N      0.0017 0.0965 27   0.018  0.9998
##  I - N     -0.1894 0.0965 27  -1.962  0.1412
##
## P value adjustment: tukey method for comparing a family of 3 estimates

confint(TukeyRes)

## $emmeans
##  Label emmean     SE df lower.CL upper.CL
##  C      -11.0 0.0682 27    -11.1    -10.9
##  I      -11.2 0.0682 27    -11.3    -11.0
##  N      -11.0 0.0682 27    -11.1    -10.9
##
```

```
## Confidence level used: 0.95
##
## $contrasts
##  contrast estimate     SE df lower.CL upper.CL
##  C - I      0.1911 0.0965 27  -0.0483    0.430
##  C - N      0.0017 0.0965 27  -0.2376    0.241
##  I - N     -0.1894 0.0965 27  -0.4287    0.050
##
## Confidence level used: 0.95
## Conf-level adjustment: tukey method for comparing a family of 3 estimates
```

```
plot(TukeyRes, comparison = T)
```

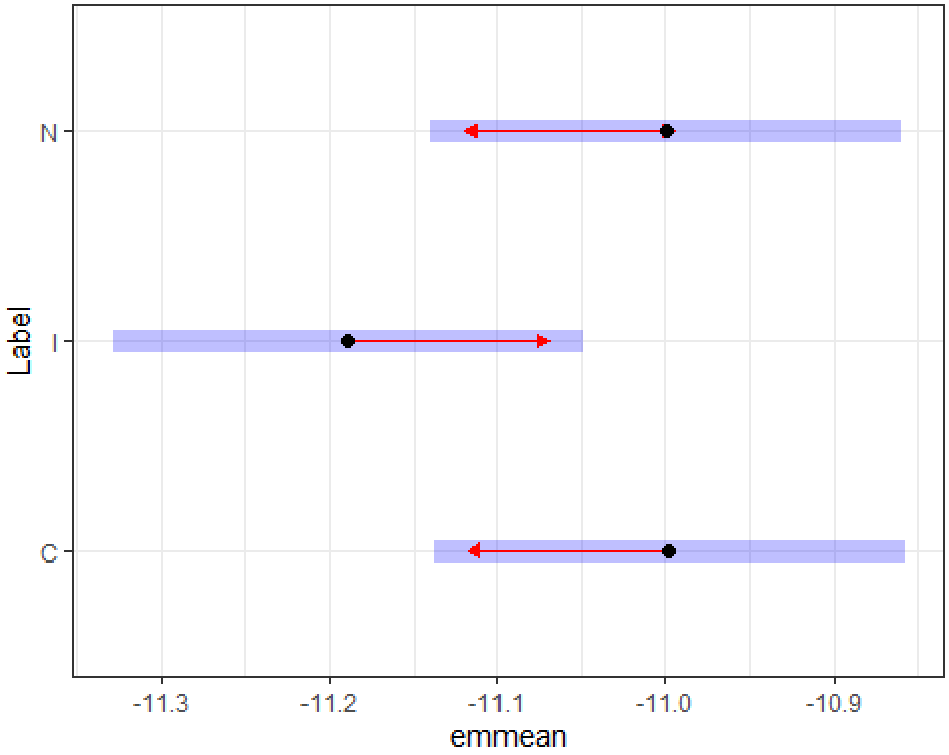


```
multcomp::cld(TukeyRes, Letters = LETTERS)
```

```
##  Label emmean     SE df lower.CL upper.CL .group
##  I      -11.2 0.0682 27    -11.3    -11.0  A
##  N      -11.0 0.0682 27    -11.1    -10.9  A
##  C      -11.0 0.0682 27    -11.1    -10.9  A
##
## Confidence level used: 0.95
## P value adjustment: tukey method for comparing a family of 3 estimates
## significance level used: alpha = 0.05
## NOTE: If two or more means share the same grouping symbol,
##       then we cannot show them to be different.
##       But we also did not show them to be the same.
```

Logged Lactic Acid

```
lm0 <- lm(LacticAcidT ~ Label+Donor, data=TICsetR2)
Anova(lm0)

## Anova Table (Type II tests)
##
## Response: LacticAcidT
##            Sum Sq Df F value Pr(>F)
## Label      0.7910  2  0.5138 0.6067
## Donor      7.4626  9  1.0773 0.4240
## Residuals 13.8545 18

resid_panel(lm0, "R")
```

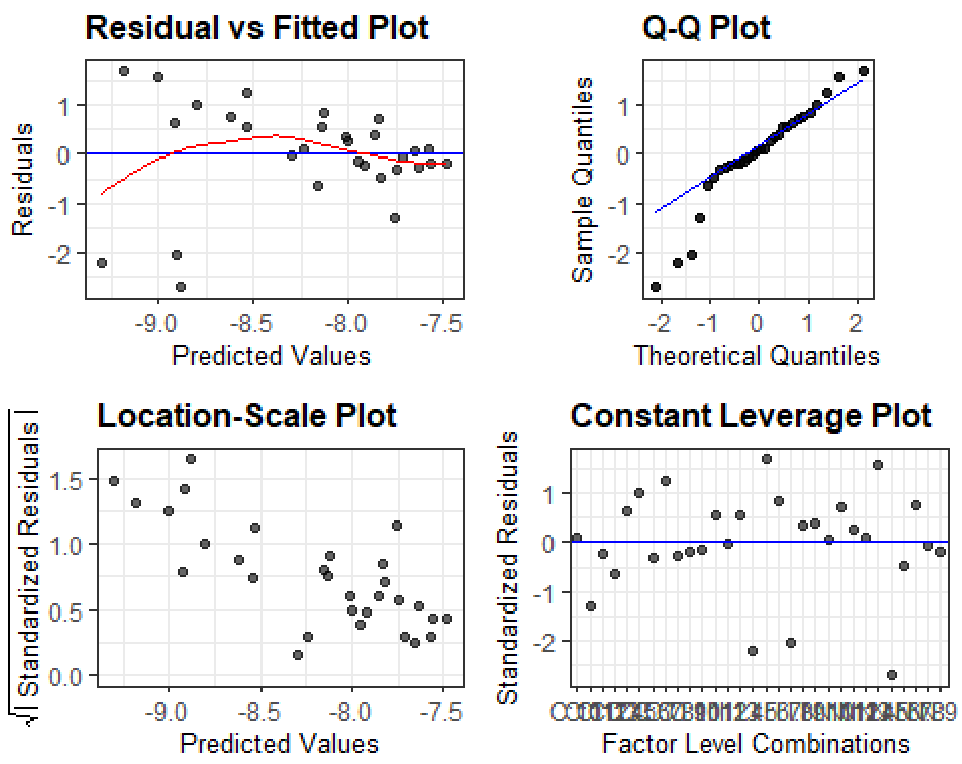


```
summary(lm0)

##
## Call:
## lm(formula = LacticAcidT ~ Label + Donor, data = TICsetR2)
##
## Residuals:
##      Min       1Q   Median       3Q      Max
## -1.84215 -0.17709  0.05164  0.41014  1.15500
##
## Coefficients:
##             Estimate Std. Error t value Pr(>|t|)
## (Intercept)  -8.9218     0.5549 -16.079 4.01e-12 ***
```

```
## LabelI       -0.3775     0.3923  -0.962   0.3487
## LabelN       -0.0802     0.3923  -0.204   0.8403
## Donor5        0.1185     0.7163   0.165   0.8705
## Donor6        1.1732     0.7163   1.638   0.1188
## Donor7        0.3873     0.7163   0.541   0.5953
## Donor8        1.2885     0.7163   1.799   0.0889 .
## Donor9        1.4387     0.7163   2.008   0.0598 .
## Donor10       1.3477     0.7163   1.881   0.0762 .
## Donor11       1.1616     0.7163   1.622   0.1223
## Donor12       1.0033     0.7163   1.401   0.1783
## Donor13       0.7632     0.7163   1.065   0.3008
## ---
## Signif. codes:  0 '***' 0.001 '**' 0.01 '*' 0.05 '.' 0.1 ' ' 1
##
## Residual standard error: 0.8773 on 18 degrees of freedom
## Multiple R-squared:  0.3733, Adjusted R-squared:  -0.009638
## F-statistic: 0.9748 on 11 and 18 DF,  p-value: 0.5009
```

```
plot(allEffects(lm0), grid = T)
```

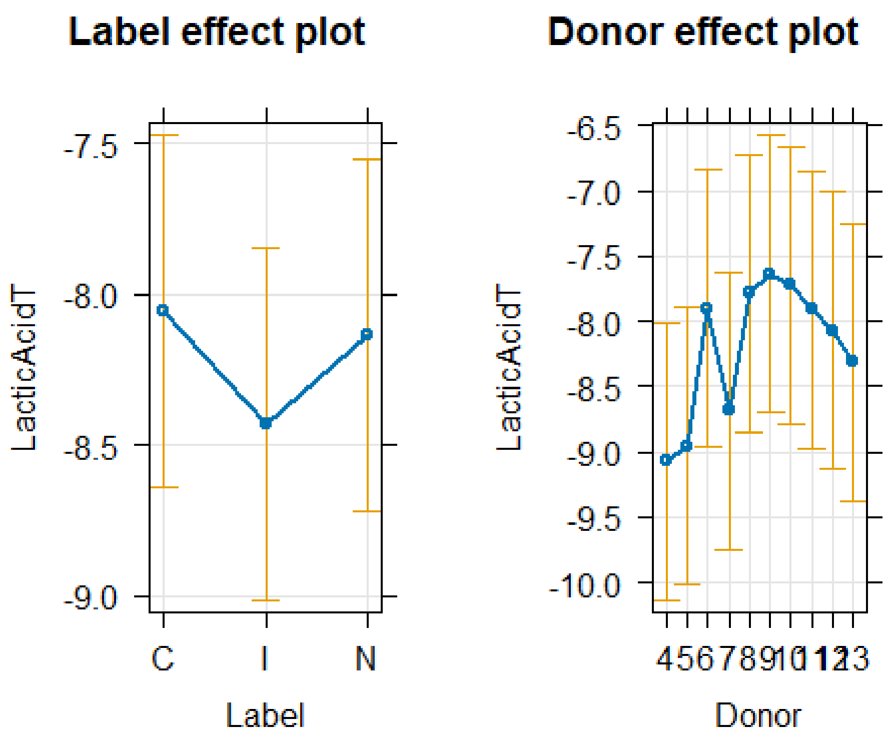


```
TukeyRes <- emmeans(lm0, pairwise ~ Label, adjust = "tukey")
TukeyRes
```

```
## $emmeans
##  Label emmean    SE df lower.CL upper.CL
##  C      -8.05 0.277 18    -8.64    -7.47
##  I      -8.43 0.277 18    -9.01    -7.85
```

```
##  N      -8.13 0.277 18    -8.72    -7.55
##
## Results are averaged over the levels of: Donor
## Confidence level used: 0.95
##
## $contrasts
##  contrast estimate    SE df t.ratio p.value
##  C - I      0.3775 0.392 18   0.962  0.6093
##  C - N      0.0802 0.392 18   0.204  0.9773
##  I - N     -0.2973 0.392 18  -0.758  0.7329
##
## Results are averaged over the levels of: Donor
## P value adjustment: tukey method for comparing a family of 3 estimates
```

```r
confint(TukeyRes)
```

```
## $emmeans
##  Label emmean    SE df lower.CL upper.CL
##  C      -8.05 0.277 18    -8.64    -7.47
##  I      -8.43 0.277 18    -9.01    -7.85
##  N      -8.13 0.277 18    -8.72    -7.55
##
## Results are averaged over the levels of: Donor
## Confidence level used: 0.95
##
## $contrasts
##  contrast estimate    SE df lower.CL upper.CL
##  C - I      0.3775 0.392 18   -0.624    1.379
##  C - N      0.0802 0.392 18   -0.921    1.082
##  I - N     -0.2973 0.392 18   -1.299    0.704
##
## Results are averaged over the levels of: Donor
## Confidence level used: 0.95
## Conf-level adjustment: tukey method for comparing a family of 3 estimates
```

```r
plot(TukeyRes, comparison = T)
```

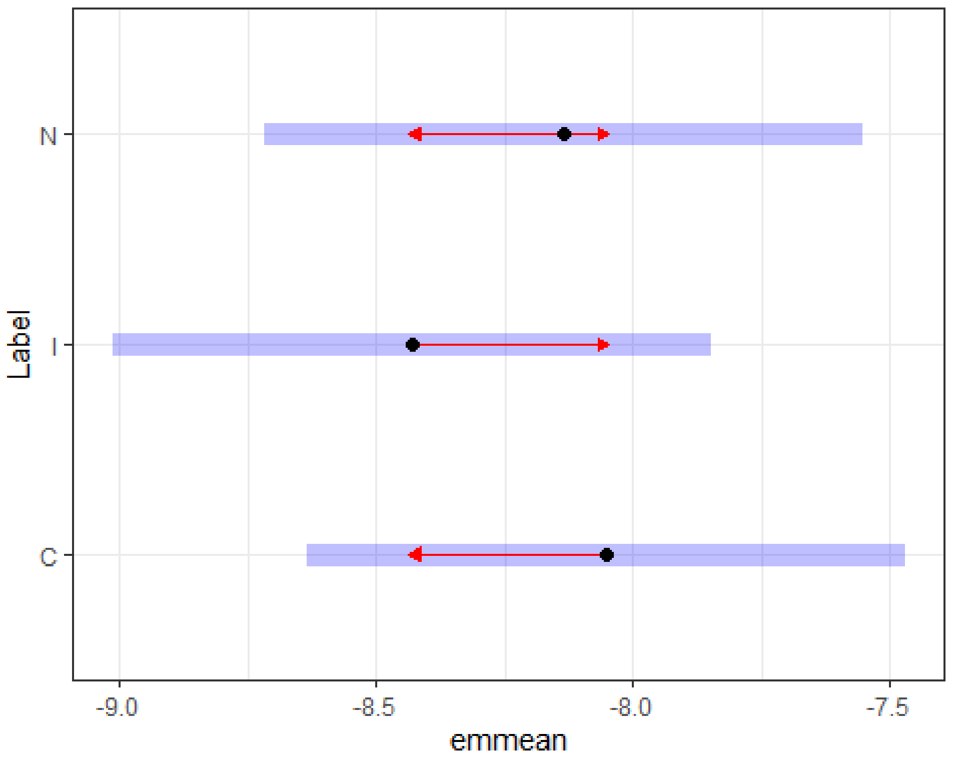


```
multcomp::cld(TukeyRes, Letters = LETTERS)
```

```
##  Label emmean    SE df lower.CL upper.CL .group
##  I       -8.43 0.277 18    -9.01    -7.85  A
##  N       -8.13 0.277 18    -8.72    -7.55  A
##  C       -8.05 0.277 18    -8.64    -7.47  A
##
## Results are averaged over the levels of: Donor
## Confidence level used: 0.95
## P value adjustment: tukey method for comparing a family of 3 estimates
## significance level used: alpha = 0.05
## NOTE: If two or more means share the same grouping symbol,
##       then we cannot show them to be different.
##       But we also did not show them to be the same.
```

Glucose

```
lm0 <- lm(GlucoseT ~ Label+Donor, data=TICsetR2)
Anova(lm0)
```

```
## Anova Table (Type II tests)
##
## Response: GlucoseT
##            Sum Sq Df F value Pr(>F)
## Label     0.04663  2  0.5393 0.5923
## Donor     0.50560  9  1.2996 0.3030
## Residuals 0.77810 18
```

```
resid_panel(lm0, "R")
```

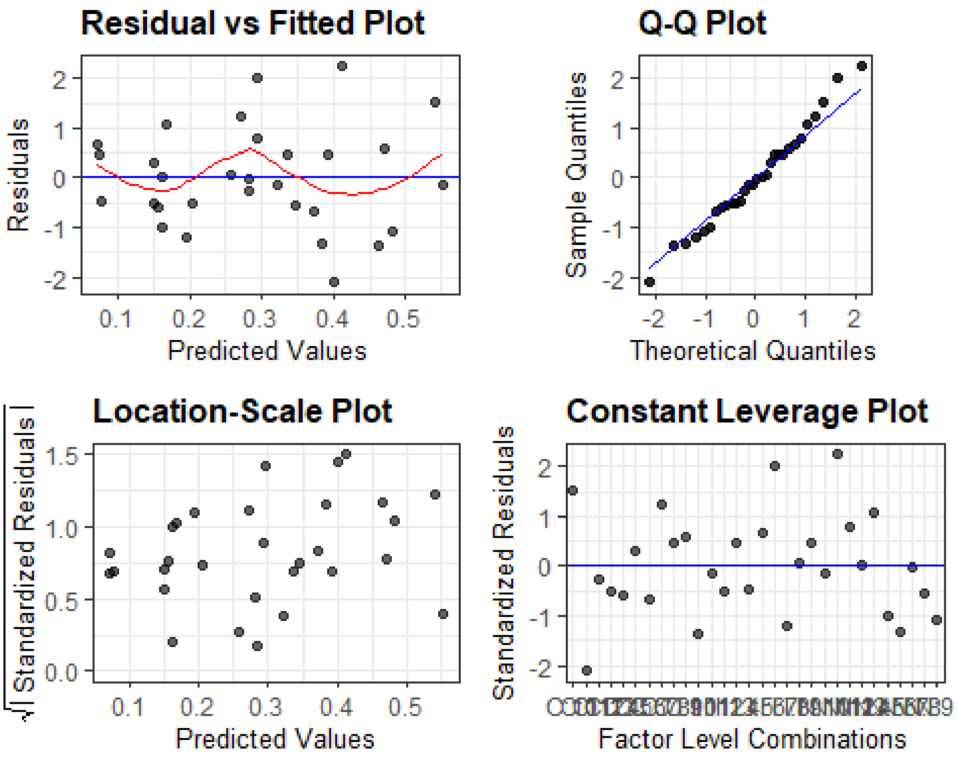


```
summary(lm0)

## 
## Call:
## lm(formula = GlucoseT ~ Label + Donor, data = TICsetR2)
## 
## Residuals:
##      Min       1Q   Median       3Q      Max 
## -0.34046 -0.09347 -0.01402  0.09179  0.36402 
## 
## Coefficients:
##              Estimate Std. Error t value Pr(>|t|)  
## (Intercept)  0.155895   0.131496   1.186   0.2512  
## LabelI      -0.077743   0.092982  -0.836   0.4141  
## LabelN       0.010741   0.092982   0.116   0.9093  
## Donor5      -0.006010   0.169760  -0.035   0.9721  
## Donor6       0.216467   0.169760   1.275   0.2185  
## Donor7       0.116597   0.169760   0.687   0.5009  
## Donor8       0.179960   0.169760   1.060   0.3031  
## Donor9       0.314681   0.169760   1.854   0.0802 .
## Donor10      0.384824   0.169760   2.267   0.0360 *
## Donor11      0.244844   0.169760   1.442   0.1664  
## Donor12      0.126370   0.169760   0.744   0.4662  
## Donor13     -0.004851   0.169760  -0.029   0.9775  
## ---
```

```
## Signif. codes:  0 '***' 0.001 '**' 0.01 '*' 0.05 '.' 0.1 ' ' 1
##
## Residual standard error: 0.2079 on 18 degrees of freedom
## Multiple R-squared:  0.4151, Adjusted R-squared:  0.05767
## F-statistic: 1.161 on 11 and 18 DF,  p-value: 0.3756
```

```
plot(allEffects(lm0), grid = T)
```

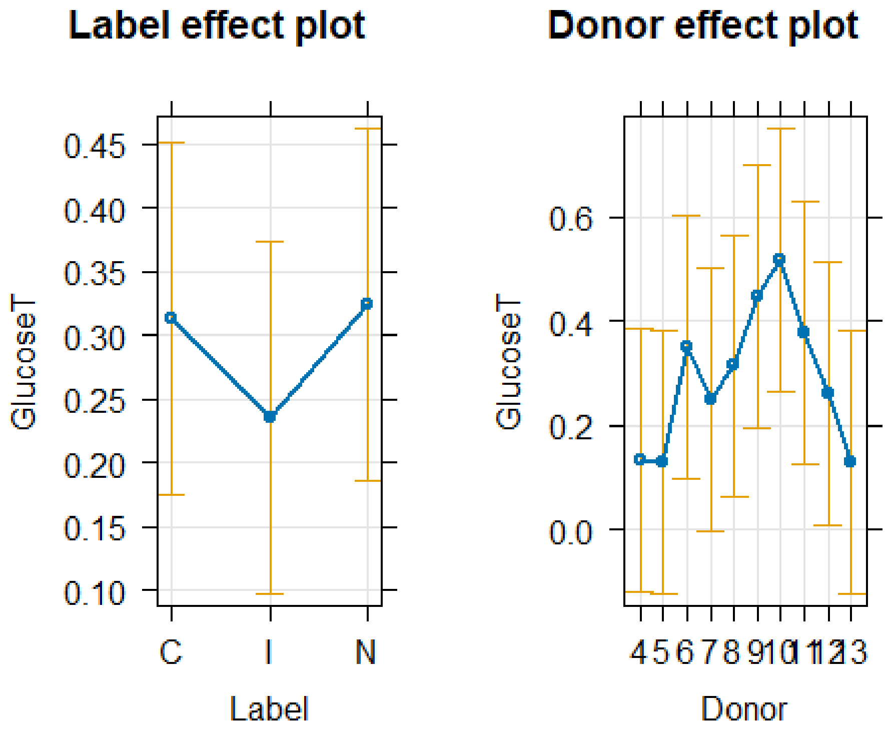


```
TukeyRes <- emmeans(lm0, pairwise ~ Label, adjust = "tukey")
TukeyRes
```

```
## $emmeans
##  Label emmean     SE df lower.CL upper.CL
##  C      0.313 0.0657 18   0.1751    0.451
##  I      0.235 0.0657 18   0.0973    0.374
##  N      0.324 0.0657 18   0.1858    0.462
##
## Results are averaged over the levels of: Donor
## Confidence level used: 0.95
##
## $contrasts
##  contrast estimate    SE df t.ratio p.value
##  C - I      0.0777 0.093 18   0.836  0.6860
##  C - N     -0.0107 0.093 18  -0.116  0.9927
##  I - N     -0.0885 0.093 18  -0.952  0.6157
##
```

```
## Results are averaged over the levels of: Donor
## P value adjustment: tukey method for comparing a family of 3 estimates
```

```r
confint(TukeyRes)
```

```
## $emmeans
##  Label emmean     SE df lower.CL upper.CL
##  C      0.313 0.0657 18   0.1751    0.451
##  I      0.235 0.0657 18   0.0973    0.374
##  N      0.324 0.0657 18   0.1858    0.462
##
## Results are averaged over the levels of: Donor
## Confidence level used: 0.95
##
## $contrasts
##  contrast estimate    SE df lower.CL upper.CL
##  C - I      0.0777 0.093 18   -0.160    0.315
##  C - N     -0.0107 0.093 18   -0.248    0.227
##  I - N     -0.0885 0.093 18   -0.326    0.149
##
## Results are averaged over the levels of: Donor
## Confidence level used: 0.95
## Conf-level adjustment: tukey method for comparing a family of 3 estimates
```

```r
plot(TukeyRes, comparison = T)
```

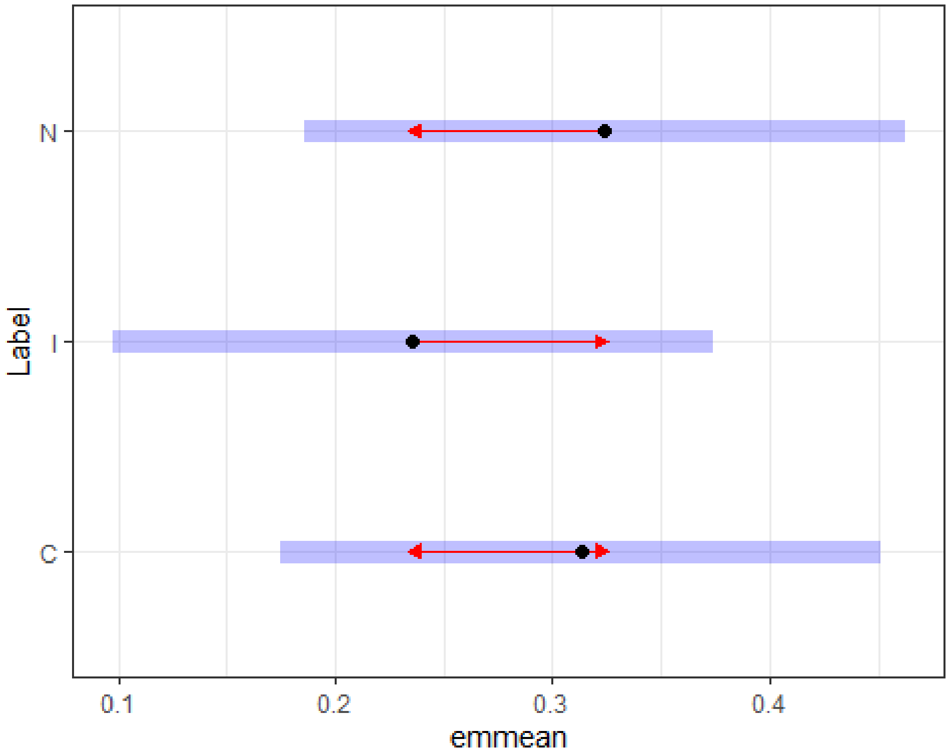


```r
multcomp::cld(TukeyRes, Letters = LETTERS)
```

```
##  Label emmean     SE df lower.CL upper.CL .group
##  I       0.235 0.0657 18   0.0973    0.374  A
##  C       0.313 0.0657 18   0.1751    0.451  A
##  N       0.324 0.0657 18   0.1858    0.462  A
##
## Results are averaged over the levels of: Donor
## Confidence level used: 0.95
## P value adjustment: tukey method for comparing a family of 3 estimates
## significance level used: alpha = 0.05
## NOTE: If two or more means share the same grouping symbol,
##       then we cannot show them to be different.
##       But we also did not show them to be the same.
```

Time Based Comparisons Binomial groups between 1/3 and 80% detection

#### Remove control to explore time by I or N interaction:

Pyruvic Acid

```
TICsetR <- TICset %>% dplyr::filter(Label != "C") %>%
  mutate(PyruvicB = factor(PyruvicB))

glm3 <- glm(PyruvicB ~ TimeF+Label, data=TICsetR, family = "binomial")

#Can't add donor - causes separation here
Anova(glm3)

## Analysis of Deviance Table (Type II tests)
##
## Response: PyruvicB
##       LR Chisq Df Pr(>Chisq)
## TimeF  23.5603  3  3.086e-05 ***
## Label   0.6467  1     0.4213
## ---
## Signif. codes:  0 '***' 0.001 '**' 0.01 '*' 0.05 '.' 0.1 ' ' 1

summary(glm3)

##
## Call:
## glm(formula = PyruvicB ~ TimeF + Label, family = "binomial",
##     data = TICsetR)
##
## Coefficients:
##             Estimate Std. Error z value Pr(>|z|)
## (Intercept)  18.7939  1450.2576   0.013    0.990
## TimeF1      -18.3749  1450.2576  -0.013    0.990
## TimeF4      -18.9880  1450.2576  -0.013    0.990
## TimeF24     -17.7212  1450.2576  -0.012    0.990
## LabelN       -0.4321     0.5392  -0.801    0.423
##
```

```
## (Dispersion parameter for binomial family taken to be 1)
##
##     Null deviance: 102.298  on 79  degrees of freedom
## Residual deviance:  78.234  on 75  degrees of freedom
## AIC: 88.234
##
## Number of Fisher Scoring iterations: 17
```

```
plot(allEffects(glm3), type = "response", grid = T)
```

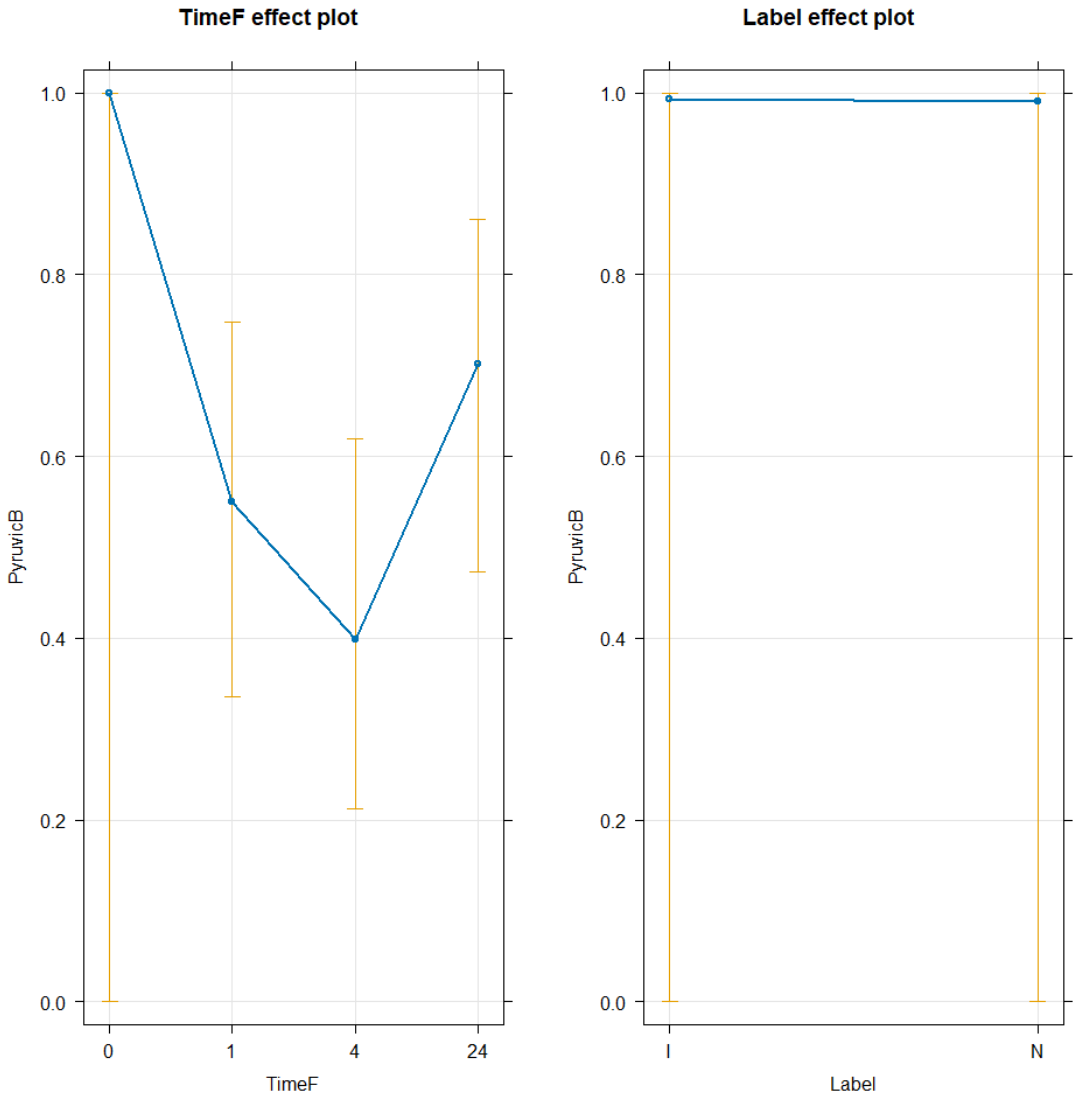


Glutemic Acid

```
TICsetR <- TICset %>% dplyr::filter(Label != "C") %>%
  mutate(GlutemicAcidB = factor(GlutemicAcidB))

glm3 <- glm(GlutemicAcidB ~ TimeF+Label, data=TICsetR, family = "binomial")
```

```r
#Can't add donor - causes separation here
Anova(glm3)

## Analysis of Deviance Table (Type II tests)
##
## Response: GlutemicAcidB
##       LR Chisq Df Pr(>Chisq)
## TimeF   5.0452  3     0.1685
## Label   0.2152  1     0.6427

summary(glm3)

##
## Call:
## glm(formula = GlutemicAcidB ~ TimeF + Label, family = "binomial",
##     data = TICsetR)
##
## Coefficients:
##               Estimate Std. Error z value Pr(>|z|)
## (Intercept) -9.360e-02  5.054e-01  -0.185   0.8531
## TimeF1       1.303e+00  6.856e-01   1.900   0.0574 .
## TimeF4      -1.530e-16  6.366e-01   0.000   1.0000
## TimeF24      4.025e-01  6.366e-01   0.632   0.5272
## LabelN      -2.153e-01  4.645e-01  -0.463   0.6430
## ---
## Signif. codes:  0 '***' 0.001 '**' 0.01 '*' 0.05 '.' 0.1 ' ' 1
##
## (Dispersion parameter for binomial family taken to be 1)
##
##     Null deviance: 110.10  on 79  degrees of freedom
## Residual deviance: 104.85  on 75  degrees of freedom
## AIC: 114.85
##
## Number of Fisher Scoring iterations: 4

plot(allEffects(glm3), type = "response", grid = T)
```

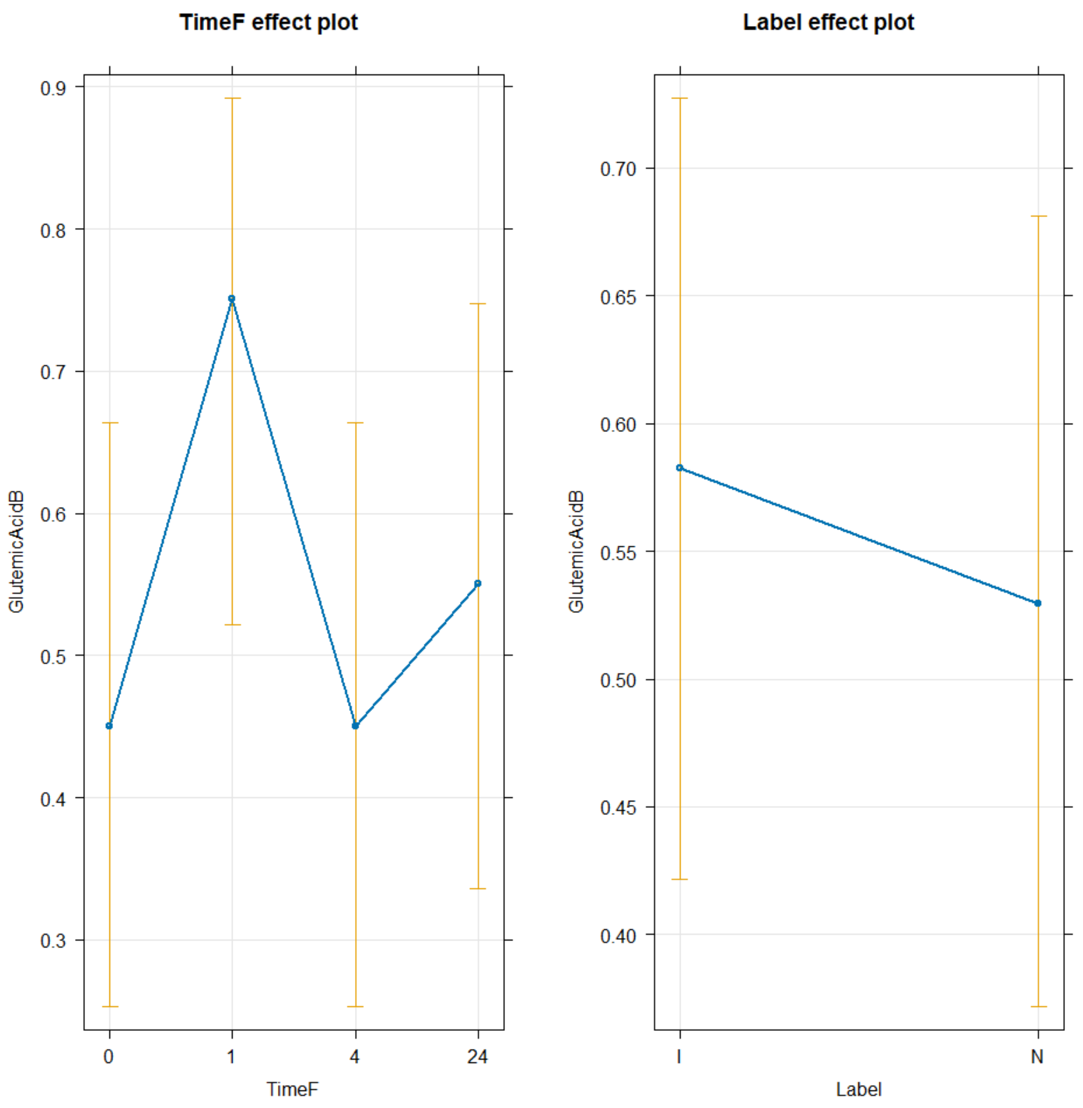


HSCOA

```
TICsetR <- TICset %>% dplyr::filter(Label != "C") %>%
  mutate(HSCOAB = factor(HSCOAB))

glm3 <- glm(HSCOAB ~ TimeF+Label, data=TICsetR, family = "binomial")

#Can't add donor - causes separation here
Anova(glm3)

## Analysis of Deviance Table (Type II tests)
##
## Response: HSCOAB
##        LR Chisq Df Pr(>Chisq)
## TimeF    3.9844  3     0.2632
## Label    0.9614  1     0.3268
```

```r
summary(glm3)

##
## Call:
## glm(formula = HSCOAB ~ TimeF + Label, family = "binomial", data = TICsetR)
##
## Coefficients:
##             Estimate Std. Error z value Pr(>|z|)
## (Intercept)  -1.6449     0.6289  -2.616   0.0089 **
## TimeF1        1.2001     0.7219   1.662   0.0964 .
## TimeF4        0.2906     0.7648   0.380   0.7040
## TimeF24       0.9924     0.7261   1.367   0.1717
## LabelN        0.4823     0.4945   0.975   0.3293
## ---
## Signif. codes:  0 '***' 0.001 '**' 0.01 '*' 0.05 '.' 0.1 ' ' 1
##
## (Dispersion parameter for binomial family taken to be 1)
##
##     Null deviance: 100.893  on 79  degrees of freedom
## Residual deviance:  95.994  on 75  degrees of freedom
## AIC: 105.99
##
## Number of Fisher Scoring iterations: 4

plot(allEffects(glm3), type = "response", grid = T)
```

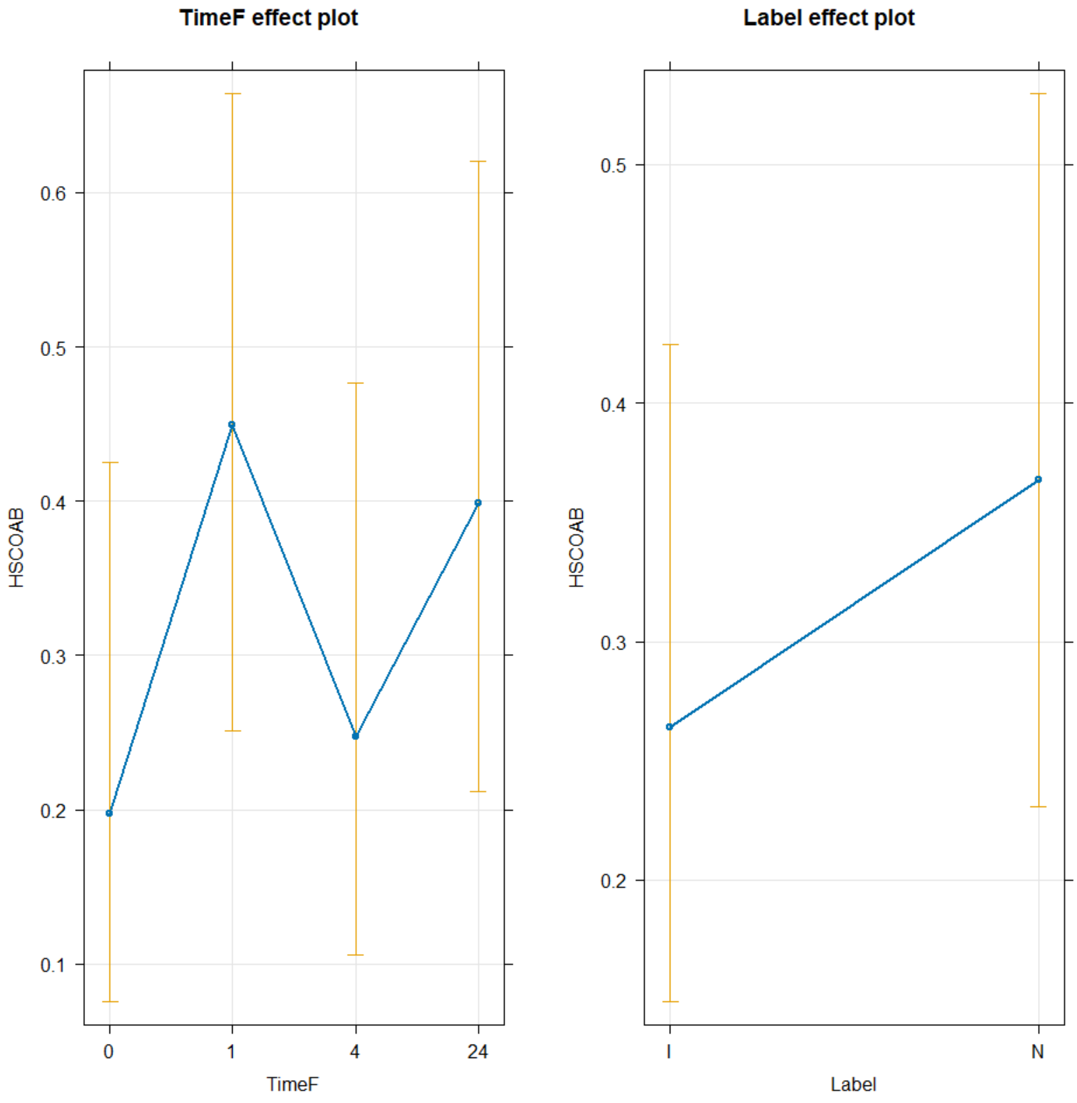


6-Phosphogluconate

```
TICsetR <- TICset %>% dplyr::filter(Label != "C") %>%
  mutate(SixPhosphoB = factor(SixPhosphoB))

glm3 <- glm(SixPhosphoB ~ TimeF+Label, data=TICsetR, family = "binomial")

#Can't add donor - causes separation here
Anova(glm3)

## Analysis of Deviance Table (Type II tests)
##
## Response: SixPhosphoB
##       LR Chisq Df Pr(>Chisq)
## TimeF  1.81607  3     0.6114
## Label  0.20522  1     0.6505
```

```
summary(glm3)

##
## Call:
## glm(formula = SixPhosphoB ~ TimeF + Label, family = "binomial",
##     data = TICsetR)
##
## Coefficients:
##                Estimate Std. Error z value Pr(>|z|)
## (Intercept) -3.039e-01  5.081e-01  -0.598    0.550
## TimeF1      -6.428e-16  6.463e-01   0.000    1.000
## TimeF4       6.077e-01  6.414e-01   0.947    0.343
## TimeF24      6.077e-01  6.414e-01   0.947    0.343
## LabelN      -2.053e-01  4.536e-01  -0.453    0.651
##
## (Dispersion parameter for binomial family taken to be 1)
##
##     Null deviance: 110.70  on 79  degrees of freedom
## Residual deviance: 108.69  on 75  degrees of freedom
## AIC: 118.69
##
## Number of Fisher Scoring iterations: 4

plot(allEffects(glm3), type = "response", grid = T)
```

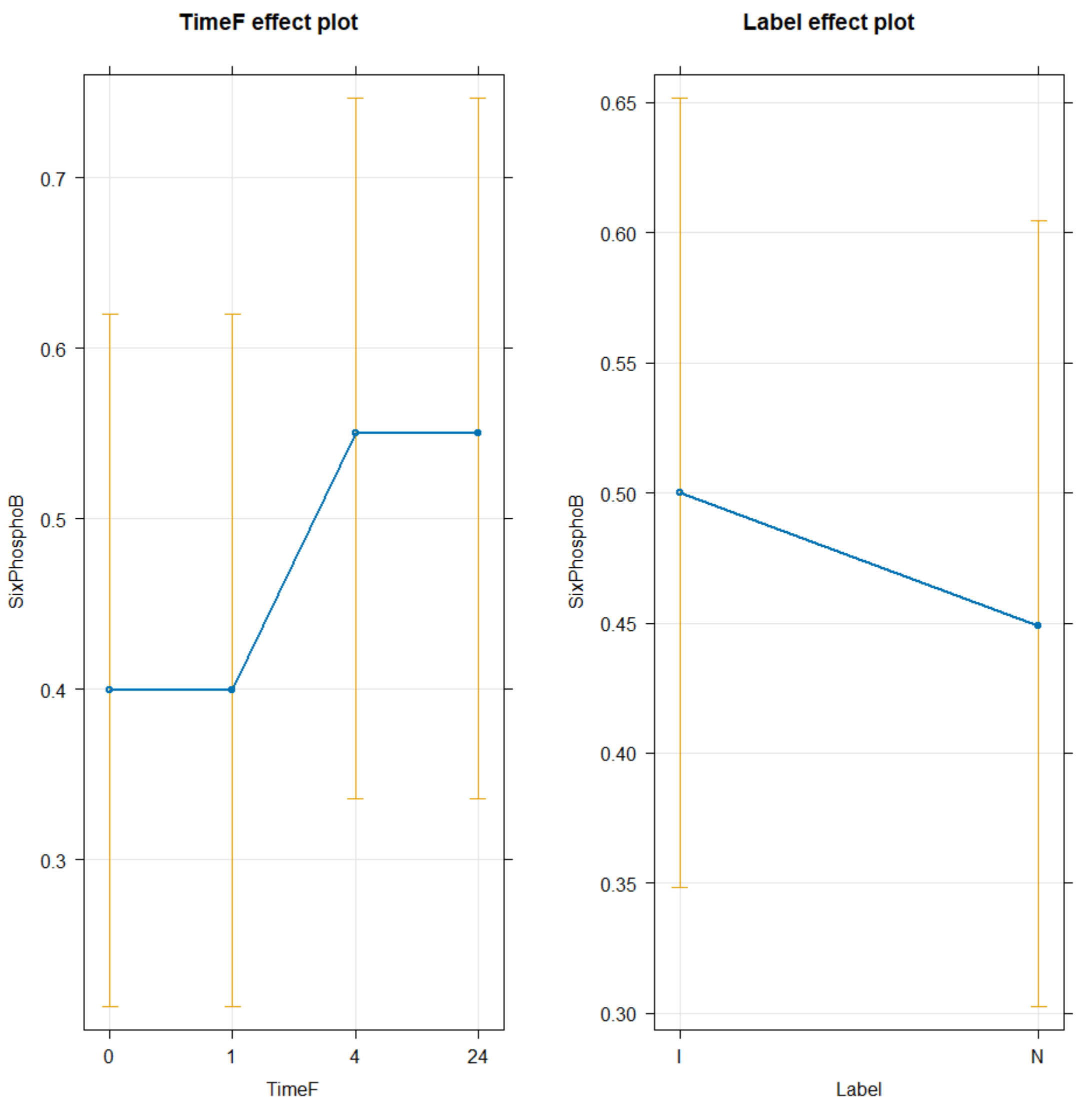


PEP

```
TICsetR <- TICset %>% dplyr::filter(Label != "C") %>%
  mutate(PhosphoenolB = factor(PhosphoenolB))

glm3 <- glm(PhosphoenolB ~ TimeF+Label, data=TICsetR, family = "binomial")

#Can't add donor - causes separation here
Anova(glm3)

## Analysis of Deviance Table (Type II tests)
##
## Response: PhosphoenolB
##       LR Chisq Df Pr(>Chisq)
## TimeF   3.5288  3     0.3170
## Label   1.6085  1     0.2047
```

```r
summary(glm3)

##
## Call:
## glm(formula = PhosphoenolB ~ TimeF + Label, family = "binomial",
##     data = TICsetR)
##
## Coefficients:
##               Estimate Std. Error z value Pr(>|z|)
## (Intercept) -8.010e-01  5.644e-01  -1.419    0.156
## TimeF1      -2.928e-01  7.678e-01  -0.381    0.703
## TimeF4       9.189e-01  6.930e-01   1.326    0.185
## TimeF24     -4.070e-16  7.374e-01   0.000    1.000
## LabelN      -6.479e-01  5.162e-01  -1.255    0.209
##
## (Dispersion parameter for binomial family taken to be 1)
##
##     Null deviance: 95.984  on 79  degrees of freedom
## Residual deviance: 90.920  on 75  degrees of freedom
## AIC: 100.92
##
## Number of Fisher Scoring iterations: 4

plot(allEffects(glm3), type = "response", grid = T)
```

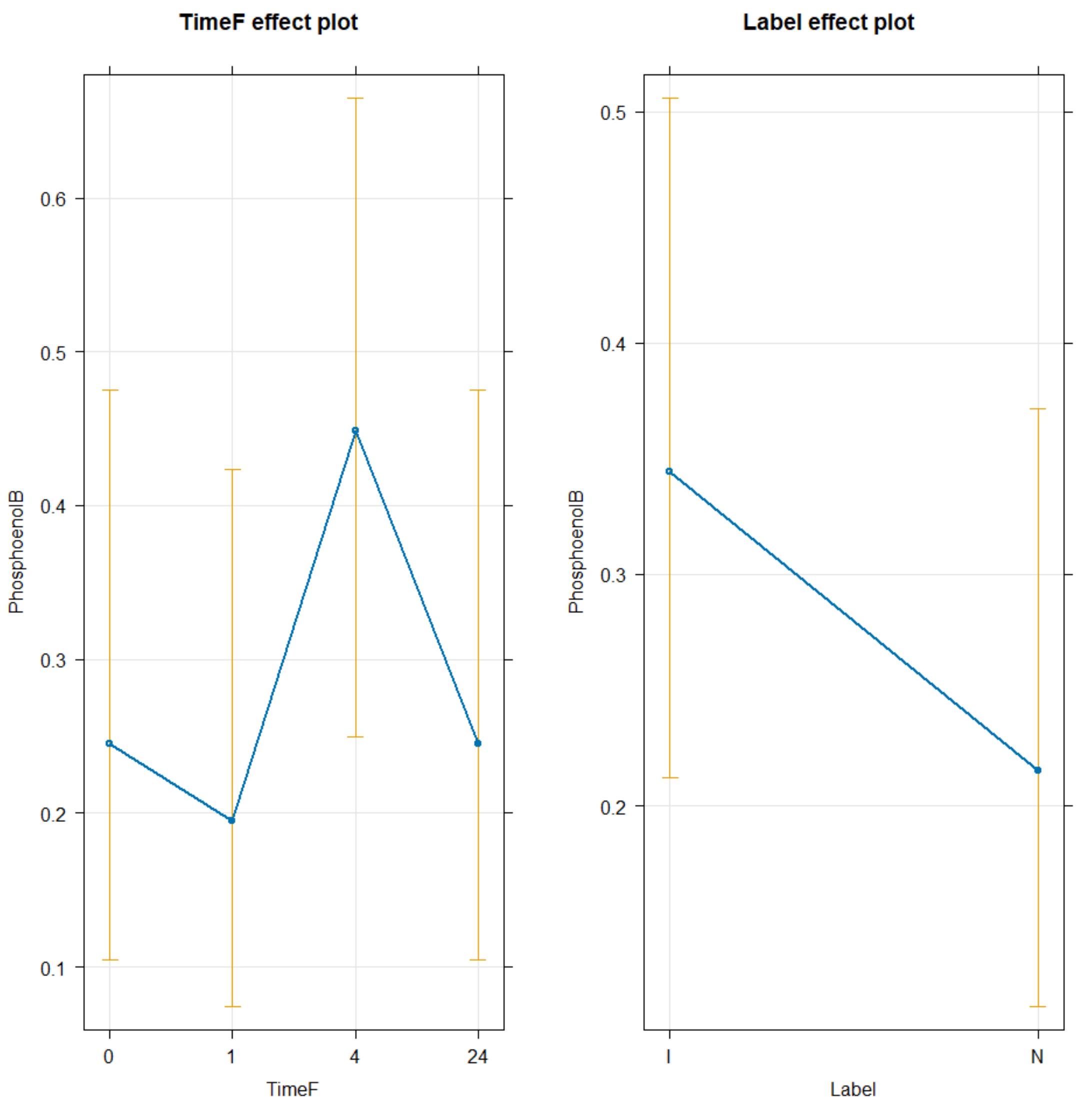


Logged NADH

```
lm1 <- lm(NADHT ~ Label+TimeF+Donor , data=TICsetR)
Anova(lm1)

## Anova Table (Type II tests)
##
## Response: NADHT
##            Sum Sq Df F value Pr(>F)
## Label     0.06765  1  1.5527 0.2171
## TimeF     0.22195  3  1.6981 0.1760
## Donor     0.31876  9  0.8129 0.6061
## Residuals 2.87557 66

lm2 <- lm(NADHT ~ Label+TimeF , data=TICsetR)
Anova(lm2)
```

```
## Anova Table (Type II tests)
##
## Response: NADHT
##           Sum Sq Df F value Pr(>F)
## Label     0.0677  1  1.5884 0.2115
## TimeF     0.2220  3  1.7371 0.1666
## Residuals 3.1943 75

#If need Tukey's in interaction model:

TukeyRes2 <- emmeans(lm2, pairwise ~ Label+TimeF, adjust = "tukey")
TukeyRes2

## $emmeans
##  Label TimeF emmean     SE df lower.CL upper.CL
##  I     0      -11.1 0.0516 75    -11.2    -11.0
##  N     0      -11.1 0.0516 75    -11.2    -11.0
##  I     1      -11.0 0.0516 75    -11.1    -10.9
##  N     1      -11.0 0.0516 75    -11.1    -10.9
##  I     4      -11.0 0.0516 75    -11.1    -10.9
##  N     4      -10.9 0.0516 75    -11.0    -10.8
##  I     24     -11.0 0.0516 75    -11.1    -10.9
##  N     24     -10.9 0.0516 75    -11.0    -10.8
##
## Confidence level used: 0.95
##
## $contrasts
##  contrast              estimate     SE df t.ratio p.value
##  I TimeF0 - N TimeF0   -0.05816 0.0461 75  -1.260  0.9103
##  I TimeF0 - I TimeF1   -0.08651 0.0653 75  -1.326  0.8862
##  I TimeF0 - N TimeF1   -0.14467 0.0799 75  -1.810  0.6154
##  I TimeF0 - I TimeF4   -0.12695 0.0653 75  -1.945  0.5251
##  I TimeF0 - N TimeF4   -0.18511 0.0799 75  -2.316  0.2990
##  I TimeF0 - I TimeF24  -0.13099 0.0653 75  -2.007  0.4841
##  I TimeF0 - N TimeF24  -0.18915 0.0799 75  -2.367  0.2730
##  N TimeF0 - I TimeF1   -0.02835 0.0799 75  -0.355  1.0000
##  N TimeF0 - N TimeF1   -0.08651 0.0653 75  -1.326  0.8862
##  N TimeF0 - I TimeF4   -0.06879 0.0799 75  -0.861  0.9886
##  N TimeF0 - N TimeF4   -0.12695 0.0653 75  -1.945  0.5251
##  N TimeF0 - I TimeF24  -0.07283 0.0799 75  -0.911  0.9841
##  N TimeF0 - N TimeF24  -0.13099 0.0653 75  -2.007  0.4841
##  I TimeF1 - N TimeF1   -0.05816 0.0461 75  -1.260  0.9103
##  I TimeF1 - I TimeF4   -0.04044 0.0653 75  -0.620  0.9985
##  I TimeF1 - N TimeF4   -0.09860 0.0799 75  -1.234  0.9192
##  I TimeF1 - I TimeF24  -0.04448 0.0653 75  -0.682  0.9973
##  I TimeF1 - N TimeF24  -0.10264 0.0799 75  -1.284  0.9019
##  N TimeF1 - I TimeF4    0.01772 0.0799 75   0.222  1.0000
##  N TimeF1 - N TimeF4   -0.04044 0.0653 75  -0.620  0.9985
##  N TimeF1 - I TimeF24   0.01368 0.0799 75   0.171  1.0000
##  N TimeF1 - N TimeF24  -0.04448 0.0653 75  -0.682  0.9973
```

```
##  I TimeF4 - N TimeF4   -0.05816 0.0461 75  -1.260  0.9103
##  I TimeF4 - I TimeF24  -0.00405 0.0653 75  -0.062  1.0000
##  I TimeF4 - N TimeF24  -0.06221 0.0799 75  -0.778  0.9938
##  N TimeF4 - I TimeF24   0.05411 0.0799 75   0.677  0.9974
##  N TimeF4 - N TimeF24  -0.00405 0.0653 75  -0.062  1.0000
##  I TimeF24 - N TimeF24 -0.05816 0.0461 75  -1.260  0.9103
##
## P value adjustment: tukey method for comparing a family of 8 estimates
```

```
confint(TukeyRes2)
```

```
## $emmeans
##  Label TimeF emmean     SE df lower.CL upper.CL
##  I     0      -11.1 0.0516 75    -11.2    -11.0
##  N     0      -11.1 0.0516 75    -11.2    -11.0
##  I     1      -11.0 0.0516 75    -11.1    -10.9
##  N     1      -11.0 0.0516 75    -11.1    -10.9
##  I     4      -11.0 0.0516 75    -11.1    -10.9
##  N     4      -10.9 0.0516 75    -11.0    -10.8
##  I     24     -11.0 0.0516 75    -11.1    -10.9
##  N     24     -10.9 0.0516 75    -11.0    -10.8
##
## Confidence level used: 0.95
##
## $contrasts
##  contrast              estimate     SE df lower.CL upper.CL
##  I TimeF0 - N TimeF0   -0.05816 0.0461 75   -0.202   0.0857
##  I TimeF0 - I TimeF1   -0.08651 0.0653 75   -0.290   0.1170
##  I TimeF0 - N TimeF1   -0.14467 0.0799 75   -0.394   0.1046
##  I TimeF0 - I TimeF4   -0.12695 0.0653 75   -0.330   0.0765
##  I TimeF0 - N TimeF4   -0.18511 0.0799 75   -0.434   0.0641
##  I TimeF0 - I TimeF24  -0.13099 0.0653 75   -0.334   0.0725
##  I TimeF0 - N TimeF24  -0.18915 0.0799 75   -0.438   0.0601
##  N TimeF0 - I TimeF1   -0.02835 0.0799 75   -0.278   0.2209
##  N TimeF0 - N TimeF1   -0.08651 0.0653 75   -0.290   0.1170
##  N TimeF0 - I TimeF4   -0.06879 0.0799 75   -0.318   0.1804
##  N TimeF0 - N TimeF4   -0.12695 0.0653 75   -0.330   0.0765
##  N TimeF0 - I TimeF24  -0.07283 0.0799 75   -0.322   0.1764
##  N TimeF0 - N TimeF24  -0.13099 0.0653 75   -0.334   0.0725
##  I TimeF1 - N TimeF1   -0.05816 0.0461 75   -0.202   0.0857
##  I TimeF1 - I TimeF4   -0.04044 0.0653 75   -0.244   0.1631
##  I TimeF1 - N TimeF4   -0.09860 0.0799 75   -0.348   0.1506
##  I TimeF1 - I TimeF24  -0.04448 0.0653 75   -0.248   0.1590
##  I TimeF1 - N TimeF24  -0.10264 0.0799 75   -0.352   0.1466
##  N TimeF1 - I TimeF4    0.01772 0.0799 75   -0.232   0.2669
##  N TimeF1 - N TimeF4   -0.04044 0.0653 75   -0.244   0.1631
##  N TimeF1 - I TimeF24   0.01368 0.0799 75   -0.236   0.2629
##  N TimeF1 - N TimeF24  -0.04448 0.0653 75   -0.248   0.1590
##  I TimeF4 - N TimeF4   -0.05816 0.0461 75   -0.202   0.0857
##  I TimeF4 - I TimeF24  -0.00405 0.0653 75   -0.208   0.1994
```

```
##  I TimeF4 - N TimeF24  -0.06221 0.0799 75   -0.311   0.1870
##  N TimeF4 - I TimeF24   0.05411 0.0799 75   -0.195   0.3033
##  N TimeF4 - N TimeF24  -0.00405 0.0653 75   -0.208   0.1994
##  I TimeF24 - N TimeF24 -0.05816 0.0461 75   -0.202   0.0857
##
## Confidence level used: 0.95
## Conf-level adjustment: tukey method for comparing a family of 8 estimates
```

```
plot(TukeyRes2, comparison = T) + coord_flip()
```

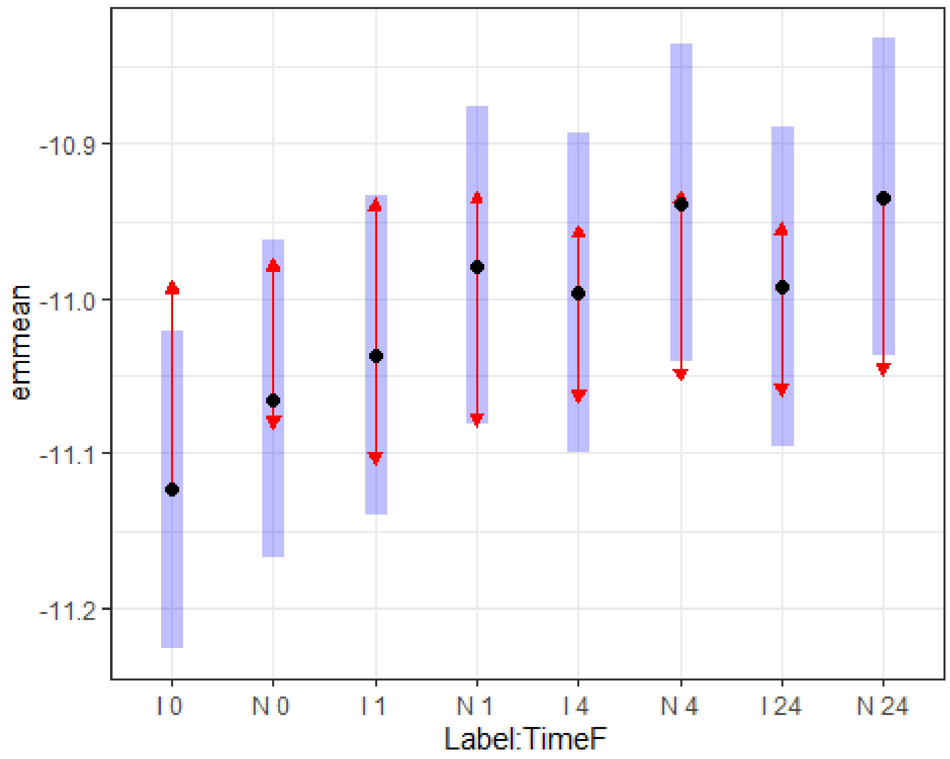


```
multcomp::cld(TukeyRes2, Letters = LETTERS)
```

```
##  Label TimeF emmean     SE df lower.CL upper.CL .group
##  I     0      -11.1 0.0516 75    -11.2    -11.0  A
##  N     0      -11.1 0.0516 75    -11.2    -11.0  A
##  I     1      -11.0 0.0516 75    -11.1    -10.9  A
##  I     4      -11.0 0.0516 75    -11.1    -10.9  A
##  I     24     -11.0 0.0516 75    -11.1    -10.9  A
##  N     1      -11.0 0.0516 75    -11.1    -10.9  A
##  N     4      -10.9 0.0516 75    -11.0    -10.8  A
##  N     24     -10.9 0.0516 75    -11.0    -10.8  A
##
## Confidence level used: 0.95
## P value adjustment: tukey method for comparing a family of 8 estimates
## significance level used: alpha = 0.05
## NOTE: If two or more means share the same grouping symbol,
```

```
##       then we cannot show them to be different.
##       But we also did not show them to be the same.

plot(allEffects(lm2), grid = T)
```

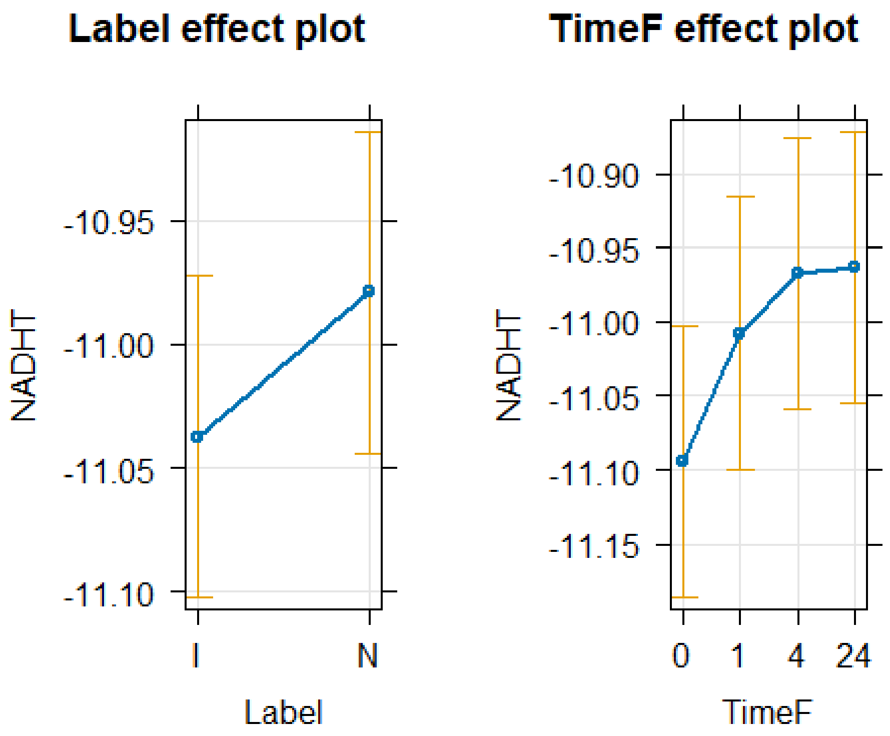


```
resid_panel(lm2, "R")
```

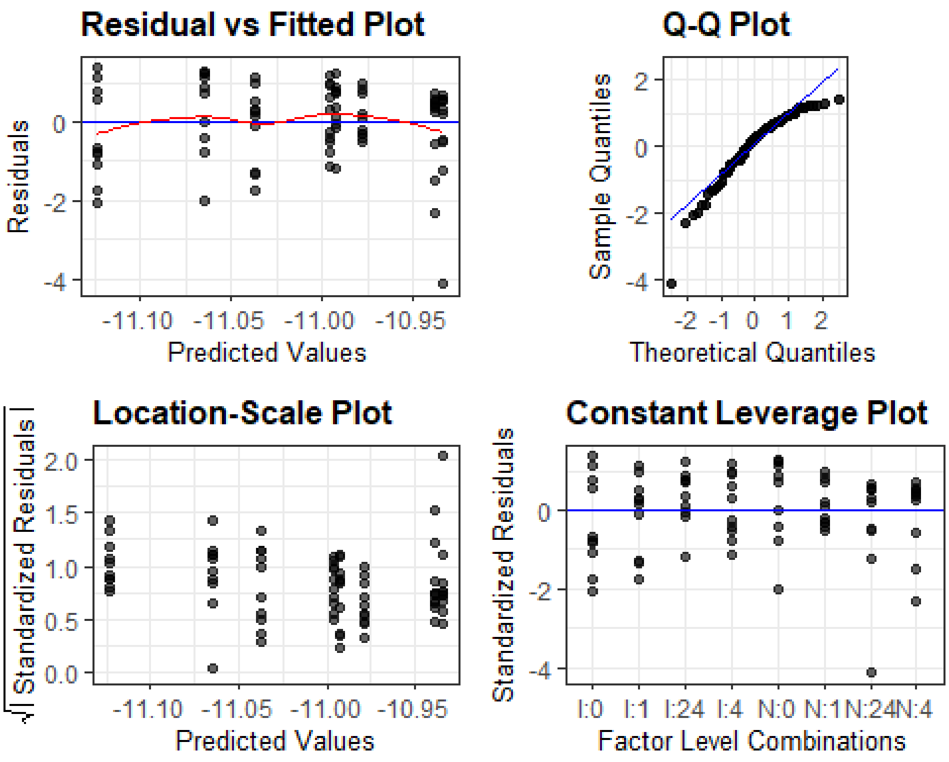


```
plot(allEffects(lm2, residuals = T), x.var = "TimeF:Label")
```

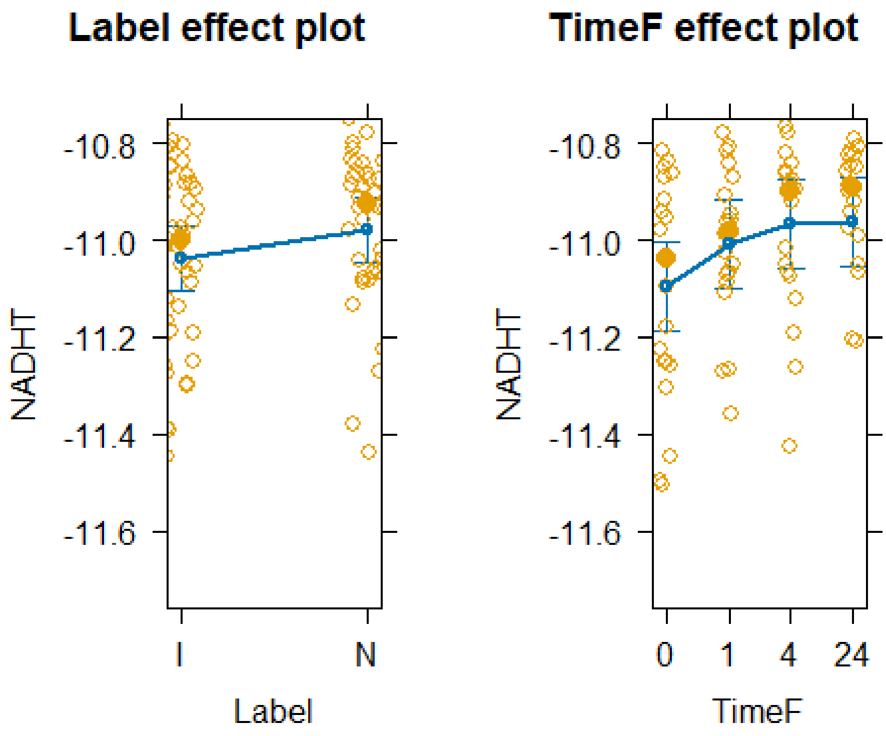


```
plot(allEffects(lm2), x.var = "Label", grid = T)
```

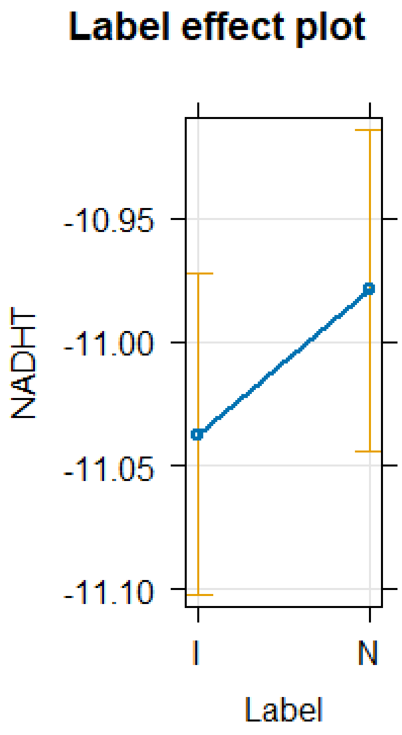


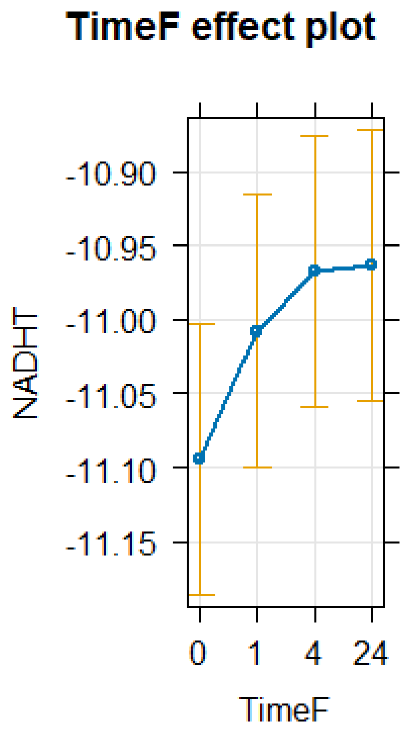


Logged Lactic Acid

```
lm1 <- lm(LacticAcidT ~ Label+TimeF+Donor , data=TICsetR)
Anova(lm1)

## Anova Table (Type II tests)
##
## Response: LacticAcidT
##           Sum Sq Df F value  Pr(>F)
## Label      3.880  1  4.3549 0.04077 *
## TimeF      1.678  3  0.6279 0.59953
## Donor     13.136  9  1.6384 0.12257
## Residuals 58.796 66
## ---
## Signif. codes:  0 '***' 0.001 '**' 0.01 '*' 0.05 '.' 0.1 ' ' 1

#If need Tukey's in interaction model:

TukeyRes2 <- emmeans(lm1, pairwise ~ Label+TimeF, adjust = "tukey")
TukeyRes2

## $emmeans
##  Label TimeF emmean    SE df lower.CL upper.CL
##  I     0      -8.50 0.236 66    -8.97    -8.03
##  N     0      -8.06 0.236 66    -8.53    -7.59
##  I     1      -8.23 0.236 66    -8.70    -7.76
##  N     1      -7.79 0.236 66    -8.26    -7.32
```

```
## I     4      -8.43 0.236 66    -8.90    -7.96
## N     4      -7.99 0.236 66    -8.46    -7.52
## I     24     -8.15 0.236 66    -8.62    -7.68
## N     24     -7.71 0.236 66    -8.18    -7.24
##
## Results are averaged over the levels of: Donor
## Confidence level used: 0.95
##
## $contrasts
##  contrast              estimate    SE df t.ratio p.value
##  I TimeF0 - N TimeF0    -0.4404 0.211 66  -2.087  0.4340
##  I TimeF0 - I TimeF1    -0.2752 0.298 66  -0.922  0.9828
##  I TimeF0 - N TimeF1    -0.7157 0.366 66  -1.958  0.5175
##  I TimeF0 - I TimeF4    -0.0709 0.298 66  -0.238  1.0000
##  I TimeF0 - N TimeF4    -0.5114 0.366 66  -1.399  0.8546
##  I TimeF0 - I TimeF24   -0.3550 0.298 66  -1.189  0.9322
##  I TimeF0 - N TimeF24   -0.7954 0.366 66  -2.176  0.3793
##  N TimeF0 - I TimeF1     0.1652 0.366 66   0.452  0.9998
##  N TimeF0 - N TimeF1    -0.2752 0.298 66  -0.922  0.9828
##  N TimeF0 - I TimeF4     0.3695 0.366 66   1.011  0.9712
##  N TimeF0 - N TimeF4    -0.0709 0.298 66  -0.238  1.0000
##  N TimeF0 - I TimeF24    0.0854 0.366 66   0.234  1.0000
##  N TimeF0 - N TimeF24   -0.3550 0.298 66  -1.189  0.9322
##  I TimeF1 - N TimeF1    -0.4404 0.211 66  -2.087  0.4340
##  I TimeF1 - I TimeF4     0.2043 0.298 66   0.685  0.9972
##  I TimeF1 - N TimeF4    -0.2361 0.366 66  -0.646  0.9980
##  I TimeF1 - I TimeF24   -0.0798 0.298 66  -0.267  1.0000
##  I TimeF1 - N TimeF24   -0.5202 0.366 66  -1.423  0.8433
##  N TimeF1 - I TimeF4     0.6447 0.366 66   1.764  0.6461
##  N TimeF1 - N TimeF4     0.2043 0.298 66   0.685  0.9972
##  N TimeF1 - I TimeF24    0.3607 0.366 66   0.987  0.9748
##  N TimeF1 - N TimeF24   -0.0798 0.298 66  -0.267  1.0000
##  I TimeF4 - N TimeF4    -0.4404 0.211 66  -2.087  0.4340
##  I TimeF4 - I TimeF24   -0.2841 0.298 66  -0.952  0.9794
##  I TimeF4 - N TimeF24   -0.7245 0.366 66  -1.982  0.5016
##  N TimeF4 - I TimeF24    0.1564 0.366 66   0.428  0.9999
##  N TimeF4 - N TimeF24   -0.2841 0.298 66  -0.952  0.9794
##  I TimeF24 - N TimeF24  -0.4404 0.211 66  -2.087  0.4340
##
## Results are averaged over the levels of: Donor
## P value adjustment: tukey method for comparing a family of 8 estimates
```

```r
confint(TukeyRes2)
```

```
## $emmeans
##  Label TimeF emmean    SE df lower.CL upper.CL
##  I     0      -8.50 0.236 66    -8.97    -8.03
##  N     0      -8.06 0.236 66    -8.53    -7.59
##  I     1      -8.23 0.236 66    -8.70    -7.76
##  N     1      -7.79 0.236 66    -8.26    -7.32
```

```
##  I     4      -8.43 0.236 66    -8.90    -7.96
##  N     4      -7.99 0.236 66    -8.46    -7.52
##  I     24     -8.15 0.236 66    -8.62    -7.68
##  N     24     -7.71 0.236 66    -8.18    -7.24
##
## Results are averaged over the levels of: Donor
## Confidence level used: 0.95
##
## $contrasts
##  contrast              estimate    SE df lower.CL upper.CL
##  I TimeF0 - N TimeF0    -0.4404 0.211 66   -1.101    0.220
##  I TimeF0 - I TimeF1    -0.2752 0.298 66   -1.210    0.659
##  I TimeF0 - N TimeF1    -0.7157 0.366 66   -1.860    0.429
##  I TimeF0 - I TimeF4    -0.0709 0.298 66   -1.005    0.863
##  I TimeF0 - N TimeF4    -0.5114 0.366 66   -1.656    0.633
##  I TimeF0 - I TimeF24   -0.3550 0.298 66   -1.289    0.579
##  I TimeF0 - N TimeF24   -0.7954 0.366 66   -1.940    0.349
##  N TimeF0 - I TimeF1     0.1652 0.366 66   -0.979    1.309
##  N TimeF0 - N TimeF1    -0.2752 0.298 66   -1.210    0.659
##  N TimeF0 - I TimeF4     0.3695 0.366 66   -0.775    1.514
##  N TimeF0 - N TimeF4    -0.0709 0.298 66   -1.005    0.863
##  N TimeF0 - I TimeF24    0.0854 0.366 66   -1.059    1.230
##  N TimeF0 - N TimeF24   -0.3550 0.298 66   -1.289    0.579
##  I TimeF1 - N TimeF1    -0.4404 0.211 66   -1.101    0.220
##  I TimeF1 - I TimeF4     0.2043 0.298 66   -0.730    1.139
##  I TimeF1 - N TimeF4    -0.2361 0.366 66   -1.380    0.908
##  I TimeF1 - I TimeF24   -0.0798 0.298 66   -1.014    0.855
##  I TimeF1 - N TimeF24   -0.5202 0.366 66   -1.664    0.624
##  N TimeF1 - I TimeF4     0.6447 0.366 66   -0.500    1.789
##  N TimeF1 - N TimeF4     0.2043 0.298 66   -0.730    1.139
##  N TimeF1 - I TimeF24    0.3607 0.366 66   -0.784    1.505
##  N TimeF1 - N TimeF24   -0.0798 0.298 66   -1.014    0.855
##  I TimeF4 - N TimeF4    -0.4404 0.211 66   -1.101    0.220
##  I TimeF4 - I TimeF24   -0.2841 0.298 66   -1.218    0.650
##  I TimeF4 - N TimeF24   -0.7245 0.366 66   -1.869    0.420
##  N TimeF4 - I TimeF24    0.1564 0.366 66   -0.988    1.301
##  N TimeF4 - N TimeF24   -0.2841 0.298 66   -1.218    0.650
##  I TimeF24 - N TimeF24  -0.4404 0.211 66   -1.101    0.220
##
## Results are averaged over the levels of: Donor
## Confidence level used: 0.95
## Conf-level adjustment: tukey method for comparing a family of 8 estimates
```

```r
plot(TukeyRes2, comparison = T) + coord_flip()
```

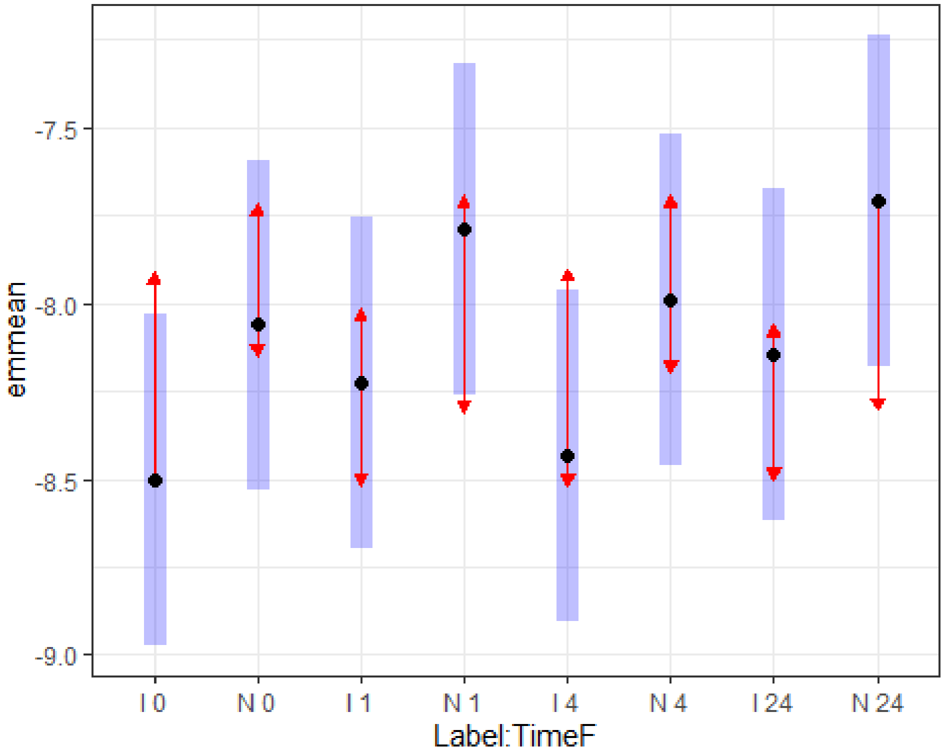


```
multcomp::cld(TukeyRes2, Letters = LETTERS)

##  Label TimeF emmean    SE df lower.CL upper.CL .group
##  I     0      -8.50 0.236 66    -8.97    -8.03  A
##  I     4      -8.43 0.236 66    -8.90    -7.96  A
##  I     1      -8.23 0.236 66    -8.70    -7.76  A
##  I     24     -8.15 0.236 66    -8.62    -7.68  A
##  N     0      -8.06 0.236 66    -8.53    -7.59  A
##  N     4      -7.99 0.236 66    -8.46    -7.52  A
##  N     1      -7.79 0.236 66    -8.26    -7.32  A
##  N     24     -7.71 0.236 66    -8.18    -7.24  A
##
## Results are averaged over the levels of: Donor
## Confidence level used: 0.95
## P value adjustment: tukey method for comparing a family of 8 estimates
## significance level used: alpha = 0.05
## NOTE: If two or more means share the same grouping symbol,
##       then we cannot show them to be different.
##       But we also did not show them to be the same.

plot(allEffects(lm1), grid = T)
```

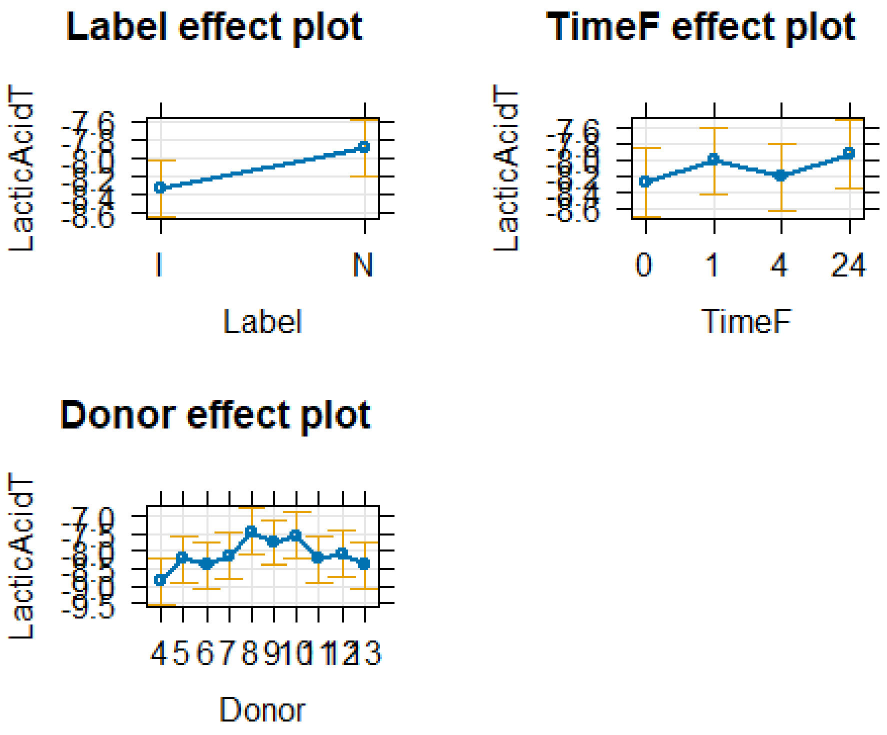


```r
resid_panel(lm1, "R")
```

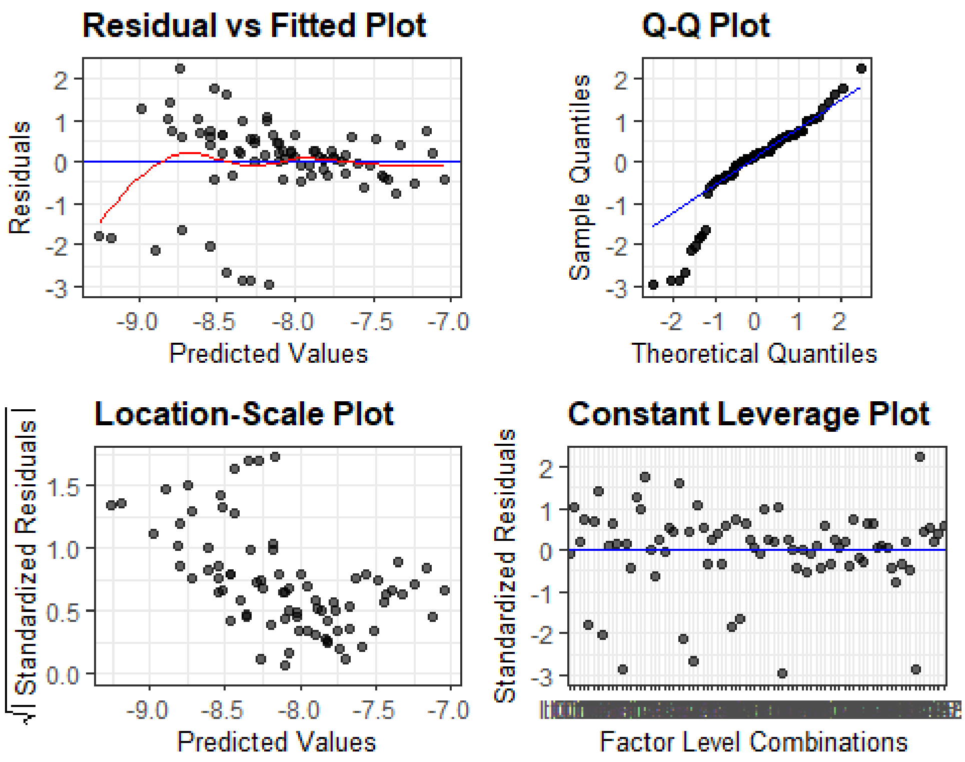


```r
plot(allEffects(lm1, residuals = T), x.var = "TimeF:Label")
```

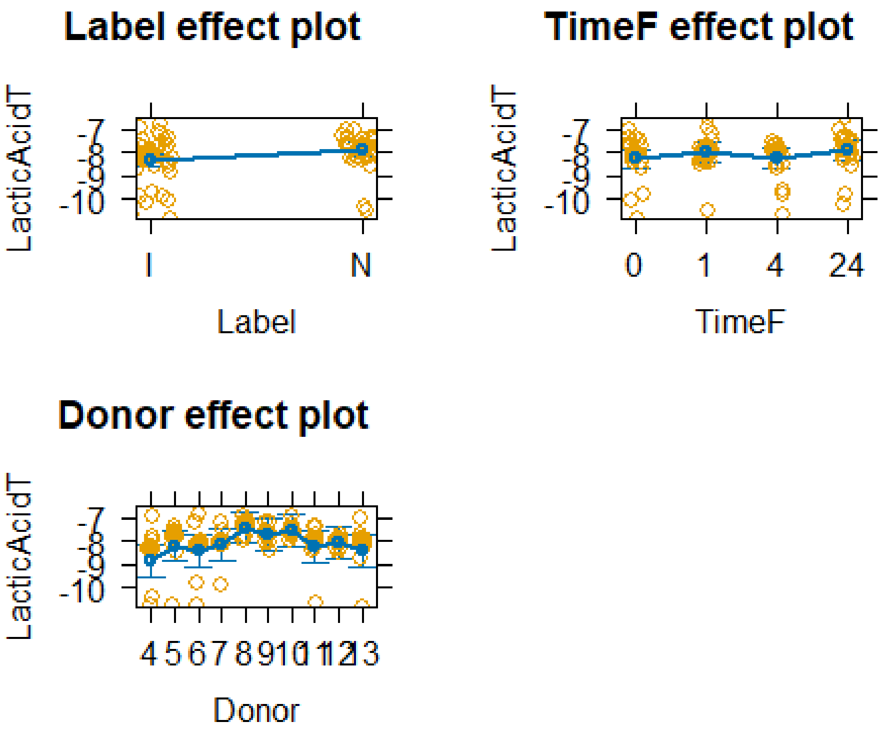


```
plot(allEffects(lm1), x.var = "Label", grid = T)
```

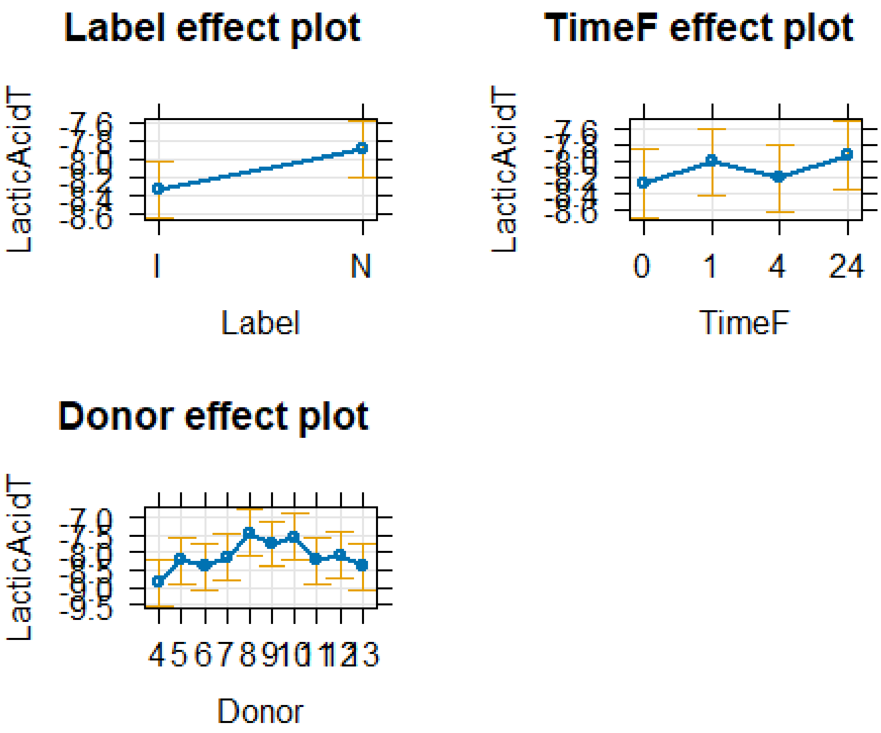


Glucose

```r
lm1 <- lm(GlucoseT ~ Label+TimeF+Donor , data=TICsetR)
Anova(lm1)

## Anova Table (Type II tests)
##
## Response: GlucoseT
##              Sum Sq Df F value  Pr(>F)
## Label     0.16715  1  4.9810 0.02903 *
## TimeF     0.09965  3  0.9898 0.40311
## Donor     0.78876  9  2.6117 0.01208 *
## Residuals 2.21475 66
## ---
## Signif. codes:  0 '***' 0.001 '**' 0.01 '*' 0.05 '.' 0.1 ' ' 1

#If need Tukey's in interaction model:

TukeyRes2 <- emmeans(lm1, pairwise ~ Label+TimeF, adjust = "tukey")
TukeyRes2

## $emmeans
##  Label TimeF emmean     SE df lower.CL upper.CL
##  I     0      0.234 0.0458 66    0.143    0.325
##  N     0      0.325 0.0458 66    0.234    0.417
##  I     1      0.235 0.0458 66    0.144    0.327
##  N     1      0.327 0.0458 66    0.235    0.418
##  I     4      0.208 0.0458 66    0.117    0.300
##  N     4      0.300 0.0458 66    0.208    0.391
##  I     24     0.303 0.0458 66    0.212    0.395
##  N     24     0.395 0.0458 66    0.303    0.486
##
## Results are averaged over the levels of: Donor
## Confidence level used: 0.95
##
## $contrasts
##  contrast                 estimate     SE df t.ratio p.value
##  I TimeF0 - N TimeF0   -0.09142 0.0410 66  -2.232  0.3469
##  I TimeF0 - I TimeF1   -0.00134 0.0579 66  -0.023  1.0000
##  I TimeF0 - N TimeF1   -0.09275 0.0709 66  -1.307  0.8929
##  I TimeF0 - I TimeF4    0.02588 0.0579 66   0.447  0.9998
##  I TimeF0 - N TimeF4   -0.06554 0.0709 66  -0.924  0.9826
##  I TimeF0 - I TimeF24  -0.06938 0.0579 66  -1.198  0.9298
##  I TimeF0 - N TimeF24  -0.16080 0.0709 66  -2.266  0.3275
##  N TimeF0 - I TimeF1    0.09008 0.0709 66   1.270  0.9067
##  N TimeF0 - N TimeF1   -0.00134 0.0579 66  -0.023  1.0000
##  N TimeF0 - I TimeF4    0.11730 0.0709 66   1.653  0.7164
##  N TimeF0 - N TimeF4    0.02588 0.0579 66   0.447  0.9998
##  N TimeF0 - I TimeF24   0.02204 0.0709 66   0.311  1.0000
##  N TimeF0 - N TimeF24  -0.06938 0.0579 66  -1.198  0.9298
##  I TimeF1 - N TimeF1   -0.09142 0.0410 66  -2.232  0.3469
##  I TimeF1 - I TimeF4    0.02722 0.0579 66   0.470  0.9998
```

```
##  I TimeF1 - N TimeF4   -0.06420 0.0709 66  -0.905  0.9846
##  I TimeF1 - I TimeF24  -0.06804 0.0579 66  -1.175  0.9363
##  I TimeF1 - N TimeF24  -0.15946 0.0709 66  -2.248  0.3380
##  N TimeF1 - I TimeF4    0.11863 0.0709 66   1.672  0.7047
##  N TimeF1 - N TimeF4    0.02722 0.0579 66   0.470  0.9998
##  N TimeF1 - I TimeF24   0.02338 0.0709 66   0.329  1.0000
##  N TimeF1 - N TimeF24  -0.06804 0.0579 66  -1.175  0.9363
##  I TimeF4 - N TimeF4   -0.09142 0.0410 66  -2.232  0.3469
##  I TimeF4 - I TimeF24  -0.09526 0.0579 66  -1.644  0.7219
##  I TimeF4 - N TimeF24  -0.18668 0.0709 66  -2.631  0.1634
##  N TimeF4 - I TimeF24  -0.00384 0.0709 66  -0.054  1.0000
##  N TimeF4 - N TimeF24  -0.09526 0.0579 66  -1.644  0.7219
##  I TimeF24 - N TimeF24 -0.09142 0.0410 66  -2.232  0.3469
##
## Results are averaged over the levels of: Donor
## P value adjustment: tukey method for comparing a family of 8 estimates
```

```
confint(TukeyRes2)
```

```
## $emmeans
##  Label TimeF emmean     SE df lower.CL upper.CL
##  I     0      0.234 0.0458 66    0.143    0.325
##  N     0      0.325 0.0458 66    0.234    0.417
##  I     1      0.235 0.0458 66    0.144    0.327
##  N     1      0.327 0.0458 66    0.235    0.418
##  I     4      0.208 0.0458 66    0.117    0.300
##  N     4      0.300 0.0458 66    0.208    0.391
##  I     24     0.303 0.0458 66    0.212    0.395
##  N     24     0.395 0.0458 66    0.303    0.486
##
## Results are averaged over the levels of: Donor
## Confidence level used: 0.95
##
## $contrasts
##  contrast              estimate     SE df lower.CL upper.CL
##  I TimeF0 - N TimeF0   -0.09142 0.0410 66   -0.220   0.0368
##  I TimeF0 - I TimeF1   -0.00134 0.0579 66   -0.183   0.1800
##  I TimeF0 - N TimeF1   -0.09275 0.0709 66   -0.315   0.1293
##  I TimeF0 - I TimeF4    0.02588 0.0579 66   -0.155   0.2072
##  I TimeF0 - N TimeF4   -0.06554 0.0709 66   -0.288   0.1565
##  I TimeF0 - I TimeF24  -0.06938 0.0579 66   -0.251   0.1119
##  I TimeF0 - N TimeF24  -0.16080 0.0709 66   -0.383   0.0613
##  N TimeF0 - I TimeF1    0.09008 0.0709 66   -0.132   0.3122
##  N TimeF0 - N TimeF1   -0.00134 0.0579 66   -0.183   0.1800
##  N TimeF0 - I TimeF4    0.11730 0.0709 66   -0.105   0.3394
##  N TimeF0 - N TimeF4    0.02588 0.0579 66   -0.155   0.2072
##  N TimeF0 - I TimeF24   0.02204 0.0709 66   -0.200   0.2441
##  N TimeF0 - N TimeF24  -0.06938 0.0579 66   -0.251   0.1119
##  I TimeF1 - N TimeF1   -0.09142 0.0410 66   -0.220   0.0368
##  I TimeF1 - I TimeF4    0.02722 0.0579 66   -0.154   0.2085
```

```
##  I TimeF1 - N TimeF4   -0.06420 0.0709 66   -0.286   0.1579
##  I TimeF1 - I TimeF24  -0.06804 0.0579 66   -0.249   0.1133
##  I TimeF1 - N TimeF24  -0.15946 0.0709 66   -0.382   0.0626
##  N TimeF1 - I TimeF4    0.11863 0.0709 66   -0.103   0.3407
##  N TimeF1 - N TimeF4    0.02722 0.0579 66   -0.154   0.2085
##  N TimeF1 - I TimeF24   0.02338 0.0709 66   -0.199   0.2455
##  N TimeF1 - N TimeF24  -0.06804 0.0579 66   -0.249   0.1133
##  I TimeF4 - N TimeF4   -0.09142 0.0410 66   -0.220   0.0368
##  I TimeF4 - I TimeF24  -0.09526 0.0579 66   -0.277   0.0861
##  I TimeF4 - N TimeF24  -0.18668 0.0709 66   -0.409   0.0354
##  N TimeF4 - I TimeF24  -0.00384 0.0709 66   -0.226   0.2182
##  N TimeF4 - N TimeF24  -0.09526 0.0579 66   -0.277   0.0861
##  I TimeF24 - N TimeF24 -0.09142 0.0410 66   -0.220   0.0368
##
## Results are averaged over the levels of: Donor
## Confidence level used: 0.95
## Conf-level adjustment: tukey method for comparing a family of 8 estimates
```

```
plot(TukeyRes2, comparison = T) + coord_flip()
```

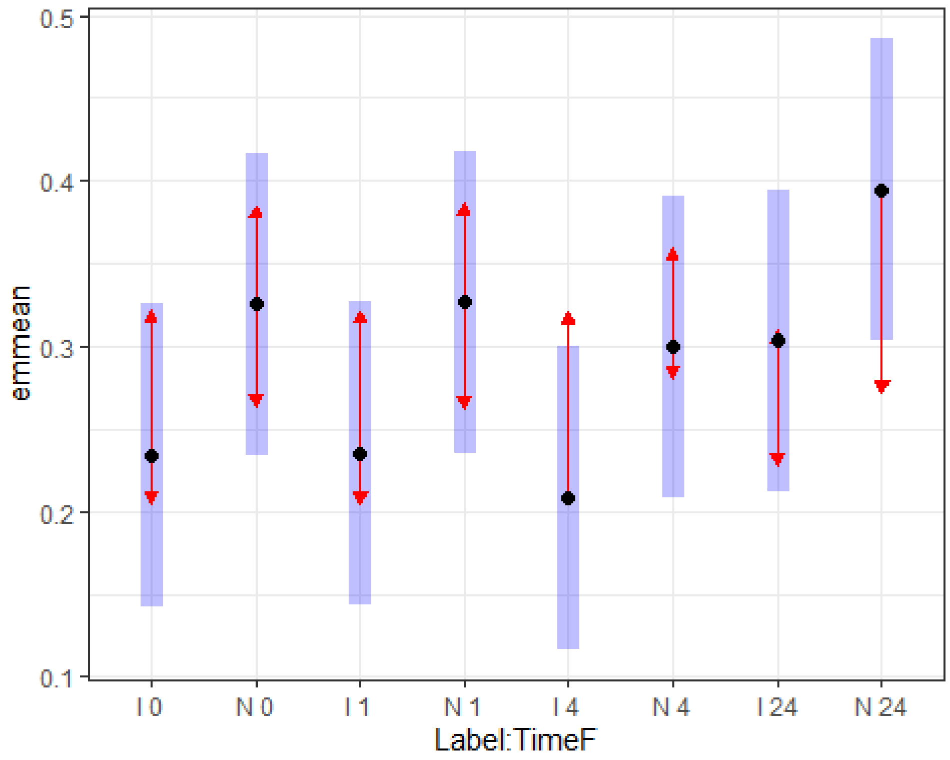


```
multcomp::cld(TukeyRes2, Letters = LETTERS)
```

```
##  Label TimeF emmean     SE df lower.CL upper.CL .group
##  I     4      0.208 0.0458 66    0.117    0.300  A
##  I     0      0.234 0.0458 66    0.143    0.325  A
##  I     1      0.235 0.0458 66    0.144    0.327  A
##  N     4      0.300 0.0458 66    0.208    0.391  A
```

```
##  I      24      0.303 0.0458 66     0.212     0.395  A
##  N      0       0.325 0.0458 66     0.234     0.417  A
##  N      1       0.327 0.0458 66     0.235     0.418  A
##  N      24      0.395 0.0458 66     0.303     0.486  A
##
## Results are averaged over the levels of: Donor
## Confidence level used: 0.95
## P value adjustment: tukey method for comparing a family of 8 estimates
## significance level used: alpha = 0.05
## NOTE: If two or more means share the same grouping symbol,
##       then we cannot show them to be different.
##       But we also did not show them to be the same.
```

```r
plot(allEffects(lm1), grid = T)
```

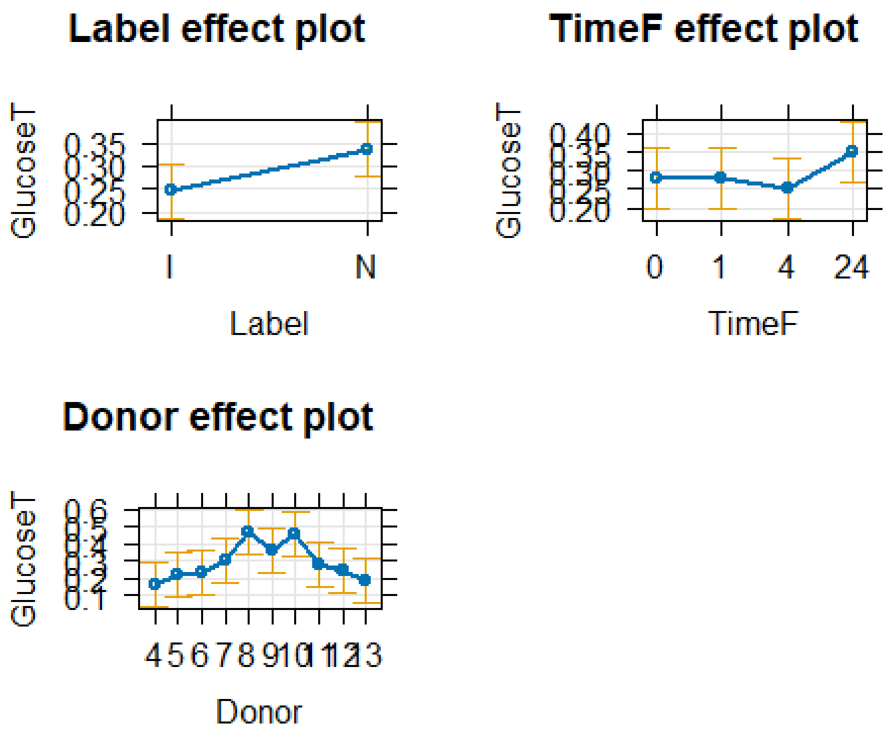


```r
resid_panel(lm1, "R")
```

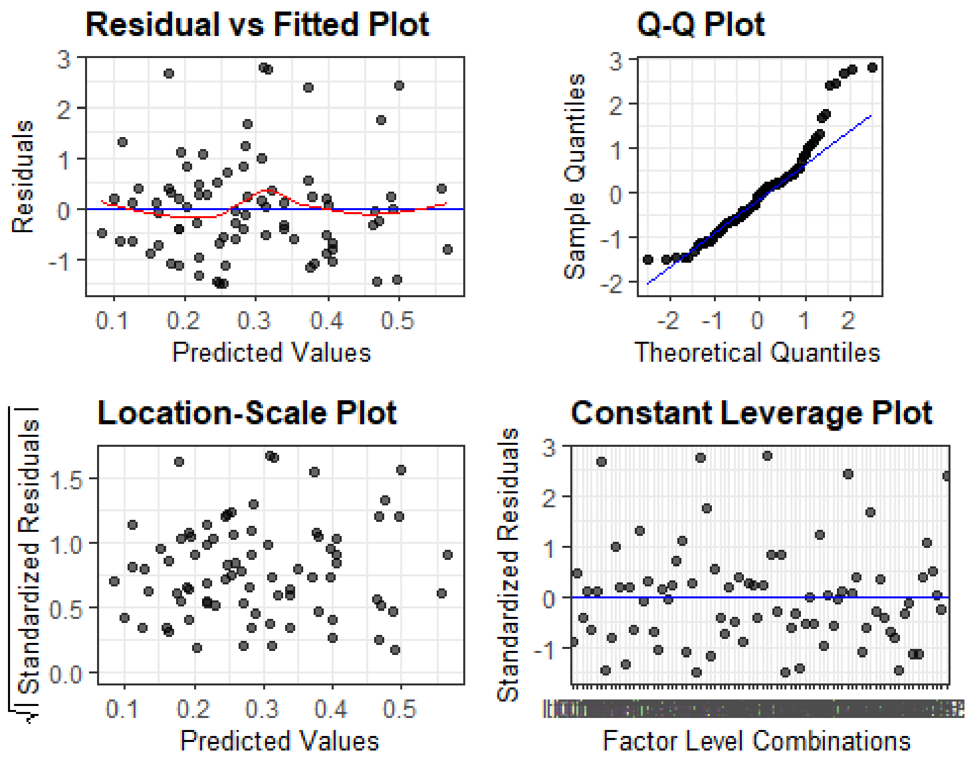


```
plot(allEffects(lm1, residuals = T), x.var = "TimeF:Label")
```

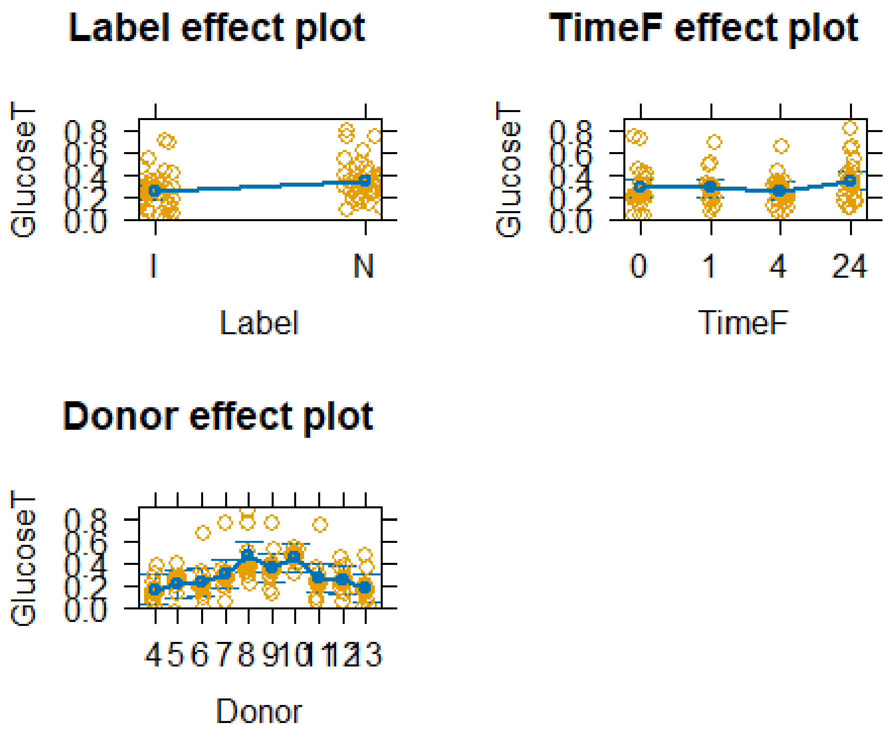


```
plot(allEffects(lm1), x.var = "Label", grid = T)
```

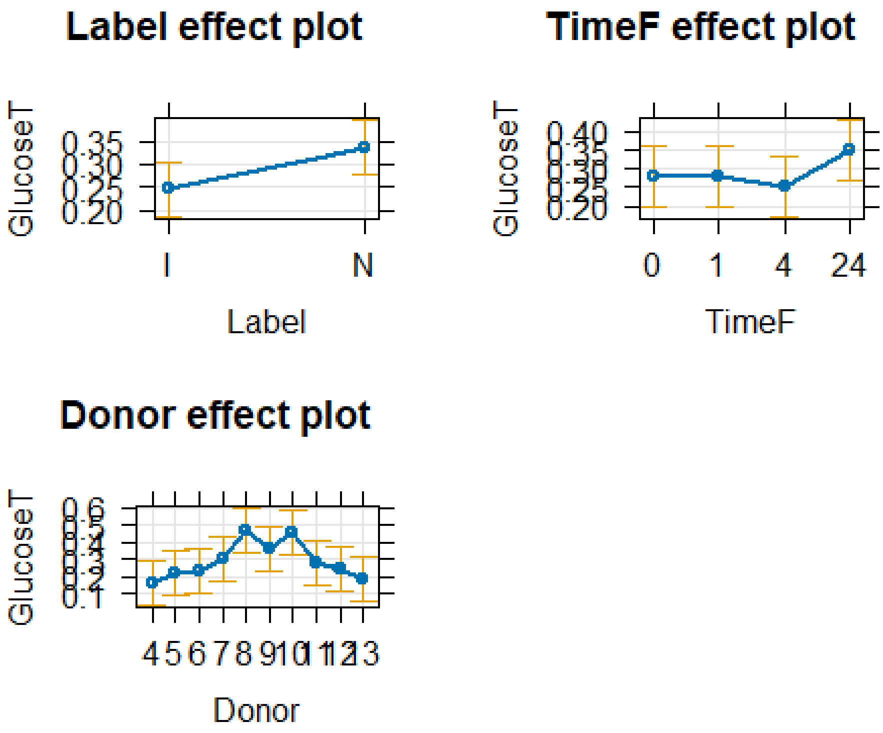


```
pdf("glutaminebovine.pdf",width=6, height=4)
pirateplot(formula=GlutamineT~Loading+Time, data=TICsetR, theme=3, inf.b.o =
0,inf.f.o = 0, ylab="TIC Corrected Glutamine*10e9", main="Glutamine
Concentration",ylim=c(0,6.5))
text(x=1,y=2.6,"A")
text(x=2,y=3.1,"AB")
text(x=4,y=2.6,"ABC")
text(x=5,y=5.3,"ABC")
text(x=7,y=3.8,"BC")
text(x=8,y=6.2,"C")
text(x=10,y=4,"ABC")
text(x=11,y=4,"ABC")
```

### TIC_Testing

Molly Piazza

2025-10-13

1) Read in your data set and run dim on it:

```
library(readxl)
TICsetOA <- read_excel("Injury_ML_HeatMap_OA_withTIC.xlsx")
dim(TICsetOA)

## [1] 90 50

tally(Time ~ Label, data = TICsetOA)

##       Label
## Time    C  I  N <NA>
##   0    10 10 10    0
##   1     0 10 10    0
##   4     0 10 10    0
##   24    0 10  9    0
##   <NA>  0  0  0    1

TICsetOA <- TICsetOA %>% mutate(TimeF = factor(Time),
                           Label.Time = str_c(Label, TimeF))
```

#### Just Time 0 observations for three group comparison:

Glutamine

```
TICsetOAR2 <- TICsetOA %>% dplyr::filter(TimeF == "0") %>%
  mutate(GlutamineB = factor(GlutamineB))


glm2 <- glm(GlutamineB ~ Label, data=TICsetOAR2, family = "binomial")

#Can't add donor - causes separation here
Anova(glm2)

## Analysis of Deviance Table (Type II tests)
##
## Response: GlutamineB
##       LR Chisq Df Pr(>Chisq)
## Label  0.29605  2     0.8624

summary(glm2)

##
## Call:
```

```
## glm(formula = GlutamineB ~ Label, family = "binomial", data = TICsetOAR2)
##
## Coefficients:
##              Estimate Std. Error z value Pr(>|z|)
## (Intercept) -8.473e-01  6.901e-01  -1.228     0.22
## LabelI       2.783e-16  9.759e-01   0.000     1.00
## LabelN       4.418e-01  9.449e-01   0.468     0.64
##
## (Dispersion parameter for binomial family taken to be 1)
##
##     Null deviance: 38.191  on 29  degrees of freedom
## Residual deviance: 37.895  on 27  degrees of freedom
## AIC: 43.895
##
## Number of Fisher Scoring iterations: 4
```

```
plot(allEffects(glm2), type = "response", grid = T)
```

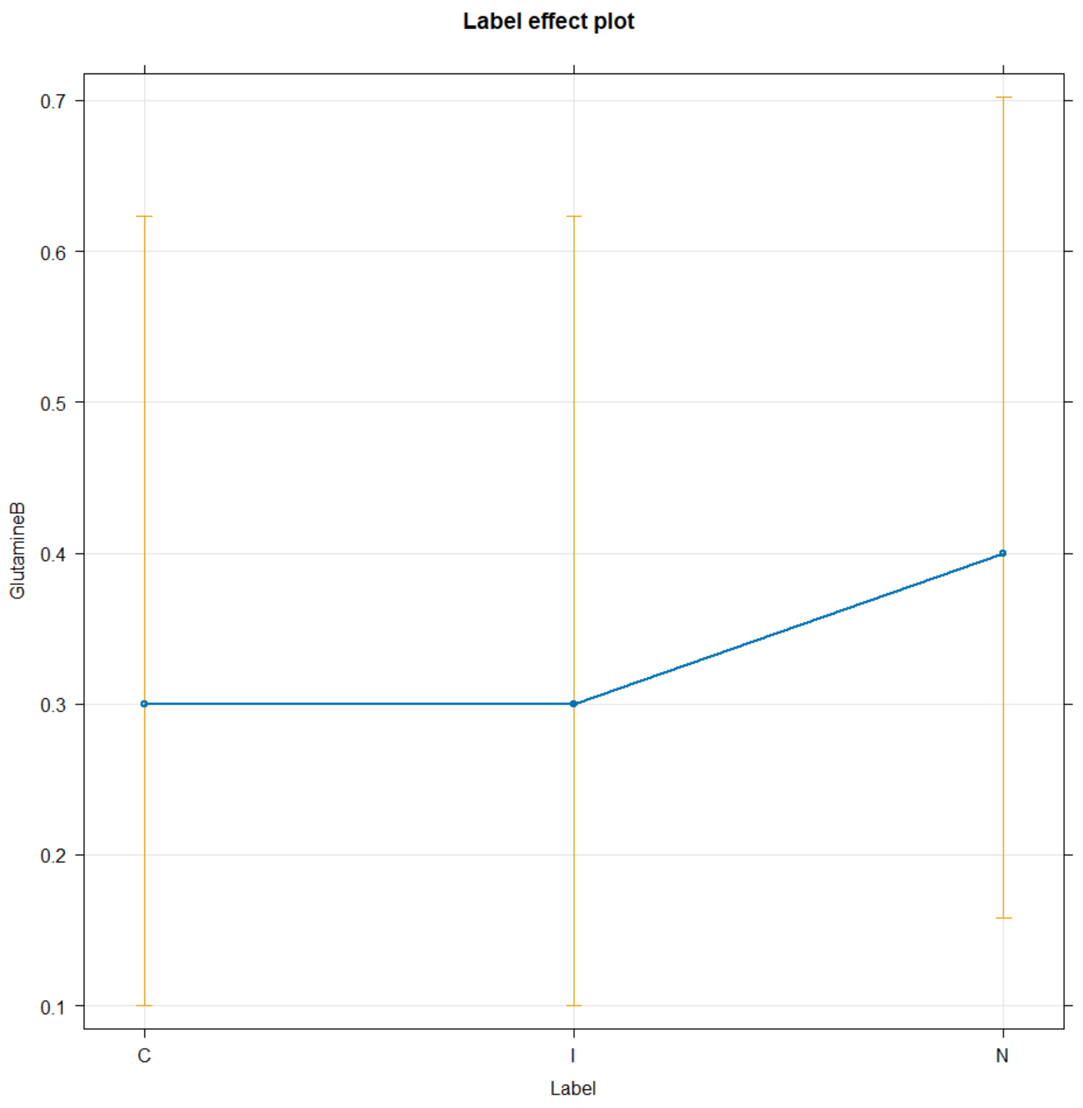

Fructose-6-Phosphate

```r
TICsetOAR2 <- TICsetOA %>% dplyr::filter(TimeF == "0") %>%
  mutate(Fructose6B = factor(Fructose6B))


glm2 <- glm(Fructose6B ~ Label, data=TICsetOAR2, family = "binomial")

#Can't add donor - causes separation here
Anova(glm2)

## Analysis of Deviance Table (Type II tests)
##
## Response: Fructose6B
##       LR Chisq Df Pr(>Chisq)
## Label        0  2          1

summary(glm2)

##
## Call:
## glm(formula = Fructose6B ~ Label, family = "binomial", data = TICsetOAR2)
##
## Coefficients:
##               Estimate Std. Error z value Pr(>|z|)
## (Intercept)  1.386e+00  7.906e-01   1.754   0.0795
## LabelI      -1.574e-16  1.118e+00   0.000   1.0000
## LabelN      -7.448e-16  1.118e+00   0.000   1.0000
##
## (Dispersion parameter for binomial family taken to be 1)
##
##     Null deviance: 30.024  on 29  degrees of freedom
## Residual deviance: 30.024  on 27  degrees of freedom
## AIC: 36.024
##
## Number of Fisher Scoring iterations: 4

plot(allEffects(glm2), type = "response", grid = T)
```

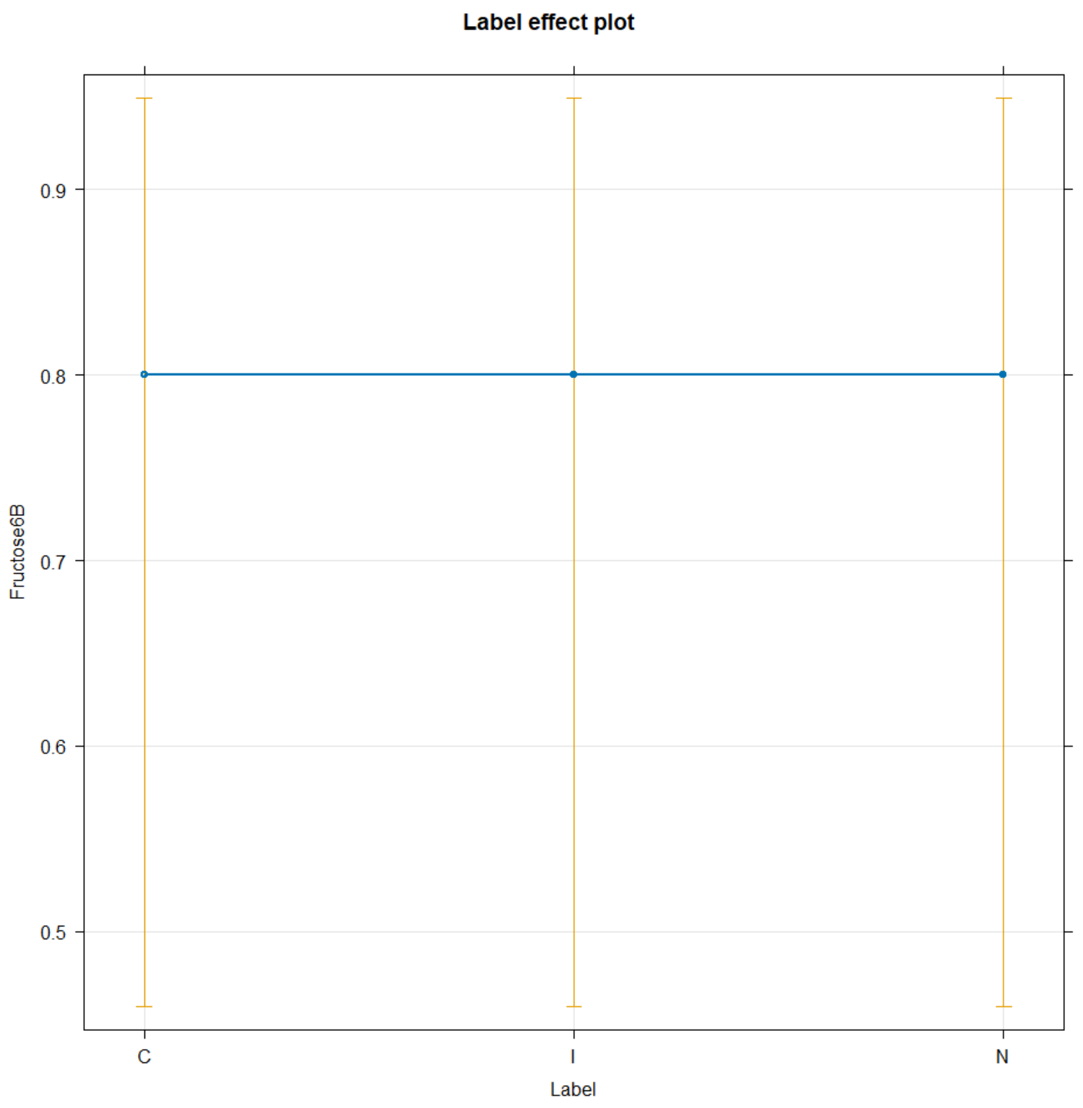


NADH

```
TICsetOAR2 <- TICsetOA %>% dplyr::filter(TimeF == "0") %>%
  mutate(NADHB = factor(NADHB))


glm2 <- glm(NADHB ~ Label, data=TICsetOAR2, family = "binomial")

#Can't add donor - causes separation here
Anova(glm2)

## Analysis of Deviance Table (Type II tests)
##
## Response: NADHB
##       LR Chisq Df Pr(>Chisq)
## Label  0.26928  2      0.874
```

```r
summary(glm2)

##
## Call:
## glm(formula = NADHB ~ Label, family = "binomial", data = TICsetOAR2)
##
## Coefficients:
##             Estimate Std. Error z value Pr(>|z|)
## (Intercept)   0.4055     0.6455   0.628    0.530
## LabelI       -0.4055     0.9037  -0.449    0.654
## LabelN       -0.4055     0.9037  -0.449    0.654
##
## (Dispersion parameter for binomial family taken to be 1)
##
##     Null deviance: 41.455  on 29  degrees of freedom
## Residual deviance: 41.186  on 27  degrees of freedom
## AIC: 47.186
##
## Number of Fisher Scoring iterations: 4

plot(allEffects(glm2), type = "response", grid = T)
```

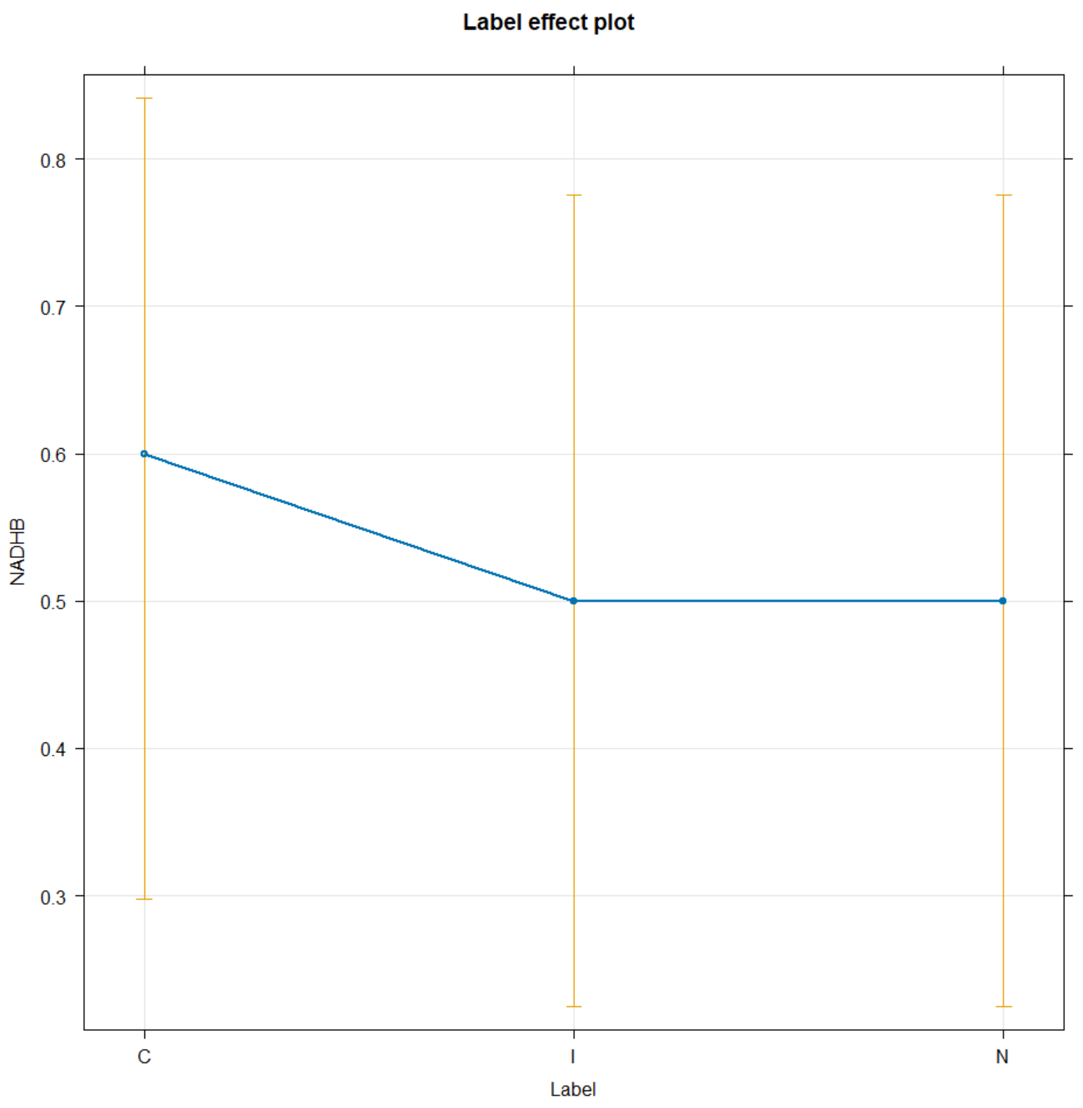


6-Phosphogluconate

```
TICsetOAR2 <- TICsetOA %>% dplyr::filter(TimeF == "0") %>%
  mutate(SixPhosphoB = factor(SixPhosphoB))


glm2 <- glm(SixPhosphoB ~ Label, data=TICsetOAR2, family = "binomial")

#Can't add donor - causes separation here
Anova(glm2)

## Analysis of Deviance Table (Type II tests)
##
## Response: SixPhosphoB
##       LR Chisq Df Pr(>Chisq)
## Label     4.07  2     0.1307
```

```r
summary(glm2)

##
## Call:
## glm(formula = SixPhosphoB ~ Label, family = "binomial", data = TICsetOAR2)
##
## Coefficients:
##             Estimate Std. Error z value Pr(>|z|)
## (Intercept)   0.8473     0.6901   1.228    0.220
## LabelI        1.3499     1.2598   1.072    0.284
## LabelN       -0.8473     0.9361  -0.905    0.365
##
## (Dispersion parameter for binomial family taken to be 1)
##
##     Null deviance: 36.652  on 29  degrees of freedom
## Residual deviance: 32.582  on 27  degrees of freedom
## AIC: 38.582
##
## Number of Fisher Scoring iterations: 4

plot(allEffects(glm2), type = "response", grid = T)
```

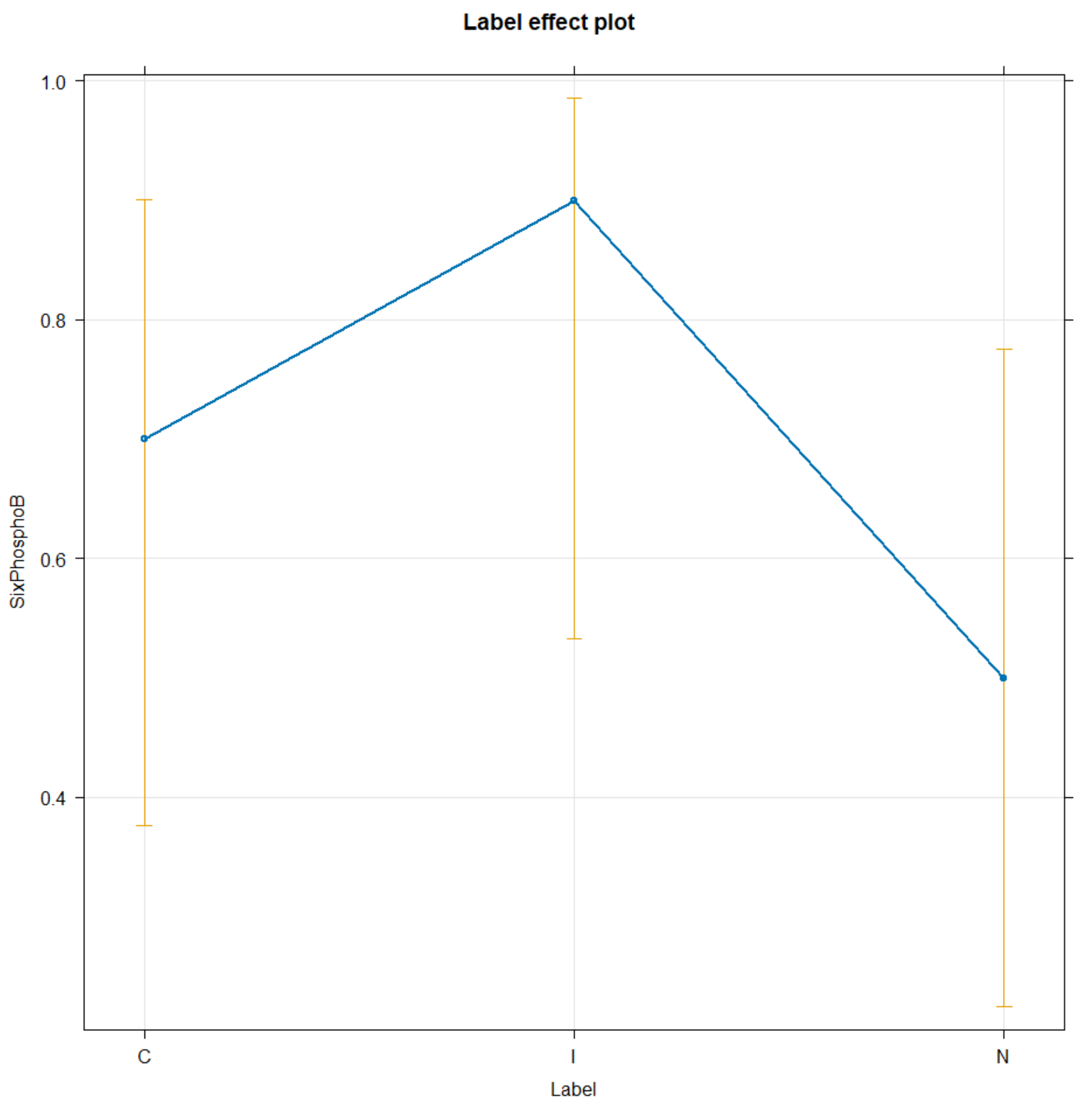


Glucose

```
TICsetOAR2 <- TICsetOA %>% dplyr::filter(TimeF == "0") %>%
  mutate(GlucoseB = factor(GlucoseB))


glm2 <- glm(GlucoseB ~ Label, data=TICsetOAR2, family = "binomial")

#Can't add donor - causes separation here
Anova(glm2)

## Analysis of Deviance Table (Type II tests)
##
## Response: GlucoseB
##       LR Chisq Df Pr(>Chisq)
## Label  0.96629  2     0.6168
```

```r
summary(glm2)

##
## Call:
## glm(formula = GlucoseB ~ Label, family = "binomial", data = TICsetOAR2)
##
## Coefficients:
##             Estimate Std. Error z value Pr(>|z|)
## (Intercept)  -1.3863     0.7906  -1.754   0.0795
## LabelI        0.5390     1.0494   0.514   0.6075
## LabelN        0.9808     1.0206   0.961   0.3365
##
## (Dispersion parameter for binomial family taken to be 1)
##
##     Null deviance: 36.652  on 29  degrees of freedom
## Residual deviance: 35.686  on 27  degrees of freedom
## AIC: 41.686
##
## Number of Fisher Scoring iterations: 4

plot(allEffects(glm2), type = "response", grid = T)
```

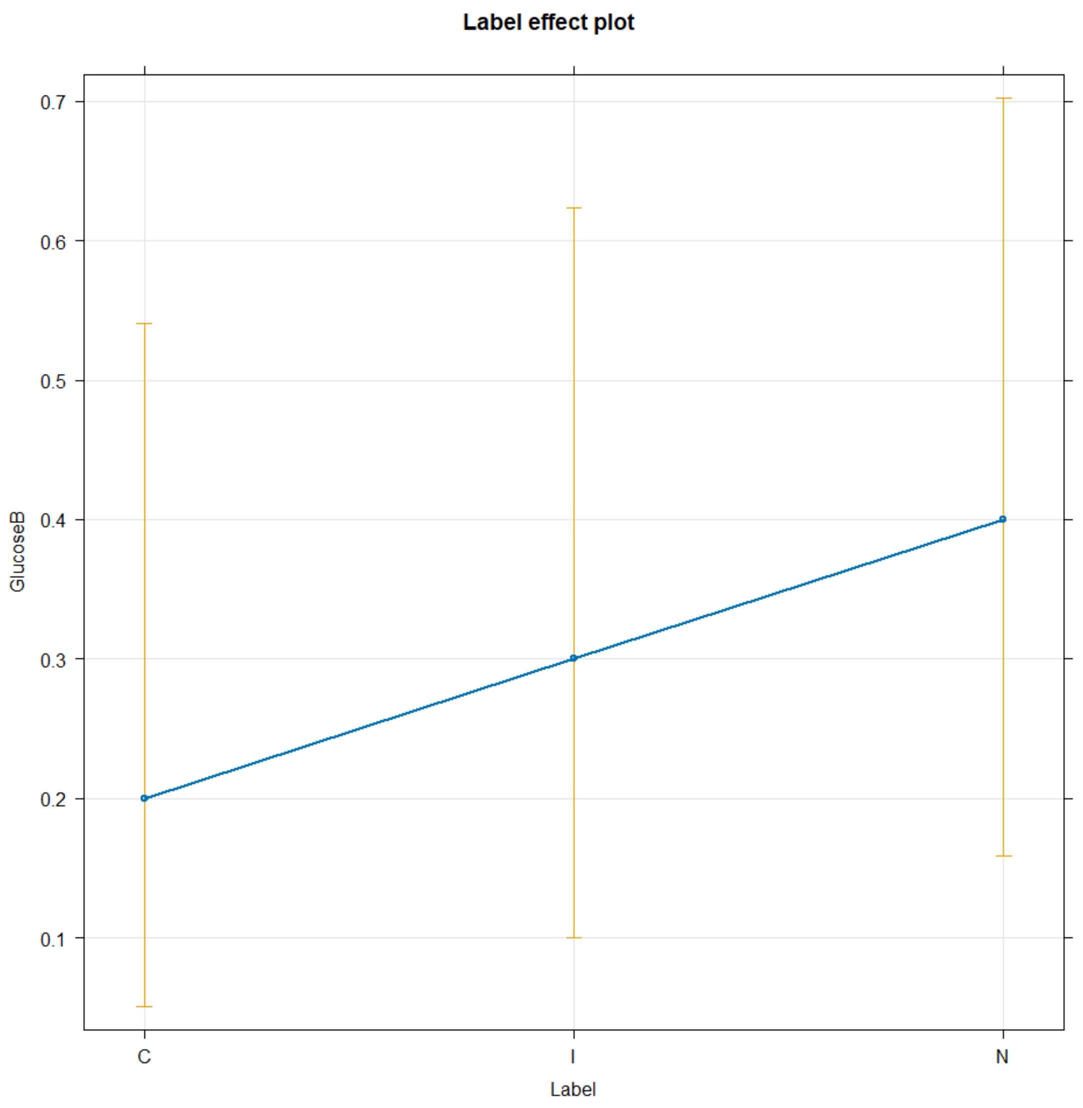


Lactic Acid

```
TICsetOAR2 <- TICsetOA %>% dplyr::filter(TimeF == "0") %>%
  mutate(LacticAcidB = factor(LacticAcidB))


glm2 <- glm(LacticAcidB ~ Label, data=TICsetOAR2, family = "binomial")

#Can't add donor - causes separation here
Anova(glm2)

## Analysis of Deviance Table (Type II tests)
##
## Response: LacticAcidB
##       LR Chisq Df Pr(>Chisq)
## Label  0.29171  2     0.8643
```

```r
summary(glm2)

##
## Call:
## glm(formula = LacticAcidB ~ Label, family = "binomial", data = TICsetOAR2)
##
## Coefficients:
##              Estimate Std. Error z value Pr(>|z|)
## (Intercept) 4.055e-01  6.455e-01   0.628     0.53
## LabelI      1.331e-16  9.129e-01   0.000     1.00
## LabelN      4.418e-01  9.449e-01   0.468     0.64
##
## (Dispersion parameter for binomial family taken to be 1)
##
##     Null deviance: 39.429  on 29  degrees of freedom
## Residual deviance: 39.138  on 27  degrees of freedom
## AIC: 45.138
##
## Number of Fisher Scoring iterations: 4

plot(allEffects(glm2), type = "response", grid = T)
```

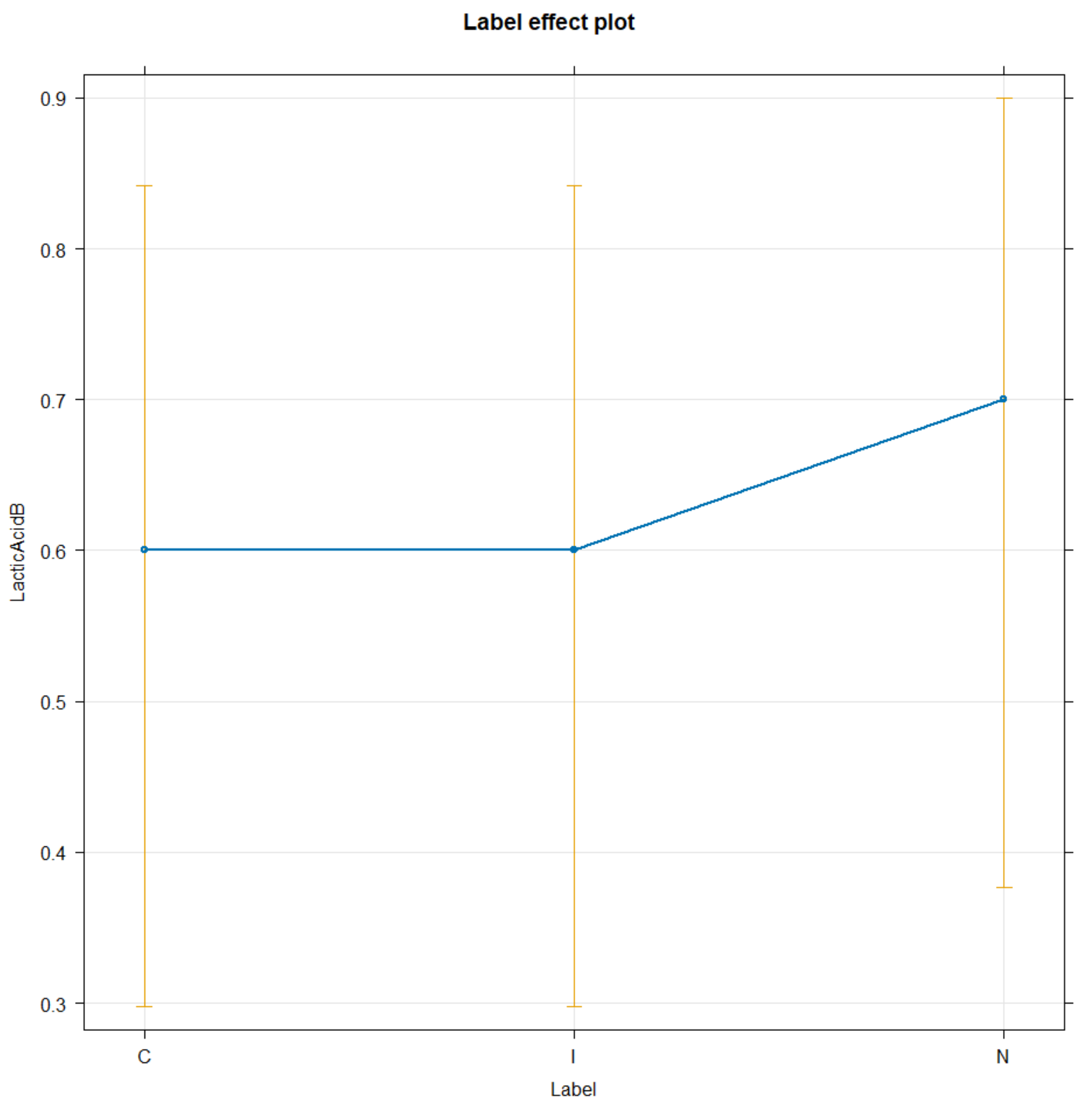


Succinic Acid

```
TICsetOAR2 <- TICsetOA %>% dplyr::filter(TimeF == "0") %>%
  mutate(SuccinicB = factor(SuccinicB))


glm2 <- glm(SuccinicB ~ Label, data=TICsetOAR2, family = "binomial")

#Can't add donor - causes separation here
Anova(glm2)

## Analysis of Deviance Table (Type II tests)
##
## Response: SuccinicB
##       LR Chisq Df Pr(>Chisq)
## Label   2.6264  2      0.269
```

```r
summary(glm2)

##
## Call:
## glm(formula = SuccinicB ~ Label, family = "binomial", data = TICsetOAR2)
##
## Coefficients:
##             Estimate Std. Error z value Pr(>|z|)
## (Intercept)   1.3863     0.7906   1.754   0.0795
## LabelI       -0.9808     1.0206  -0.961   0.3365
## LabelN        0.8109     1.3175   0.615   0.5382
##
## (Dispersion parameter for binomial family taken to be 1)
##
##     Null deviance: 32.596  on 29  degrees of freedom
## Residual deviance: 29.970  on 27  degrees of freedom
## AIC: 35.97
##
## Number of Fisher Scoring iterations: 4

plot(allEffects(glm2), type = "response", grid = T)
```

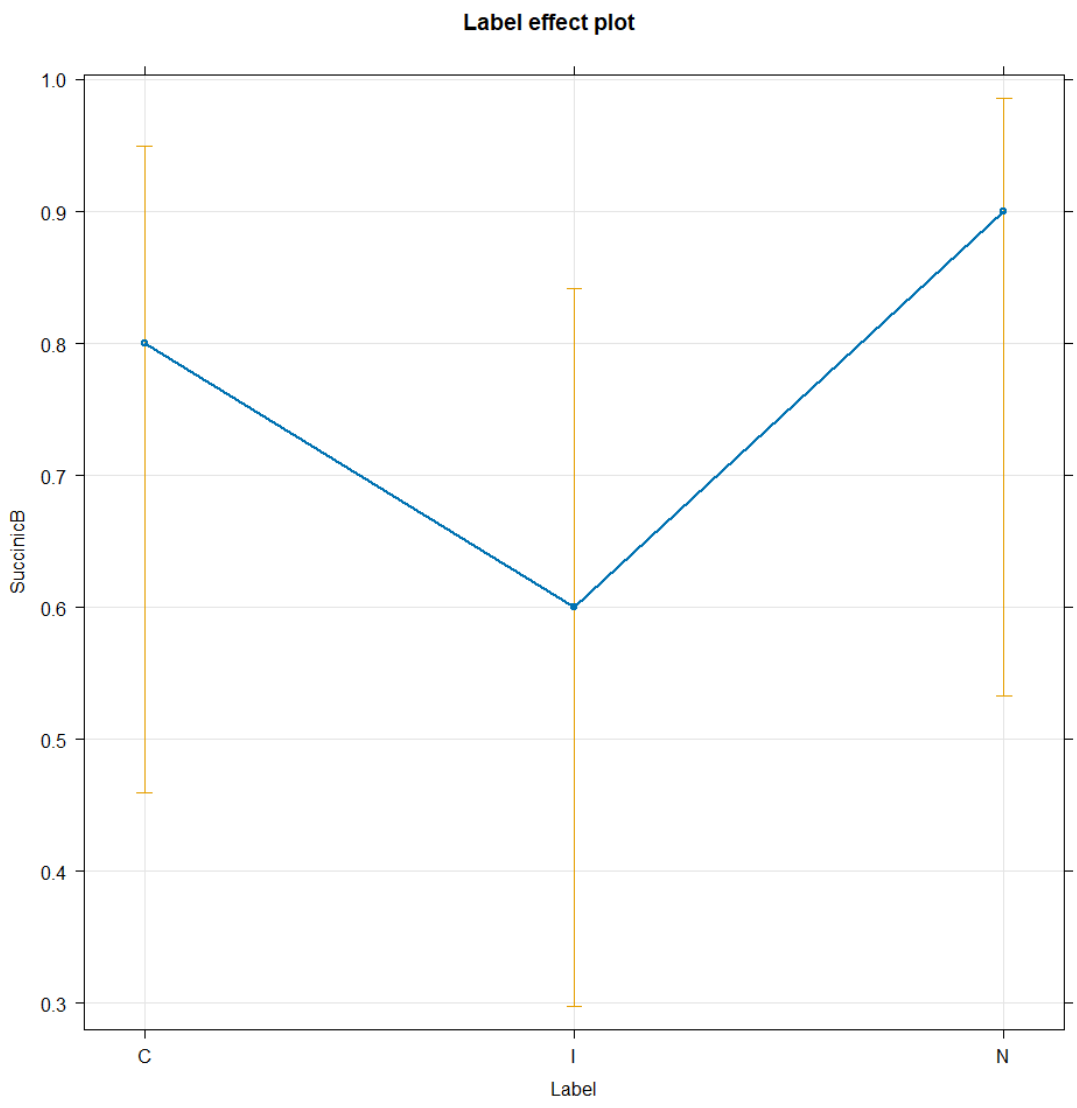


#### Remove control to explore time by I or N interaction:

Loading and Time

Glutamine

```
TICsetOAR <- TICsetOA %>% dplyr::filter(Label != "C") %>%
  mutate(GlutamineB = factor(GlutamineB))

glm3 <- glm(GlutamineB ~ Label+TimeF, data=TICsetOAR, family = "binomial")

#Can't add donor - causes separation here
Anova(glm3)

## Analysis of Deviance Table (Type II tests)
##
## Response: GlutamineB
```

```
##       LR Chisq Df Pr(>Chisq)
## Label  0.68417  1     0.4082
## TimeF  1.57679  3     0.6647
```

```
summary(glm3)
```

```
##
## Call:
## glm(formula = GlutamineB ~ Label + TimeF, family = "binomial",
##     data = TICsetOAR)
##
## Coefficients:
##             Estimate Std. Error z value Pr(>|z|)
## (Intercept)  -0.8396     0.5462  -1.537    0.124
## LabelN        0.4274     0.5189   0.824    0.410
## TimeF1       -0.2305     0.6801  -0.339    0.735
## TimeF4       -0.7741     0.7329  -1.056    0.291
## TimeF24      -0.6978     0.7356  -0.949    0.343
##
## (Dispersion parameter for binomial family taken to be 1)
##
##     Null deviance: 91.491  on 78  degrees of freedom
## Residual deviance: 89.221  on 74  degrees of freedom
## AIC: 99.221
##
## Number of Fisher Scoring iterations: 4
```

```
plot(allEffects(glm3), type = "response", grid = T)
```

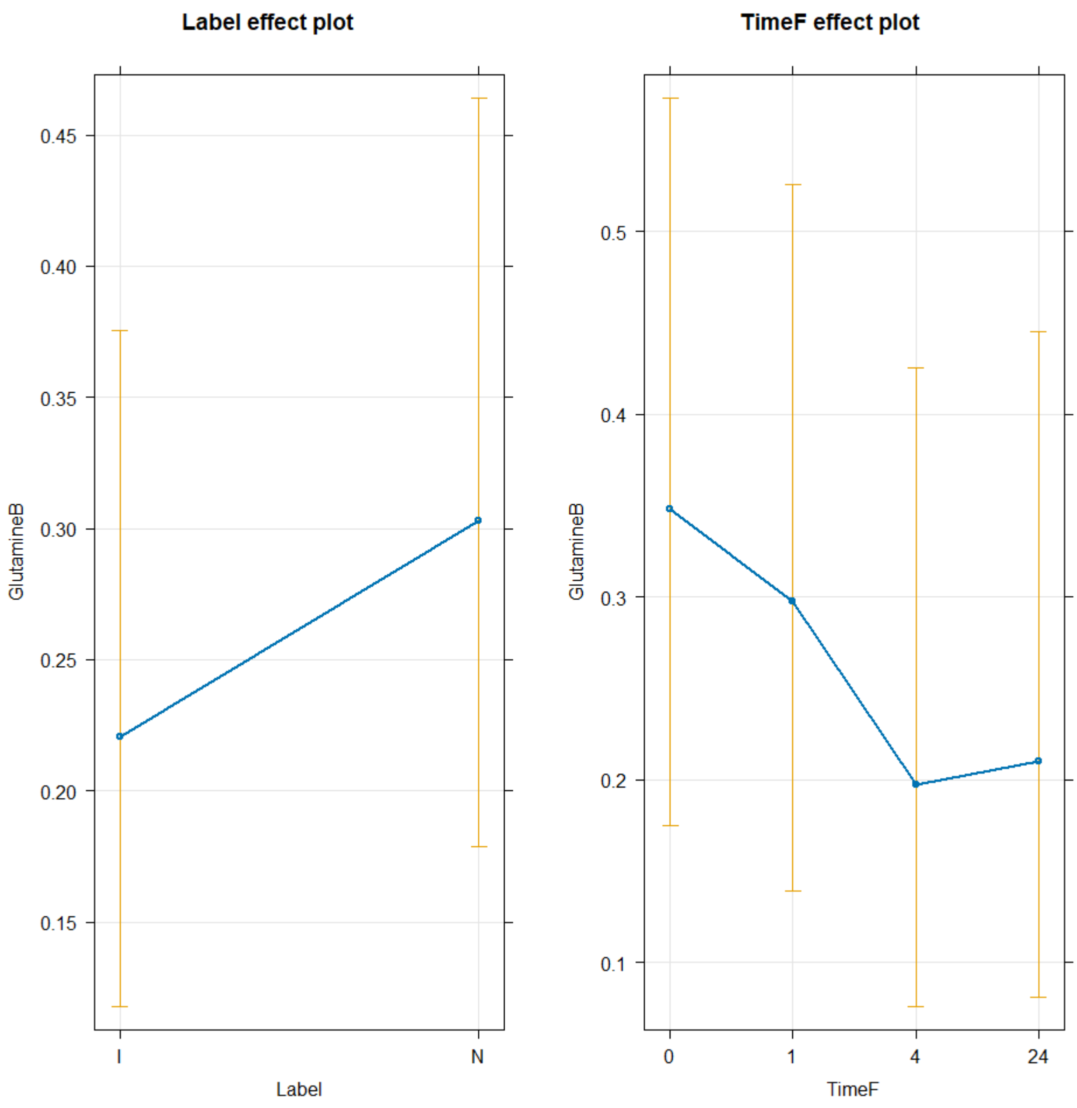


```
TukeyRes2 <- emmeans(glm3, pairwise ~ Label+TimeF, adjust = "tukey")
TukeyRes2

## $emmeans
##  Label TimeF emmean    SE  df asymp.LCL asymp.UCL
##  I     0     -0.840 0.546 Inf     -1.91  0.230937
##  N     0     -0.412 0.530 Inf     -1.45  0.626690
##  I     1     -1.070 0.565 Inf     -2.18  0.038125
##  N     1     -0.643 0.545 Inf     -1.71  0.424694
##  I     4     -1.614 0.633 Inf     -2.85 -0.373421
##  N     4     -1.186 0.605 Inf     -2.37 -0.000687
##  I     24    -1.537 0.629 Inf     -2.77 -0.303615
##  N     24    -1.110 0.614 Inf     -2.31  0.093728
##
## Results are given on the logit (not the response) scale.
## Confidence level used: 0.95
##
```

```
## $contrasts
##  contrast               estimate    SE  df z.ratio p.value
##  I TimeF0 - N TimeF0     -0.4274 0.519 Inf  -0.824  0.9918
##  I TimeF0 - I TimeF1      0.2305 0.680 Inf   0.339  1.0000
##  I TimeF0 - N TimeF1     -0.1968 0.852 Inf  -0.231  1.0000
##  I TimeF0 - I TimeF4      0.7741 0.733 Inf   1.056  0.9655
##  I TimeF0 - N TimeF4      0.3467 0.888 Inf   0.390  0.9999
##  I TimeF0 - I TimeF24     0.6978 0.736 Inf   0.949  0.9812
##  I TimeF0 - N TimeF24     0.2704 0.899 Inf   0.301  1.0000
##  N TimeF0 - I TimeF1      0.6579 0.859 Inf   0.766  0.9948
##  N TimeF0 - N TimeF1      0.2305 0.680 Inf   0.339  1.0000
##  N TimeF0 - I TimeF4      1.2014 0.907 Inf   1.324  0.8900
##  N TimeF0 - N TimeF4      0.7741 0.733 Inf   1.056  0.9655
##  N TimeF0 - I TimeF24     1.1251 0.901 Inf   1.249  0.9172
##  N TimeF0 - N TimeF24     0.6978 0.736 Inf   0.949  0.9812
##  I TimeF1 - N TimeF1     -0.4274 0.519 Inf  -0.824  0.9918
##  I TimeF1 - I TimeF4      0.5435 0.745 Inf   0.729  0.9961
##  I TimeF1 - N TimeF4      0.1162 0.902 Inf   0.129  1.0000
##  I TimeF1 - I TimeF24     0.4672 0.748 Inf   0.625  0.9986
##  I TimeF1 - N TimeF24     0.0399 0.913 Inf   0.044  1.0000
##  N TimeF1 - I TimeF4      0.9709 0.914 Inf   1.062  0.9645
##  N TimeF1 - N TimeF4      0.5435 0.745 Inf   0.729  0.9961
##  N TimeF1 - I TimeF24     0.8946 0.908 Inf   0.985  0.9766
##  N TimeF1 - N TimeF24     0.4672 0.748 Inf   0.625  0.9986
##  I TimeF4 - N TimeF4     -0.4274 0.519 Inf  -0.824  0.9918
##  I TimeF4 - I TimeF24    -0.0763 0.796 Inf  -0.096  1.0000
##  I TimeF4 - N TimeF24    -0.5037 0.959 Inf  -0.525  0.9995
##  N TimeF4 - I TimeF24     0.3511 0.942 Inf   0.373  1.0000
##  N TimeF4 - N TimeF24    -0.0763 0.796 Inf  -0.096  1.0000
##  I TimeF24 - N TimeF24   -0.4274 0.519 Inf  -0.824  0.9918
##
## Results are given on the log odds ratio (not the response) scale.
## P value adjustment: tukey method for comparing a family of 8 estimates
```

```
confint(TukeyRes2)
```

```
## $emmeans
##  Label TimeF emmean    SE  df asymp.LCL asymp.UCL
##  I     0     -0.840 0.546 Inf     -1.91  0.230937
##  N     0     -0.412 0.530 Inf     -1.45  0.626690
##  I     1     -1.070 0.565 Inf     -2.18  0.038125
##  N     1     -0.643 0.545 Inf     -1.71  0.424694
##  I     4     -1.614 0.633 Inf     -2.85 -0.373421
##  N     4     -1.186 0.605 Inf     -2.37 -0.000687
##  I     24    -1.537 0.629 Inf     -2.77 -0.303615
##  N     24    -1.110 0.614 Inf     -2.31  0.093728
##
## Results are given on the logit (not the response) scale.
## Confidence level used: 0.95
##
```

```
## $contrasts
##  contrast               estimate    SE  df asymp.LCL asymp.UCL
##  I TimeF0 - N TimeF0    -0.4274 0.519 Inf     -2.00      1.15
##  I TimeF0 - I TimeF1     0.2305 0.680 Inf     -1.83      2.29
##  I TimeF0 - N TimeF1    -0.1968 0.852 Inf     -2.78      2.39
##  I TimeF0 - I TimeF4     0.7741 0.733 Inf     -1.45      3.00
##  I TimeF0 - N TimeF4     0.3467 0.888 Inf     -2.35      3.04
##  I TimeF0 - I TimeF24    0.6978 0.736 Inf     -1.53      2.93
##  I TimeF0 - N TimeF24    0.2704 0.899 Inf     -2.46      3.00
##  N TimeF0 - I TimeF1     0.6579 0.859 Inf     -1.94      3.26
##  N TimeF0 - N TimeF1     0.2305 0.680 Inf     -1.83      2.29
##  N TimeF0 - I TimeF4     1.2014 0.907 Inf     -1.55      3.95
##  N TimeF0 - N TimeF4     0.7741 0.733 Inf     -1.45      3.00
##  N TimeF0 - I TimeF24    1.1251 0.901 Inf     -1.61      3.86
##  N TimeF0 - N TimeF24    0.6978 0.736 Inf     -1.53      2.93
##  I TimeF1 - N TimeF1    -0.4274 0.519 Inf     -2.00      1.15
##  I TimeF1 - I TimeF4     0.5435 0.745 Inf     -1.71      2.80
##  I TimeF1 - N TimeF4     0.1162 0.902 Inf     -2.62      2.85
##  I TimeF1 - I TimeF24    0.4672 0.748 Inf     -1.80      2.73
##  I TimeF1 - N TimeF24    0.0399 0.913 Inf     -2.73      2.81
##  N TimeF1 - I TimeF4     0.9709 0.914 Inf     -1.80      3.74
##  N TimeF1 - N TimeF4     0.5435 0.745 Inf     -1.71      2.80
##  N TimeF1 - I TimeF24    0.8946 0.908 Inf     -1.86      3.65
##  N TimeF1 - N TimeF24    0.4672 0.748 Inf     -1.80      2.73
##  I TimeF4 - N TimeF4    -0.4274 0.519 Inf     -2.00      1.15
##  I TimeF4 - I TimeF24   -0.0763 0.796 Inf     -2.49      2.34
##  I TimeF4 - N TimeF24   -0.5037 0.959 Inf     -3.41      2.40
##  N TimeF4 - I TimeF24    0.3511 0.942 Inf     -2.50      3.21
##  N TimeF4 - N TimeF24   -0.0763 0.796 Inf     -2.49      2.34
##  I TimeF24 - N TimeF24  -0.4274 0.519 Inf     -2.00      1.15
##
## Results are given on the log odds ratio (not the response) scale.
## Confidence level used: 0.95
## Conf-level adjustment: tukey method for comparing a family of 8 estimates
```

```
plot(TukeyRes2, comparison = T) + coord_flip()
```

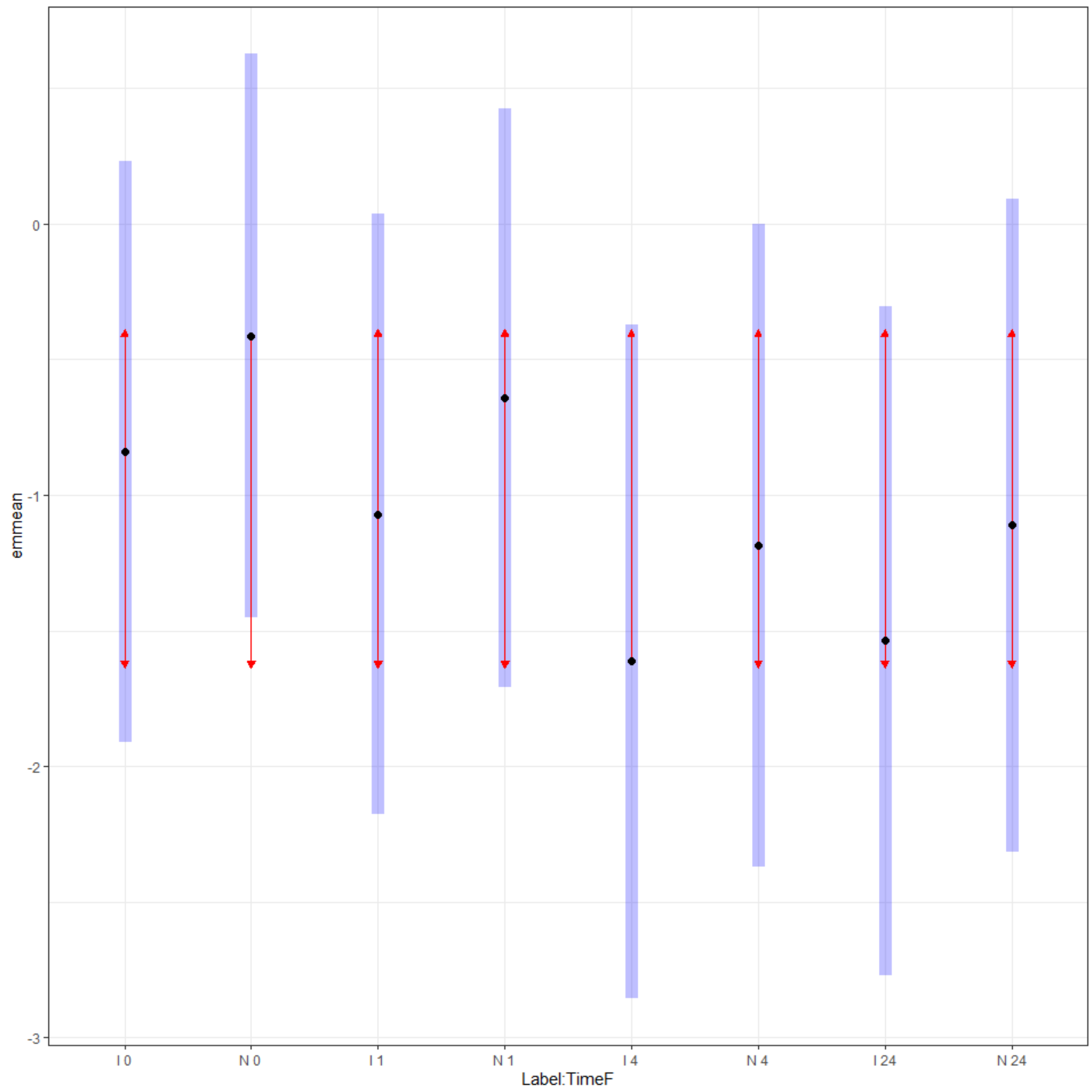


```
multcomp::cld(TukeyRes2, Letters = LETTERS)

##  Label TimeF emmean    SE  df asymp.LCL asymp.UCL .group
##  I     4     -1.614 0.633 Inf     -2.85 -0.373421  A
##  I     24    -1.537 0.629 Inf     -2.77 -0.303615  A
##  N     4     -1.186 0.605 Inf     -2.37 -0.000687  A
##  N     24    -1.110 0.614 Inf     -2.31  0.093728  A
##  I     1     -1.070 0.565 Inf     -2.18  0.038125  A
##  I     0     -0.840 0.546 Inf     -1.91  0.230937  A
##  N     1     -0.643 0.545 Inf     -1.71  0.424694  A
##  N     0     -0.412 0.530 Inf     -1.45  0.626690  A
##
## Results are given on the logit (not the response) scale.
## Confidence level used: 0.95
## Results are given on the log odds ratio (not the response) scale.
## P value adjustment: tukey method for comparing a family of 8 estimates
## significance level used: alpha = 0.05
```

```
## NOTE: If two or more means share the same grouping symbol,
##       then we cannot show them to be different.
##       But we also did not show them to be the same.
```

NADH

```
TICsetOAR <- TICsetOA %>% dplyr::filter(Label != "C") %>%
  mutate(NADHB = factor(NADHB))

glm3 <- glm(NADHB ~ Label+TimeF, data=TICsetOAR, family = "binomial")

#Can't add donor - causes separation here
Anova(glm3)

## Analysis of Deviance Table (Type II tests)
##
## Response: NADHB
##       LR Chisq Df Pr(>Chisq)
## Label   0.1163  1     0.7331
## TimeF   4.3780  3     0.2234

summary(glm3)

##
## Call:
## glm(formula = NADHB ~ Label + TimeF, family = "binomial", data =
TICsetOAR)
##
## Coefficients:
##             Estimate Std. Error z value Pr(>|z|)
## (Intercept) -0.08186    0.50795  -0.161   0.8720
## LabelN       0.16372    0.48033   0.341   0.7332
## TimeF1      -1.38830    0.71646  -1.938   0.0527
## TimeF4      -0.62004    0.64845  -0.956   0.3390
## TimeF24     -0.31467    0.64551  -0.487   0.6259
##
## (Dispersion parameter for binomial family taken to be 1)
##
##     Null deviance: 103.867  on 78  degrees of freedom
## Residual deviance:  99.387  on 74  degrees of freedom
## AIC: 109.39
##
## Number of Fisher Scoring iterations: 4

plot(allEffects(glm3), type = "response", grid = T)
```

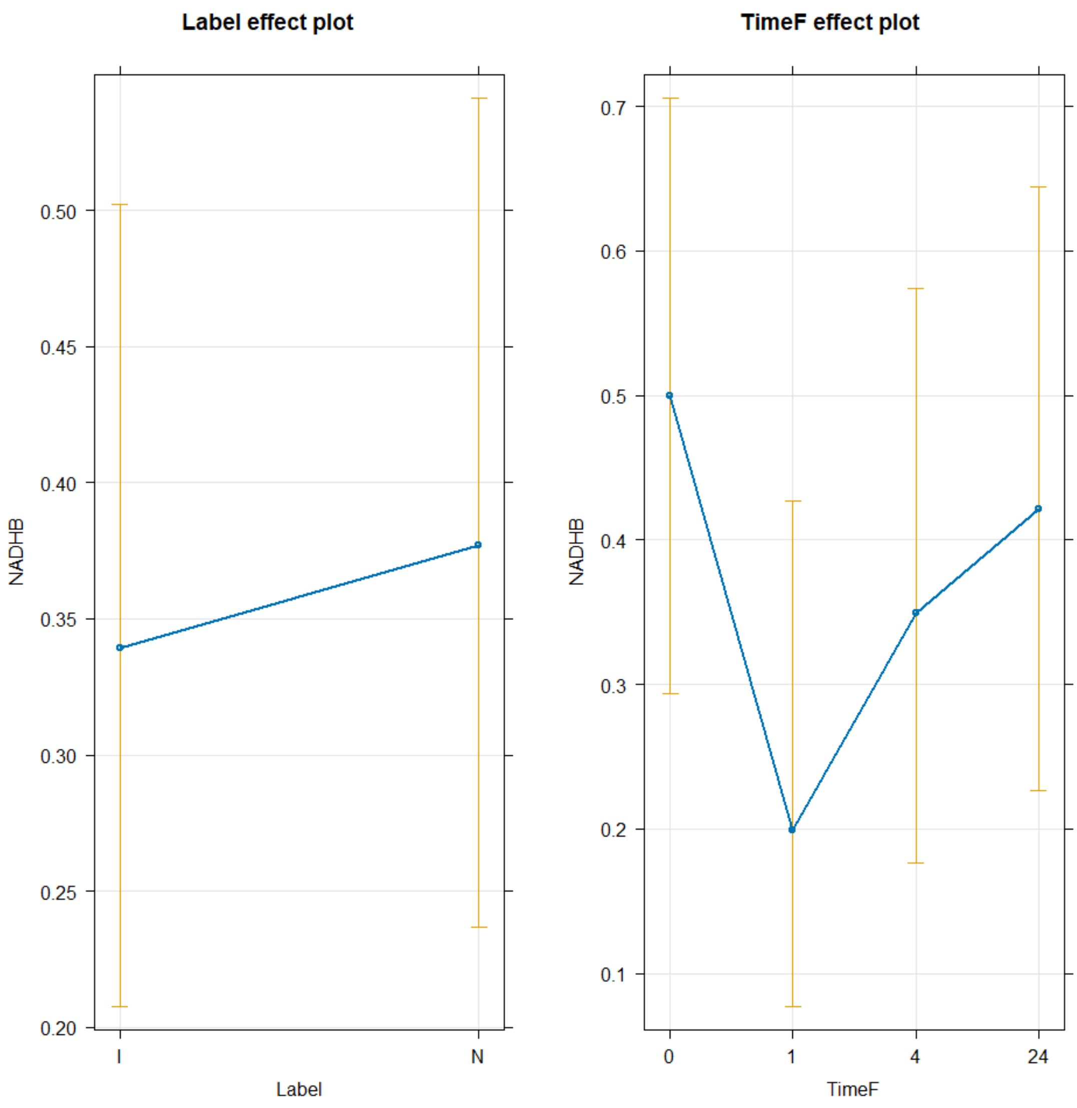


```
TukeyRes2 <- emmeans(glm3, pairwise ~ Label+TimeF, adjust = "tukey")
TukeyRes2
```

```
## $emmeans
##  Label TimeF  emmean    SE  df asymp.LCL asymp.UCL
##  I     0     -0.0819 0.508 Inf    -1.077     0.914
##  N     0      0.0819 0.508 Inf    -0.914     1.077
##  I     1     -1.4702 0.613 Inf    -2.672    -0.268
##  N     1     -1.3064 0.604 Inf    -2.491    -0.122
##  I     4     -0.7019 0.530 Inf    -1.740     0.336
##  N     4     -0.5382 0.524 Inf    -1.566     0.490
##  I     24    -0.3965 0.519 Inf    -1.414     0.621
##  N     24    -0.2328 0.528 Inf    -1.267     0.802
##
## Results are given on the logit (not the response) scale.
## Confidence level used: 0.95
##
```

```
## $contrasts
##  contrast              estimate    SE  df z.ratio p.value
##  I TimeF0 - N TimeF0     -0.164 0.480 Inf  -0.341  1.0000
##  I TimeF0 - I TimeF1      1.388 0.716 Inf   1.938  0.5247
##  I TimeF0 - N TimeF1      1.225 0.856 Inf   1.431  0.8432
##  I TimeF0 - I TimeF4      0.620 0.648 Inf   0.956  0.9803
##  I TimeF0 - N TimeF4      0.456 0.803 Inf   0.568  0.9992
##  I TimeF0 - I TimeF24     0.315 0.646 Inf   0.487  0.9997
##  I TimeF0 - N TimeF24     0.151 0.810 Inf   0.186  1.0000
##  N TimeF0 - I TimeF1      1.552 0.869 Inf   1.786  0.6298
##  N TimeF0 - N TimeF1      1.388 0.716 Inf   1.938  0.5247
##  N TimeF0 - I TimeF4      0.784 0.810 Inf   0.967  0.9790
##  N TimeF0 - N TimeF4      0.620 0.648 Inf   0.956  0.9803
##  N TimeF0 - I TimeF24     0.478 0.799 Inf   0.599  0.9989
##  N TimeF0 - N TimeF24     0.315 0.646 Inf   0.487  0.9997
##  I TimeF1 - N TimeF1     -0.164 0.480 Inf  -0.341  1.0000
##  I TimeF1 - I TimeF4     -0.768 0.730 Inf  -1.052  0.9662
##  I TimeF1 - N TimeF4     -0.932 0.877 Inf  -1.063  0.9644
##  I TimeF1 - I TimeF24    -1.074 0.728 Inf  -1.475  0.8208
##  I TimeF1 - N TimeF24    -1.237 0.884 Inf  -1.400  0.8575
##  N TimeF1 - I TimeF4     -0.605 0.871 Inf  -0.694  0.9972
##  N TimeF1 - N TimeF4     -0.768 0.730 Inf  -1.052  0.9662
##  N TimeF1 - I TimeF24    -0.910 0.860 Inf  -1.058  0.9652
##  N TimeF1 - N TimeF24    -1.074 0.728 Inf  -1.475  0.8208
##  I TimeF4 - N TimeF4     -0.164 0.480 Inf  -0.341  1.0000
##  I TimeF4 - I TimeF24    -0.305 0.661 Inf  -0.462  0.9998
##  I TimeF4 - N TimeF24    -0.469 0.826 Inf  -0.568  0.9992
##  N TimeF4 - I TimeF24    -0.142 0.808 Inf  -0.175  1.0000
##  N TimeF4 - N TimeF24    -0.305 0.661 Inf  -0.462  0.9998
##  I TimeF24 - N TimeF24   -0.164 0.480 Inf  -0.341  1.0000
##
## Results are given on the log odds ratio (not the response) scale.
## P value adjustment: tukey method for comparing a family of 8 estimates
```

```r
confint(TukeyRes2)
```

```
## $emmeans
##  Label TimeF  emmean    SE  df asymp.LCL asymp.UCL
##  I     0     -0.0819 0.508 Inf    -1.077     0.914
##  N     0      0.0819 0.508 Inf    -0.914     1.077
##  I     1     -1.4702 0.613 Inf    -2.672    -0.268
##  N     1     -1.3064 0.604 Inf    -2.491    -0.122
##  I     4     -0.7019 0.530 Inf    -1.740     0.336
##  N     4     -0.5382 0.524 Inf    -1.566     0.490
##  I     24    -0.3965 0.519 Inf    -1.414     0.621
##  N     24    -0.2328 0.528 Inf    -1.267     0.802
##
## Results are given on the logit (not the response) scale.
## Confidence level used: 0.95
##
```

```
## $contrasts
##  contrast              estimate    SE  df asymp.LCL asymp.UCL
##  I TimeF0 - N TimeF0     -0.164 0.480 Inf    -1.620      1.29
##  I TimeF0 - I TimeF1      1.388 0.716 Inf    -0.783      3.56
##  I TimeF0 - N TimeF1      1.225 0.856 Inf    -1.370      3.82
##  I TimeF0 - I TimeF4      0.620 0.648 Inf    -1.345      2.59
##  I TimeF0 - N TimeF4      0.456 0.803 Inf    -1.979      2.89
##  I TimeF0 - I TimeF24     0.315 0.646 Inf    -1.642      2.27
##  I TimeF0 - N TimeF24     0.151 0.810 Inf    -2.305      2.61
##  N TimeF0 - I TimeF1      1.552 0.869 Inf    -1.082      4.19
##  N TimeF0 - N TimeF1      1.388 0.716 Inf    -0.783      3.56
##  N TimeF0 - I TimeF4      0.784 0.810 Inf    -1.673      3.24
##  N TimeF0 - N TimeF4      0.620 0.648 Inf    -1.345      2.59
##  N TimeF0 - I TimeF24     0.478 0.799 Inf    -1.943      2.90
##  N TimeF0 - N TimeF24     0.315 0.646 Inf    -1.642      2.27
##  I TimeF1 - N TimeF1     -0.164 0.480 Inf    -1.620      1.29
##  I TimeF1 - I TimeF4     -0.768 0.730 Inf    -2.981      1.44
##  I TimeF1 - N TimeF4     -0.932 0.877 Inf    -3.590      1.73
##  I TimeF1 - I TimeF24    -1.074 0.728 Inf    -3.279      1.13
##  I TimeF1 - N TimeF24    -1.237 0.884 Inf    -3.916      1.44
##  N TimeF1 - I TimeF4     -0.605 0.871 Inf    -3.243      2.03
##  N TimeF1 - N TimeF4     -0.768 0.730 Inf    -2.981      1.44
##  N TimeF1 - I TimeF24    -0.910 0.860 Inf    -3.517      1.70
##  N TimeF1 - N TimeF24    -1.074 0.728 Inf    -3.279      1.13
##  I TimeF4 - N TimeF4     -0.164 0.480 Inf    -1.620      1.29
##  I TimeF4 - I TimeF24    -0.305 0.661 Inf    -2.308      1.70
##  I TimeF4 - N TimeF24    -0.469 0.826 Inf    -2.972      2.03
##  N TimeF4 - I TimeF24    -0.142 0.808 Inf    -2.590      2.31
##  N TimeF4 - N TimeF24    -0.305 0.661 Inf    -2.308      1.70
##  I TimeF24 - N TimeF24   -0.164 0.480 Inf    -1.620      1.29
##
## Results are given on the log odds ratio (not the response) scale.
## Confidence level used: 0.95
## Conf-level adjustment: tukey method for comparing a family of 8 estimates
```

```
plot(TukeyRes2, comparison = T) + coord_flip()
```

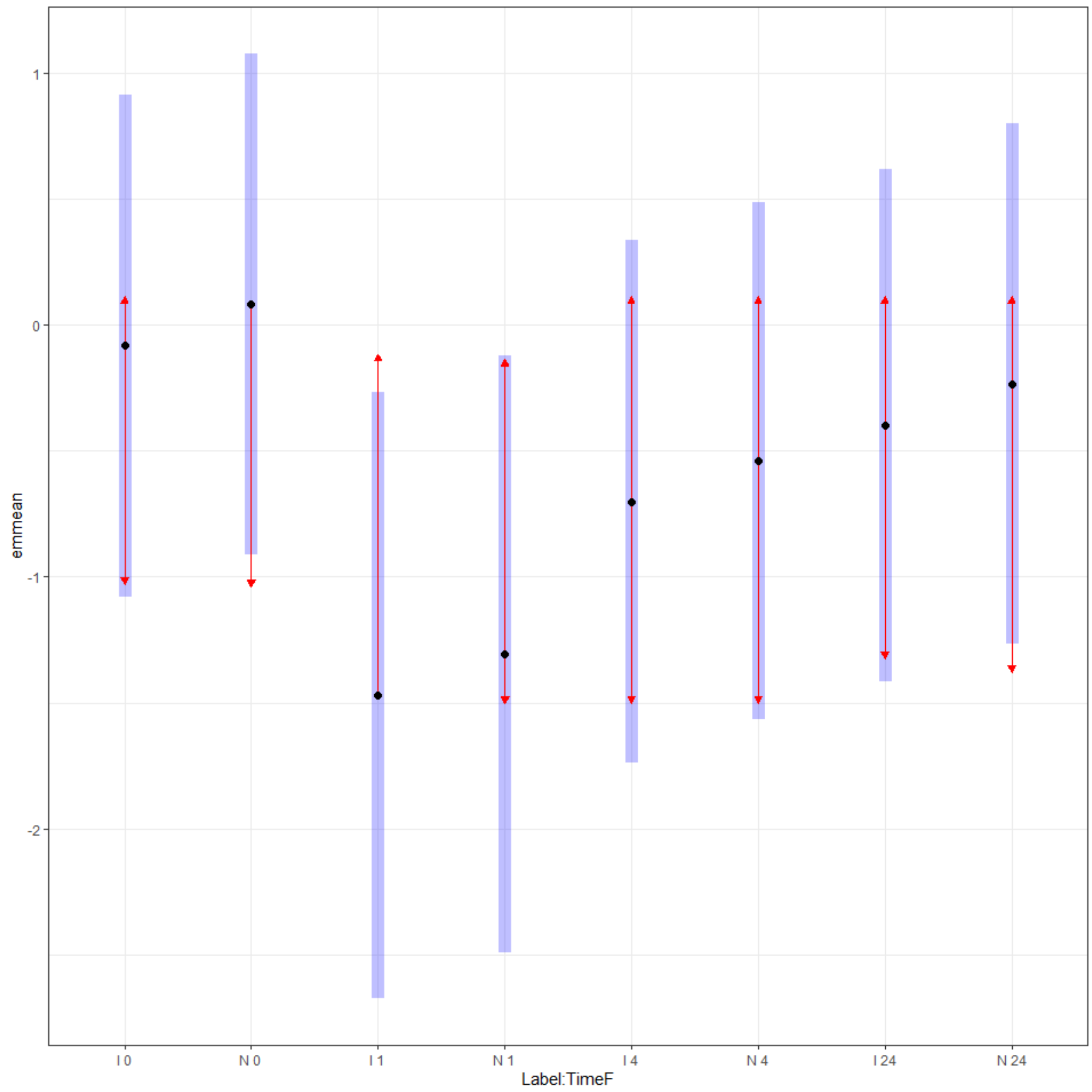


```
multcomp::cld(TukeyRes2, Letters = LETTERS)

##  Label TimeF  emmean    SE  df asymp.LCL asymp.UCL .group
##  I     1     -1.4702 0.613 Inf    -2.672    -0.268  A
##  N     1     -1.3064 0.604 Inf    -2.491    -0.122  A
##  I     4     -0.7019 0.530 Inf    -1.740     0.336  A
##  N     4     -0.5382 0.524 Inf    -1.566     0.490  A
##  I     24    -0.3965 0.519 Inf    -1.414     0.621  A
##  N     24    -0.2328 0.528 Inf    -1.267     0.802  A
##  I     0     -0.0819 0.508 Inf    -1.077     0.914  A
##  N     0      0.0819 0.508 Inf    -0.914     1.077  A
##
## Results are given on the logit (not the response) scale.
## Confidence level used: 0.95
## Results are given on the log odds ratio (not the response) scale.
## P value adjustment: tukey method for comparing a family of 8 estimates
## significance level used: alpha = 0.05
```

```
## NOTE: If two or more means share the same grouping symbol,
##       then we cannot show them to be different.
##       But we also did not show them to be the same.
```

6 Phosphogluconate

```
TICsetOAR <- TICsetOA %>% dplyr::filter(Label != "C") %>%
  mutate(SixPhosphoB = factor(SixPhosphoB))

glm3 <- glm(SixPhosphoB ~ Label+TimeF, data=TICsetOAR, family = "binomial")

#Can't add donor - causes separation here
Anova(glm3)
```

```
## Analysis of Deviance Table (Type II tests)
##
## Response: SixPhosphoB
##       LR Chisq Df Pr(>Chisq)
## Label  0.15688  1     0.6920
## TimeF  2.13842  3     0.5442
```

```
summary(glm3)
```

```
##
## Call:
## glm(formula = SixPhosphoB ~ Label + TimeF, family = "binomial",
##     data = TICsetOAR)
##
## Coefficients:
##             Estimate Std. Error z value Pr(>|z|)
## (Intercept)  0.75583    0.53845   1.404    0.160
## LabelN       0.18641    0.47100   0.396    0.692
## TimeF1      -0.44270    0.66882  -0.662    0.508
## TimeF4      -0.84904    0.66260  -1.281    0.200
## TimeF24     -0.06934    0.69477  -0.100    0.921
##
## (Dispersion parameter for binomial family taken to be 1)
##
##     Null deviance: 104.90  on 78  degrees of freedom
## Residual deviance: 102.62  on 74  degrees of freedom
## AIC: 112.62
##
## Number of Fisher Scoring iterations: 4
```

```
plot(allEffects(glm3), type = "response", grid = T)
```

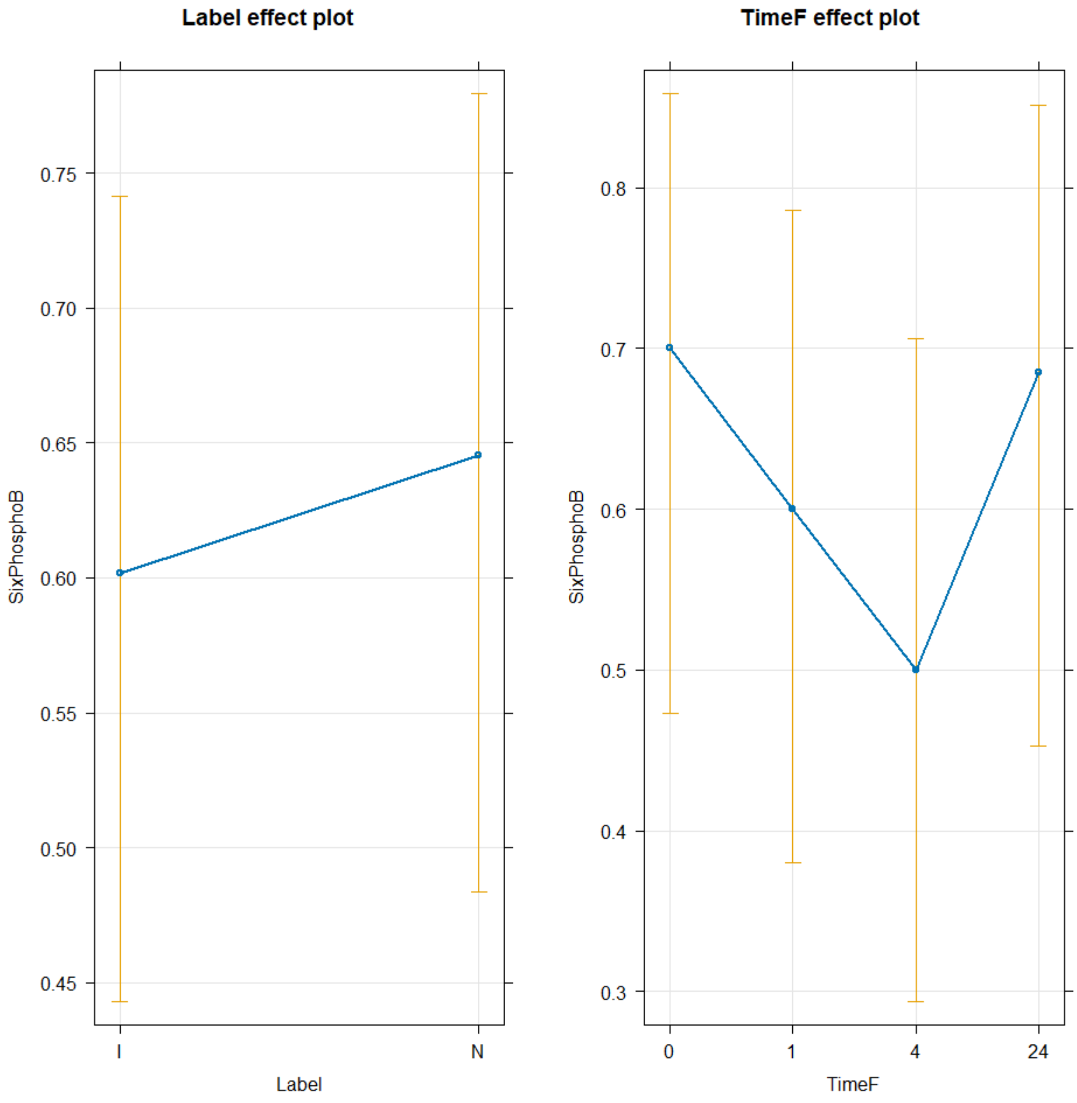


```
TukeyRes2 <- emmeans(glm3, pairwise ~ Label+TimeF, adjust = "tukey")
TukeyRes2

## $emmeans
##  Label TimeF  emmean    SE  df asymp.LCL asymp.UCL
##  I     0      0.7558 0.538 Inf    -0.300     1.811
##  N     0      0.9422 0.546 Inf    -0.128     2.013
##  I     1      0.3131 0.512 Inf    -0.690     1.317
##  N     1      0.4995 0.516 Inf    -0.512     1.511
##  I     4     -0.0932 0.506 Inf    -1.085     0.898
##  N     4      0.0932 0.506 Inf    -0.898     1.085
##  I     24     0.6865 0.539 Inf    -0.369     1.742
##  N     24     0.8729 0.556 Inf    -0.218     1.963
##
## Results are given on the logit (not the response) scale.
## Confidence level used: 0.95
##
```

```
## $contrasts
##  contrast              estimate    SE  df z.ratio p.value
##  I TimeF0 - N TimeF0    -0.1864 0.471 Inf  -0.396  0.9999
##  I TimeF0 - I TimeF1     0.4427 0.669 Inf   0.662  0.9979
##  I TimeF0 - N TimeF1     0.2563 0.815 Inf   0.314  1.0000
##  I TimeF0 - I TimeF4     0.8490 0.663 Inf   1.281  0.9060
##  I TimeF0 - N TimeF4     0.6626 0.808 Inf   0.820  0.9920
##  I TimeF0 - I TimeF24    0.0693 0.695 Inf   0.100  1.0000
##  I TimeF0 - N TimeF24   -0.1171 0.846 Inf  -0.138  1.0000
##  N TimeF0 - I TimeF1     0.6291 0.821 Inf   0.767  0.9947
##  N TimeF0 - N TimeF1     0.4427 0.669 Inf   0.662  0.9979
##  N TimeF0 - I TimeF4     1.0354 0.818 Inf   1.266  0.9114
##  N TimeF0 - N TimeF4     0.8490 0.663 Inf   1.281  0.9060
##  N TimeF0 - I TimeF24    0.2557 0.833 Inf   0.307  1.0000
##  N TimeF0 - N TimeF24    0.0693 0.695 Inf   0.100  1.0000
##  I TimeF1 - N TimeF1    -0.1864 0.471 Inf  -0.396  0.9999
##  I TimeF1 - I TimeF4     0.4063 0.640 Inf   0.635  0.9984
##  I TimeF1 - N TimeF4     0.2199 0.792 Inf   0.278  1.0000
##  I TimeF1 - I TimeF24   -0.3734 0.673 Inf  -0.555  0.9993
##  I TimeF1 - N TimeF24   -0.5598 0.831 Inf  -0.674  0.9977
##  N TimeF1 - I TimeF4     0.5927 0.797 Inf   0.744  0.9956
##  N TimeF1 - N TimeF4     0.4063 0.640 Inf   0.635  0.9984
##  N TimeF1 - I TimeF24   -0.1870 0.812 Inf  -0.230  1.0000
##  N TimeF1 - N TimeF24   -0.3734 0.673 Inf  -0.555  0.9993
##  I TimeF4 - N TimeF4    -0.1864 0.471 Inf  -0.396  0.9999
##  I TimeF4 - I TimeF24   -0.7797 0.667 Inf  -1.169  0.9407
##  I TimeF4 - N TimeF24   -0.9661 0.828 Inf  -1.166  0.9414
##  N TimeF4 - I TimeF24   -0.5933 0.805 Inf  -0.737  0.9959
##  N TimeF4 - N TimeF24   -0.7797 0.667 Inf  -1.169  0.9407
##  I TimeF24 - N TimeF24  -0.1864 0.471 Inf  -0.396  0.9999
##
## Results are given on the log odds ratio (not the response) scale.
## P value adjustment: tukey method for comparing a family of 8 estimates
```

```
confint(TukeyRes2)
```

```
## $emmeans
##  Label TimeF  emmean    SE  df asymp.LCL asymp.UCL
##  I     0      0.7558 0.538 Inf    -0.300     1.811
##  N     0      0.9422 0.546 Inf    -0.128     2.013
##  I     1      0.3131 0.512 Inf    -0.690     1.317
##  N     1      0.4995 0.516 Inf    -0.512     1.511
##  I     4     -0.0932 0.506 Inf    -1.085     0.898
##  N     4      0.0932 0.506 Inf    -0.898     1.085
##  I     24     0.6865 0.539 Inf    -0.369     1.742
##  N     24     0.8729 0.556 Inf    -0.218     1.963
##
## Results are given on the logit (not the response) scale.
## Confidence level used: 0.95
##
```

```
## $contrasts
##  contrast              estimate    SE  df asymp.LCL asymp.UCL
##  I TimeF0 - N TimeF0    -0.1864 0.471 Inf     -1.61      1.24
##  I TimeF0 - I TimeF1     0.4427 0.669 Inf     -1.58      2.47
##  I TimeF0 - N TimeF1     0.2563 0.815 Inf     -2.22      2.73
##  I TimeF0 - I TimeF4     0.8490 0.663 Inf     -1.16      2.86
##  I TimeF0 - N TimeF4     0.6626 0.808 Inf     -1.79      3.11
##  I TimeF0 - I TimeF24    0.0693 0.695 Inf     -2.04      2.18
##  I TimeF0 - N TimeF24   -0.1171 0.846 Inf     -2.68      2.45
##  N TimeF0 - I TimeF1     0.6291 0.821 Inf     -1.86      3.12
##  N TimeF0 - N TimeF1     0.4427 0.669 Inf     -1.58      2.47
##  N TimeF0 - I TimeF4     1.0354 0.818 Inf     -1.44      3.51
##  N TimeF0 - N TimeF4     0.8490 0.663 Inf     -1.16      2.86
##  N TimeF0 - I TimeF24    0.2557 0.833 Inf     -2.27      2.78
##  N TimeF0 - N TimeF24    0.0693 0.695 Inf     -2.04      2.18
##  I TimeF1 - N TimeF1    -0.1864 0.471 Inf     -1.61      1.24
##  I TimeF1 - I TimeF4     0.4063 0.640 Inf     -1.53      2.35
##  I TimeF1 - N TimeF4     0.2199 0.792 Inf     -2.18      2.62
##  I TimeF1 - I TimeF24   -0.3734 0.673 Inf     -2.41      1.67
##  I TimeF1 - N TimeF24   -0.5598 0.831 Inf     -3.08      1.96
##  N TimeF1 - I TimeF4     0.5927 0.797 Inf     -1.82      3.01
##  N TimeF1 - N TimeF4     0.4063 0.640 Inf     -1.53      2.35
##  N TimeF1 - I TimeF24   -0.1870 0.812 Inf     -2.65      2.27
##  N TimeF1 - N TimeF24   -0.3734 0.673 Inf     -2.41      1.67
##  I TimeF4 - N TimeF4    -0.1864 0.471 Inf     -1.61      1.24
##  I TimeF4 - I TimeF24   -0.7797 0.667 Inf     -2.80      1.24
##  I TimeF4 - N TimeF24   -0.9661 0.828 Inf     -3.48      1.54
##  N TimeF4 - I TimeF24   -0.5933 0.805 Inf     -3.03      1.85
##  N TimeF4 - N TimeF24   -0.7797 0.667 Inf     -2.80      1.24
##  I TimeF24 - N TimeF24  -0.1864 0.471 Inf     -1.61      1.24
##
## Results are given on the log odds ratio (not the response) scale.
## Confidence level used: 0.95
## Conf-level adjustment: tukey method for comparing a family of 8 estimates
```

```
plot(TukeyRes2, comparison = T) + coord_flip()
```

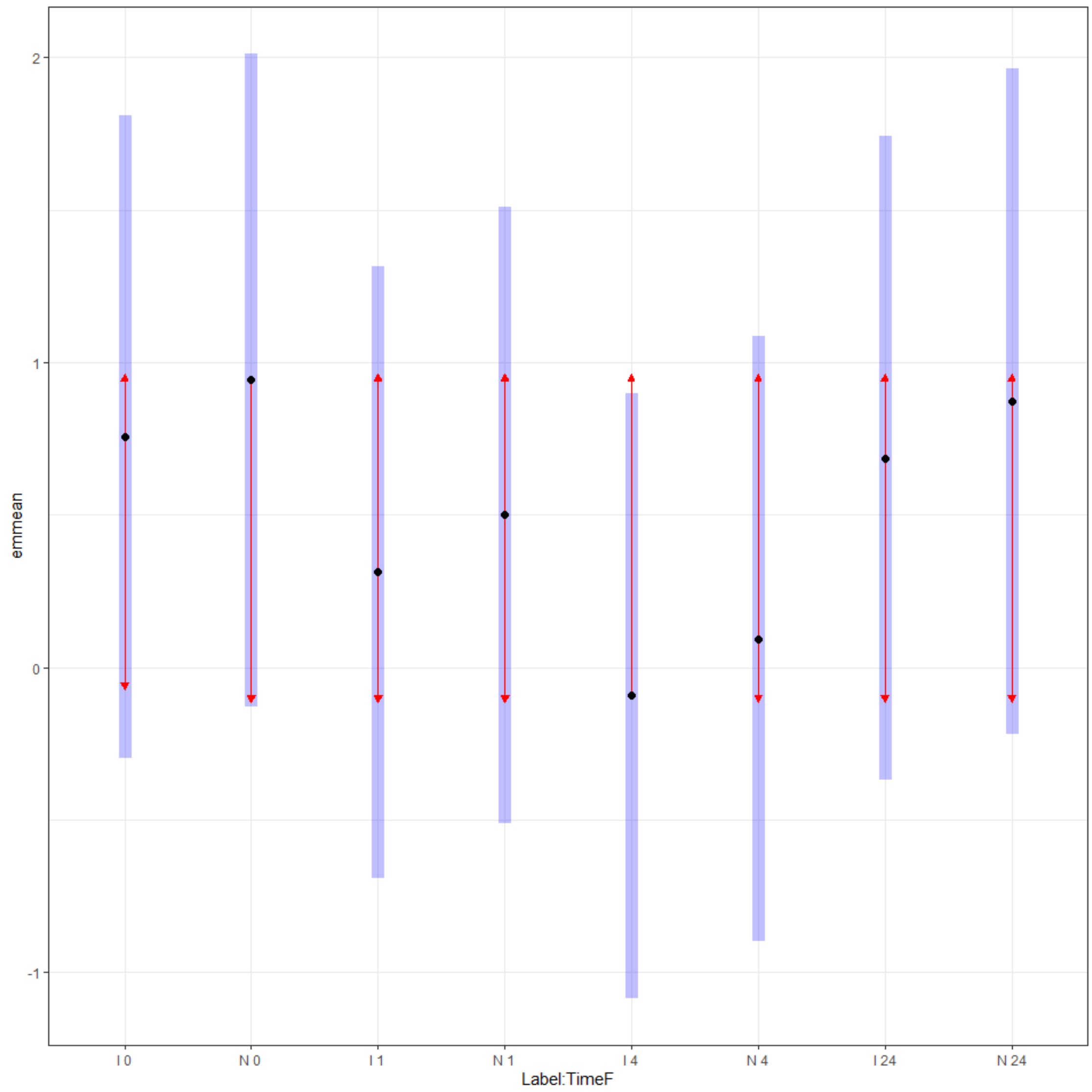


```
multcomp::cld(TukeyRes2, Letters = LETTERS)

##  Label TimeF  emmean    SE  df asymp.LCL asymp.UCL .group
##  I     4     -0.0932 0.506 Inf    -1.085     0.898  A
##  N     4      0.0932 0.506 Inf    -0.898     1.085  A
##  I     1      0.3131 0.512 Inf    -0.690     1.317  A
##  N     1      0.4995 0.516 Inf    -0.512     1.511  A
##  I     24     0.6865 0.539 Inf    -0.369     1.742  A
##  I     0      0.7558 0.538 Inf    -0.300     1.811  A
##  N     24     0.8729 0.556 Inf    -0.218     1.963  A
##  N     0      0.9422 0.546 Inf    -0.128     2.013  A
##
## Results are given on the logit (not the response) scale.
## Confidence level used: 0.95
## Results are given on the log odds ratio (not the response) scale.
## P value adjustment: tukey method for comparing a family of 8 estimates
## significance level used: alpha = 0.05
```

```
## NOTE: If two or more means share the same grouping symbol,
##       then we cannot show them to be different.
##       But we also did not show them to be the same.
```

Fructose-6-Phosphate

```
TICsetOAR <- TICsetOA %>% dplyr::filter(Label != "C") %>%
  mutate(Fructose6B = factor(Fructose6B))

glm3 <- glm(Fructose6B ~ Label+TimeF, data=TICsetOAR, family = "binomial")

#Can't add donor - causes separation here
Anova(glm3)

## Analysis of Deviance Table (Type II tests)
##
## Response: Fructose6B
##       LR Chisq Df Pr(>Chisq)
## Label   4.2361  1    0.03957
## TimeF   6.0730  3    0.10811

summary(glm3)

##
## Call:
## glm(formula = Fructose6B ~ Label + TimeF, family = "binomial",
##     data = TICsetOAR)
##
## Coefficients:
##             Estimate Std. Error z value Pr(>|z|)
## (Intercept)   1.9789     0.6605   2.996  0.00273
## LabelN       -1.0273     0.5104  -2.013  0.04414
## TimeF1       -1.6794     0.7409  -2.267  0.02341
## TimeF4       -0.5646     0.7595  -0.743  0.45722
## TimeF24      -0.9190     0.7541  -1.219  0.22297
##
## (Dispersion parameter for binomial family taken to be 1)
##
##     Null deviance: 102.723  on 78  degrees of freedom
## Residual deviance:  92.748  on 74  degrees of freedom
## AIC: 102.75
##
## Number of Fisher Scoring iterations: 4

plot(allEffects(glm3), type = "response", grid = T)
```

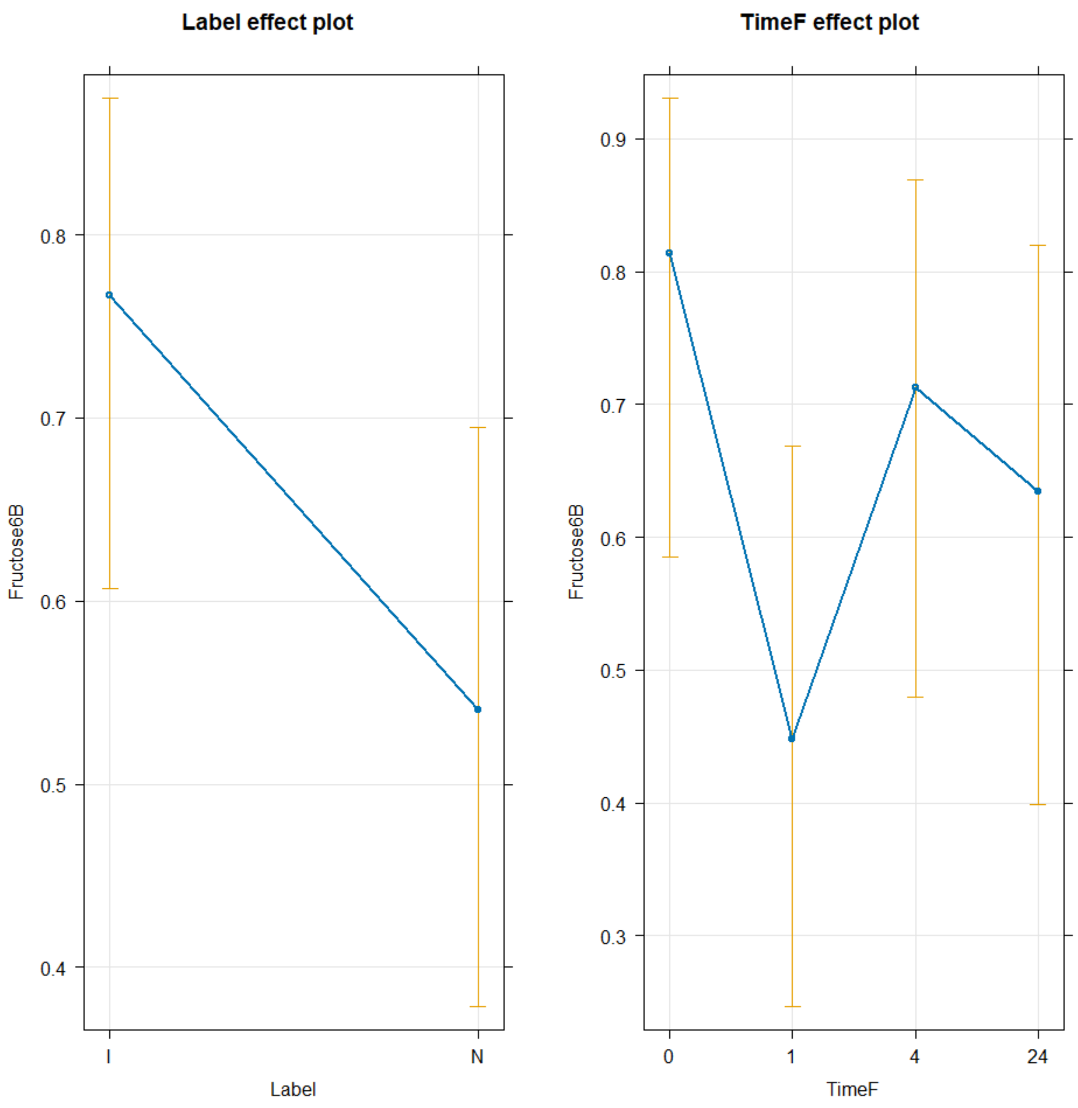


```
TukeyRes2 <- emmeans(glm3, pairwise ~ Label+TimeF, adjust = "tukey")
TukeyRes2

## $emmeans
##  Label TimeF  emmean    SE  df asymp.LCL asymp.UCL
##  I     0      1.9789 0.660 Inf    0.6844     3.273
##  N     0      0.9516 0.597 Inf   -0.2184     2.122
##  I     1      0.2995 0.523 Inf   -0.7261     1.325
##  N     1     -0.7278 0.536 Inf   -1.7792     0.324
##  I     4      1.4143 0.589 Inf    0.2606     2.568
##  N     4      0.3870 0.540 Inf   -0.6718     1.446
##  I     24     1.0599 0.563 Inf   -0.0428     2.163
##  N     24     0.0326 0.543 Inf   -1.0325     1.098
##
## Results are given on the logit (not the response) scale.
## Confidence level used: 0.95
##
```

```
## $contrasts
##  contrast              estimate    SE  df z.ratio p.value
##  I TimeF0 - N TimeF0     1.0273 0.510 Inf   2.013  0.4731
##  I TimeF0 - I TimeF1     1.6794 0.741 Inf   2.267  0.3122
##  I TimeF0 - N TimeF1     2.7067 0.950 Inf   2.848  0.0836
##  I TimeF0 - I TimeF4     0.5646 0.759 Inf   0.743  0.9957
##  I TimeF0 - N TimeF4     1.5919 0.929 Inf   1.714  0.6780
##  I TimeF0 - I TimeF24    0.9190 0.754 Inf   1.219  0.9267
##  I TimeF0 - N TimeF24    1.9463 0.942 Inf   2.066  0.4375
##  N TimeF0 - I TimeF1     0.6521 0.846 Inf   0.771  0.9946
##  N TimeF0 - N TimeF1     1.6794 0.741 Inf   2.267  0.3122
##  N TimeF0 - I TimeF4    -0.4627 0.901 Inf  -0.513  0.9996
##  N TimeF0 - N TimeF4     0.5646 0.759 Inf   0.743  0.9957
##  N TimeF0 - I TimeF24   -0.1083 0.878 Inf  -0.123  1.0000
##  N TimeF0 - N TimeF24    0.9190 0.754 Inf   1.219  0.9267
##  I TimeF1 - N TimeF1     1.0273 0.510 Inf   2.013  0.4731
##  I TimeF1 - I TimeF4    -1.1148 0.686 Inf  -1.624  0.7358
##  I TimeF1 - N TimeF4    -0.0875 0.814 Inf  -0.107  1.0000
##  I TimeF1 - I TimeF24   -0.7604 0.676 Inf  -1.125  0.9516
##  I TimeF1 - N TimeF24    0.2669 0.826 Inf   0.323  1.0000
##  N TimeF1 - I TimeF4    -2.1421 0.895 Inf  -2.395  0.2438
##  N TimeF1 - N TimeF4    -1.1148 0.686 Inf  -1.624  0.7358
##  N TimeF1 - I TimeF24   -1.7877 0.868 Inf  -2.061  0.4408
##  N TimeF1 - N TimeF24   -0.7604 0.676 Inf  -1.125  0.9516
##  I TimeF4 - N TimeF4     1.0273 0.510 Inf   2.013  0.4731
##  I TimeF4 - I TimeF24    0.3544 0.702 Inf   0.505  0.9996
##  I TimeF4 - N TimeF24    1.3817 0.887 Inf   1.558  0.7754
##  N TimeF4 - I TimeF24   -0.6729 0.848 Inf  -0.793  0.9935
##  N TimeF4 - N TimeF24    0.3544 0.702 Inf   0.505  0.9996
##  I TimeF24 - N TimeF24   1.0273 0.510 Inf   2.013  0.4731
##
## Results are given on the log odds ratio (not the response) scale.
## P value adjustment: tukey method for comparing a family of 8 estimates

confint(TukeyRes2)

## $emmeans
##  Label TimeF  emmean    SE  df asymp.LCL asymp.UCL
##  I     0      1.9789 0.660 Inf    0.6844     3.273
##  N     0      0.9516 0.597 Inf   -0.2184     2.122
##  I     1      0.2995 0.523 Inf   -0.7261     1.325
##  N     1     -0.7278 0.536 Inf   -1.7792     0.324
##  I     4      1.4143 0.589 Inf    0.2606     2.568
##  N     4      0.3870 0.540 Inf   -0.6718     1.446
##  I     24     1.0599 0.563 Inf   -0.0428     2.163
##  N     24     0.0326 0.543 Inf   -1.0325     1.098
##
## Results are given on the logit (not the response) scale.
## Confidence level used: 0.95
##
```

```
## $contrasts
##  contrast               estimate    SE  df asymp.LCL asymp.UCL
##  I TimeF0 - N TimeF0     1.0273 0.510 Inf    -0.520     2.574
##  I TimeF0 - I TimeF1     1.6794 0.741 Inf    -0.566     3.925
##  I TimeF0 - N TimeF1     2.7067 0.950 Inf    -0.174     5.587
##  I TimeF0 - I TimeF4     0.5646 0.759 Inf    -1.737     2.866
##  I TimeF0 - N TimeF4     1.5919 0.929 Inf    -1.223     4.407
##  I TimeF0 - I TimeF24    0.9190 0.754 Inf    -1.367     3.205
##  I TimeF0 - N TimeF24    1.9463 0.942 Inf    -0.910     4.802
##  N TimeF0 - I TimeF1     0.6521 0.846 Inf    -1.912     3.216
##  N TimeF0 - N TimeF1     1.6794 0.741 Inf    -0.566     3.925
##  N TimeF0 - I TimeF4    -0.4627 0.901 Inf    -3.194     2.269
##  N TimeF0 - N TimeF4     0.5646 0.759 Inf    -1.737     2.866
##  N TimeF0 - I TimeF24   -0.1083 0.878 Inf    -2.769     2.552
##  N TimeF0 - N TimeF24    0.9190 0.754 Inf    -1.367     3.205
##  I TimeF1 - N TimeF1     1.0273 0.510 Inf    -0.520     2.574
##  I TimeF1 - I TimeF4    -1.1148 0.686 Inf    -3.195     0.966
##  I TimeF1 - N TimeF4    -0.0875 0.814 Inf    -2.555     2.380
##  I TimeF1 - I TimeF24   -0.7604 0.676 Inf    -2.809     1.289
##  I TimeF1 - N TimeF24    0.2669 0.826 Inf    -2.237     2.771
##  N TimeF1 - I TimeF4    -2.1421 0.895 Inf    -4.853     0.569
##  N TimeF1 - N TimeF4    -1.1148 0.686 Inf    -3.195     0.966
##  N TimeF1 - I TimeF24   -1.7877 0.868 Inf    -4.417     0.842
##  N TimeF1 - N TimeF24   -0.7604 0.676 Inf    -2.809     1.289
##  I TimeF4 - N TimeF4     1.0273 0.510 Inf    -0.520     2.574
##  I TimeF4 - I TimeF24    0.3544 0.702 Inf    -1.773     2.482
##  I TimeF4 - N TimeF24    1.3817 0.887 Inf    -1.306     4.070
##  N TimeF4 - I TimeF24   -0.6729 0.848 Inf    -3.244     1.898
##  N TimeF4 - N TimeF24    0.3544 0.702 Inf    -1.773     2.482
##  I TimeF24 - N TimeF24   1.0273 0.510 Inf    -0.520     2.574
##
## Results are given on the log odds ratio (not the response) scale.
## Confidence level used: 0.95
## Conf-level adjustment: tukey method for comparing a family of 8 estimates
```

```
plot(TukeyRes2, comparison = T) + coord_flip()
```

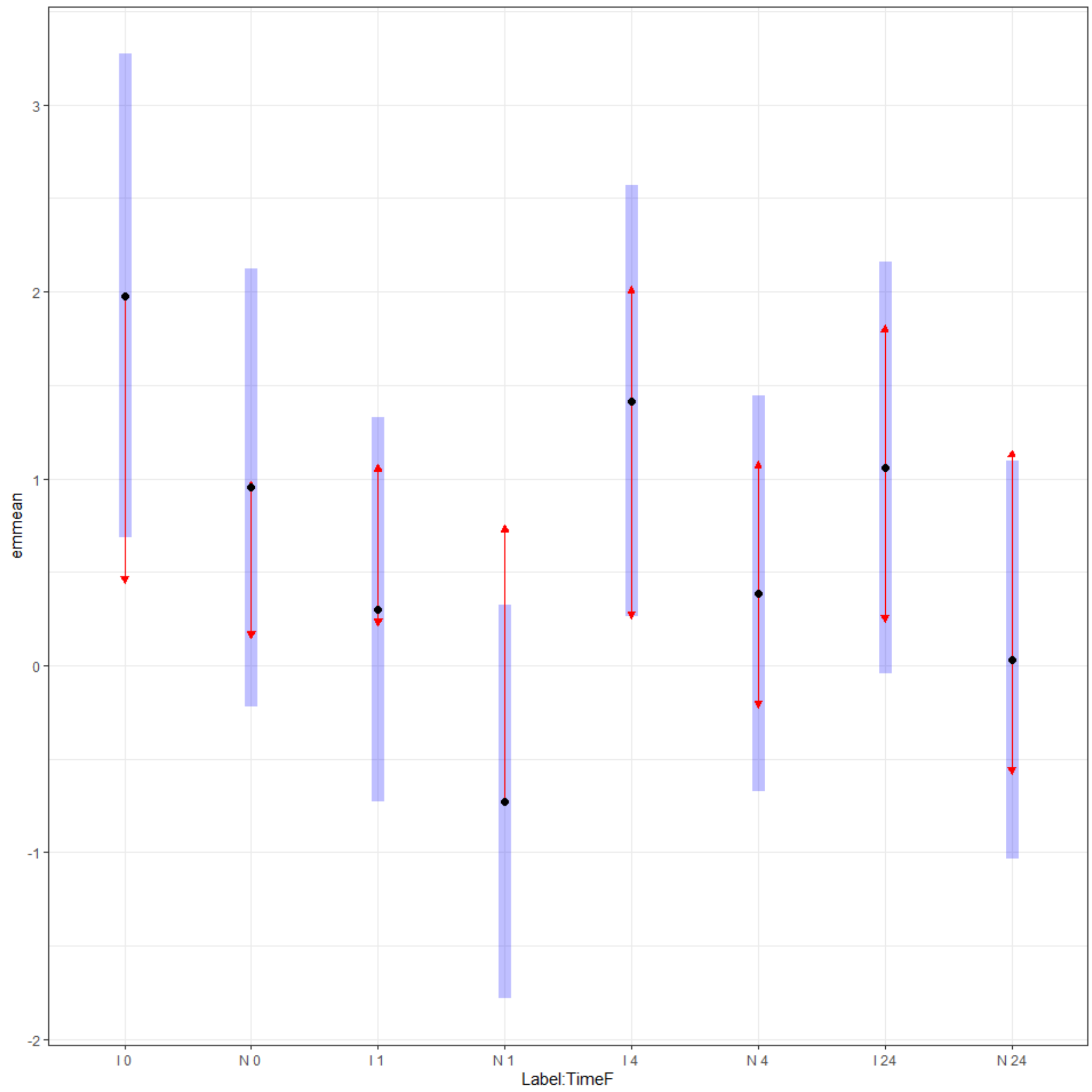


```
multcomp::cld(TukeyRes2, Letters = LETTERS)

##  Label TimeF  emmean    SE  df asymp.LCL asymp.UCL .group
##  N     1     -0.7278 0.536 Inf   -1.7792     0.324  A
##  N     24     0.0326 0.543 Inf   -1.0325     1.098  A
##  I     1      0.2995 0.523 Inf   -0.7261     1.325  A
##  N     4      0.3870 0.540 Inf   -0.6718     1.446  A
##  N     0      0.9516 0.597 Inf   -0.2184     2.122  A
##  I     24     1.0599 0.563 Inf   -0.0428     2.163  A
##  I     4      1.4143 0.589 Inf    0.2606     2.568  A
##  I     0      1.9789 0.660 Inf    0.6844     3.273  A
##
## Results are given on the logit (not the response) scale.
## Confidence level used: 0.95
## Results are given on the log odds ratio (not the response) scale.
## P value adjustment: tukey method for comparing a family of 8 estimates
## significance level used: alpha = 0.05
```

```
## NOTE: If two or more means share the same grouping symbol,
##       then we cannot show them to be different.
##       But we also did not show them to be the same.
```

Glucose

```r
TICsetOAR <- TICsetOA %>% dplyr::filter(Label != "C") %>%
  mutate(GlucoseB = factor(GlucoseB))

glm3 <- glm(GlucoseB ~ Label+TimeF, data=TICsetOAR, family = "binomial")

#Can't add donor - causes separation here
Anova(glm3)
```

```
## Analysis of Deviance Table (Type II tests)
##
## Response: GlucoseB
##       LR Chisq Df Pr(>Chisq)
## Label   1.1262  1     0.2886
## TimeF   0.5087  3     0.9170
```

```r
summary(glm3)
```

```
##
## Call:
## glm(formula = GlucoseB ~ Label + TimeF, family = "binomial",
##     data = TICsetOAR)
##
## Coefficients:
##             Estimate Std. Error z value Pr(>|z|)
## (Intercept)  -0.8913     0.5431  -1.641    0.101
## LabelN        0.5239     0.4966   1.055    0.291
## TimeF1       -0.2317     0.6817  -0.340    0.734
## TimeF4       -0.4864     0.7025  -0.692    0.489
## TimeF24      -0.1426     0.6859  -0.208    0.835
##
## (Dispersion parameter for binomial family taken to be 1)
##
##     Null deviance: 97.020  on 78  degrees of freedom
## Residual deviance: 95.399  on 74  degrees of freedom
## AIC: 105.4
##
## Number of Fisher Scoring iterations: 4
```

```r
plot(allEffects(glm3), type = "response", grid = T)
```

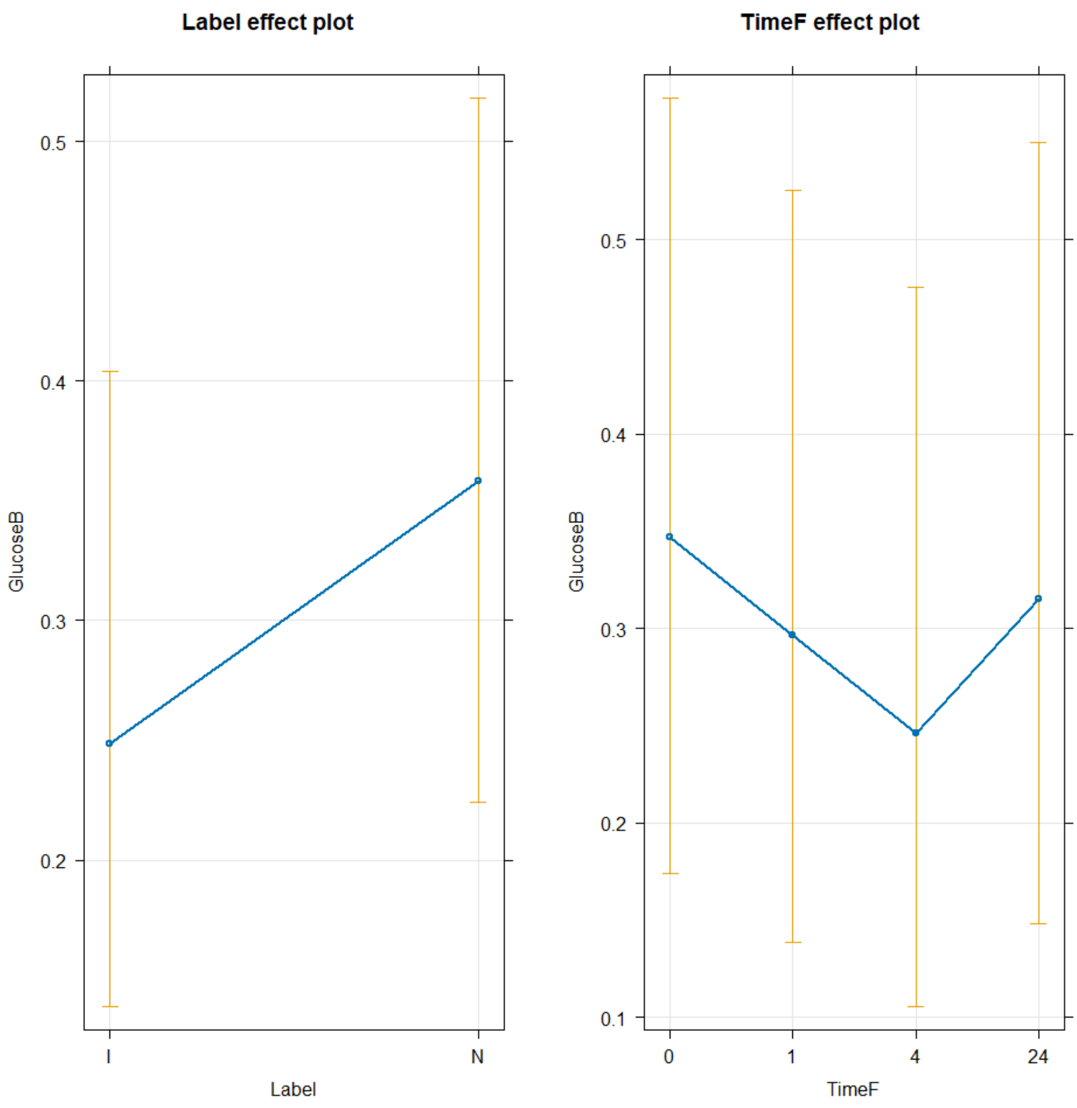


```
TukeyRes2 <- emmeans(glm3, pairwise ~ Label+TimeF, adjust = "tukey")
TukeyRes2
```

```
## $emmeans
##  Label TimeF emmean    SE  df asymp.LCL asymp.UCL
##  I     0     -0.891 0.543 Inf     -1.96    0.1732
##  N     0     -0.367 0.525 Inf     -1.40    0.6613
##  I     1     -1.123 0.563 Inf     -2.23   -0.0198
##  N     1     -0.599 0.539 Inf     -1.66    0.4579
##  I     4     -1.378 0.591 Inf     -2.54   -0.2199
##  N     4     -0.854 0.563 Inf     -1.96    0.2489
##  I     24    -1.034 0.561 Inf     -2.13    0.0649
##  N     24    -0.510 0.551 Inf     -1.59    0.5700
##
## Results are given on the logit (not the response) scale.
## Confidence level used: 0.95
##
```

```
## $contrasts
##  contrast              estimate    SE  df z.ratio p.value
##  I TimeF0 - N TimeF0    -0.5239 0.497 Inf  -1.055  0.9657
##  I TimeF0 - I TimeF1     0.2317 0.682 Inf   0.340  1.0000
##  I TimeF0 - N TimeF1    -0.2922 0.840 Inf  -0.348  1.0000
##  I TimeF0 - I TimeF4     0.4864 0.702 Inf   0.692  0.9972
##  I TimeF0 - N TimeF4    -0.0375 0.853 Inf  -0.044  1.0000
##  I TimeF0 - I TimeF24    0.1426 0.686 Inf   0.208  1.0000
##  I TimeF0 - N TimeF24   -0.3813 0.852 Inf  -0.448  0.9998
##  N TimeF0 - I TimeF1     0.7556 0.847 Inf   0.892  0.9869
##  N TimeF0 - N TimeF1     0.2317 0.682 Inf   0.340  1.0000
##  N TimeF0 - I TimeF4     1.0103 0.868 Inf   1.164  0.9419
##  N TimeF0 - N TimeF4     0.4864 0.702 Inf   0.692  0.9972
##  N TimeF0 - I TimeF24    0.6665 0.842 Inf   0.792  0.9936
##  N TimeF0 - N TimeF24    0.1426 0.686 Inf   0.208  1.0000
##  I TimeF1 - N TimeF1    -0.5239 0.497 Inf  -1.055  0.9657
##  I TimeF1 - I TimeF4     0.2547 0.715 Inf   0.356  1.0000
##  I TimeF1 - N TimeF4    -0.2691 0.867 Inf  -0.310  1.0000
##  I TimeF1 - I TimeF24   -0.0891 0.699 Inf  -0.127  1.0000
##  I TimeF1 - N TimeF24   -0.6130 0.867 Inf  -0.707  0.9968
##  N TimeF1 - I TimeF4     0.7786 0.874 Inf   0.890  0.9870
##  N TimeF1 - N TimeF4     0.2547 0.715 Inf   0.356  1.0000
##  N TimeF1 - I TimeF24    0.4348 0.849 Inf   0.512  0.9996
##  N TimeF1 - N TimeF24   -0.0891 0.699 Inf  -0.127  1.0000
##  I TimeF4 - N TimeF4    -0.5239 0.497 Inf  -1.055  0.9657
##  I TimeF4 - I TimeF24   -0.3438 0.720 Inf  -0.478  0.9998
##  I TimeF4 - N TimeF24   -0.8677 0.887 Inf  -0.979  0.9775
##  N TimeF4 - I TimeF24    0.1800 0.862 Inf   0.209  1.0000
##  N TimeF4 - N TimeF24   -0.3438 0.720 Inf  -0.478  0.9998
##  I TimeF24 - N TimeF24  -0.5239 0.497 Inf  -1.055  0.9657
##
## Results are given on the log odds ratio (not the response) scale.
## P value adjustment: tukey method for comparing a family of 8 estimates
```

```r
confint(TukeyRes2)
```

```
## $emmeans
##  Label TimeF emmean    SE  df asymp.LCL asymp.UCL
##  I     0     -0.891 0.543 Inf     -1.96    0.1732
##  N     0     -0.367 0.525 Inf     -1.40    0.6613
##  I     1     -1.123 0.563 Inf     -2.23   -0.0198
##  N     1     -0.599 0.539 Inf     -1.66    0.4579
##  I     4     -1.378 0.591 Inf     -2.54   -0.2199
##  N     4     -0.854 0.563 Inf     -1.96    0.2489
##  I     24    -1.034 0.561 Inf     -2.13    0.0649
##  N     24    -0.510 0.551 Inf     -1.59    0.5700
##
## Results are given on the logit (not the response) scale.
## Confidence level used: 0.95
##
```

```
## $contrasts
##  contrast              estimate    SE  df asymp.LCL asymp.UCL
##  I TimeF0 - N TimeF0    -0.5239 0.497 Inf     -2.03     0.981
##  I TimeF0 - I TimeF1     0.2317 0.682 Inf     -1.83     2.298
##  I TimeF0 - N TimeF1    -0.2922 0.840 Inf     -2.84     2.252
##  I TimeF0 - I TimeF4     0.4864 0.702 Inf     -1.64     2.615
##  I TimeF0 - N TimeF4    -0.0375 0.853 Inf     -2.62     2.547
##  I TimeF0 - I TimeF24    0.1426 0.686 Inf     -1.94     2.222
##  I TimeF0 - N TimeF24   -0.3813 0.852 Inf     -2.96     2.201
##  N TimeF0 - I TimeF1     0.7556 0.847 Inf     -1.81     3.323
##  N TimeF0 - N TimeF1     0.2317 0.682 Inf     -1.83     2.298
##  N TimeF0 - I TimeF4     1.0103 0.868 Inf     -1.62     3.640
##  N TimeF0 - N TimeF4     0.4864 0.702 Inf     -1.64     2.615
##  N TimeF0 - I TimeF24    0.6665 0.842 Inf     -1.88     3.217
##  N TimeF0 - N TimeF24    0.1426 0.686 Inf     -1.94     2.222
##  I TimeF1 - N TimeF1    -0.5239 0.497 Inf     -2.03     0.981
##  I TimeF1 - I TimeF4     0.2547 0.715 Inf     -1.91     2.423
##  I TimeF1 - N TimeF4    -0.2691 0.867 Inf     -2.90     2.359
##  I TimeF1 - I TimeF24   -0.0891 0.699 Inf     -2.21     2.030
##  I TimeF1 - N TimeF24   -0.6130 0.867 Inf     -3.24     2.013
##  N TimeF1 - I TimeF4     0.7786 0.874 Inf     -1.87     3.429
##  N TimeF1 - N TimeF4     0.2547 0.715 Inf     -1.91     2.423
##  N TimeF1 - I TimeF24    0.4348 0.849 Inf     -2.14     3.007
##  N TimeF1 - N TimeF24   -0.0891 0.699 Inf     -2.21     2.030
##  I TimeF4 - N TimeF4    -0.5239 0.497 Inf     -2.03     0.981
##  I TimeF4 - I TimeF24   -0.3438 0.720 Inf     -2.52     1.837
##  I TimeF4 - N TimeF24   -0.8677 0.887 Inf     -3.55     1.819
##  N TimeF4 - I TimeF24    0.1800 0.862 Inf     -2.43     2.792
##  N TimeF4 - N TimeF24   -0.3438 0.720 Inf     -2.52     1.837
##  I TimeF24 - N TimeF24  -0.5239 0.497 Inf     -2.03     0.981
##
## Results are given on the log odds ratio (not the response) scale.
## Confidence level used: 0.95
## Conf-level adjustment: tukey method for comparing a family of 8 estimates
```

```r
plot(TukeyRes2, comparison = T) + coord_flip()
```

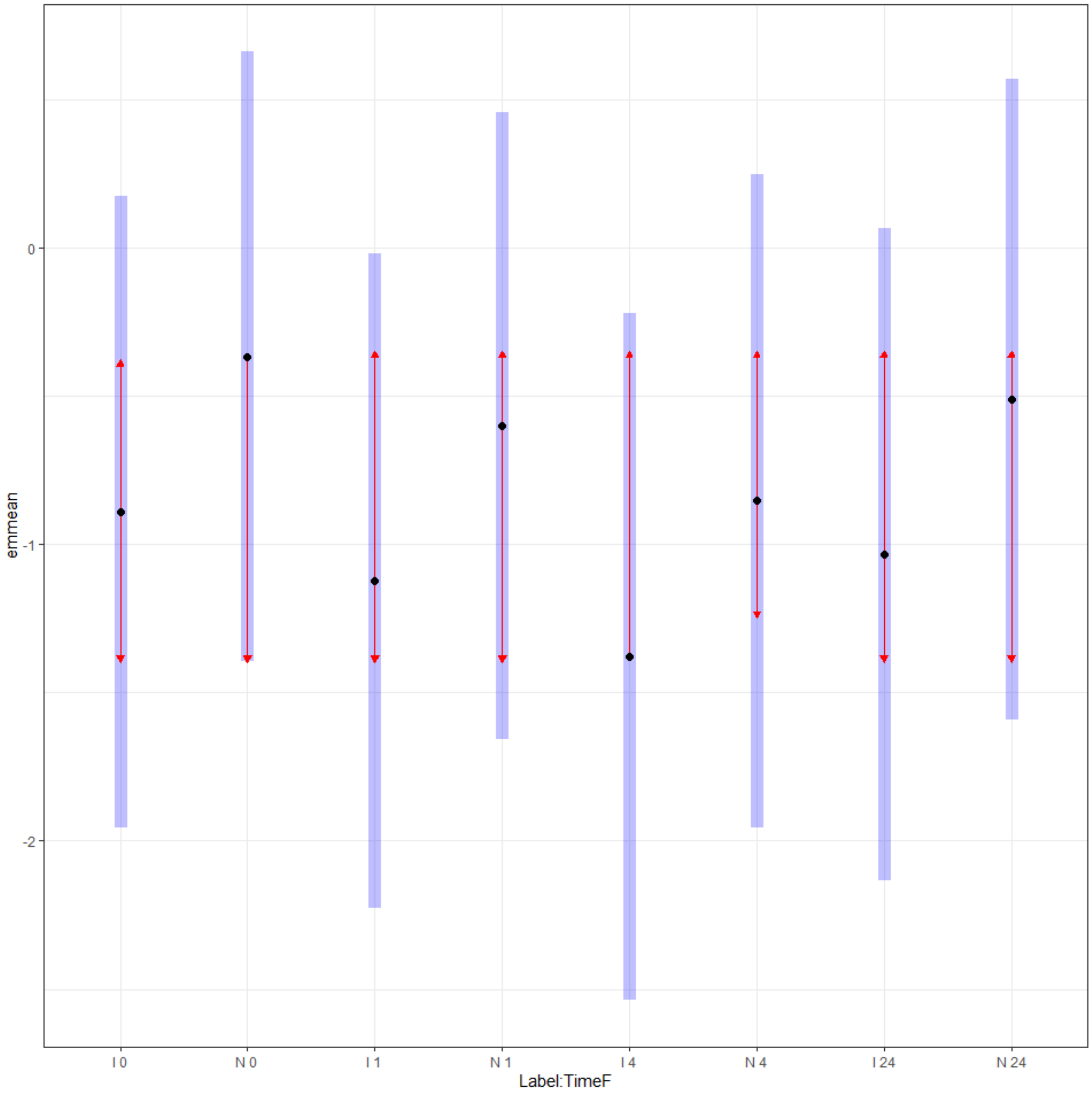


```
multcomp::cld(TukeyRes2, Letters = LETTERS)

##  Label TimeF emmean    SE  df asymp.LCL asymp.UCL .group
##  I     4     -1.378 0.591 Inf     -2.54   -0.2199  A
##  I     1     -1.123 0.563 Inf     -2.23   -0.0198  A
##  I     24    -1.034 0.561 Inf     -2.13    0.0649  A
##  I     0     -0.891 0.543 Inf     -1.96    0.1732  A
##  N     4     -0.854 0.563 Inf     -1.96    0.2489  A
##  N     1     -0.599 0.539 Inf     -1.66    0.4579  A
##  N     24    -0.510 0.551 Inf     -1.59    0.5700  A
##  N     0     -0.367 0.525 Inf     -1.40    0.6613  A
##
## Results are given on the logit (not the response) scale.
## Confidence level used: 0.95
## Results are given on the log odds ratio (not the response) scale.
## P value adjustment: tukey method for comparing a family of 8 estimates
## significance level used: alpha = 0.05
```

```
## NOTE: If two or more means share the same grouping symbol,
##       then we cannot show them to be different.
##       But we also did not show them to be the same.
```

Lactic Acid

```r
TICsetOAR <- TICsetOA %>% dplyr::filter(Label != "C") %>%
  mutate(LacticAcidB = factor(LacticAcidB))

glm3 <- glm(LacticAcidB ~ Label+TimeF, data=TICsetOAR, family = "binomial")

#Can't add donor - causes separation here
Anova(glm3)
```

```
## Analysis of Deviance Table (Type II tests)
##
## Response: LacticAcidB
##       LR Chisq Df Pr(>Chisq)
## Label   1.6891  1     0.1937
## TimeF   5.0800  3     0.1660
```

```r
summary(glm3)
```

```
##
## Call:
## glm(formula = LacticAcidB ~ Label + TimeF, family = "binomial",
##     data = TICsetOAR)
##
## Coefficients:
##             Estimate Std. Error z value Pr(>|z|)
## (Intercept)   0.3288     0.5198   0.633   0.5270
## LabelN        0.6084     0.4715   1.290   0.1969
## TimeF1       -1.2660     0.6714  -1.886   0.0593
## TimeF4       -1.0478     0.6624  -1.582   0.1137
## TimeF24      -0.2910     0.6672  -0.436   0.6628
##
## (Dispersion parameter for binomial family taken to be 1)
##
##     Null deviance: 109.50  on 78  degrees of freedom
## Residual deviance: 102.89  on 74  degrees of freedom
## AIC: 112.89
##
## Number of Fisher Scoring iterations: 4
```

```r
plot(allEffects(glm3), type = "response", grid = T)
```

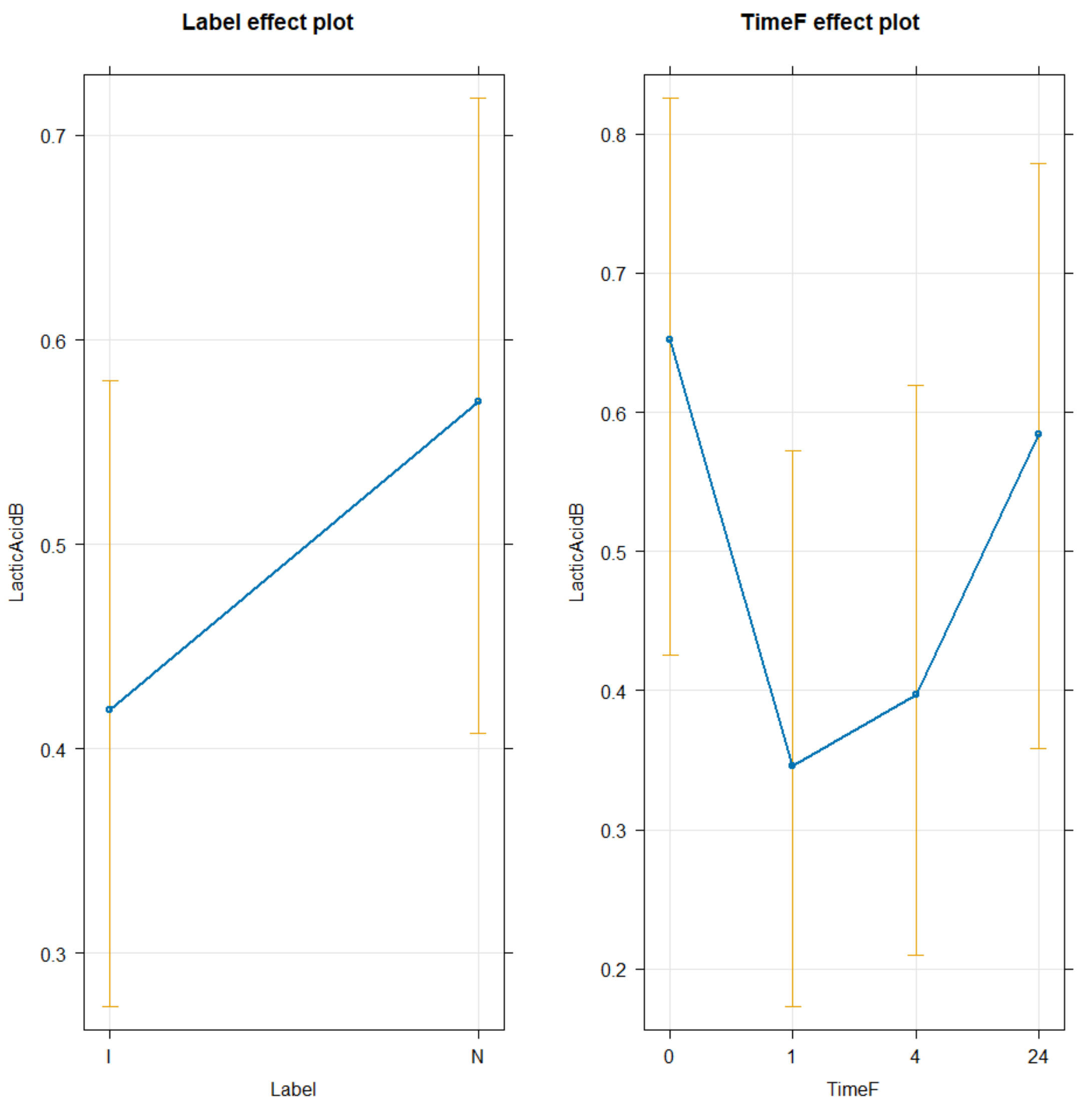


```
TukeyRes2 <- emmeans(glm3, pairwise ~ Label+TimeF, adjust = "tukey")
TukeyRes2

## $emmeans
##  Label TimeF  emmean    SE  df asymp.LCL asymp.UCL
##  I     0      0.3288 0.520 Inf    -0.690     1.348
##  N     0      0.9372 0.539 Inf    -0.120     1.994
##  I     1     -0.9372 0.539 Inf    -1.994     0.120
##  N     1     -0.3288 0.520 Inf    -1.348     0.690
##  I     4     -0.7190 0.525 Inf    -1.748     0.310
##  N     4     -0.1106 0.512 Inf    -1.114     0.892
##  I     24     0.0378 0.515 Inf    -0.972     1.048
##  N     24     0.6462 0.537 Inf    -0.406     1.699
##
## Results are given on the logit (not the response) scale.
## Confidence level used: 0.95
##
```

```
## $contrasts
##  contrast              estimate    SE  df z.ratio p.value
##  I TimeF0 - N TimeF0     -0.608 0.471 Inf  -1.290  0.9027
##  I TimeF0 - I TimeF1      1.266 0.671 Inf   1.886  0.5608
##  I TimeF0 - N TimeF1      0.658 0.795 Inf   0.827  0.9916
##  I TimeF0 - I TimeF4      1.048 0.662 Inf   1.582  0.7614
##  I TimeF0 - N TimeF4      0.439 0.792 Inf   0.555  0.9993
##  I TimeF0 - I TimeF24     0.291 0.667 Inf   0.436  0.9999
##  I TimeF0 - N TimeF24    -0.317 0.819 Inf  -0.388  0.9999
##  N TimeF0 - I TimeF1      1.874 0.845 Inf   2.218  0.3406
##  N TimeF0 - N TimeF1      1.266 0.671 Inf   1.886  0.5608
##  N TimeF0 - I TimeF4      1.656 0.834 Inf   1.986  0.4911
##  N TimeF0 - N TimeF4      1.048 0.662 Inf   1.582  0.7614
##  N TimeF0 - I TimeF24     0.899 0.815 Inf   1.103  0.9564
##  N TimeF0 - N TimeF24     0.291 0.667 Inf   0.436  0.9999
##  I TimeF1 - N TimeF1     -0.608 0.471 Inf  -1.290  0.9027
##  I TimeF1 - I TimeF4     -0.218 0.661 Inf  -0.330  1.0000
##  I TimeF1 - N TimeF4     -0.827 0.816 Inf  -1.013  0.9727
##  I TimeF1 - I TimeF24    -0.975 0.669 Inf  -1.458  0.8298
##  I TimeF1 - N TimeF24    -1.583 0.844 Inf  -1.875  0.5682
##  N TimeF1 - I TimeF4      0.390 0.808 Inf   0.483  0.9997
##  N TimeF1 - N TimeF4     -0.218 0.661 Inf  -0.330  1.0000
##  N TimeF1 - I TimeF24    -0.367 0.791 Inf  -0.463  0.9998
##  N TimeF1 - N TimeF24    -0.975 0.669 Inf  -1.458  0.8298
##  I TimeF4 - N TimeF4     -0.608 0.471 Inf  -1.290  0.9027
##  I TimeF4 - I TimeF24    -0.757 0.660 Inf  -1.147  0.9463
##  I TimeF4 - N TimeF24    -1.365 0.833 Inf  -1.638  0.7268
##  N TimeF4 - I TimeF24    -0.148 0.788 Inf  -0.188  1.0000
##  N TimeF4 - N TimeF24    -0.757 0.660 Inf  -1.147  0.9463
##  I TimeF24 - N TimeF24   -0.608 0.471 Inf  -1.290  0.9027
##
## Results are given on the log odds ratio (not the response) scale.
## P value adjustment: tukey method for comparing a family of 8 estimates
```

```
confint(TukeyRes2)
```

```
## $emmeans
##  Label TimeF  emmean    SE  df asymp.LCL asymp.UCL
##  I     0      0.3288 0.520 Inf    -0.690     1.348
##  N     0      0.9372 0.539 Inf    -0.120     1.994
##  I     1     -0.9372 0.539 Inf    -1.994     0.120
##  N     1     -0.3288 0.520 Inf    -1.348     0.690
##  I     4     -0.7190 0.525 Inf    -1.748     0.310
##  N     4     -0.1106 0.512 Inf    -1.114     0.892
##  I     24     0.0378 0.515 Inf    -0.972     1.048
##  N     24     0.6462 0.537 Inf    -0.406     1.699
##
## Results are given on the logit (not the response) scale.
## Confidence level used: 0.95
##
```

```
## $contrasts
##  contrast              estimate    SE  df asymp.LCL asymp.UCL
##  I TimeF0 - N TimeF0     -0.608 0.471 Inf    -2.037     0.821
##  I TimeF0 - I TimeF1      1.266 0.671 Inf    -0.769     3.301
##  I TimeF0 - N TimeF1      0.658 0.795 Inf    -1.752     3.067
##  I TimeF0 - I TimeF4      1.048 0.662 Inf    -0.960     3.055
##  I TimeF0 - N TimeF4      0.439 0.792 Inf    -1.960     2.839
##  I TimeF0 - I TimeF24     0.291 0.667 Inf    -1.731     2.313
##  I TimeF0 - N TimeF24    -0.317 0.819 Inf    -2.798     2.163
##  N TimeF0 - I TimeF1      1.874 0.845 Inf    -0.687     4.436
##  N TimeF0 - N TimeF1      1.266 0.671 Inf    -0.769     3.301
##  N TimeF0 - I TimeF4      1.656 0.834 Inf    -0.871     4.183
##  N TimeF0 - N TimeF4      1.048 0.662 Inf    -0.960     3.055
##  N TimeF0 - I TimeF24     0.899 0.815 Inf    -1.572     3.371
##  N TimeF0 - N TimeF24     0.291 0.667 Inf    -1.731     2.313
##  I TimeF1 - N TimeF1     -0.608 0.471 Inf    -2.037     0.821
##  I TimeF1 - I TimeF4     -0.218 0.661 Inf    -2.223     1.786
##  I TimeF1 - N TimeF4     -0.827 0.816 Inf    -3.301     1.648
##  I TimeF1 - I TimeF24    -0.975 0.669 Inf    -3.002     1.052
##  I TimeF1 - N TimeF24    -1.583 0.844 Inf    -4.143     0.976
##  N TimeF1 - I TimeF4      0.390 0.808 Inf    -2.059     2.839
##  N TimeF1 - N TimeF4     -0.218 0.661 Inf    -2.223     1.786
##  N TimeF1 - I TimeF24    -0.367 0.791 Inf    -2.765     2.032
##  N TimeF1 - N TimeF24    -0.975 0.669 Inf    -3.002     1.052
##  I TimeF4 - N TimeF4     -0.608 0.471 Inf    -2.037     0.821
##  I TimeF4 - I TimeF24    -0.757 0.660 Inf    -2.756     1.243
##  I TimeF4 - N TimeF24    -1.365 0.833 Inf    -3.891     1.160
##  N TimeF4 - I TimeF24    -0.148 0.788 Inf    -2.537     2.240
##  N TimeF4 - N TimeF24    -0.757 0.660 Inf    -2.756     1.243
##  I TimeF24 - N TimeF24   -0.608 0.471 Inf    -2.037     0.821
##
## Results are given on the log odds ratio (not the response) scale.
## Confidence level used: 0.95
## Conf-level adjustment: tukey method for comparing a family of 8 estimates

plot(TukeyRes2, comparison = T) + coord_flip()
```

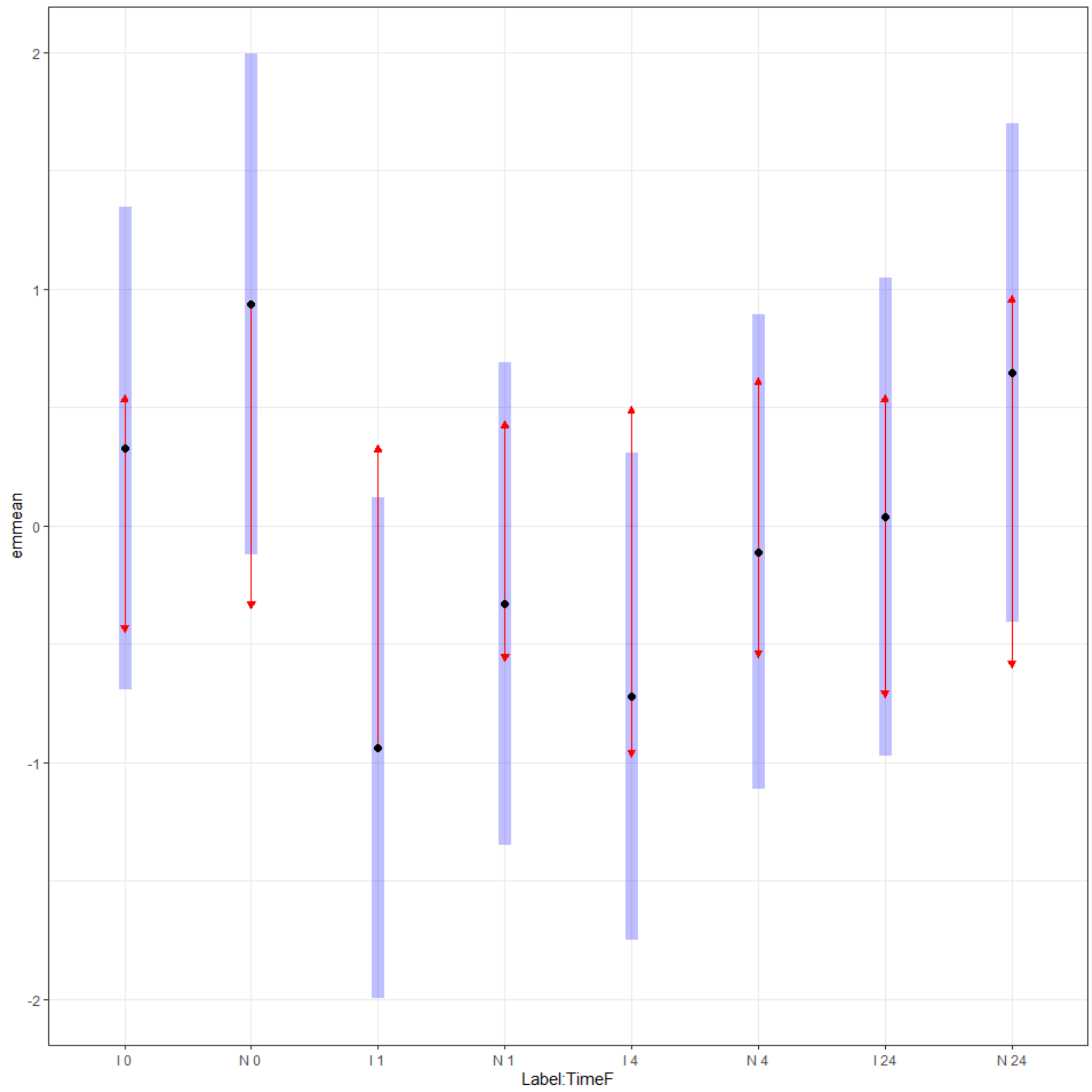


```
multcomp::cld(TukeyRes2, Letters = LETTERS)

##  Label TimeF  emmean    SE  df asymp.LCL asymp.UCL .group
##  I     1     -0.9372 0.539 Inf    -1.994     0.120  A
##  I     4     -0.7190 0.525 Inf    -1.748     0.310  A
##  N     1     -0.3288 0.520 Inf    -1.348     0.690  A
##  N     4     -0.1106 0.512 Inf    -1.114     0.892  A
##  I     24     0.0378 0.515 Inf    -0.972     1.048  A
##  I     0      0.3288 0.520 Inf    -0.690     1.348  A
##  N     24     0.6462 0.537 Inf    -0.406     1.699  A
##  N     0      0.9372 0.539 Inf    -0.120     1.994  A
##
## Results are given on the logit (not the response) scale.
## Confidence level used: 0.95
## Results are given on the log odds ratio (not the response) scale.
## P value adjustment: tukey method for comparing a family of 8 estimates
## significance level used: alpha = 0.05
```

```
## NOTE: If two or more means share the same grouping symbol,
##       then we cannot show them to be different.
##       But we also did not show them to be the same.
```

Succinic Acid

```
TICsetOAR <- TICsetOA %>% dplyr::filter(Label != "C") %>%
  mutate(SuccinicB = factor(SuccinicB))

glm3 <- glm(SuccinicB ~ Label+TimeF, data=TICsetOAR, family = "binomial")

#Can't add donor - causes separation here
Anova(glm3)
```

```
## Analysis of Deviance Table (Type II tests)
##
## Response: SuccinicB
##       LR Chisq Df Pr(>Chisq)
## Label   1.4143  1     0.2343
## TimeF   1.5380  3     0.6735
```

```
summary(glm3)
```

```
##
## Call:
## glm(formula = SuccinicB ~ Label + TimeF, family = "binomial",
##     data = TICsetOAR)
##
## Coefficients:
##               Estimate Std. Error z value Pr(>|z|)
## (Intercept)  8.207e-01  5.640e-01   1.455    0.146
## LabelN       6.010e-01  5.101e-01   1.178    0.239
## TimeF1       2.644e-16  7.364e-01   0.000    1.000
## TimeF4      -7.067e-01  6.961e-01  -1.015    0.310
## TimeF24     -5.434e-02  7.398e-01  -0.073    0.941
##
## (Dispersion parameter for binomial family taken to be 1)
##
##     Null deviance: 95.301  on 78  degrees of freedom
## Residual deviance: 92.394  on 74  degrees of freedom
## AIC: 102.39
##
## Number of Fisher Scoring iterations: 4
```

```
plot(allEffects(glm3), type = "response", grid = T)
```

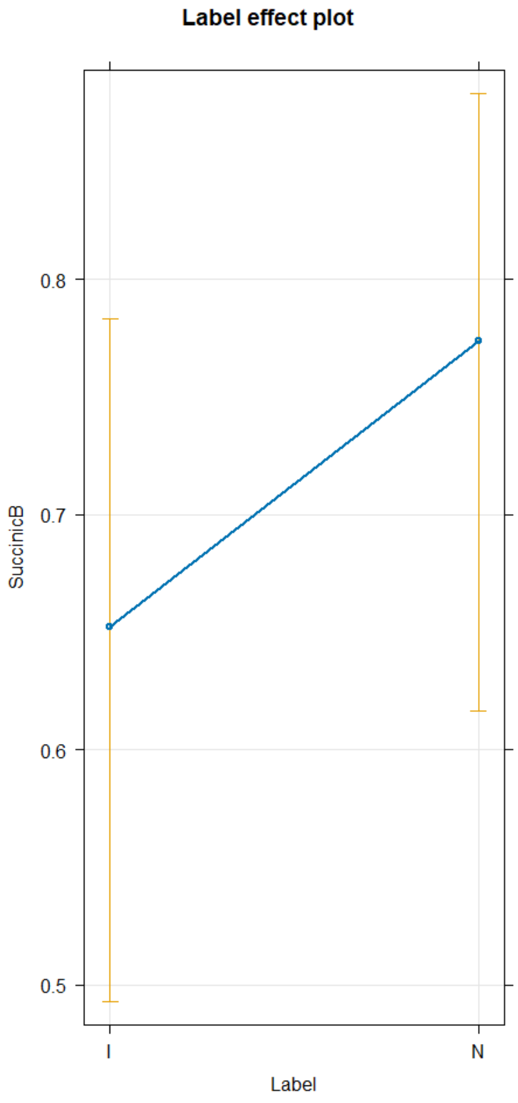


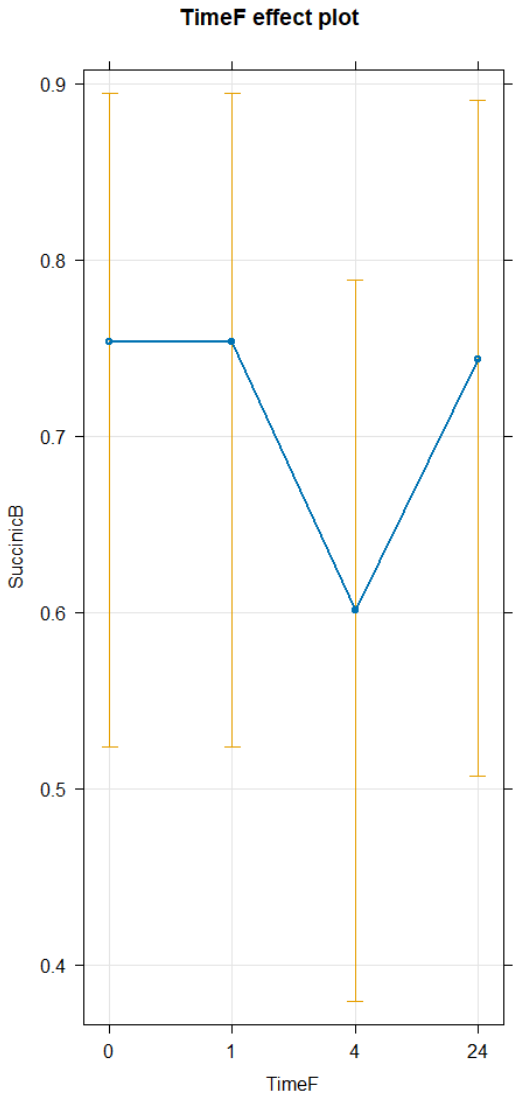


```
TukeyRes2 <- emmeans(glm3, pairwise ~ Label+TimeF, adjust = "tukey")
TukeyRes2
```

```
## $emmeans
##  Label TimeF emmean    SE  df asymp.LCL asymp.UCL
##  I     0      0.821 0.564 Inf    -0.285      1.93
##  N     0      1.422 0.598 Inf     0.250      2.59
##  I     1      0.821 0.564 Inf    -0.285      1.93
##  N     1      1.422 0.598 Inf     0.250      2.59
##  I     4      0.114 0.520 Inf    -0.905      1.13
##  N     4      0.715 0.535 Inf    -0.333      1.76
##  I     24     0.766 0.564 Inf    -0.339      1.87
##  N     24     1.367 0.608 Inf     0.177      2.56
##
## Results are given on the logit (not the response) scale.
## Confidence level used: 0.95
##
```

```
## $contrasts
##  contrast              estimate    SE  df z.ratio p.value
##  I TimeF0 - N TimeF0    -0.6010 0.510 Inf  -1.178  0.9383
##  I TimeF0 - I TimeF1     0.0000 0.736 Inf   0.000  1.0000
##  I TimeF0 - N TimeF1    -0.6010 0.896 Inf  -0.671  0.9977
##  I TimeF0 - I TimeF4     0.7067 0.696 Inf   1.015  0.9723
##  I TimeF0 - N TimeF4     0.1057 0.849 Inf   0.124  1.0000
##  I TimeF0 - I TimeF24    0.0543 0.740 Inf   0.073  1.0000
##  I TimeF0 - N TimeF24   -0.5467 0.905 Inf  -0.604  0.9988
##  N TimeF0 - I TimeF1     0.6010 0.896 Inf   0.671  0.9977
##  N TimeF0 - N TimeF1     0.0000 0.736 Inf   0.000  1.0000
##  N TimeF0 - I TimeF4     1.3077 0.876 Inf   1.492  0.8121
##  N TimeF0 - N TimeF4     0.7067 0.696 Inf   1.015  0.9723
##  N TimeF0 - I TimeF24    0.6554 0.892 Inf   0.735  0.9960
##  N TimeF0 - N TimeF24    0.0543 0.740 Inf   0.073  1.0000
##  I TimeF1 - N TimeF1    -0.6010 0.510 Inf  -1.178  0.9383
##  I TimeF1 - I TimeF4     0.7067 0.696 Inf   1.015  0.9723
##  I TimeF1 - N TimeF4     0.1057 0.849 Inf   0.124  1.0000
##  I TimeF1 - I TimeF24    0.0543 0.740 Inf   0.073  1.0000
##  I TimeF1 - N TimeF24   -0.5467 0.905 Inf  -0.604  0.9988
##  N TimeF1 - I TimeF4     1.3077 0.876 Inf   1.492  0.8121
##  N TimeF1 - N TimeF4     0.7067 0.696 Inf   1.015  0.9723
##  N TimeF1 - I TimeF24    0.6554 0.892 Inf   0.735  0.9960
##  N TimeF1 - N TimeF24    0.0543 0.740 Inf   0.073  1.0000
##  I TimeF4 - N TimeF4    -0.6010 0.510 Inf  -1.178  0.9383
##  I TimeF4 - I TimeF24   -0.6523 0.700 Inf  -0.932  0.9830
##  I TimeF4 - N TimeF24   -1.2533 0.886 Inf  -1.414  0.8511
##  N TimeF4 - I TimeF24   -0.0513 0.846 Inf  -0.061  1.0000
##  N TimeF4 - N TimeF24   -0.6523 0.700 Inf  -0.932  0.9830
##  I TimeF24 - N TimeF24  -0.6010 0.510 Inf  -1.178  0.9383
##
## Results are given on the log odds ratio (not the response) scale.
## P value adjustment: tukey method for comparing a family of 8 estimates
```

```
confint(TukeyRes2)
```

```
## $emmeans
##  Label TimeF emmean    SE  df asymp.LCL asymp.UCL
##  I     0      0.821 0.564 Inf    -0.285      1.93
##  N     0      1.422 0.598 Inf     0.250      2.59
##  I     1      0.821 0.564 Inf    -0.285      1.93
##  N     1      1.422 0.598 Inf     0.250      2.59
##  I     4      0.114 0.520 Inf    -0.905      1.13
##  N     4      0.715 0.535 Inf    -0.333      1.76
##  I     24     0.766 0.564 Inf    -0.339      1.87
##  N     24     1.367 0.608 Inf     0.177      2.56
##
## Results are given on the logit (not the response) scale.
## Confidence level used: 0.95
##
```

```
## $contrasts
##  contrast              estimate    SE  df asymp.LCL asymp.UCL
##  I TimeF0 - N TimeF0    -0.6010 0.510 Inf     -2.15     0.945
##  I TimeF0 - I TimeF1     0.0000 0.736 Inf     -2.23     2.232
##  I TimeF0 - N TimeF1    -0.6010 0.896 Inf     -3.32     2.114
##  I TimeF0 - I TimeF4     0.7067 0.696 Inf     -1.40     2.816
##  I TimeF0 - N TimeF4     0.1057 0.849 Inf     -2.47     2.680
##  I TimeF0 - I TimeF24    0.0543 0.740 Inf     -2.19     2.297
##  I TimeF0 - N TimeF24   -0.5467 0.905 Inf     -3.29     2.197
##  N TimeF0 - I TimeF1     0.6010 0.896 Inf     -2.11     3.316
##  N TimeF0 - N TimeF1     0.0000 0.736 Inf     -2.23     2.232
##  N TimeF0 - I TimeF4     1.3077 0.876 Inf     -1.35     3.964
##  N TimeF0 - N TimeF4     0.7067 0.696 Inf     -1.40     2.816
##  N TimeF0 - I TimeF24    0.6554 0.892 Inf     -2.05     3.359
##  N TimeF0 - N TimeF24    0.0543 0.740 Inf     -2.19     2.297
##  I TimeF1 - N TimeF1    -0.6010 0.510 Inf     -2.15     0.945
##  I TimeF1 - I TimeF4     0.7067 0.696 Inf     -1.40     2.816
##  I TimeF1 - N TimeF4     0.1057 0.849 Inf     -2.47     2.680
##  I TimeF1 - I TimeF24    0.0543 0.740 Inf     -2.19     2.297
##  I TimeF1 - N TimeF24   -0.5467 0.905 Inf     -3.29     2.197
##  N TimeF1 - I TimeF4     1.3077 0.876 Inf     -1.35     3.964
##  N TimeF1 - N TimeF4     0.7067 0.696 Inf     -1.40     2.816
##  N TimeF1 - I TimeF24    0.6554 0.892 Inf     -2.05     3.359
##  N TimeF1 - N TimeF24    0.0543 0.740 Inf     -2.19     2.297
##  I TimeF4 - N TimeF4    -0.6010 0.510 Inf     -2.15     0.945
##  I TimeF4 - I TimeF24   -0.6523 0.700 Inf     -2.77     1.470
##  I TimeF4 - N TimeF24   -1.2533 0.886 Inf     -3.94     1.433
##  N TimeF4 - I TimeF24   -0.0513 0.846 Inf     -2.61     2.512
##  N TimeF4 - N TimeF24   -0.6523 0.700 Inf     -2.77     1.470
##  I TimeF24 - N TimeF24  -0.6010 0.510 Inf     -2.15     0.945
##
## Results are given on the log odds ratio (not the response) scale.
## Confidence level used: 0.95
## Conf-level adjustment: tukey method for comparing a family of 8 estimates
```

```r
plot(TukeyRes2, comparison = T) + coord_flip()
```

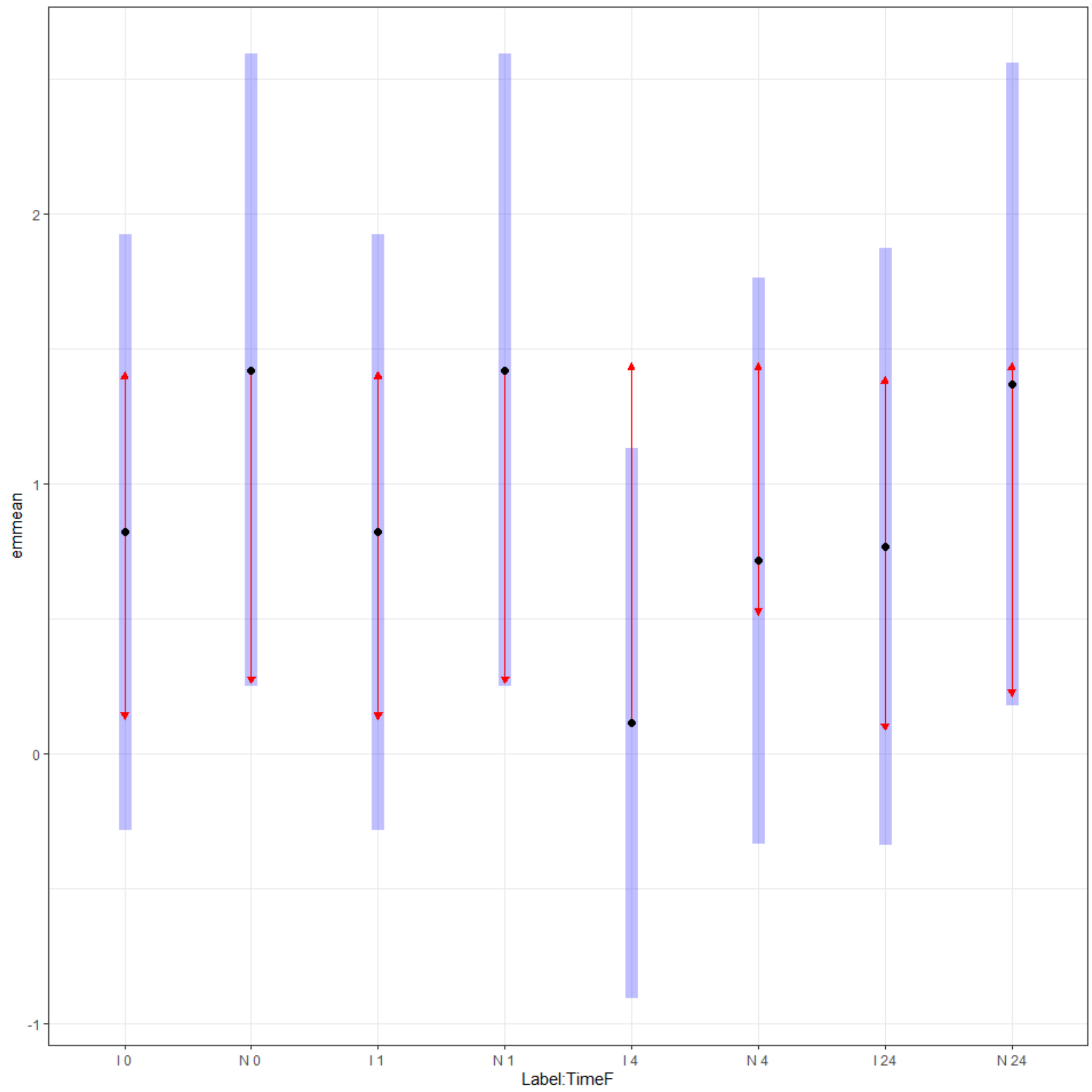


```
multcomp::cld(TukeyRes2, Letters = LETTERS)

##  Label TimeF emmean    SE  df asymp.LCL asymp.UCL .group
##  I     4      0.114 0.520 Inf    -0.905      1.13  A
##  N     4      0.715 0.535 Inf    -0.333      1.76  A
##  I     24     0.766 0.564 Inf    -0.339      1.87  A
##  I     0      0.821 0.564 Inf    -0.285      1.93  A
##  I     1      0.821 0.564 Inf    -0.285      1.93  A
##  N     24     1.367 0.608 Inf     0.177      2.56  A
##  N     0      1.422 0.598 Inf     0.250      2.59  A
##  N     1      1.422 0.598 Inf     0.250      2.59  A
##
## Results are given on the logit (not the response) scale.
## Confidence level used: 0.95
## Results are given on the log odds ratio (not the response) scale.
## P value adjustment: tukey method for comparing a family of 8 estimates
## significance level used: alpha = 0.05
```

```
## NOTE: If two or more means share the same grouping symbol,
##       then we cannot show them to be different.
##       But we also did not show them to be the same.
```

```
Frucdone<- TICsetOAR %>%
  ggplot() +
  geom_mosaic(aes(x=product(TimedLoading),fill=Fructose6B), offset=0.02) +
labs(y="Fructose-6-Phosphate", x= "Loading and Time")

Frucdone
```

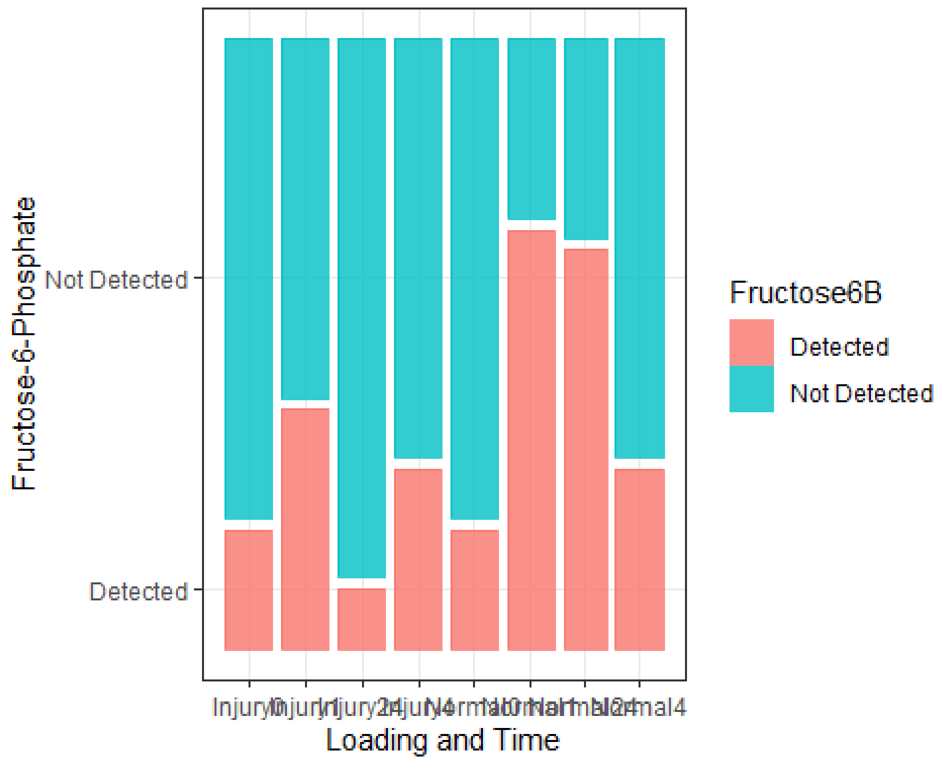


```
pirateplot(formula=Glutamineplot~Loading+Time, data=TICsetOAR, theme=3,
inf.b.o = 0,inf.f.o = 0, ylab="TIC Corrected Glutamine*10e9", main="Glutamine
Concentration")
```

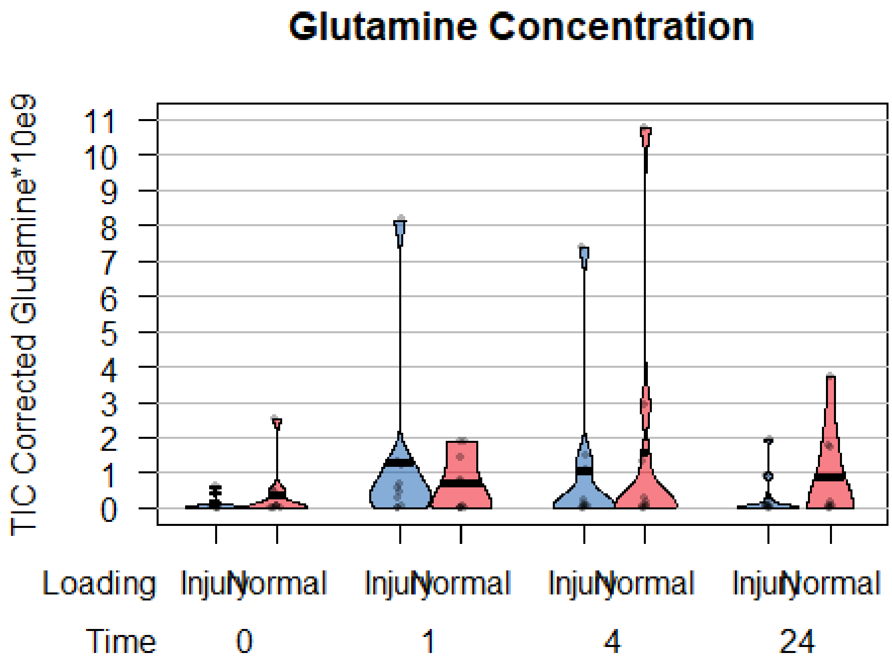

Glutamine Concentration
TIC Corrected Glutamine*10e9
11
10
9
8
7
6
5
4
3
2
1
0
Loading
Injury
Normal
Injury
Normal
Injury
Normal
Injury
Normal
Time
0
1
4
24
